# A Comprehensive Mathematical Model Simulating the Adaptive Immune Response

**DOI:** 10.1101/2025.10.14.682330

**Authors:** Zhaobin Xu, Dongqing Wei

## Abstract

Adaptive immunity plays a crucial role in defending against invading pathogenic microorganisms. It encompasses both humoral and cellular immune responses. For the first time, we have systematically developed a mathematical model of adaptive immunity that comprehensively incorporates the effects of both humoral and cellular immunity. Our model successfully explains several key immunological phenomena, including the formation of immune memory (involving both B-cell and T-cell memory), the mechanism of original antigenic sin, the basis of secondary infections, and the processes of activation and exhaustion in cellular immunity. Furthermore, we employed a discrete time-scale agent-based modeling approach to simulate the dynamics of the adaptive immune response following pathogen invasion, with a specific focus on elucidating the mechanisms underlying chronic infection. Finally, we have established a novel mathematical model of the interaction between cancer cells and the immune system, providing a more robust theoretical foundation for cancer immunotherapy.

## Introduction

The increase in human life expectancy is closely related to the development of immunology, and the surge in the world’s population shows a very strong positive correlation with vaccination ^[1–2]^.

In the 20th century, biology has made significant progress in immunology, particularly in the study of adaptive immunity. Scientists have revealed the mechanisms underlying the generation of BCR diversity, which is primarily responsible for antibody production, and TCR diversity, which is generated by helper T cells (CD4+ T cells) and cytotoxic T cells (CD8+ T cells) ^[3–5]^. Flow cytometry and single-cell sequencing technologies allow us to obtain massive amounts of immune cell sequence information ^[6–8]^. Spatial omics sequencing technologies allow us to capture immune cell complexes in different tissue structures and spatial locations, including the cell composition and gene expression characteristics of germinal centers ^[9–10]^. Although experimental methods have made significant progress, relying solely on them cannot provide a macroscopic assessment of complex systems, especially for dynamic processes such as viral infection. Furthermore, deep learning methods based on big data are far from sufficient to decode complex immune processes. Therefore, it is necessary to establish a universal model that comprehensively considers both cellular and humoral immunity. Furthermore, the mathematical model must be supported by rigorous physical mechanisms; otherwise, it will fall into the trap of overfitting. Establishing a reliable mathematical model can accurately reflect the immune system’s stress response to foreign microbial invasion and reveal the causes of many immunological phenomena from a dynamic perspective, including the occurrence of secondary infection ^[11–12]^, the maintenance of memory cells ^[13–14]^, the exhaustion of CD8+ T cells ^[15–16],^ and the mechanisms of long-term chronic infection ^[17–18]^.

However, current research on adaptive immunity lacks this universal, macroscopic mathematical model. Previous mathematical models fail to account for the polymorphism of B and T cells, nor for the remodeling of the host antibody population following viral infection. Therefore, we aim to develop a large-scale mathematical model that explicitly accounts for antibody and TCR polymorphism, thereby more accurately simulating the adaptive immune response.

We have developed a novel mathematical model based on biological mechanisms. This model considers BCR and antibody diversity and comprehensively integrates both cellular and humoral immunity. Our model not only simulates the dynamics of adaptive immunity during pathogen infection but also simulates the dynamics of adaptive immunity during cancer development. Ultimately, we hope to use this mathematical model to explain immunological phenomena and guide vaccine development, treatment of chronic infections, and immunotherapy strategies for cancer.

## Results

### 2.1 Modeling of Simple Humoral Immunity in Adaptive Immunity (Only B cells are considered, not CD4 + T cells)

A diagram of humoral immunity principle is illustrated in Figure 1A

**Figure 1A:**
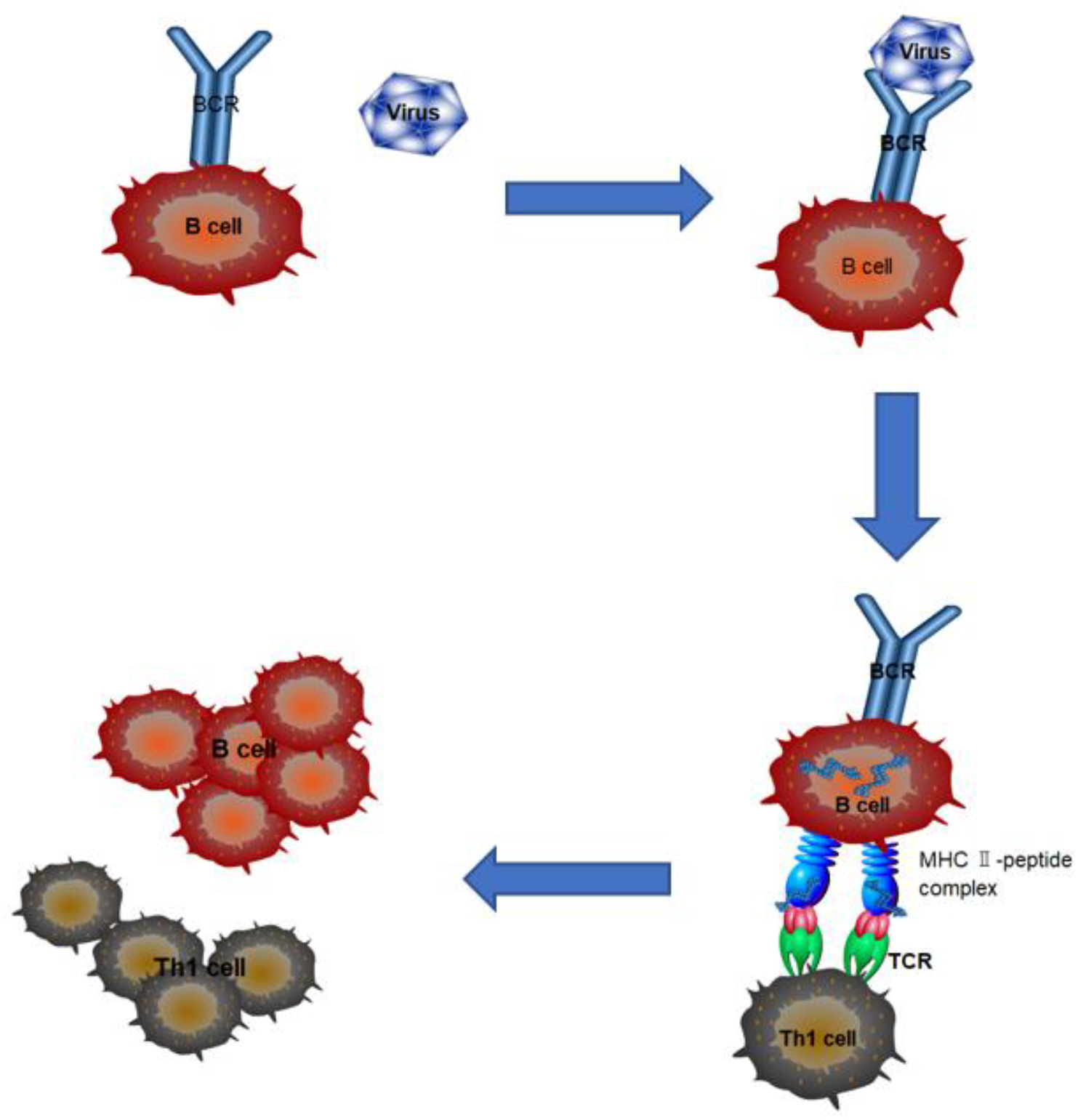
Diagram of humoral immune response

The basic framework of our humoral immunity model is shown in Figure 1A. When the virus binds to the BCR on the surface of the B cell, the antigen-antibody complex will be further processed by the B cell, and the antigen will be degraded into peptides and presented to the CD4+ T cell on the surface of the MHC-II complex. This antigen presentation complex further stimulates the CD4 + T cell to secrete cytokines to promote the division and proliferation of both, thereby achieving the following function: B cell with a stronger affinity BCR can present more antigen to the CD4 + T cell, thereby achieving a faster proliferation effect. Although the traditional view is that dendritic cell (DC) is the main antigen-presenting cell, recent studies have shown that in humoral immunity, B cells play a dominant role in antigen presentation ^[19–21]^, and studies have shown that the proliferation ability of B cells is significantly positively correlated with the affinity of their BCR and antigen ^[22–24]^. Our model is based on physical background. It explicitly expresses the ability of the antigen-antibody complex to stimulate antibody regeneration, while considering the kinetic characteristics of the antigen-BCR binding process. Therefore, it can more accurately reflect the kinetic process of antibody-virus interaction in humoral immunity.

Mathematical analysis is provided in the supplementary materials. To further verify the correctness of this model, we have expanded on this basic model. We use three cases to demonstrate the correctness of this model. These three cases are the induced proliferation process of specific antibodies, the viral rebound phenomenon in monoclonal antibody treatment, and the viral rebound case in small molecule viral inhibitor treatment. In recent years, the expansion of antibodies has been studied and simulated in more detail experimentally ^[25–26]^, but there is currently no good kinetic model that can explain the process of the immune system selecting and cloning high-affinity antibodies. We used model 3.1.3 to simulate the kinetic changes of BCR (or antibodies) with different kinetic parameters during the virus interaction process. The results are shown in Figure 1B. The red solid line in Figure 1B represents the kinetic change characteristics of the virus. It can be seen that the virus is effectively eliminated as the antibody proliferates. The four antibodies have the same initial concentration level. Antibodies 1, 3, and 4 have the same binding energy, but the positive binding coefficient of antibody 1 is greater than that of antibody 3, and the positive binding coefficient of antibody 3 is greater than that of antibody 4. Antibody 1 has a lower binding energy than antibody 2, but their positive binding coefficients are the same, and the negative dissociation coefficient of antibody 2 is greater than that of antibody 1. The specific parameters and initial values are provided in the method section and supplementary materials. As can be seen from the figure, antibodies with strong binding ability can achieve greater proliferation. For antibodies with the same binding energy, antibodies with large positive binding coefficients can achieve greater proliferation. This means that when the immune system faces external antigen stimulation, it not only significantly proliferates antibodies with good binding affinity, but also significantly proliferates antibodies that bind quickly.

The second case is to explain the phenomenon of viral rebound in some patients during monoclonal antibody treatment. This phenomenon has been observed in many clinical trials ^[27]^. This viral rebound affects the efficacy of monoclonal antibody treatment. Establishing a correct mathematical model can not only explain the humoral immune mechanism, but also provide theoretical guidance for drug dosage. Alan Perelson et al. believe that this viral rebound is caused by the generation of mutant strains ^[28]^. We put forward a different view here. We use model 3.1.4 to explain this phenomenon. Like autologous BCR and antibodies, exogenous monoclonal antibodies also have strong virus binding ability, so the binding kinetics with viruses are no different from autologous BCR. However, unlike autologous BCR, the complex formed by the binding of exogenous antibodies and viruses cannot be captured by the corresponding B cells and ultimately presented to CD4 +T cells, so it cannot further stimulate its own proliferation. Therefore, exogenous monoclonal antibodies are a one-way consumption process. In addition to binding to viruses, they also degrade in the body. Therefore, when the injection dose just reaches a certain range, it can quickly bind to the virus in the short term, causing the viral concentration to drop rapidly after its injection. However, when it is exhausted, the viral concentration will rise again, and the virus will eventually be eliminated after the massive proliferation of autoantibodies. The rebound of the virus is not an inevitable event. When monoclonal antibodies are injected in large doses or multiple injections, the virus may be completely eliminated by the combined action of exogenous antibodies and autoantibodies, so there will be no secondary rebound. Another scenario is that the patient’s own immunity is strong and can rapidly proliferate autoantibodies. After the exogenous antibodies are consumed, the autoantibodies have reached a high level, and the virus will not rebound. Otherwise, we will see the viral rebound phenomenon described in Figure 1C. In this case, we can also see that the input of exogenous antibodies will interfere with the rapid proliferation of self-antibodies.

The third case is to explain the phenomenon of viral rebound in some patients treated with small molecule viral inhibitors. This phenomenon has also been observed in clinical trials ^[29]^, which affects the therapeutic effect of small molecule inhibitors. Alan Perelson et al. believed that viral rebound is caused by incomplete depletion of susceptible cells ^[30]^. We use model 3.1.5 to explain this phenomenon. We believe that the reason why some patients experience a rebound in viral count after drug withdrawal is the insufficient antibody level. Drug-virus complex formed after the small molecule inhibitor binds to the virus temporarily inhibits viral replication, but the specific antibodies are not fully stimulated at this time. Therefore, after stopping the drug, viral replication can be further continued. The results are shown in Figure 1D. We administered the drug at the 20th time point. After administration, the concentration of the drug decreased significantly because the drug can bind to the virus to form a complex. At the same time, the drug itself has a degradation rate. However, the red antibodies are not fully stimulated at this time. Therefore, after stopping the drug, the viral concentration may rebound (as indicated by the blue solid line in the figure). Only some patients will experience viral rebound. This depends on the relationship between the virus, the host and the drug. Large doses or long-term continuous administration are not likely to cause viral rebound. Patients with strong humoral immune function can stimulate antibody proliferation more quickly and are not likely to cause viral rebound. Viral rebound is only a sporadic event, occurring when the dosage and the patient’s immunity are within a threshold range. Establishing a scientific mathematical model can better explain the mechanism of viral rebound and, at the same time, allow for scientific planning of dosage and time intervals for administration to prevent the occurrence of viral rebound. Although small molecule inhibitors can cause viral rebound in some individuals, small molecule inhibitors, like monoclonal antibody therapy, also inhibit the activation of individual humoral immunity. However, both methods can effectively control the peak concentration of the virus, thereby fundamentally protecting patients. We present those two case studies to substantiate the theoretical underpinnings of our modeling framework.

In order to more accurately describe the humoral immune process, we added more compartments to the simple model. The most important of these is the addition of B cell typing, which divides B cells into non-plasma cell and antibody secreting plasma. As shown in Figure 1E, non-plasma B cell B cells can be further divided into memory IgG-BCR B cells and non-memory IgM-BCR B cells. IgM-BCR B cells are continuously replenished by stem cell division, while IgG-BCR B cells do not have this pathway. We believe that both IgM and IgG are produced by plasma cells. In this model, we particularly emphasize the role of environmental antigens in maintaining the homeostasis of various B cells. Unlike viral antigens, environmental antigens are not a fixed component but a complex of all antigens. Therefore, their affinity for the BCR of different B cells is the same. In the non-infected state, IgM-BCR B cells are replenished from two sources: stem cell differentiation and the regeneration of new IgM-BCR B cells in response to environmental antigens. In the non-infected state, IgG-BCR B cells are replenished from a single source: the regeneration of new IgG-BCR B cells in response to environmental antigens. In the non-infected state, the replenishment of IgM-ASC cells is mainly through the transformation of a part of IgM-BCR B cells after being stimulated by environmental antigens; the replenishment of IgG-ASC cells is mainly through the transformation of a part of IgG-BCR B cells after being stimulated by environmental antigens.

After infection, the mutual transformation relationship of various cells is as follows: the complex formed after IgM-BCR binds to the virus is presented to CD4+ T cells through the MHC-II complex, which will eventually stimulate the proliferation of its own B cells. At the same time, some cells will differentiate into IgM-ASC cells, while some will be converted into IgG-BCR B cells.

The complex formed after IgG-BCR binds to the virus is presented to CD4+ T cells through the MHC-II complex, which will eventually stimulate the proliferation of its own B cells. At the same time, some cells will differentiate into IgG-ASC cells.

Based on model 3.1.6, we analyzed two phenomena: the impact of vaccination strategies on antibody dynamics and the antibody production disorder (germinal center expansion disorder) in severe patients. A large number of experimental statistical data show ^[31–33]^ that vaccination with a short interval often leads to the obstruction of secondary antibody elevation, and even causes a significant decrease in antibodies after the second vaccination. Increasing the vaccination interval or increasing the vaccination dose during the second vaccination can significantly play a role in secondary antibody enhancement. The main reason for this phenomenon is that the large amount of IgM and IgG antibodies produced after the first vaccination competitively interfere with the binding of IgM-BCR or IgG-BCR with antigens, and the binding of IgM or IgG with antigens can only quickly reduce the antibody content in the body fluids. After vaccination, the rate of decline of free antibodies such as IgM and IgG in the body fluids is significantly faster than the decay rate of the corresponding BCR B cells. Therefore, when the interval is long enough, this competitive interference effect will be reduced. This will further ensure that sufficient BCRs are in contact with antigens, thereby stimulating the production of more ASCs and corresponding antibodies. Increasing the second vaccination dose can also play the same role. As simulated in Figure 1F, extending the vaccinations interval and increasing the secondary vaccination dose can overcome the competitive interference of antibodies, which has been found in some other experiments ^[89–90]^. Our mathematical model successfully explains the kinetic mechanism of this interference. In addition, we used model 3.1.6 to explain another clinical phenomenon, that is, excessive antigen stimulation may cause an excessive proportion of ASCs to be produced during the early response of B cells in GC, thereby leading to the proliferation disorder of non-plasma B cells. This will eventually result in the failure of the response of the germinal center in the later stage and the occurrence of severe disease. Clinical experiments have shown ^[34–35]^ that severe COVID-19 patients often have severe germinal center dysfunction. As shown in Figure 1G, for people with strong immunity (antigen-antibody complexes stimulate antibody regeneration with a larger regeneration coefficient), highly toxic viruses (larger virus replication coefficients) can stimulate the level of specific antibodies more strongly than weakly toxic viruses. However, for people with weak immunity (antigen-antibody complexes stimulate antibody regeneration with a smaller regeneration coefficient), highly toxic viruses often lead to antibody production disorders. This is partly due to the dynamics of virus-antibody interactions (detailed analysis is provided in the supplementary materials). Furthermore, excessive antigen stimulation causes a large number of B cells in the germinal center to undergo ASC conversion, leading to a shortage of non-plasma B cells. The BCRs on the surface of non-plasma B cells sense antigen stimulation and serve as the source of further differentiation into ASCs and antibody production. When non-plasma B cells are insufficient, germinal centers atrophy, exacerbating the dysfunction of antibody production.

To further investigate the dynamics of the entire antibody repertoire during viral infection, we further expanded this model, using model 3.1.7 to simulate the effects of viral infection on the entire antibody repertoire. We assumed that the initial antibody repertoire consisted of antibodies and BCRs with different binding kinetics to the viral antigen, and that the distribution of their binding parameters followed a normal distribution. As shown in Figure 1F, during viral infection, IgM-BCR B cells with strong binding affinity rapidly proliferated, ultimately exhibiting a bimodal pattern. However, due to their rapid decay coefficient and the continuous replenishment of IgM-BCR B cells from stem cells, IgM-BCR B cells rapidly decayed after infection, ultimately returning to their original equilibrium state. As can be seen from figure 1H, IgM-BCR peaks in the high-binding capacity region rapidly decayed, corresponding to the decay of IgM-ASC cells and IgM antibodies. Therefore, IgM-BCR and IgM lack immune memory function. The carrier of immune memory is IgG-BCR B cells. As can be seen from the figure, the proliferation of IgG-BCR lags significantly behind that of IgM-BCR, and the increase in IgG also lags behind that of IgM. This lag is mainly due to two reasons: the binding affinity of IgG-BCR is less than that of IgM-BCR. On the cell membrane, IgM-BCR mostly exists in the form of polymers ^[36–37]^, which leads to its binding kinetics significantly better than IgG-BCR. This will cause IgM-BCR and IgM to be increased at a faster speed and larger amplitude. At the same time, part of IgG-BCR comes from the conversion of IgM-BCR, so its increase has a significant lag. A large number of experiments have shown ^[38–39]^ that for the first viral infection, the increase of IgM often dominates, and its elevation amplitude and speed are significantly higher than the increase amplitude and speed of IgG. However, in the secondary infection, the increase of IgG often dominates. For a long time, people have been unable to give a scientific and reasonable explanation for this. Our model can explain this phenomenon very well, and the existence of this phenomenon also verifies the correctness of our model.

Using this model, we further explain the dynamic mechanism of immune memory formation. If we extend the timeline, we can obtain Figure 1I. From Figure 1I, we can see that after a long enough time, the distribution of IgM-ASC and IgM will return to the original state, while IgG-ASC and IgG will form a small peak for a long time in the area with better binding activity. The maintenance of immune memory is a dynamic process. The specific mathematical analysis is in the supplementary materials. In the maintenance of immune memory, the presence of environmental antigenic substances is crucial. Experiments have shown that B cells without BCR will die rapidly ^[40]^, which illustrates the role of BCR in maintaining B cell proliferation. In the non-infection state, BCR can only bind to environmental antigenic substances. Therefore, environmental antigenic substances are the root in maintaining immune memory. As described above, environmental antigens exhibit nearly identical kinetic binding parameters for different antibodies (this does not mean that each individual self-antigen has identical binding kinetic parameters for different antibodies; the diversity of environmental antigens leads to a consistent overall binding kinetic). Therefore, the concentration of environmental antigens maintains the steady-state distribution of IgM and IgG in the non-infected state. Specifically, for IgG, the concentration of environmental antigens maintains a constant overall IgG level. Following infection, as specific IgG levels rise, overall IgG levels also increase. During this period, as the population approaches the new steady-state, each IgG type decreases proportionally. Therefore, even if specific IgG levels decrease, the magnitude of the decrease is equal to the overall IgG level, preventing a return to the initial state and allowing immune memory to form. The situation is different for IgM. Although IgM can also be produced through autologous proliferation, IgM-BCRs are continuously produced from stem cells, and IgM exhibits faster decay kinetics. Therefore, from a mathematical equilibrium analysis, the distribution of IgM will eventually return to its pre-infection state, and IgM does not serve as a carrier of immune memory.

**Figure 1B:**
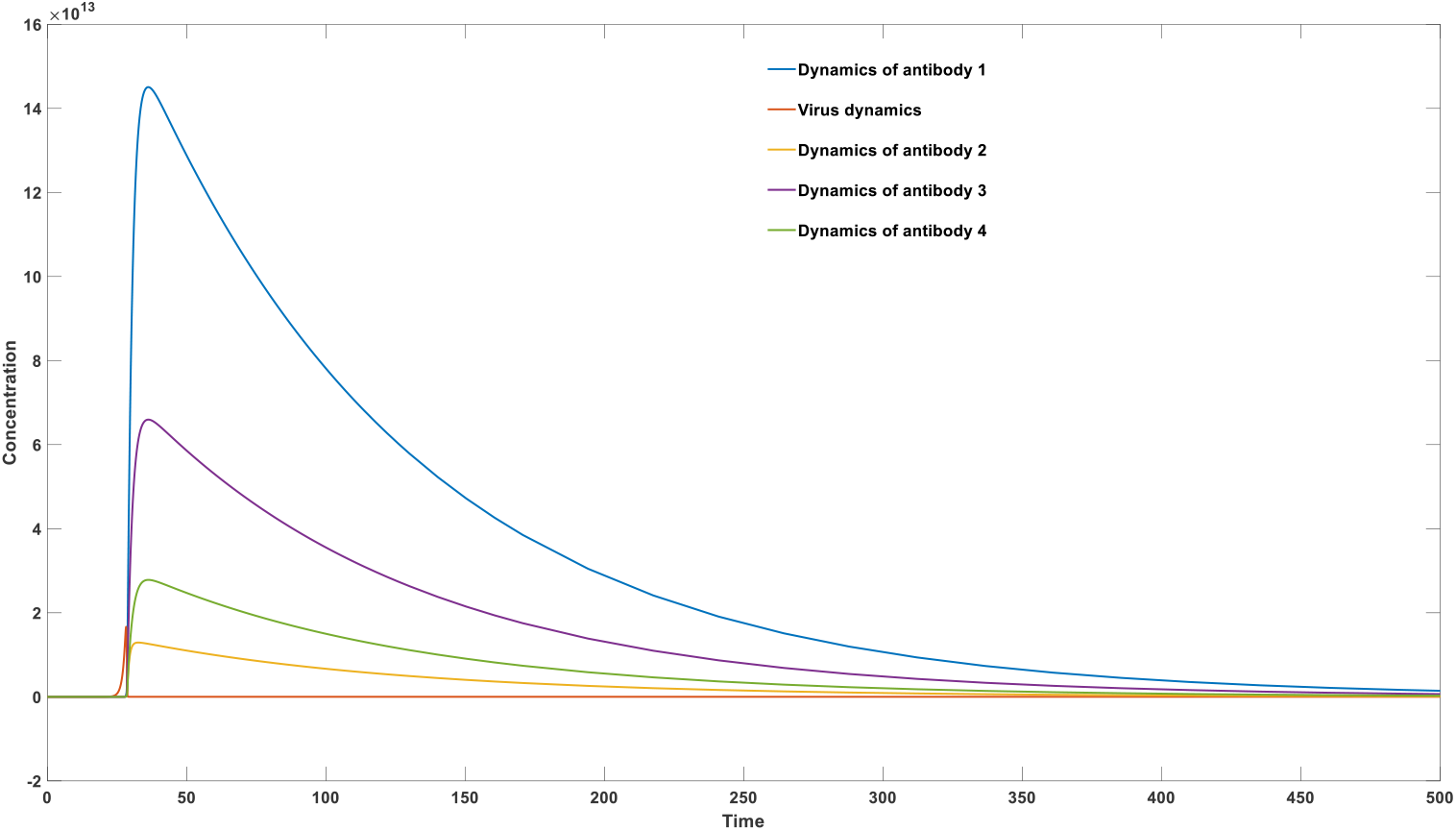
mechanism of specific antibody clonal expansion after infection

**Figure 1C:**
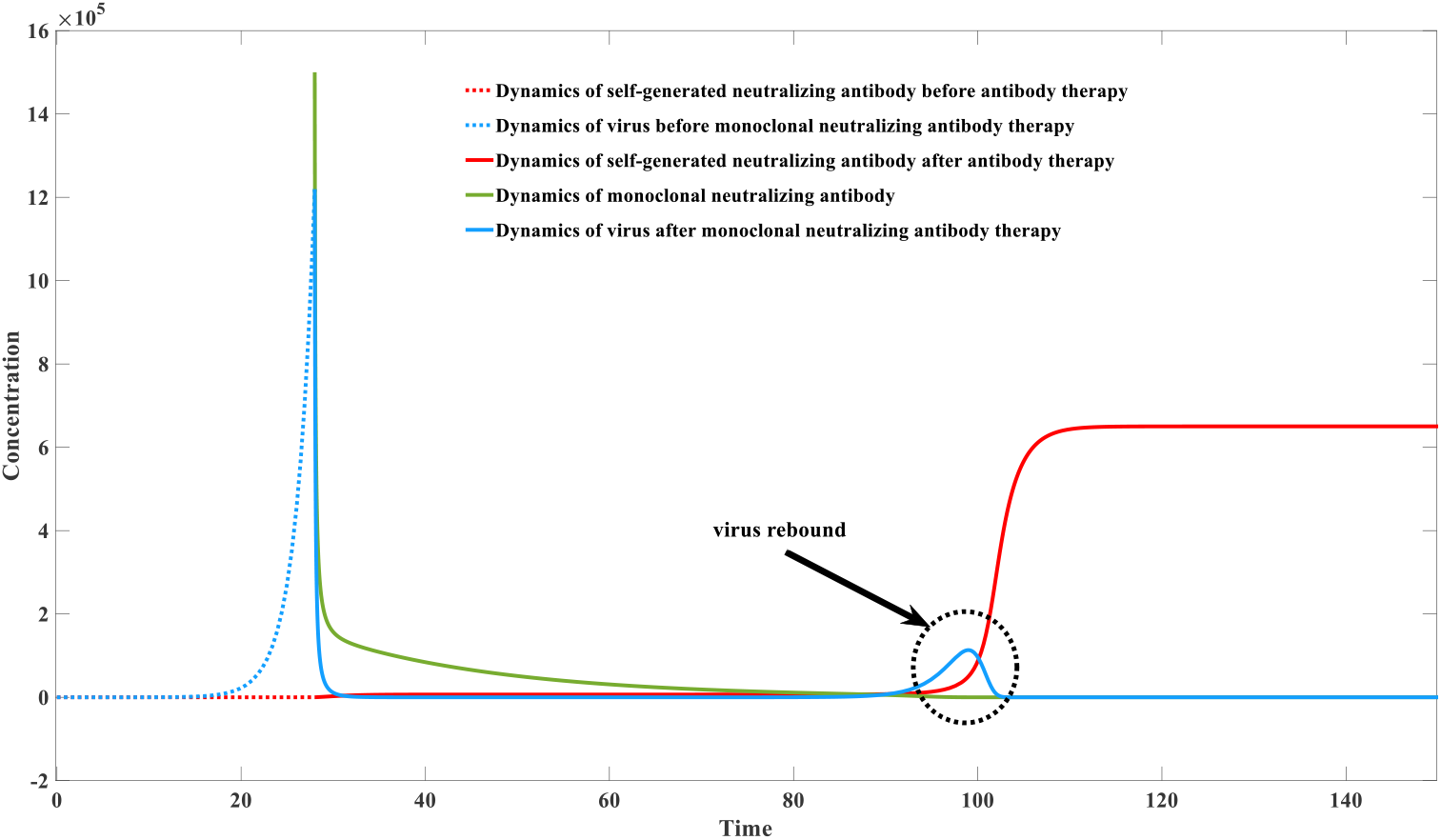
mechanism of virus rebound after monoclonal neutralizing antibody therapy

**Figure 1D:**
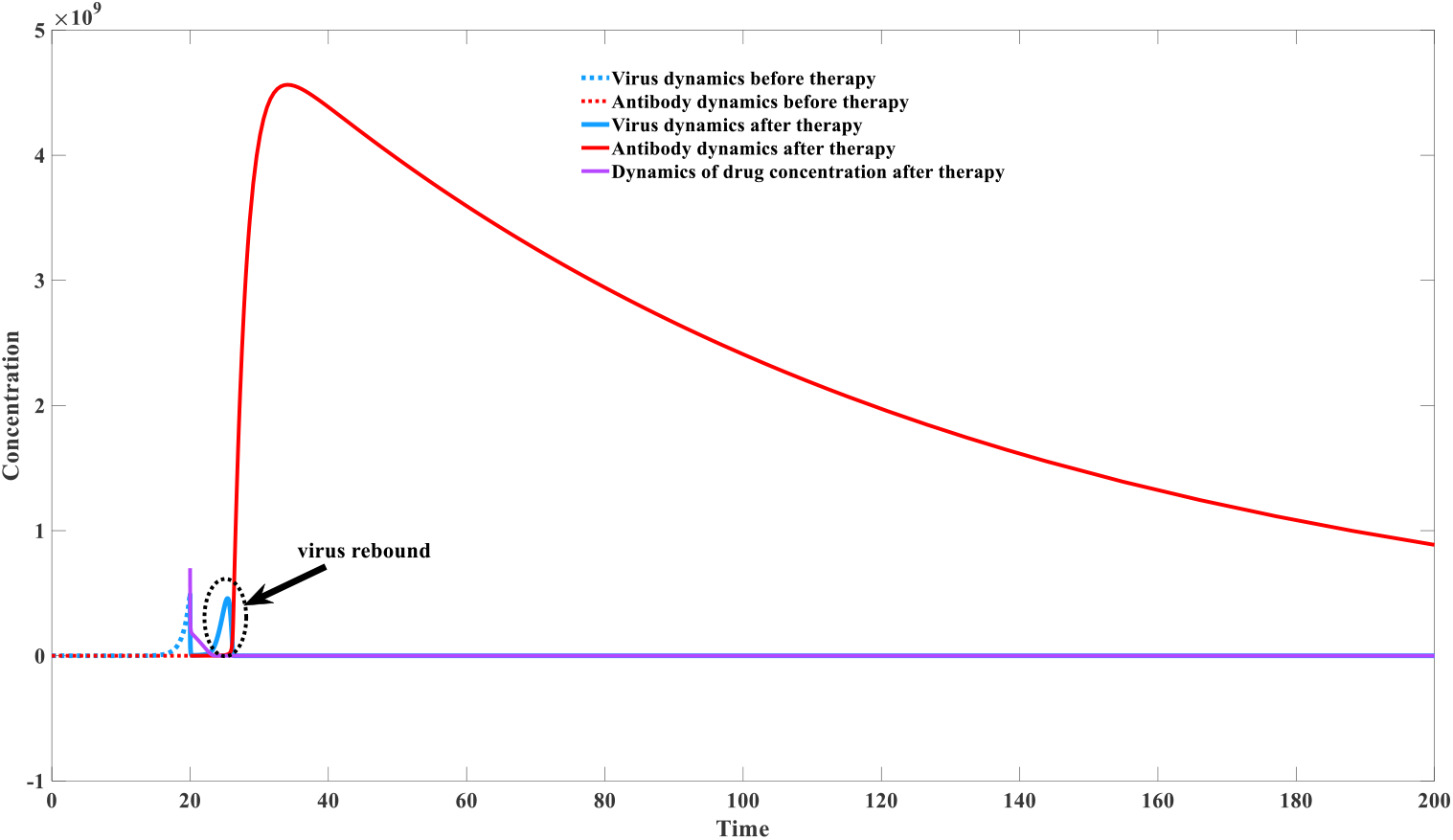
mechanism of virus rebound after nirmatrelvir-ritonavir therapy

**Figure 1E:**
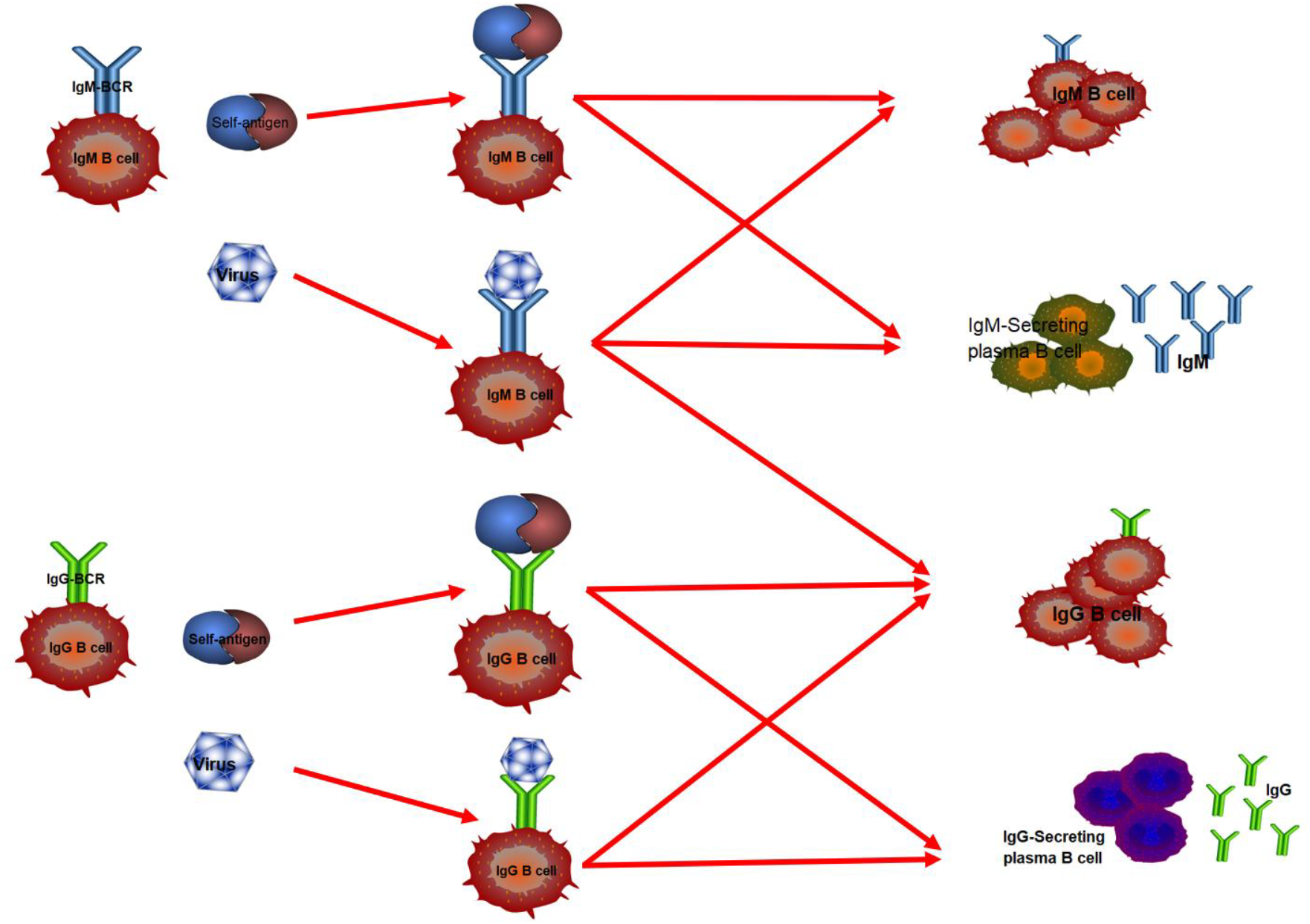
Diagram of humoral response considering antibody secreting plasma cell

**Figure 1F:**
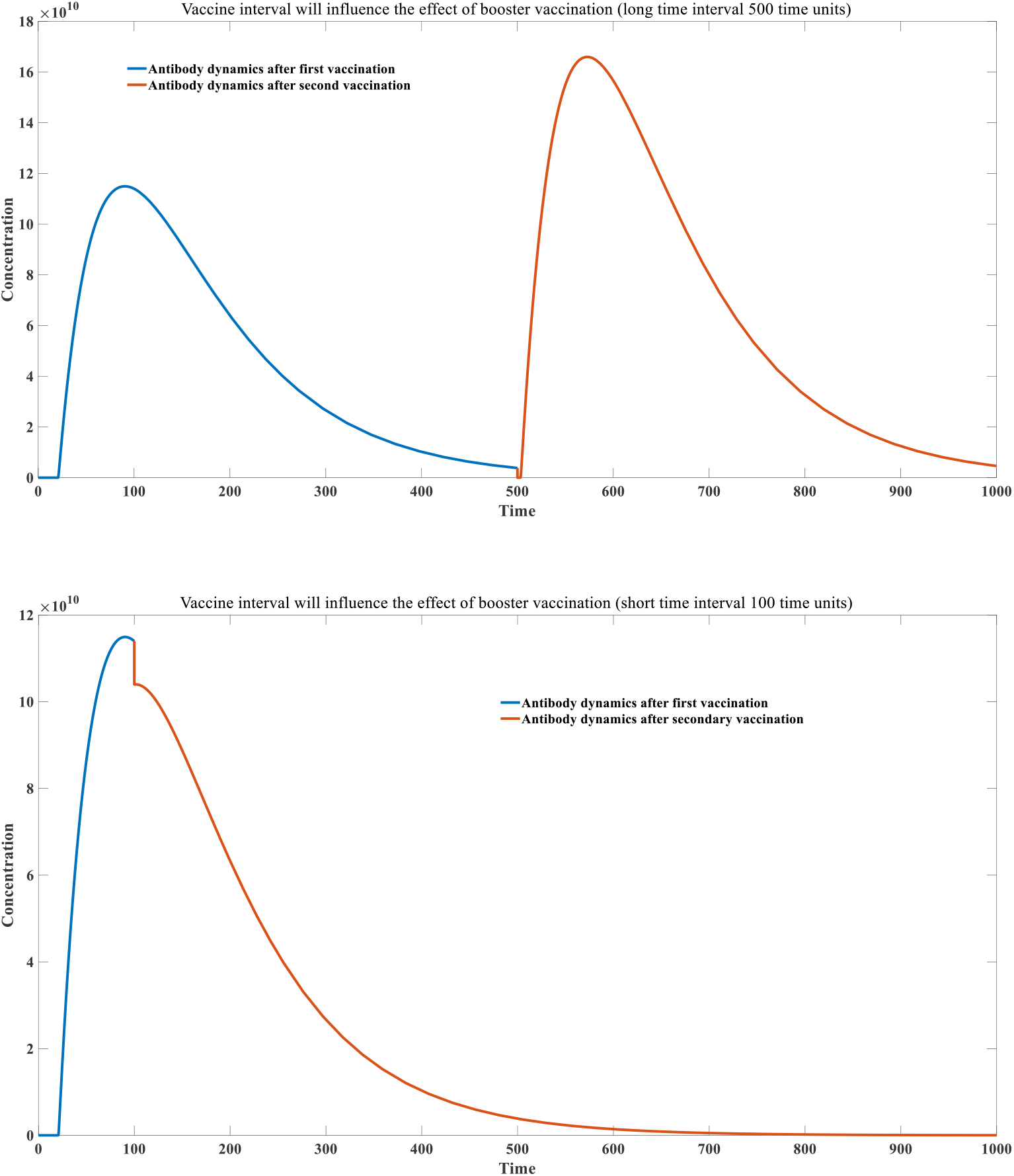
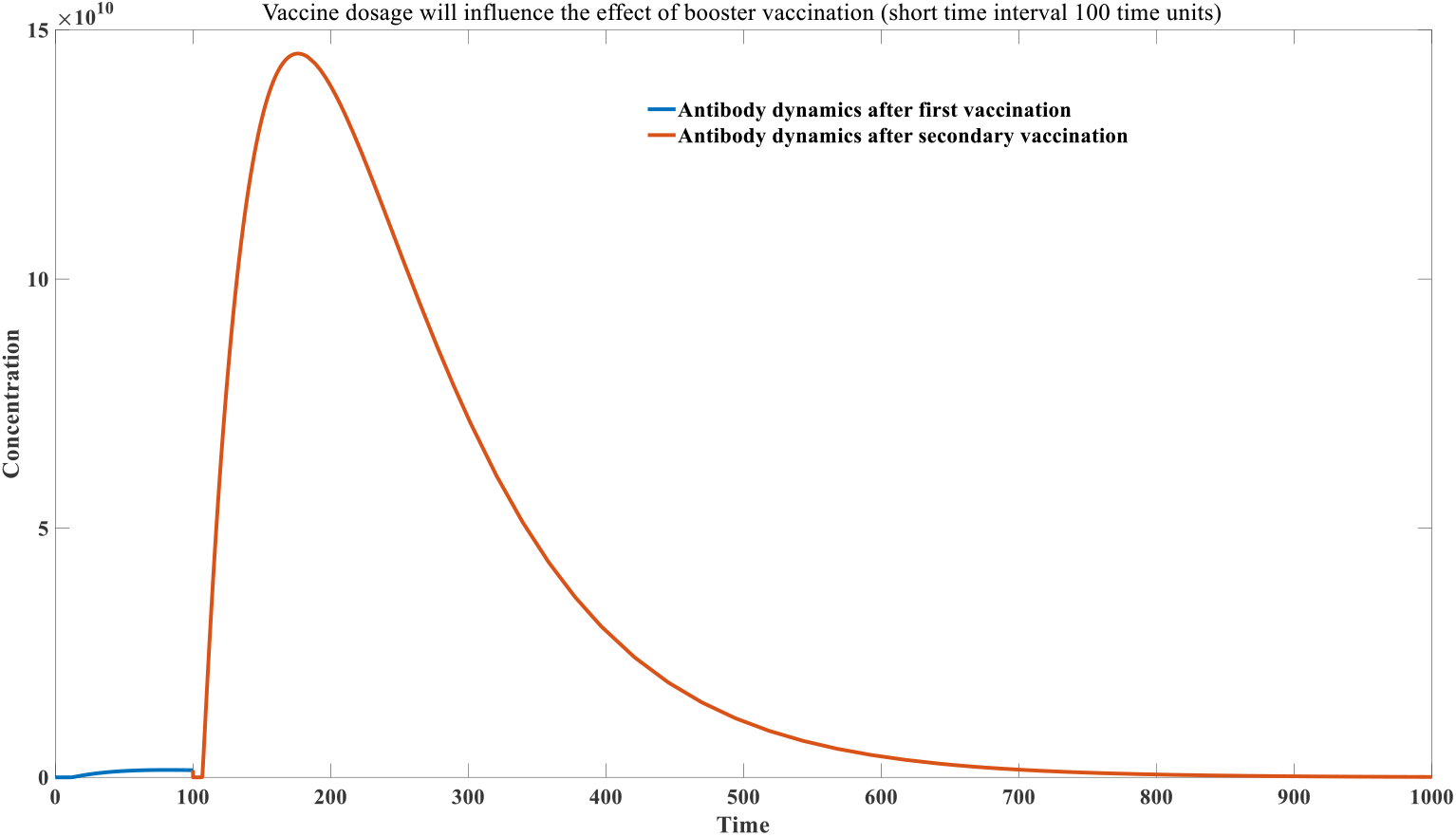
Effect of vaccine interval and dosage on the antibody dynamics

**Figure 1G:**
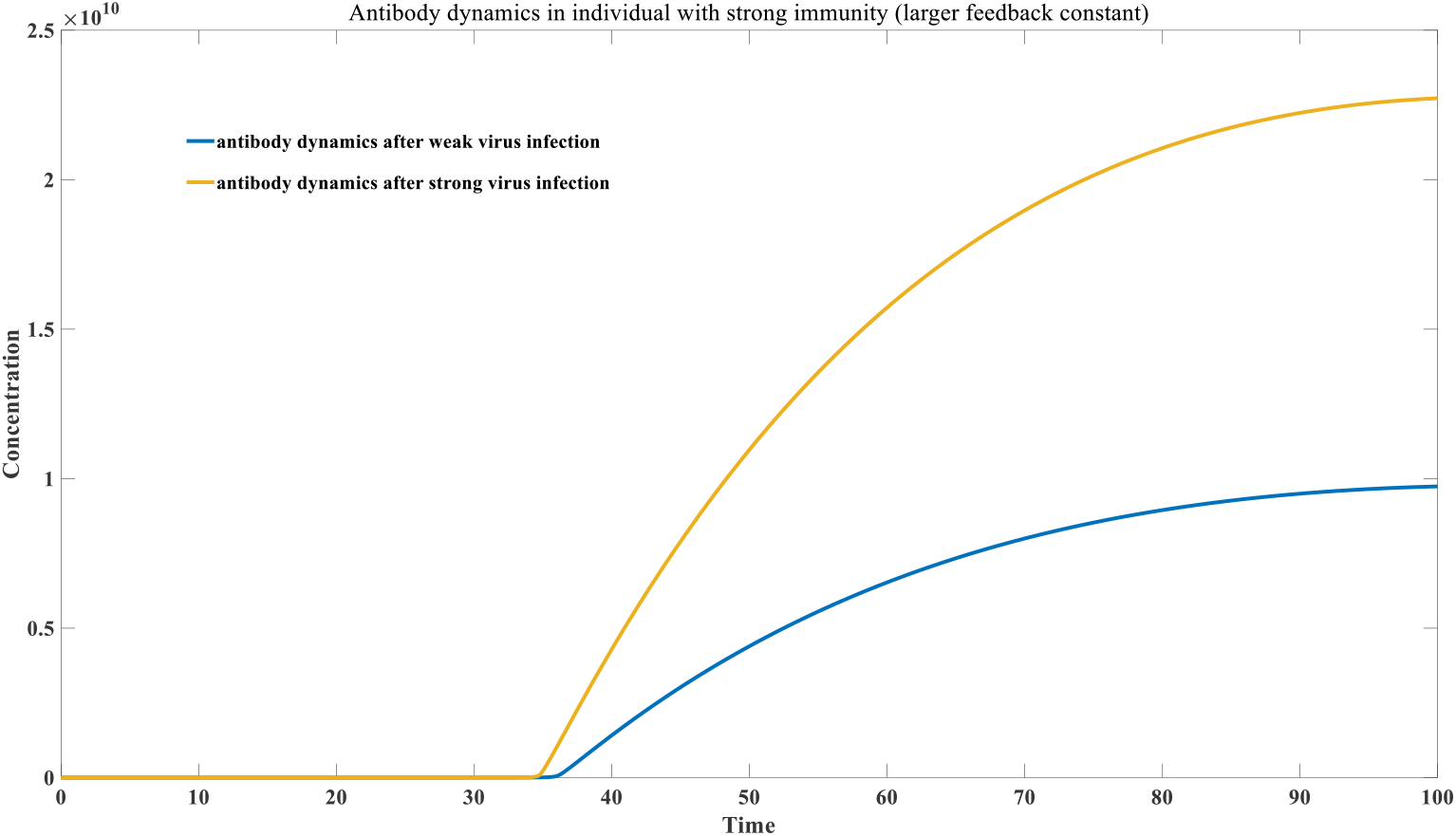
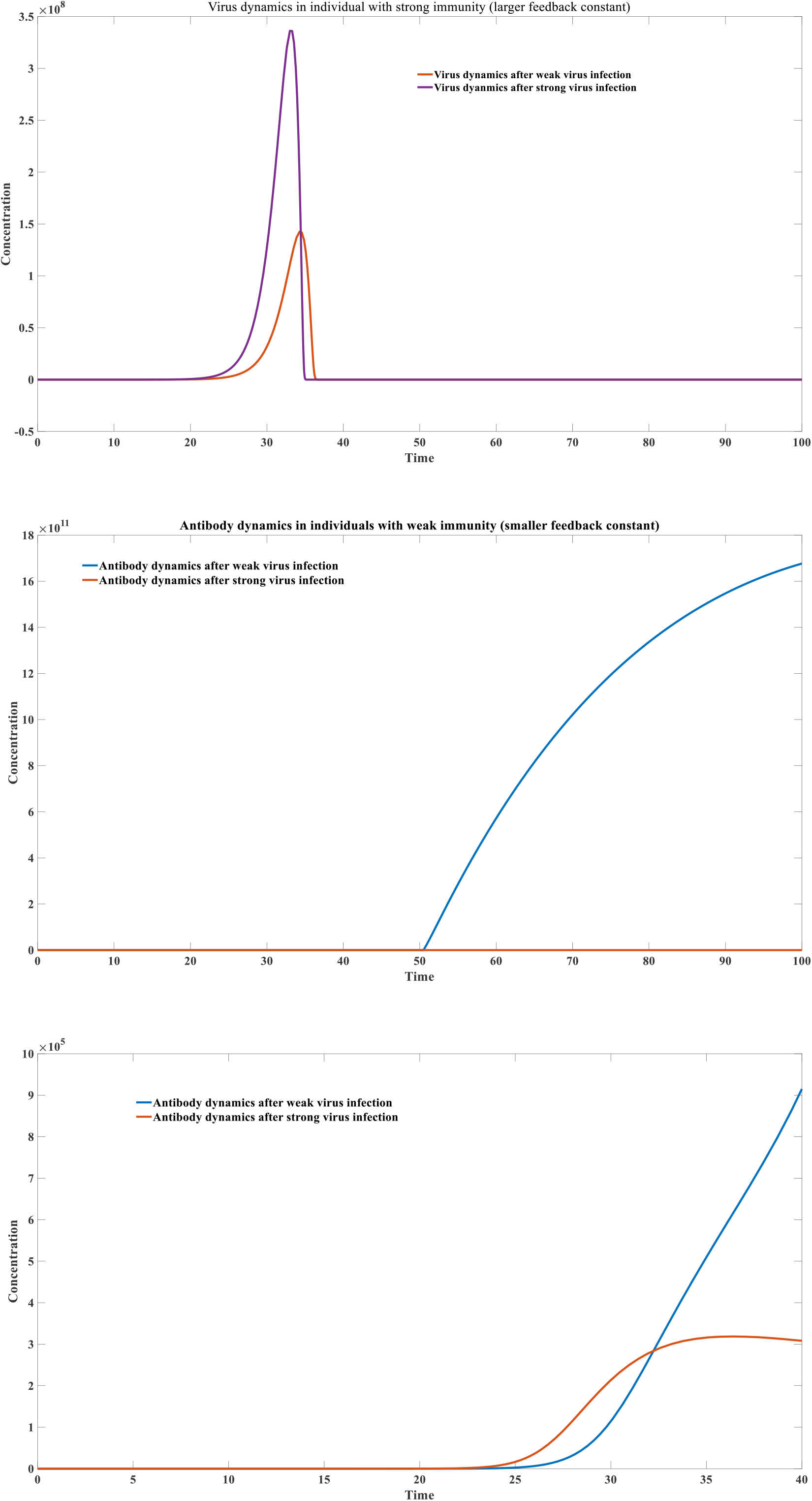
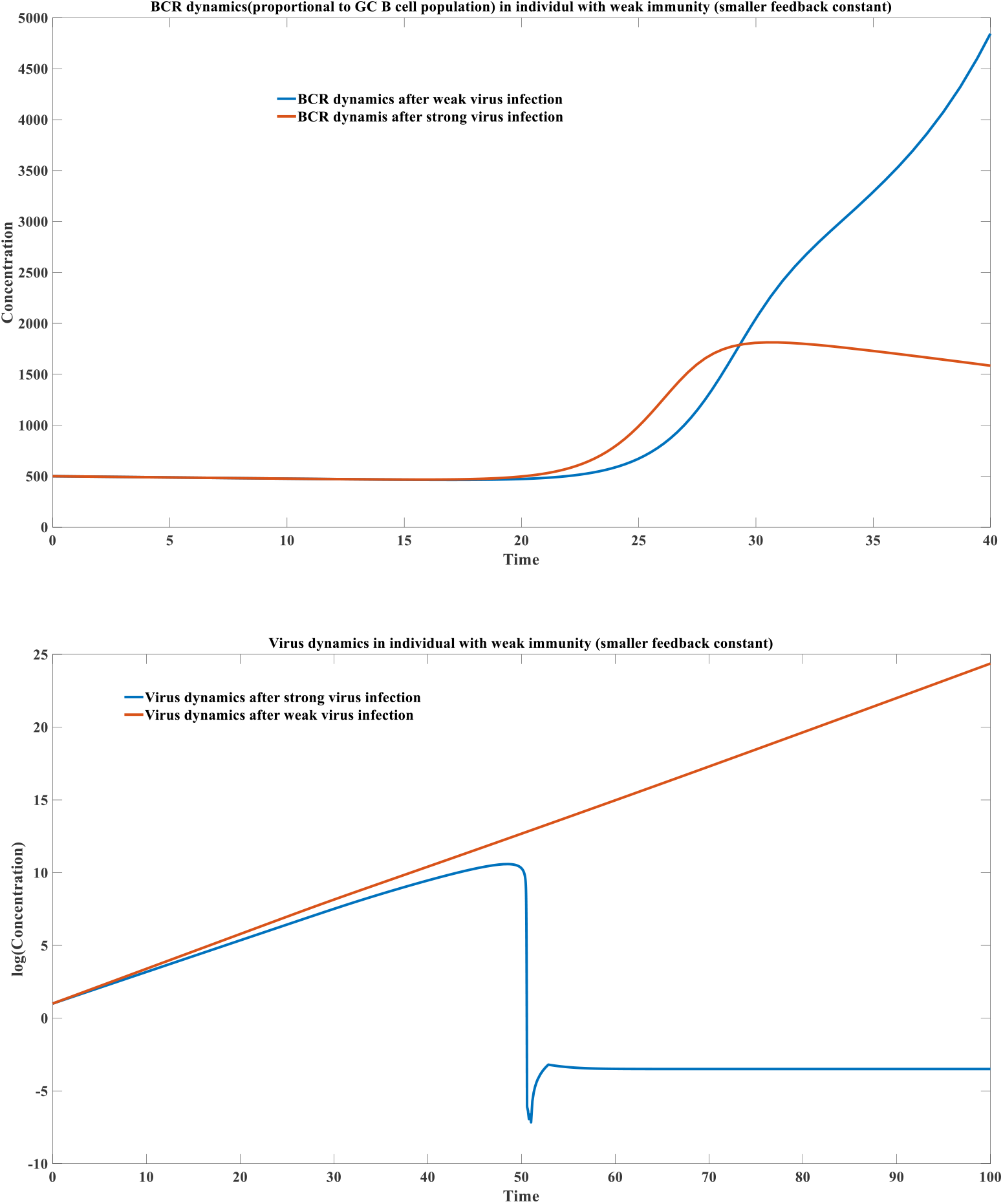
For immunocompromised individuals, excessive antigen stimulation may trigger an early-phase stress response in germinal center (GC) B cells, leading to disproportionate differentiation into plasma cells.

**Figure 1H:**
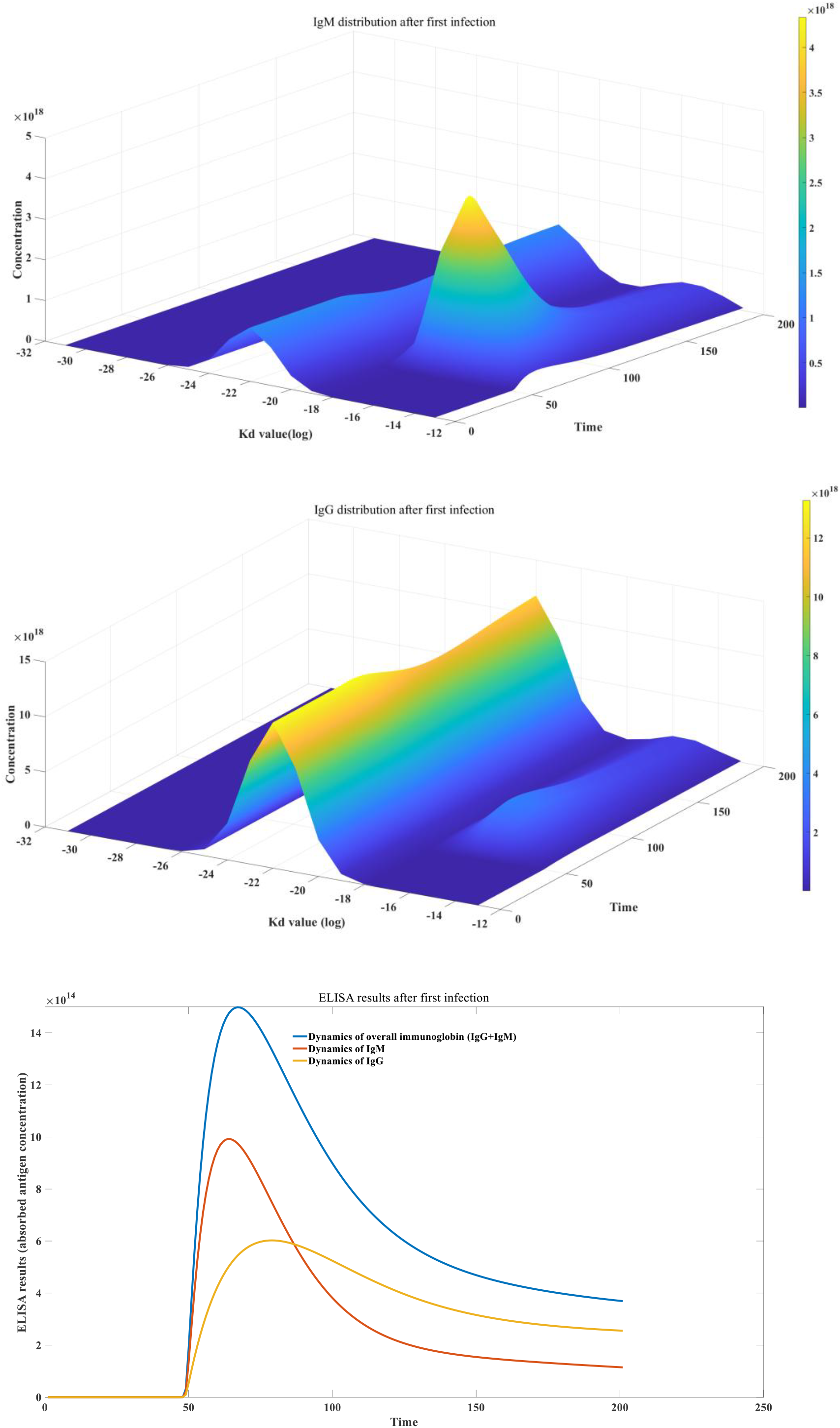
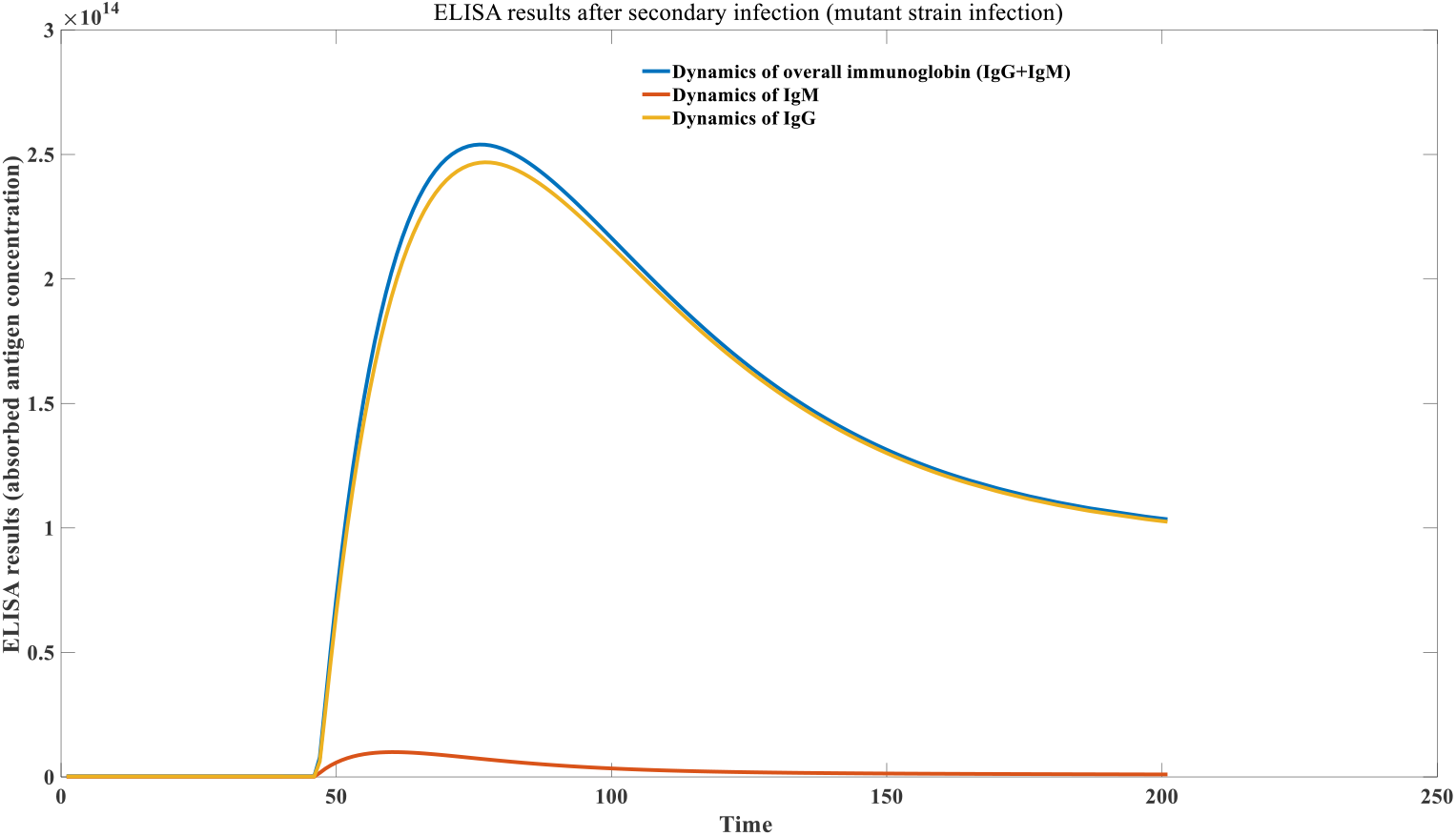
Different dynamics of IgM and IgG in primary infection and secondary infection

**Figure 1I:**
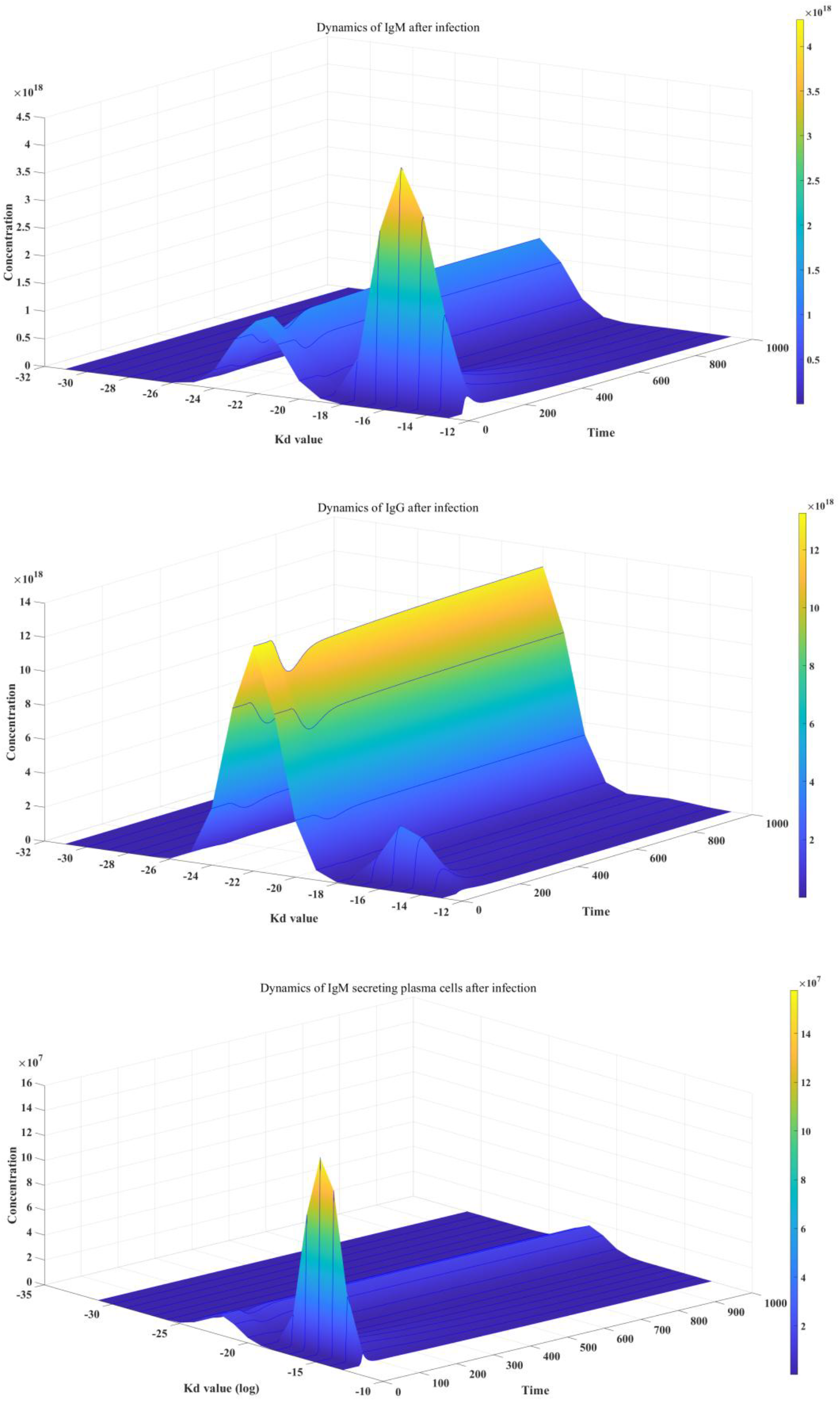
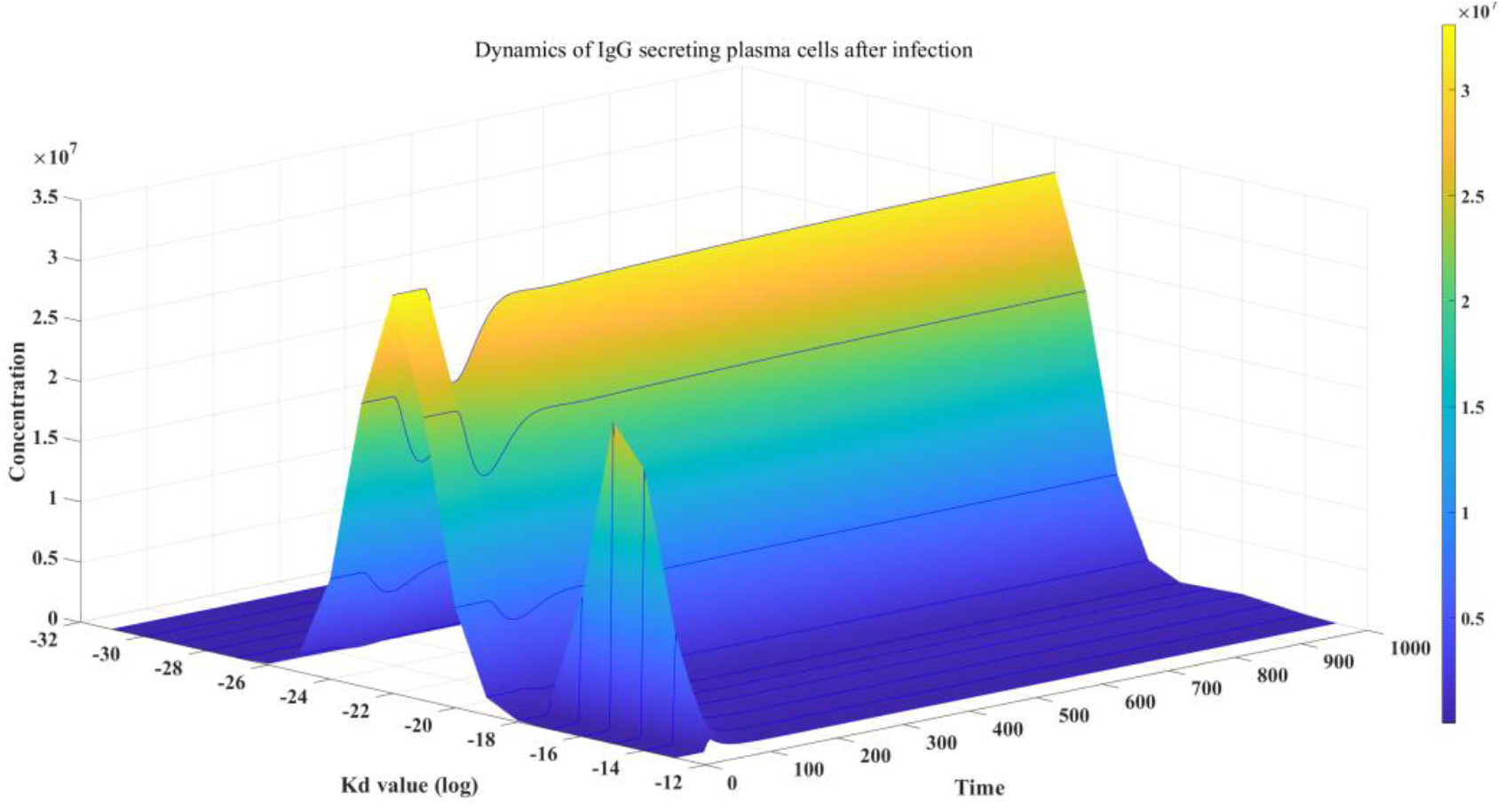
Mechanism of immune memory

The antibody attenuation observed in the experiment, especially the long-lasting antibody IgG, will continue to decline with the end of infection. This has been observed in many experiments ^[41–43]^. Interestingly, the decline of IgG is not an exponential decline process, and its decline rate gradually decreases with time. Our model believes that the late decline of specific neutralizing antibodies is a discontinuous decline, which depends on the infection of other exogenous pathogens in the later stage. If there is no subsequent infection with other pathogens, the specific neutralizing antibody IgG will be able to permanently stabilize at a level as described above, bringing an infinite protection period. However, the reality is that we will continue to be invaded by other exogenous microorganisms. When we are infected by another non-homologous microorganism, humoral immunity will be stimulated again. The final IgG-BCR has two sources: one is the expansion of self-IgG-BCR (*p*), and the other is the conversion of IgM-BCR (*1 - p*), as shown in Figure 1J. If the proportion of specific neutralizing antibodies against the first virus in total IgG is *q*, and specific IgM decays over time to a very low level, meaning its proportion of total IgM is approximately zero, then upon reinfection with a heterologous virus, the proportion of specific IgG against the first virus in the total IgG pool will decrease from *q* to *q * p*. This means that after infection with a new, non-homologous virus, the level of specific IgG against the original virus will experience a discontinuous decline. However, during infection with this new virus, total IgG levels will experience a temporary, significant increase, which we can consider to be *r*. Therefore, in the brief period following infection with the new virus or vaccination, the specific antibodies against the original virus may reach *q * p * r*, at which point this value may significantly exceed *q*. However, once infection or vaccination is complete, the specific antibodies against the original virus will eventually return to a steady state of *q * p*. The more virulent the foreign virus (the larger the replication coefficient), the higher the proportion of the new IgG antibody repertoire derived from IgM-BCR conversion, meaning that the smaller the *p* value, the greater the decline in the original antibody pool. At the same time, the more times the exogenous microorganisms are infected, the greater the attenuation of antibodies against the initial virus. This is the principle of immune memory and the discontinuous attenuation theory of specific antibodies that we proposed. Future experimenters can design simple experiments to verify this theory. The clinical data on the changes in the level of specific antibodies against the original virus after the new virus pandemic ^[44–45]^ also well verify our theory. One study found ^[45]^ that antibodies against traditional pathogens (such as syncytial virus, common coronavirus, etc.) were at a relatively stable high level before large-scale vaccination or new coronavirus infection, which led to a lower incidence of these traditional pathogens in the population. However, after the COVID-19 pandemic, these antibody levels have dropped significantly, resulting in an increased risk of infection with these traditional pathogens. When we carefully investigated their data, we found that their results perfectly verified our model. The data from March 2021 showed that the antibody level had been significantly improved compared to March 2020. New Zealand’s large-scale COVID-19 vaccination work began in February 2021. Therefore, the increase in antibody levels in March 2021 was the same as the previous increase shown by the yellow solid line in Figure 2. It was a short-term increase in specific antibodies against the original virus due to heterologous vaccination. Finally, in the surveys in 2022 and 2023, it was found that these antibody levels had dropped significantly compared to 2020.

**Figure 1J:**
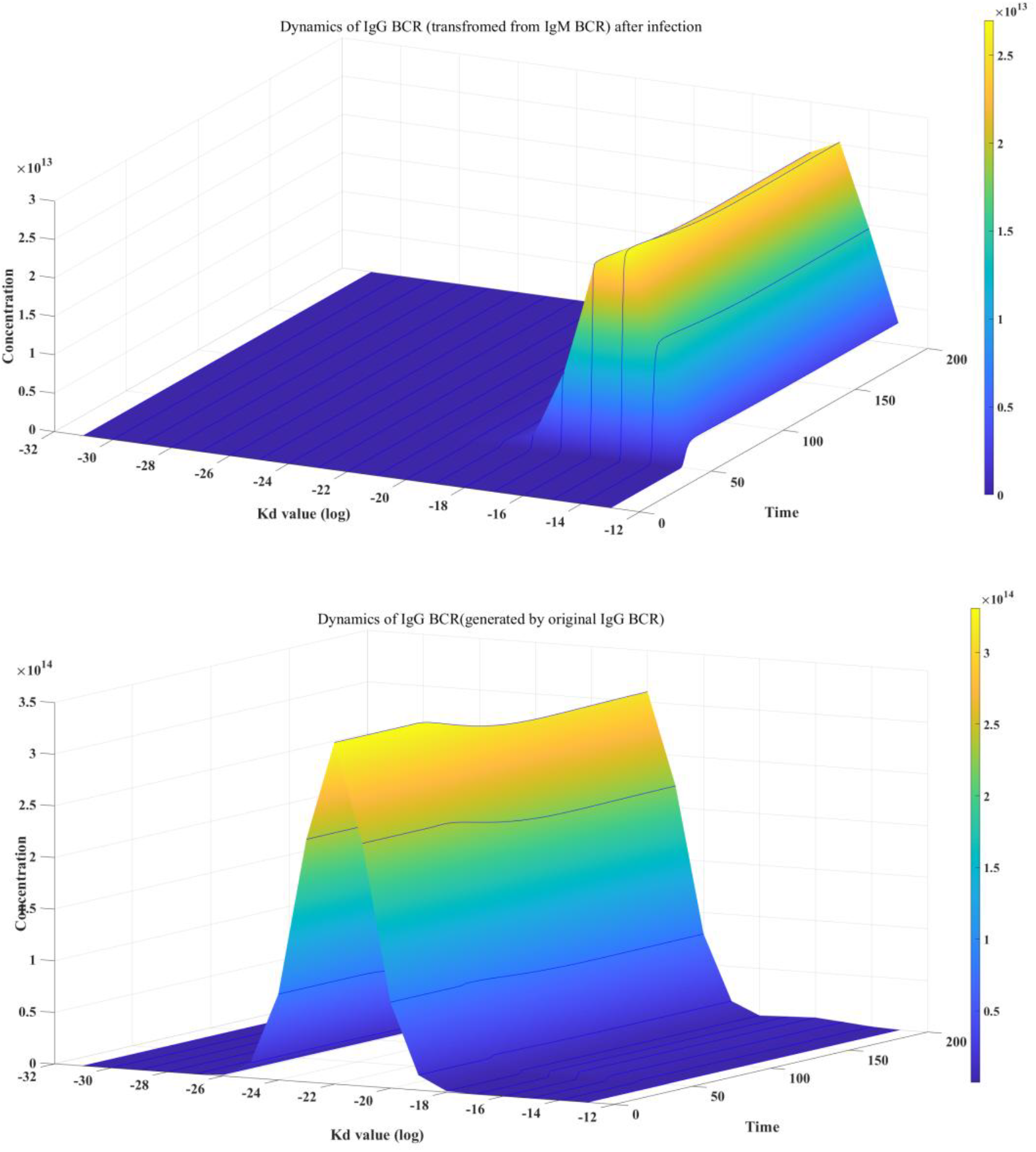
Discontinuous Decay Theory of Antigen-Specific Antibodies

**Figure 1K:**
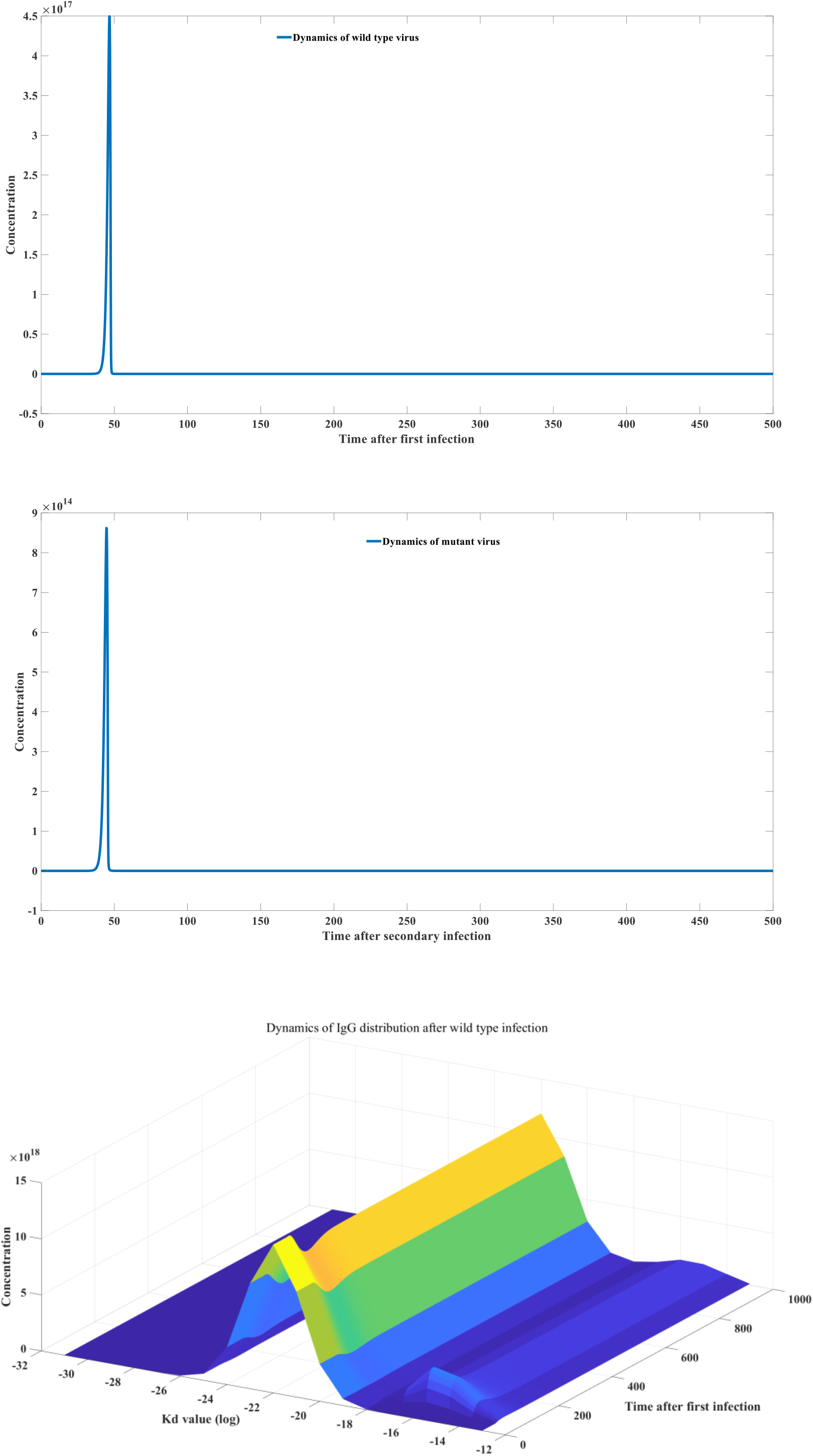
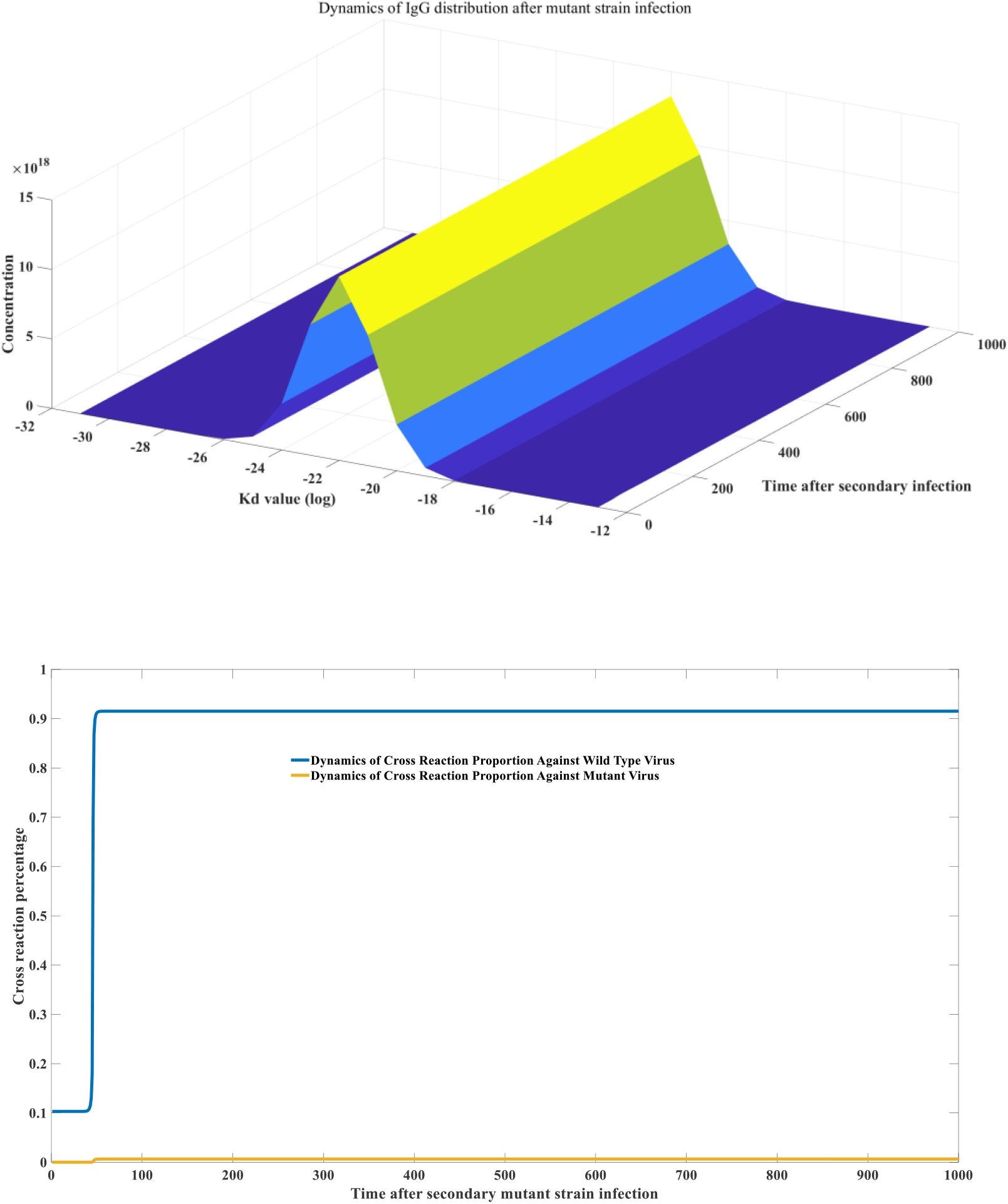
mathematical modeling of immune imprinting

Immunoblotting Imprinting, also known as the original antigenic trace, is an immunological concept that describes a phenomenon in which the immune memory generated by the body after the first encounter with a pathogen (such as a virus) has a strong and biased influence on the immune response to subsequent encounters with similar but mutated versions of the pathogen.

Simply put, the immune system will “remember” the appearance of the pathogen it first saw (the original antigen). When it encounters a “relative” or variant of the pathogen, it still tends to use antibodies and immune cells produced against the original version to respond, rather than producing a new optimized response based on the characteristics of the new version. Immunoblotting is widely present in various viral infections ^[46–47]^. However, the study of the mechanism of immunoblotting formation is still a blank field. Our model can well explain the formation principle of immunoblotting from the perspective of antibody dynamics, and at the same time determine whether a mutant strain can cause secondary infection. In addition, we can also conduct quantitative research on immune imprinting and determine the degree of immune imprinting based on viral homology, that is, calculate the proportion of B cells that can cross-bind. When the virus variation coefficient is equal to 5, the dynamic changes of the mutant virus and the change map of the new antibody library are shown in Figure 1K. At the same time, Figure 1K also gives the ratio of BCR cross-linking between the two, so as to calculate the amplitude of immune imprinting. We explained the formation mechanism of immune imprinting in the previous model ^[48]^. We have improved upon the original model by introducing antibody typing and AST cells, making it more scientific and comprehensive. Simply put, the principle of original antigenic sin is that after infection with the original virus, the antibody profile shifts toward high-binding regions, forming a bimodal profile. If the new virus is completely homologous to the original virus (sequence identity), the antibody profile will remain bimodal. Kinetic calculations indicate that secondary infection is impossible, and the immune imprinting level can be considered 100%. If the new virus is completely different from the original virus, the antibody profile returns to the original normal distribution. Kinetic calculations can simulate the kinetic changes in various compartments during infection with the new virus. However, as depicted in Figure 1J, even when the homology between the two viruses is zero, immune imprinting still exists, and BCR B cells specific for the new virus will cross-bind with the original virus at a ratio of *q * p*. When the homology between the new strain and the original strain is between 0 and 1, the antibody profile formed after infection with the original virus will be in a state intermediate between a normal distribution and a bimodal distribution. The higher the homology, the greater the proportion of the bimodal distribution. By calculating this, we can calculate the coefficient of variation of the virus that causes secondary infection and determine the specific value of the immunoblotting during the secondary infection process. The specific definition of the viral homology coefficient and the specific immune imprinting calculation method are provided in the supplementary materials.

### 2.2 Model construction of cellular immunity in adaptive immunity

The principle of cellular immunity is illustrated in Figure 2A.

DC cells, are the primary initiators of humoral immunity during viral infection, nearly all experiments tend to assume that DC cells are the primary presenters of cellular immunity. Figure 2A illustrates the basic principles of our cellular immunity model and its relationship to humoral immunity. Figure 2A shows two types of viruses, Virus 1 and Virus 2. Virus 1 is less immunogenic, lacking sufficient surface glycosylation and therefore not directly recognized by CLRs or TLRs on the surface of DC cells. Some viruses, such as LCMV and neoantigens, fall into this category of antigens. For this type of antigen, it must first bind to an antibody to form an antigen-antibody complex. This binding alters the antibody’s Fc conformation, enabling it to bind to Fc receptors on the surface of DC cells. This antigen is then transported to the DC cell for degradation and ultimately presentation to the MHC-I complex. The MHC-I-peptide complex ultimately binds to the TCR receptor on the surface of specific CD8+ T cells, ultimately activating CD8+ T cells. Therefore, for this type of antigen, the antibodies produced by humoral immunity play a very important role in stimulating cellular immunity. Infection or blocking the production of antibodies can inhibit cellular immunity, which is very significant in LCMV virus ^[49].^ At the same time, for autoimmune diseases, reducing the level of specific neutralizing antibodies can also reduce the killing effect of CD8+ T cells on their own cells, playing a role in alleviating the disease ^[50–51]^. For virus 2, the situation is more complicated, because the virus itself has very strong immunogenicity, and its protein structure may be significantly glycosylated. Therefore, receptors such as CLR and TLR on the surface of DC cells can directly bind to the virus and present it to CD8+ T cells. At the same time, the antibody-virus complex can also be captured by Fc receptors and presented to CD8+ T cells. Therefore, although humoral immunity is involved in the process of cellular immunity, blocking humoral immunity does not directly block the stimulation of cellular immunity. Interestingly, sometimes the blocking effect on humoral immunity can actually enhance the function of cellular immunity. For example, there are reports that for patients treated with rituximab, the humoral immune function of SARS-CoV-2 vaccine was affected, but the cellular immunity was improved ^[52]^. Using model 3.2.2, we conducted a simulation. As shown in Figure 1B, for patients with blocked antibody production, vaccination with SARS-CoV-2 vaccine can lead to obstacles in stimulating specific antibody levels, but can obtain higher levels of specific CD8 +T cells. Figure 2A also shows CD8 + T cell exhaustion. CD8+ T cell exhaustion is a very common phenomenon in chronic infection. When CD8+ T cells bind to infected cells or cancer cells, they initiate cytotoxic activity. In addition to the lysis of infected cells, the expression of receptors such as PD-1 on the surface of CD8+ T cells increases, resulting in a decrease in the ability of CD8+ T cells to re-recognize the infected cells. Therefore, PD-1 inhibitors are widely used in cancer treatment and chronic infection treatment to prevent the exhaustion effect caused by long-term CD8 + T cell contact with target cells.

**Figure 2A:**
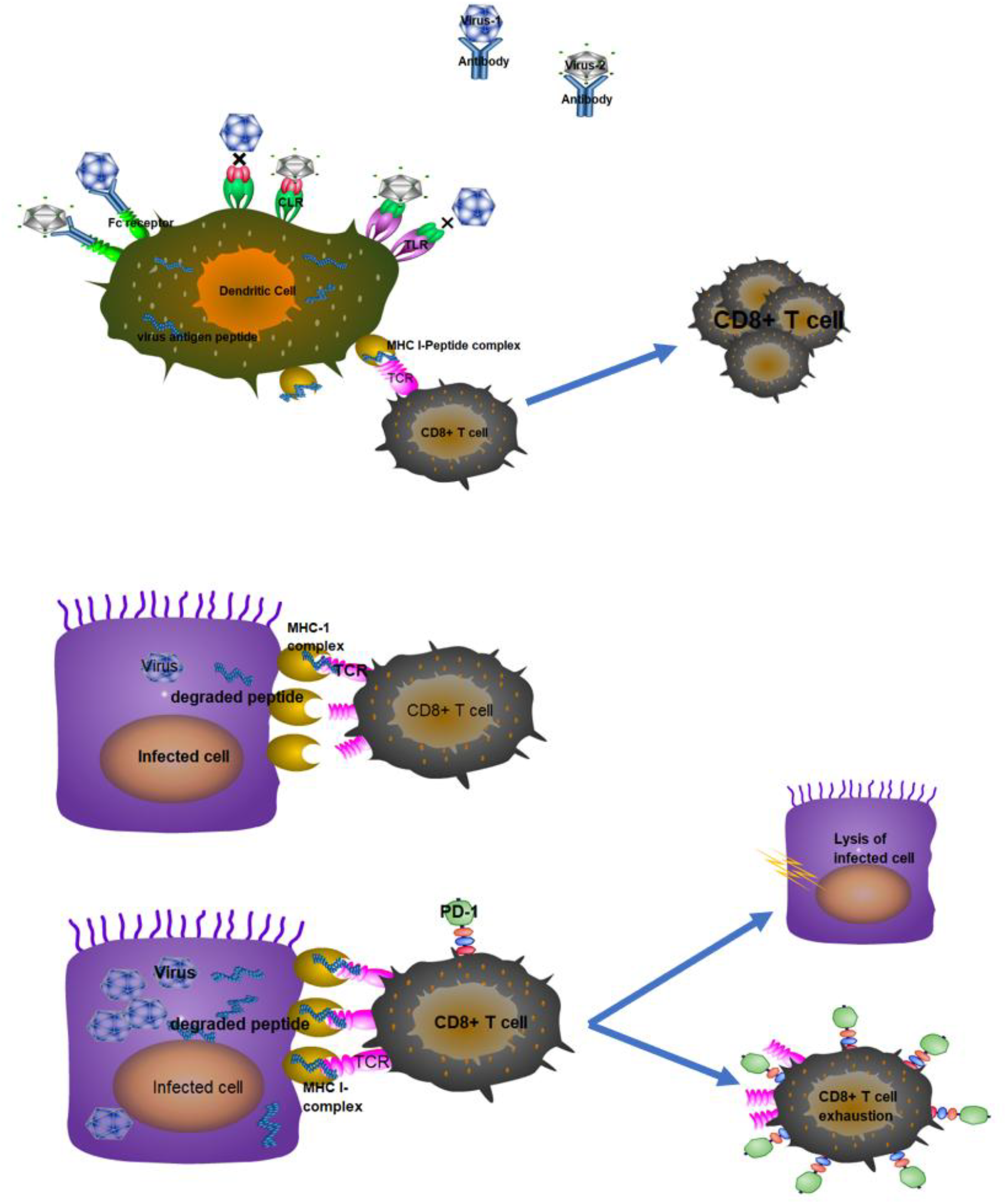
Diagram of cellular immune response

In humoral immunity, the cytotoxic effect of CD8+ T cells on target cells primarily depends on their binding affinity. The strength of this affinity is determined by two factors: the first is the sequence and corresponding structure of the TCR. CD8+ T cells with strong affinity for MHC-I-peptide complexes are activated and proliferate rapidly. The second factor is the concentration of MHC-I complexes on the target cell surface. This concentration is positively correlated with the concentration of intracellular antigen. For cancer cells, the intracellular concentration of neoantigens can remain relatively stable, so the cytotoxic effect of CD8 + T cells depends solely on the type of neoantigen and the type of CD8+ TCR. However, for virally infected target cells, the situation is more complicated. The concentration of viral antigens in target cells increases significantly over time, resulting in CD8+ T cells only recognizing target cells in the middle and late stages of infection and failing to bind to and lyse target cells in the early stages of infection. We use model 3.2.1 to simulate this phenomenon, which is important for our agent-based comprehensive humoral immunity model below. The model is very important as it can quantify factors that causes the infection to turn into a long-term chronic infection. Simply put, at a specific level of humoral antibodies and CD8+T cells, the target cells will only be lysed when the intracellular virus concentration reaches a certain threshold, then a large amount of virus will be released into the body fluid. When the specific antibody level in the body fluid is lower than a certain threshold, these viruses will not be completely neutralized, but some will be transfected into susceptible cells, causing new infections. However, due to the low virus concentration in the newly infected cells, they will not be recognized and lysed by the immune system in the early stages. The virus can continue to replicate in the cells, thus entering a new round of circulation. Our model allows for the precise quantification of two distinct immune effectors on target cell lysis: ADCC and CD8+ T cell-mediated cytotoxicity. As shown in Figure 2C, during acute infection, we can observe the dynamics of infected cells at different times. Early-infected cells lyse earlier than later-infected cells. We can also observe changes in intracellular viral concentration within infected cell populations and the dynamics of the complexes formed by viral binding to CD8+ T cells. This allows us to quantitatively characterize the cytolytic effects of CD8+ T cells and ADCC. This model can also be used to study population changes in specific CD8+ T cells during chronic infection, particularly CD8+ T cell exhaustion. We will discuss the mechanisms of chronic infection in detail in the agent-based model. Here, we primarily demonstrate the temporal evolution of CD8+ T cell-driven cell cytotoxicity in chronic infection, as shown in Figure 2D. Unlike acute infection, the number of infected cells in chronic infection undergoes a brief decline before rising again to a relatively stable plateau. Early target cell lysis is primarily driven by the ADCC effect and CD8+ T cell immunity, while later target cell lysis is primarily mediated by CD8+ T cells. We also observed that chronic infection leads to CD8+ T cell exhaustion, indicating a decrease in the level of highly active CD8+ T cells.

**Figure 2B:**
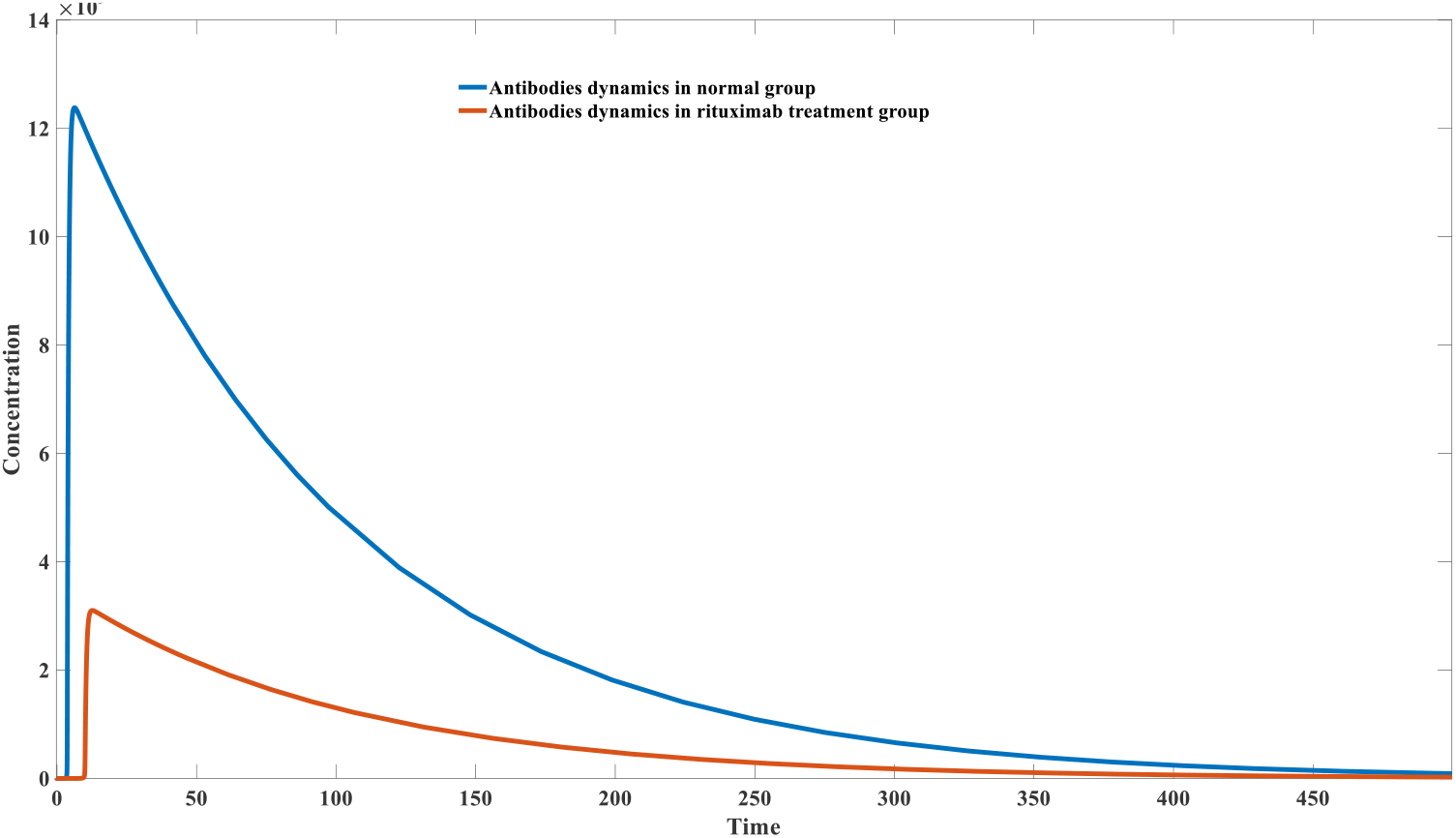
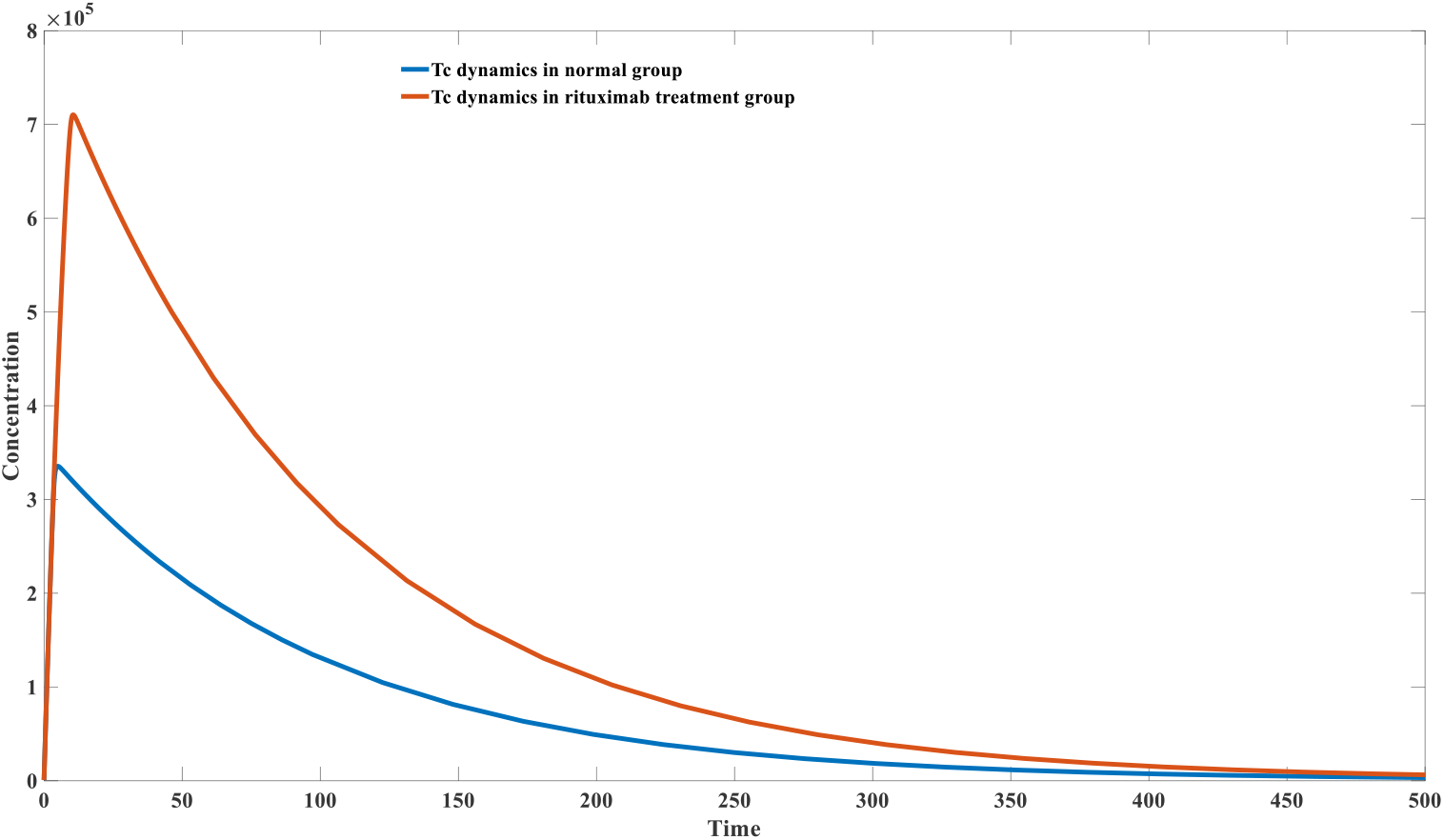
Opposite trend of humoral immune response and cellular immune response in rituximab treatment group after SARS-CoV-2 vaccination

**Figure 2C:**
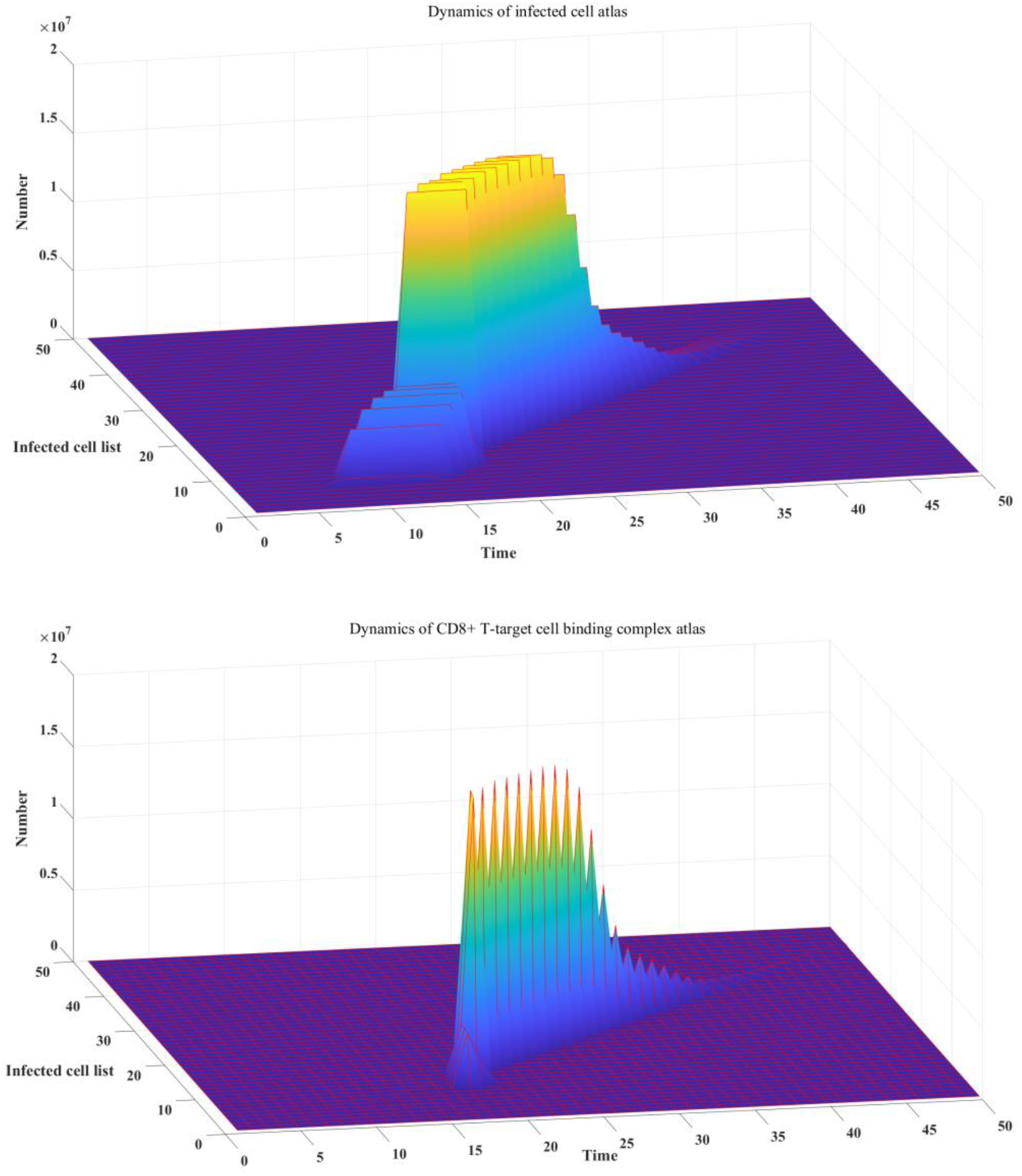
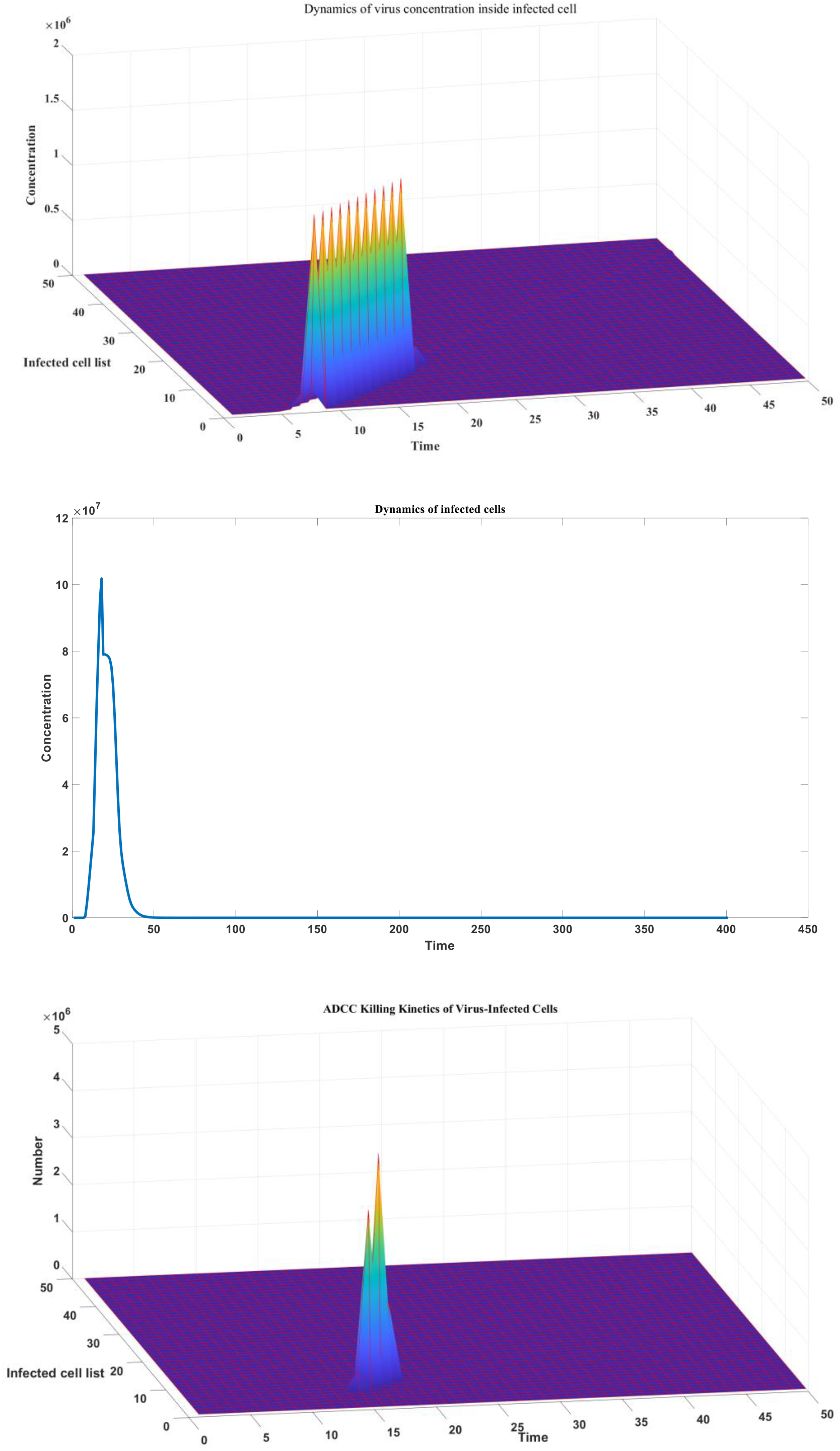
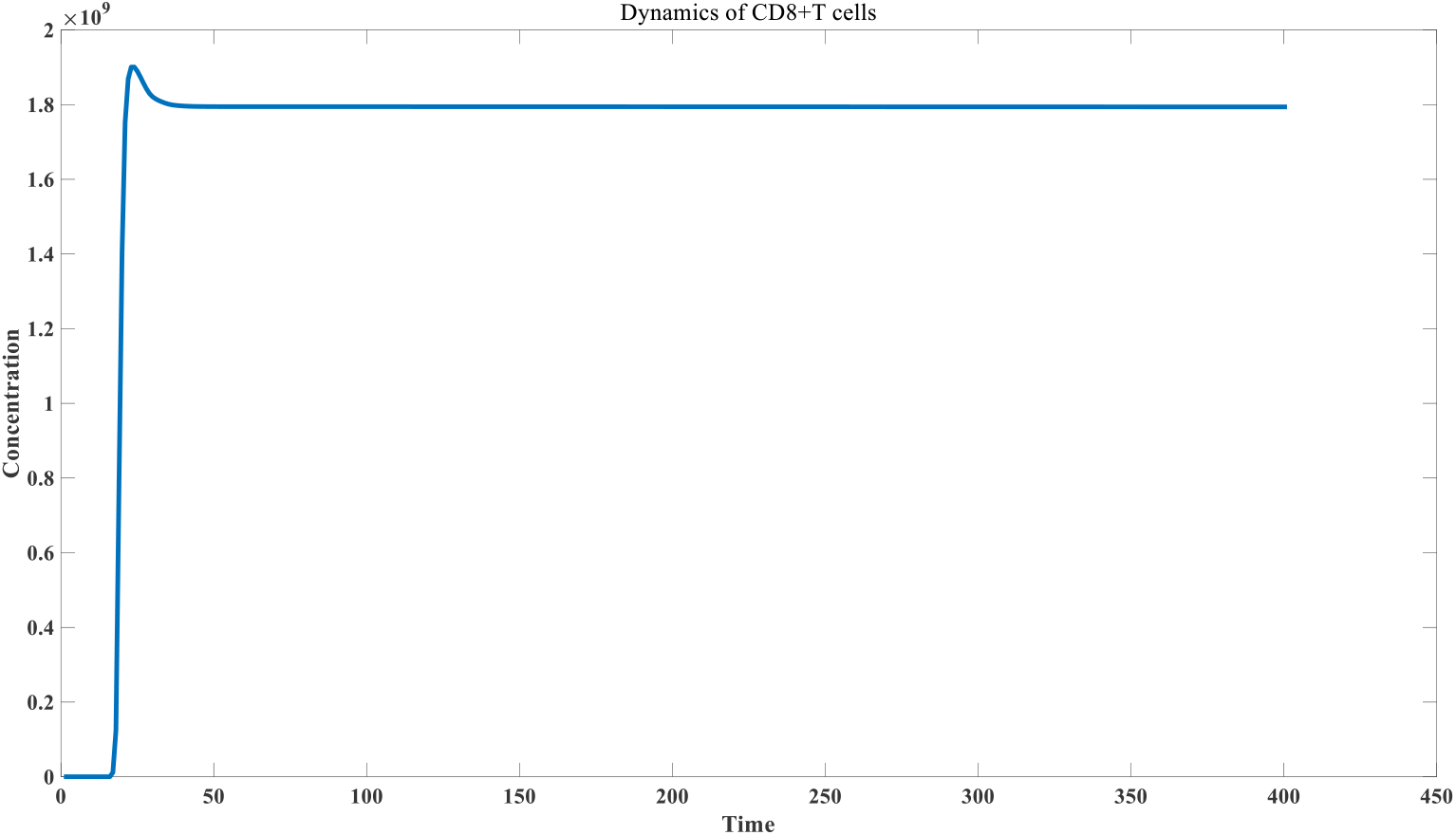
Effect of cellular immune response in acute infection

**Figure 2D:**
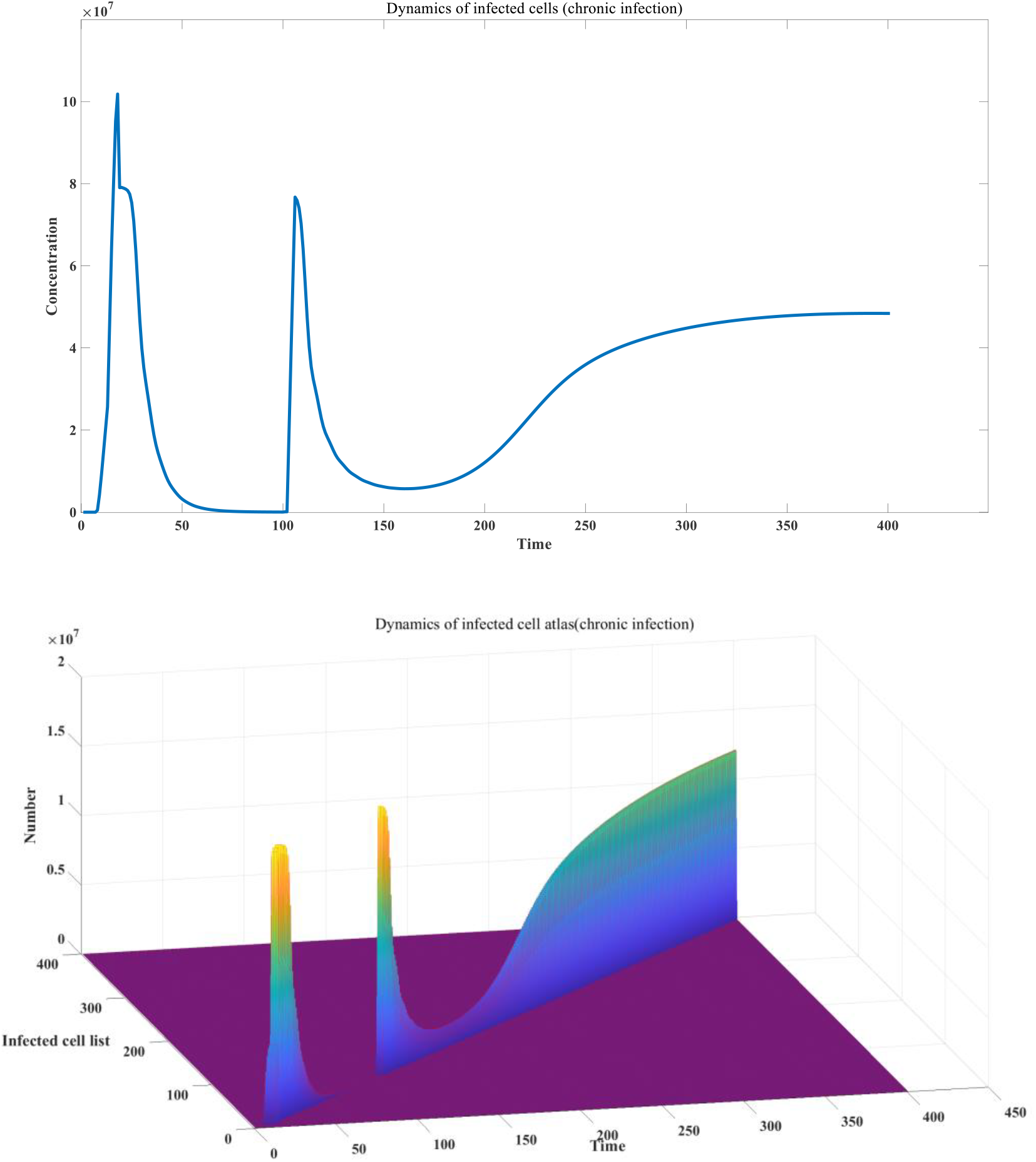
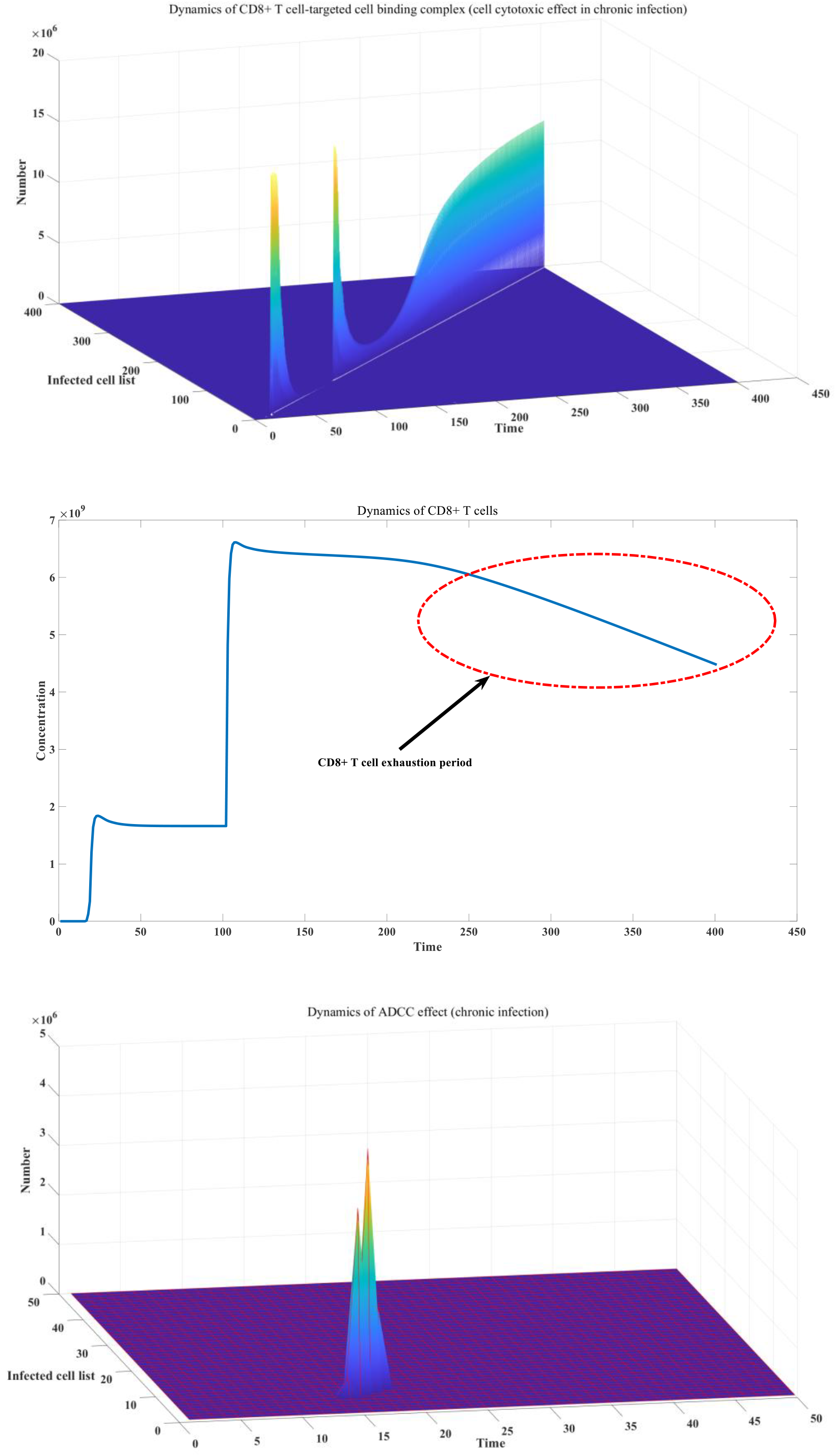
: Effect of cellular immune response in chronic infection

### 2.3 Principles of immune memory maintained by the interaction between B cells and CD4 + T cells

The antigen-presenting complex formed by the interaction between B cells and CD4+ T cells is the main feature of the germinal center. Figure 3A describes eight basic scenarios. For simplicity, we do not consider ASC cells separately. The eight scenarios are: non-memory B cells (IgM-BCR) presenting environmental antigens to effector CD4+ T cells; non-memory B cells (IgM-BCR) presenting environmental antigens to memory CD4+ T cells; memory B cells (IgG-BCR) presenting environmental antigens to effector CD4+ T cells; memory B cells (IgG-BCR) presenting environmental antigens to memory CD4+ T cells; non-memory B cells (IgM-BCR) presenting viral antigens to effector CD4+ T cells; non-memory B cells (IgM-BCR) presenting viral antigens to memory CD4+ T cells; memory B cells (IgG-BCR) presenting viral antigens to effector CD4+ T cells; memory B cells (IgG-BCR) presenting viral antigens to memory CD4+ T cells. CD4+ T cells; the specific relationship is described in detail in Model 3.3.1. As can be seen in Figure 3A, effector B cells can be replenished from stem cell differentiation, and effector CD4+ T cells can also be replenished from stem cell differentiation. However, memory B cells and memory CD4+ T cells cannot be replenished from stem cell differentiation. Their homeostasis can only be maintained by stimulating their own proliferation through environmental antigens. Therefore, the formation principle of CD4+ T cell immune memory is very similar to the immune memory of B cells discussed previously. The detailed mathematical analysis process is provided in the supplementary materials. This raises an interesting question: if the immune system can achieve the proliferation of specific neutralizing antibodies solely through B cells, why is it necessary to introduce another type of CD4+ T cell to assist in the proliferation of antibodies? Our simulation results indicate that using B cell-CD4+ T cell interactions offers two major advantages. The first is a more rapid amplification of antibodies. If antibody amplification is achieved solely through B cells, as described in Model 3.1.1, the rate of antibody production is linearly correlated with the antigen-BCR complex. However, when using the B cell-CD4+ T cell interaction model, the rate of antibody production becomes linearly correlated with the concentration of the antigen-BCR-CD4+ T cell complex. This rate of increase is much higher than the former, falling between the first and second powers of the antigen-BCR complex (the final simulation result is approximately 1.2). This rapid increase in antibody production is crucial for effective resistance to viral infection. This is particularly important when we introduce cells as the carriers of infection. Due to computational complexity limitations, we cannot currently incorporate the B cell-CD4+ T cell interaction process into our agent-based model. If we use the simple model of antibody regeneration, all infections will inevitably transition to a chronic stage in agent-based model. In reality, only a fraction of infections progress to the chronic stage during the long process of host-virus evolution, while the vast majority are completely cleared. This is largely due to the presence of CD4+ T cells. The interaction between CD4+ T cells and B cells provides a stronger driving force for antibody production. This rapid increase in antibodies helps break the antibody concentration threshold for chronic infection, thereby achieving complete eradication of the pathogen. This interaction model offers another advantage: it can cope with rapidly mutating variant strains. B cell antigens primarily target the tertiary structure of viral antigens, and relatively few point mutations can significantly alter the affinity of the BCR for the original epitope. However, T cell memory formed by CD4+ T cells targets viral peptide sequences, which are relatively conserved primary sequences and do not undergo significant changes with viral mutations. Therefore, T cell memory formed by CD4+ T cells is insensitive to viral mutations, and these memory CD4+ T cells can provide a large amount of CD4+ T cell support for antibody production during secondary infection with mutant strains.

**Figure 3A:**
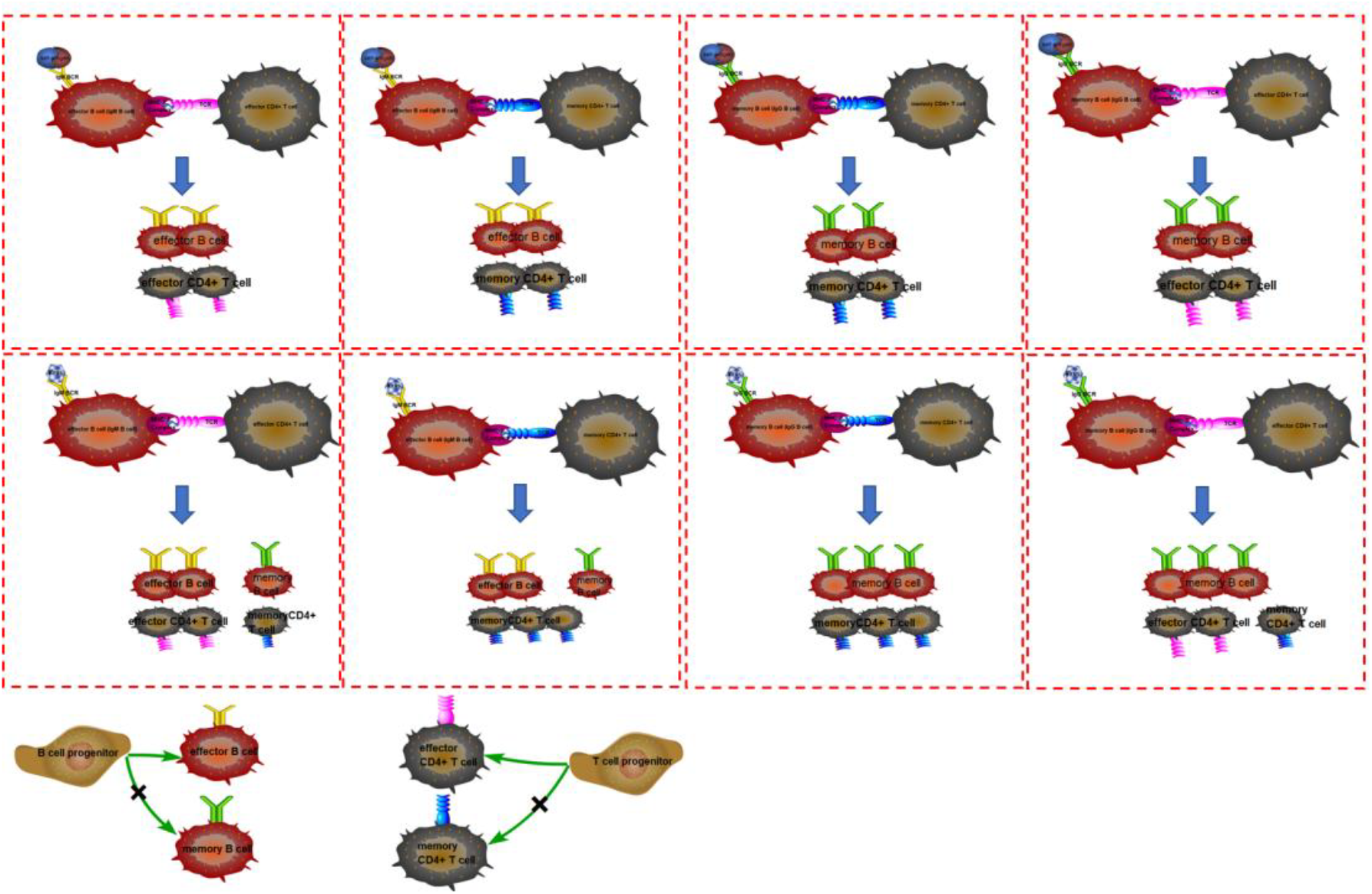
Diagram of humoral immune response considering CD4+T cell and B cell interaction

Due to computational limitations, we cannot currently calculate a large number of BCR and TCR combinations simultaneously. For example, if there are *N* types of BCRs and *M* types of TCRs with binding kinetic parameters, there will be *8*N*M* types of B-CD4+ T cell complexes. When the values of *N* and *M* are large, this poses a significant challenge for solving conventional ODE equations. Therefore, in practice, we restrict the diversity to one type. Figure 3B examines CD4+ T cell diversity. This model includes two types of BCRs (strong and weak binding) and 10*10 types of TCRs (combinations of 10 different positive binding coefficients and 10 different negative dissociation constants). The changes in virus concentration are shown in figure 3B. Furthermore, we can see that after viral infection, the distribution of CD4+ T cells changes significantly, with a clear peak in the high binding range. This is the mechanism by which CD4+ T cell immune memory is generated.

**Figure 3B:**
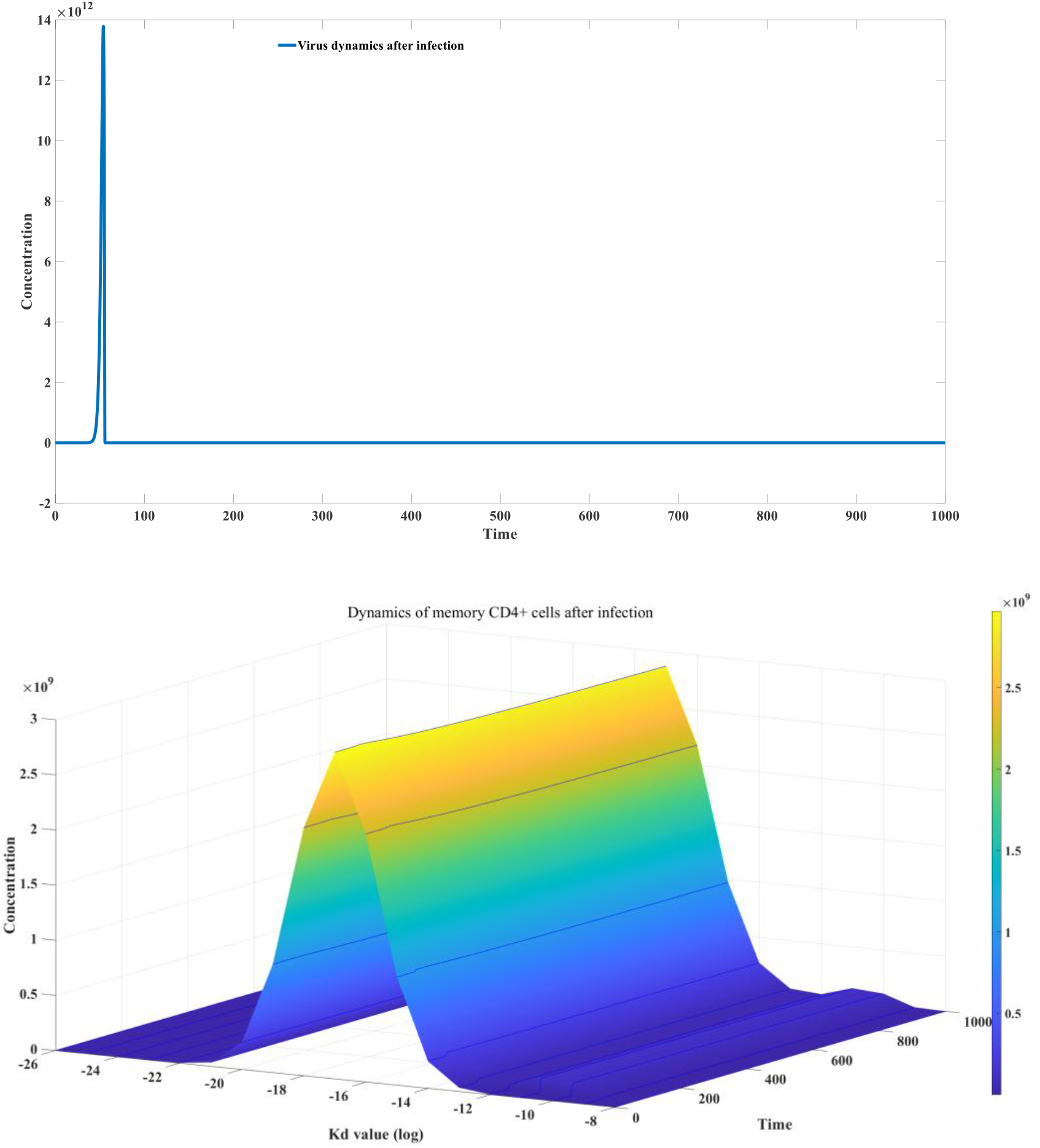
Dynamics of CD4+ T cells after infection

Figure 3C examines B cell diversity. This model includes two types of TCRs (strong binding and weak binding) and 10*10 types of BCRs (a combination of 10 different positive binding coefficients and 10 different negative dissociation constants). The changes in viral concentration are shown in the figure. We can see that after viral infection, the distribution of B cells changes significantly. IgM-BCR produces a clear peak in the high binding range, but this peak disappears rapidly over time, indicating that IgM-BCR is not the carrier of B cell immune memory. IgG-BCR also produces a small peak in the high binding range, which is maintained for a long time.

We further studied the differences between B cell memory and T cell memory. For the two models mentioned above, only one-sided memory was retained to observe its effect on secondary infection. Figure 3D shows the results of the CD4+T cell diversity model when the distribution of CD4+T cells after infection was compared with the original CD4+T cells. It can be seen from the figure that CD4+T cells can indeed reduce the peak concentration of the virus in secondary infection, thanks to its ability to quickly help the proliferation of specific antibodies, but this range is very limited and cannot effectively prevent the occurrence of secondary infection. When those rapidly mutating RNA viruses undergo point mutations, although the body retains almost complete memory CD4+T cells, it still cannot prevent the occurrence of secondary infection. The symptoms of secondary infection may be significantly weaker than the initial infection. This is consistent with the infection of SARS-CoV-2 and influenza variants ^[53–55]^. The protective effect of B cell memory against secondary infection of the original strain is more direct and effective. As shown in Figure 3E, when the B cell diversity model is used, when the distribution of B cells after infection is retained, the virus cannot be effectively proliferated after the secondary virus invasion; instead it is quickly cleared. This shows that the protective effect of B cell memory is more direct and efficient. However, as described above, B cell memory is very sensitive to viral mutations. This can be quantitatively reflected in the calculation of immune imprinting. When the virus undergoes point mutations, the dynamic distribution of BCRs for the new strain will greatly deviate from the original normal distribution. Therefore, B cell memory cannot effectively respond to rapidly mutating strains. Despite this, the partially retained B cell memory and the resulting immune imprinting will still greatly reduce the peak viral concentration during secondary infection. The short-term effect of B cell memory and the long-term effect of T cell memory have been very clearly reflected in SARS-CoV-2 infection ^[56–58]^.

**Figure 3C:**
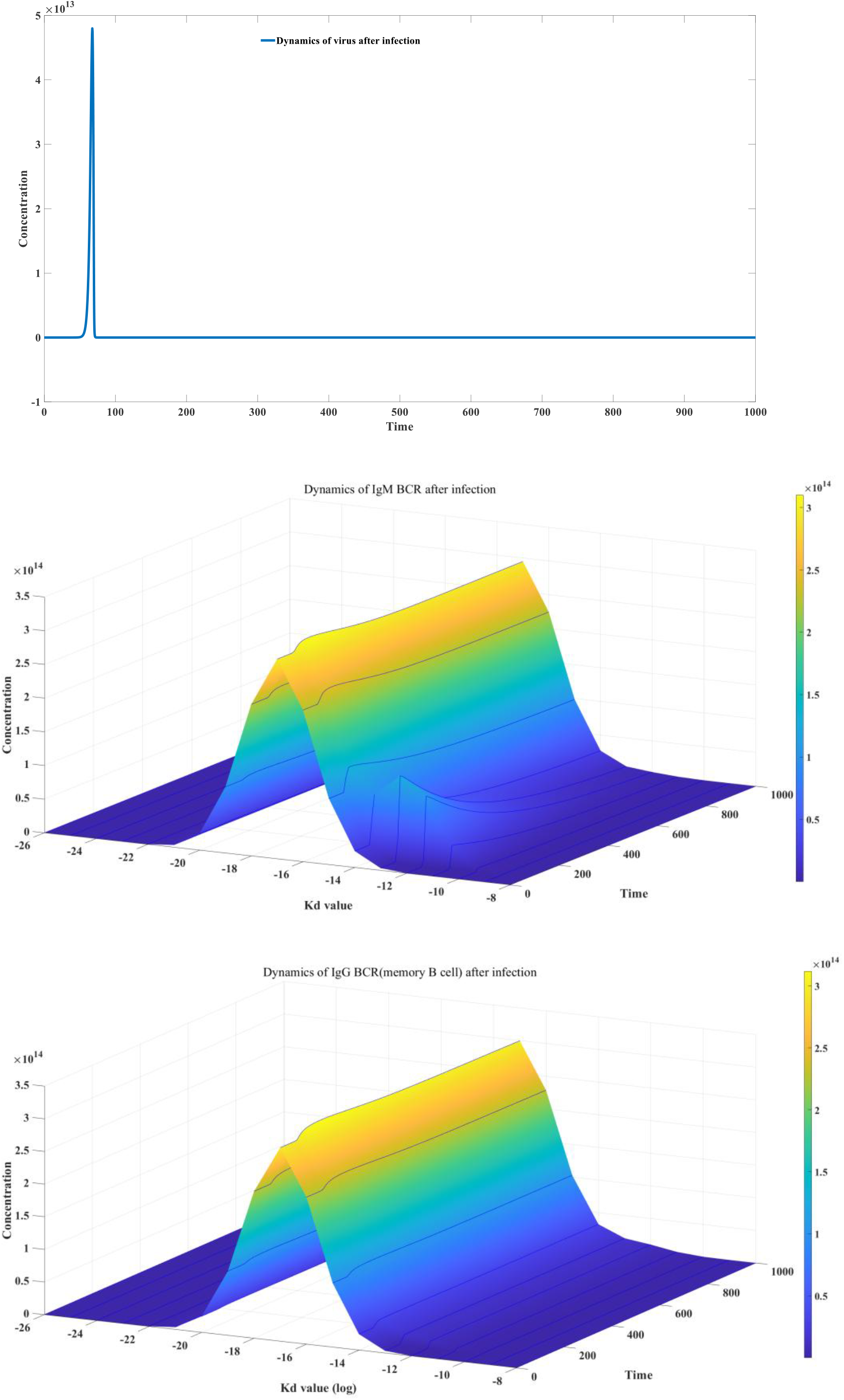
dynamics of B cell response after infection

**Figure 3D:**
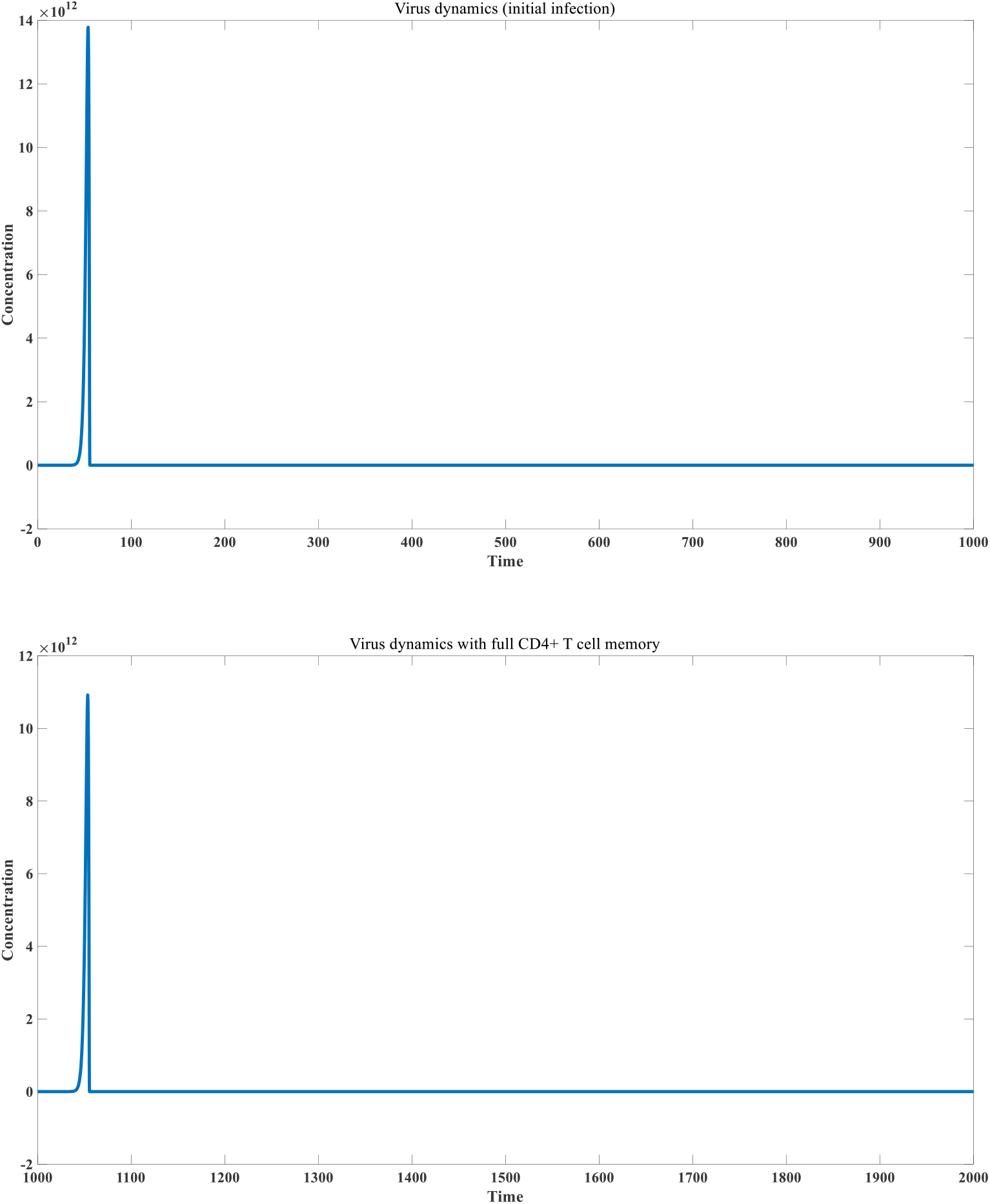
effect of CD4+ T cell memory in preventing secondary infection

**Figure 3E:**
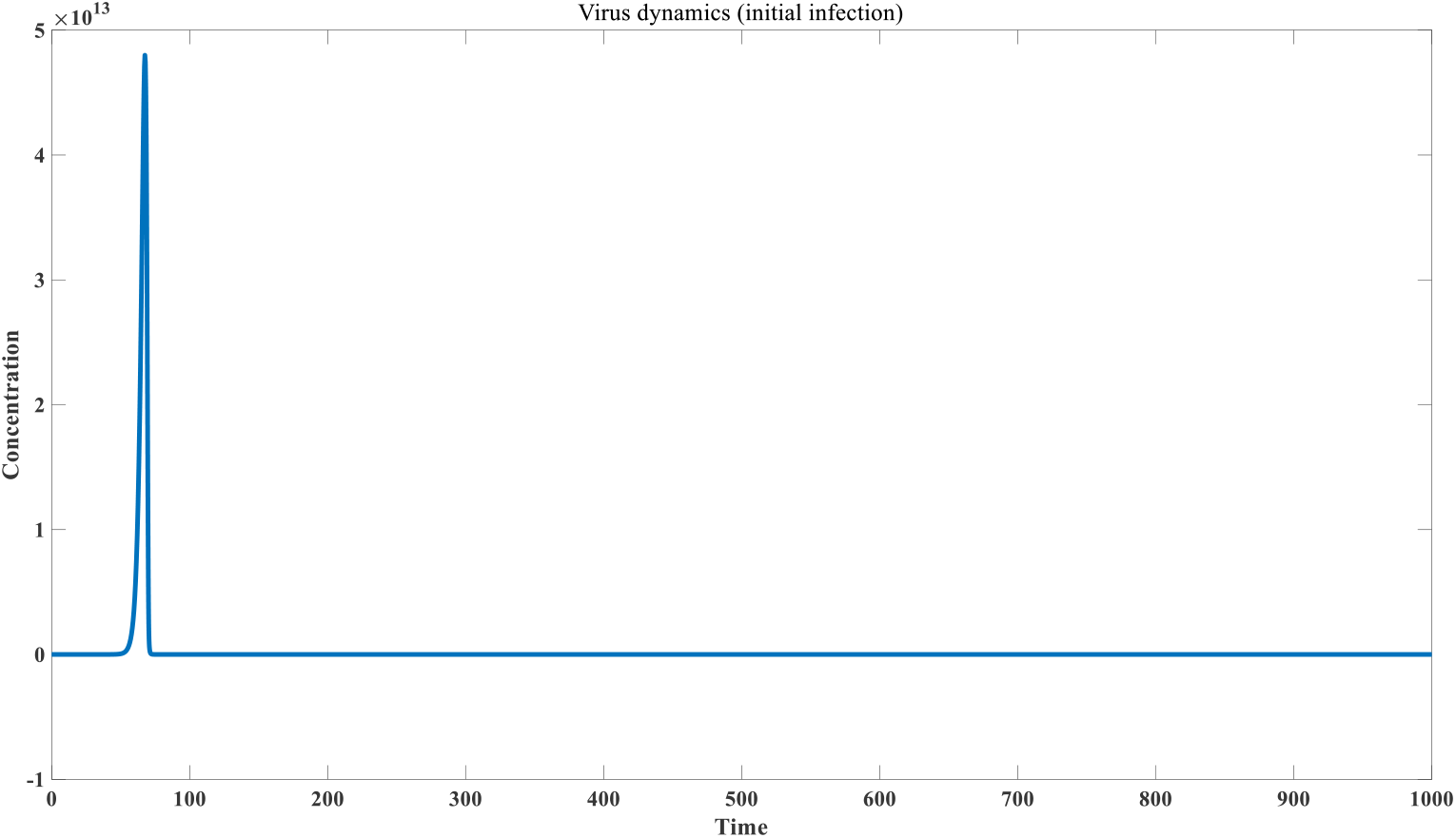
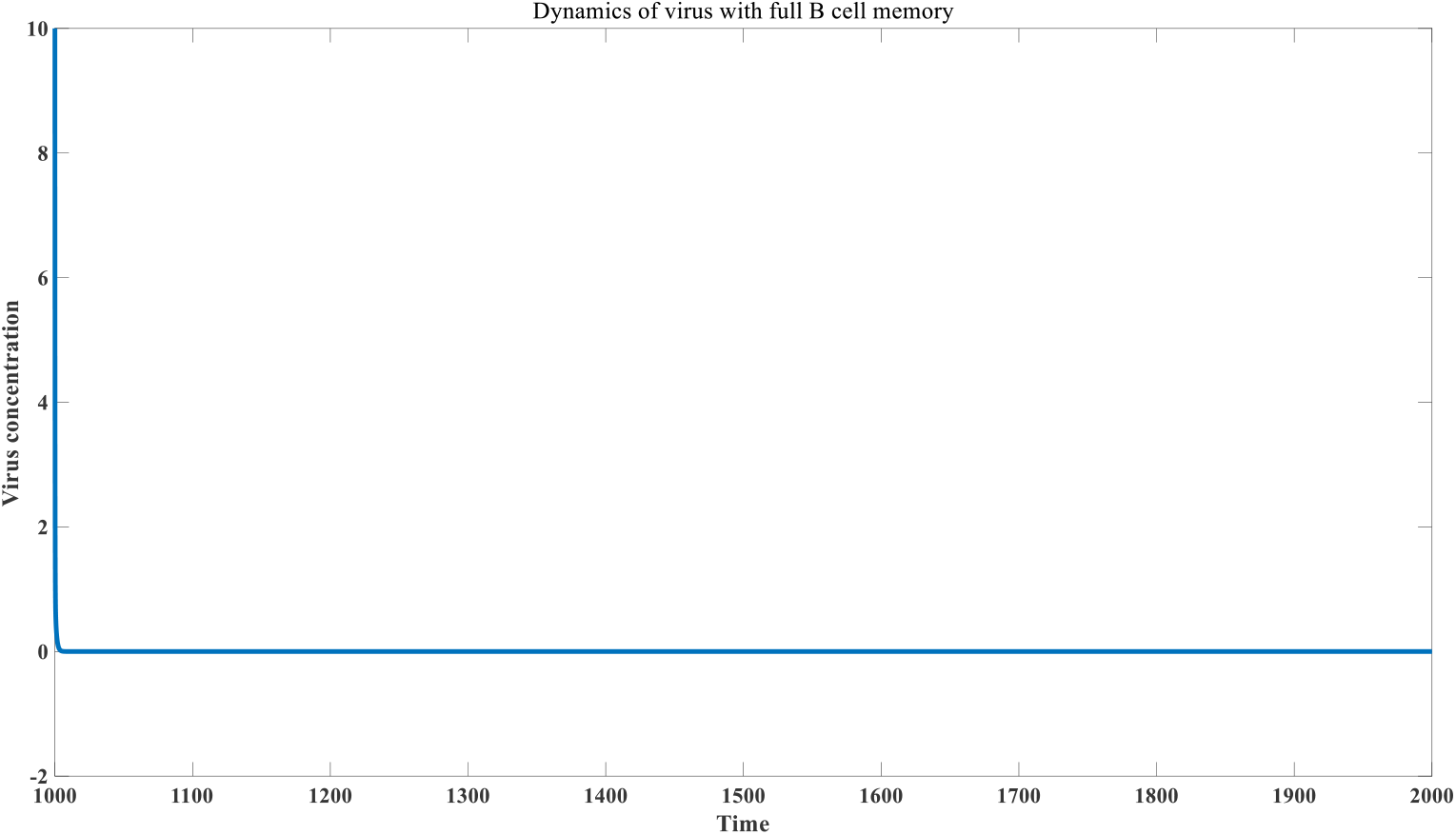
Effect of B cell memory in preventing secondary infection

Therefore, we need to have a deeper understanding of vaccines. For rapidly mutating viruses, the key factor in secondary infection lies in viral mutation, not antibody decay. For these viruses, vaccination effectiveness cannot simply be measured by the duration of protection. As the virus mutates further, secondary or even multiple infections may occur, but the severity of these secondary infections is significantly less than that of the initial infection. For infection prevention, vaccination with B cell antigens with a fixed tertiary structure can be more effective because they not only reshape the distribution of the B cell antibody repertoire but also the distribution of CD4+ T cells. Even if the antigen epitope drifts later, the immunological imprinting effect can still provide strong protection. However, compared to CD4+ T cell peptide antigens (or MHC class II antigens), B cell antigens can deplete antibodies and BCRs. Therefore, injecting large amounts of B cell antigens after infection generally does not have a positive effect. This is why B cell antigens are generally not used in cancer therapeutic neoantigens. This point will be discussed in detail in Section 5.

### 2.4 An agent - based model that comprehensively considers cellular and humoral immune processes

To more accurately investigate the activation of humoral and cellular immunity following viral infection, we constructed a complex agent-based model, treating infected cells at different moments as distinct agents. As shown in Figure 4A, the viral concentration within target cells increases over time after infection. Cell lysis occurs when the combined viral load and antibody distribution in vitro meet the ADCC threshold. Cell lysis also occurs when the combined viral load and specific CD8+ T cell population in vitro meet the CD8+ T cell killing threshold. After cell lysis, the virus is released into the body fluids, where it interacts with antibodies. Strongly binding antibodies (yellow antibodies in the figure) are present in smaller numbers, but compete with the antigen for a significant proportion. Antibodies with moderate binding (purple antibodies in the figure) are present in larger numbers and can also bind to a large number of virus particles, forming virus-antibody complexes. At this point, the virus population decreases (from 5 to 2 virus particles), but the virus cannot be completely neutralized. Viruses can still infect new cells, leading to a new round of infection. The infected cell population at time *i* forms a new agent. In addition to being eventually degraded, the antigen-antibody complex will also feedback stimulate the generation of more corresponding antibodies. If the regeneration coefficient is 2, as shown in the figure, the number of yellow high-binding activity antibodies becomes 2, and the medium-binding antibodies are also improved to 5, but the level of low-binding antibodies has not changed. This change in the antibody distribution map will affect the neutralizing ability of the next round of antibodies. When the level of high-binding antibodies rises to a certain level, the viruses released into the body fluids by somatic cell lysis can all be neutralized by antibodies. At this time, no new infection will occur and the virus has been completely eliminated. Introducing a cell model can more accurately simulate the virus infection process, because cells have the function of compartments. Once the virus enters the cell, it enters an environment blocked by antibodies. Antibodies cannot directly act on the virus inside the infected cell. This is the mechanism of chronic infection. Furthermore, both ADCC and CD8+ T cell-induced cell killing are positively correlated with intracellular viral concentration. Since the virus continues to proliferate within infected cells, defining the infected cell population by the time of infection is reasonable, as the number of cells infected at time *i* will remain constant or decrease in the future, while the intracellular viral concentration will only increase until it suddenly reaches zero after lysis. When antibody levels rise slowly (possibly due to three factors: weak humoral immunity, secondary infection caused by viral mutants, and low viral replication activity), this virus-host interaction may reach a steady state. In other words, a state can be reached where, under the combined influence of ADCC and CD8+ T cell killing, the intracellular viral concentration in infected cells can only reach a low threshold, at which point the cells lyse. However, the concentration of specific neutralizing antibodies in the body fluids cannot guarantee complete viral clearance, leading to a new round of infection. This situation can approach a long-term stable state, resulting in chronic infection. During chronic infection, viral concentrations are significantly lower than during acute infection. However, the harm of chronic infection is equally devastating because infected cells continue to die, and the immune system’s ability to kill infected cells is no less potent than during acute infection. Our agent-based model allows for more quantitative analysis of how various components change over time. This includes changes in the kinetics of infected cells (including newly infected and lysed cell kinetics), viral dynamics in body fluids, viral dynamics within infected cells, dynamics of IgM-BCR and IgG-BCR B cells, changes in IgM and IgG distribution patterns, dynamics of various antibody-antigen complexes, dynamics of environmental antigens, dynamics of specific CD8+ T cells, dynamics of infected cell lysis by ADCC, dynamics of infected cell lysis by cellular immunity, and dynamics of susceptible cell populations. This will help elevate immunological research to a new level of precision and quantitative analysis. Figure 4B depicts the dynamics of various components during acute infection.

**Figure 4A:**
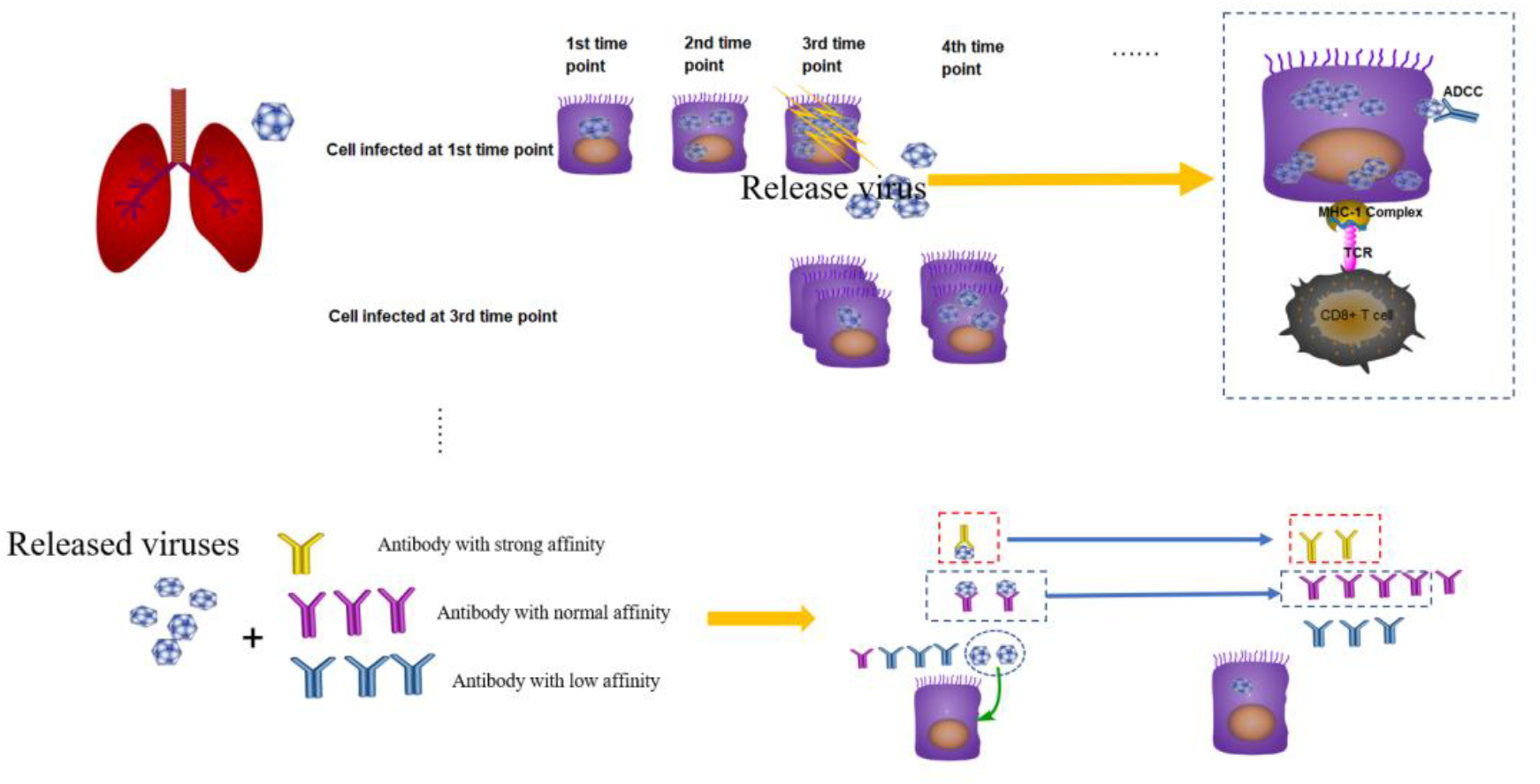
Overall Methodology Flowchart

**Figure 4B:**
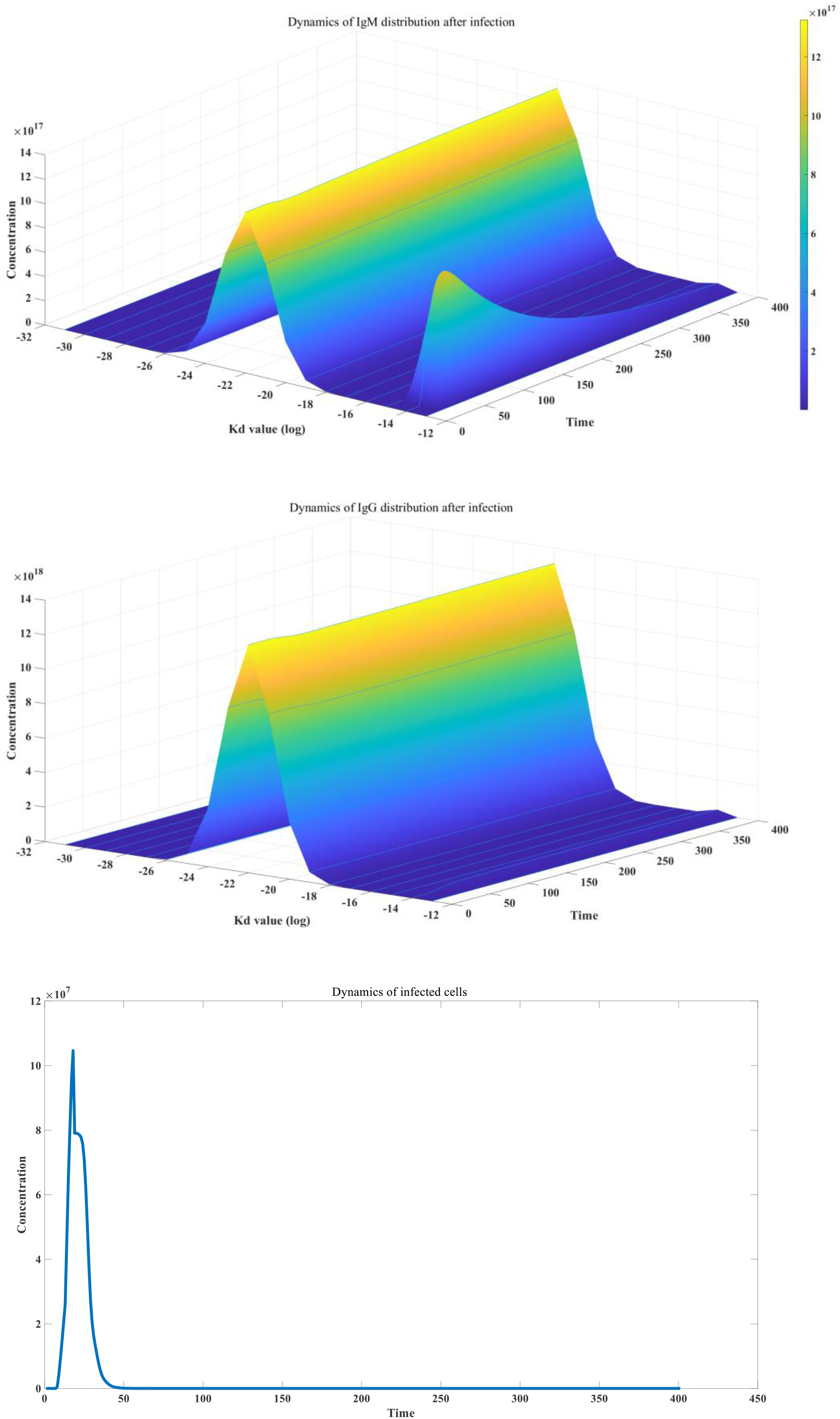
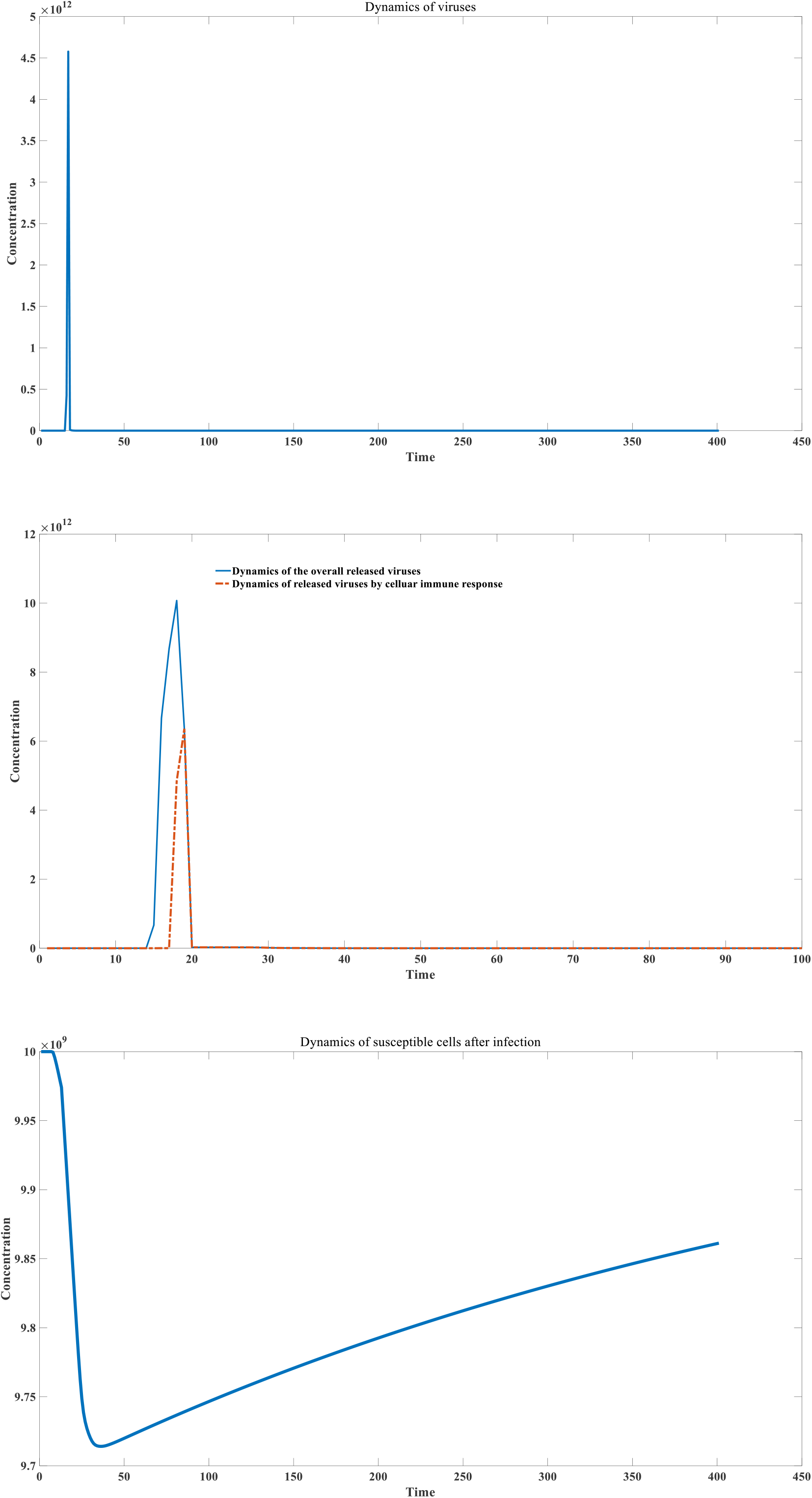
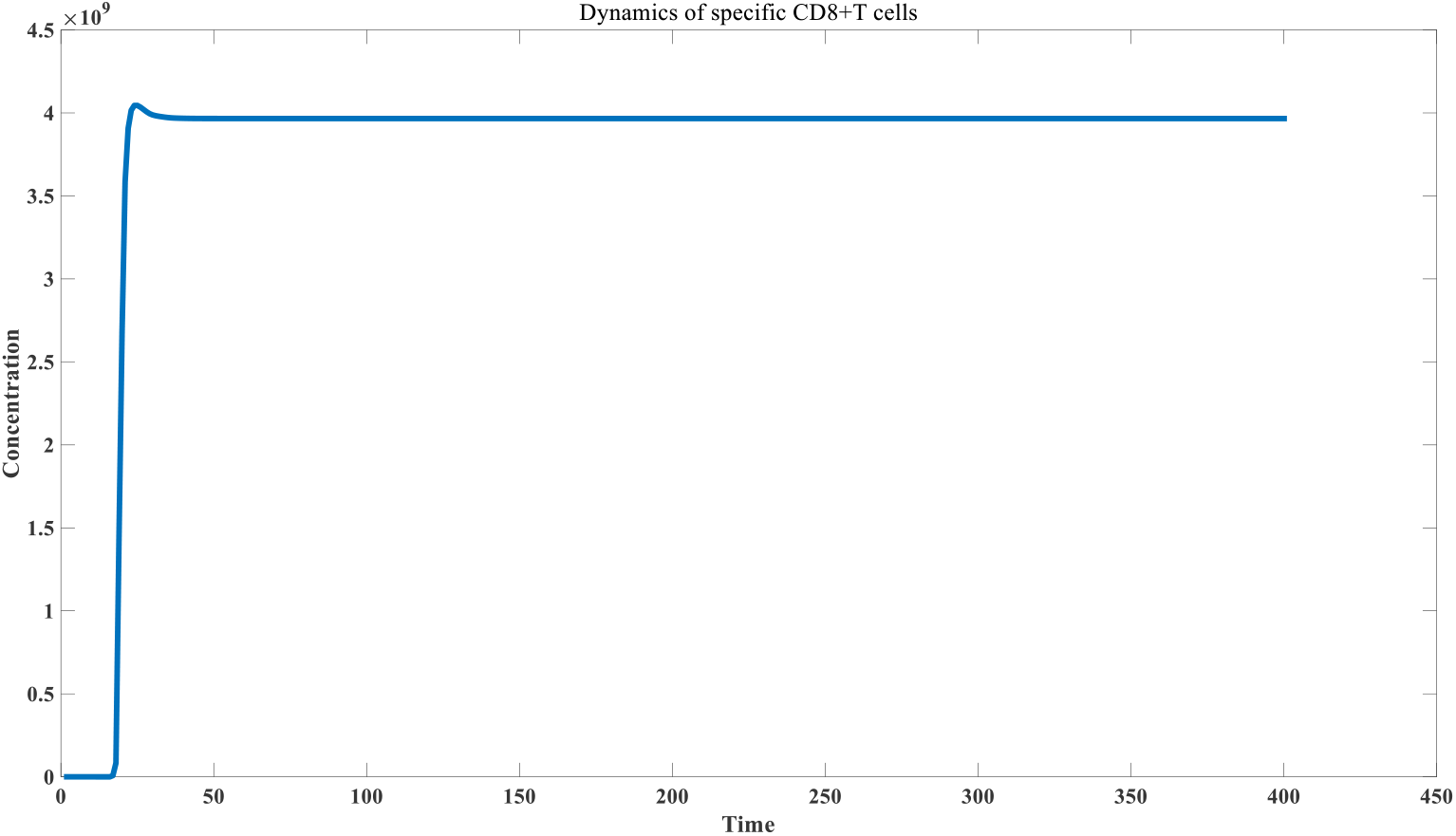
Virus-host interaction in acute infection

We further discussed the conditions for the occurrence of chronic infection. Chronic infection is a very common phenomenon. Chronic infection caused by certain viruses, such as HBV, has been extensively studied ^[59–60]^. Many recent experiments have shown that sequelae caused by acute viral infection, such as COVID-19, may also be caused by chronic infection ^[61–63]^. Here, we simulated three situations that are prone to chronic infection. The first is when the body’s immunity is low. In our model, this is reflected in a low antigen-antibody complex feedback coefficient, as shown in Figure 4C. When the body’s immunity is low, it is easy to turn into a chronic infection stage after acute infection. The characteristics of chronic infection are that the number of infected cells remains at a high level, but at this time the virus concentration in the infected cells is very low, the virus concentration in the body fluids is also low, and CD8+ T cell exhaustion occurs. Compared with people with strong immunity, an important reason for chronic infection caused by low immunity is insufficient antibody (including IgM and IgG) induction level, which can be seen by comparing Figure 4B and Figure 4C.

**Figure 4C:**
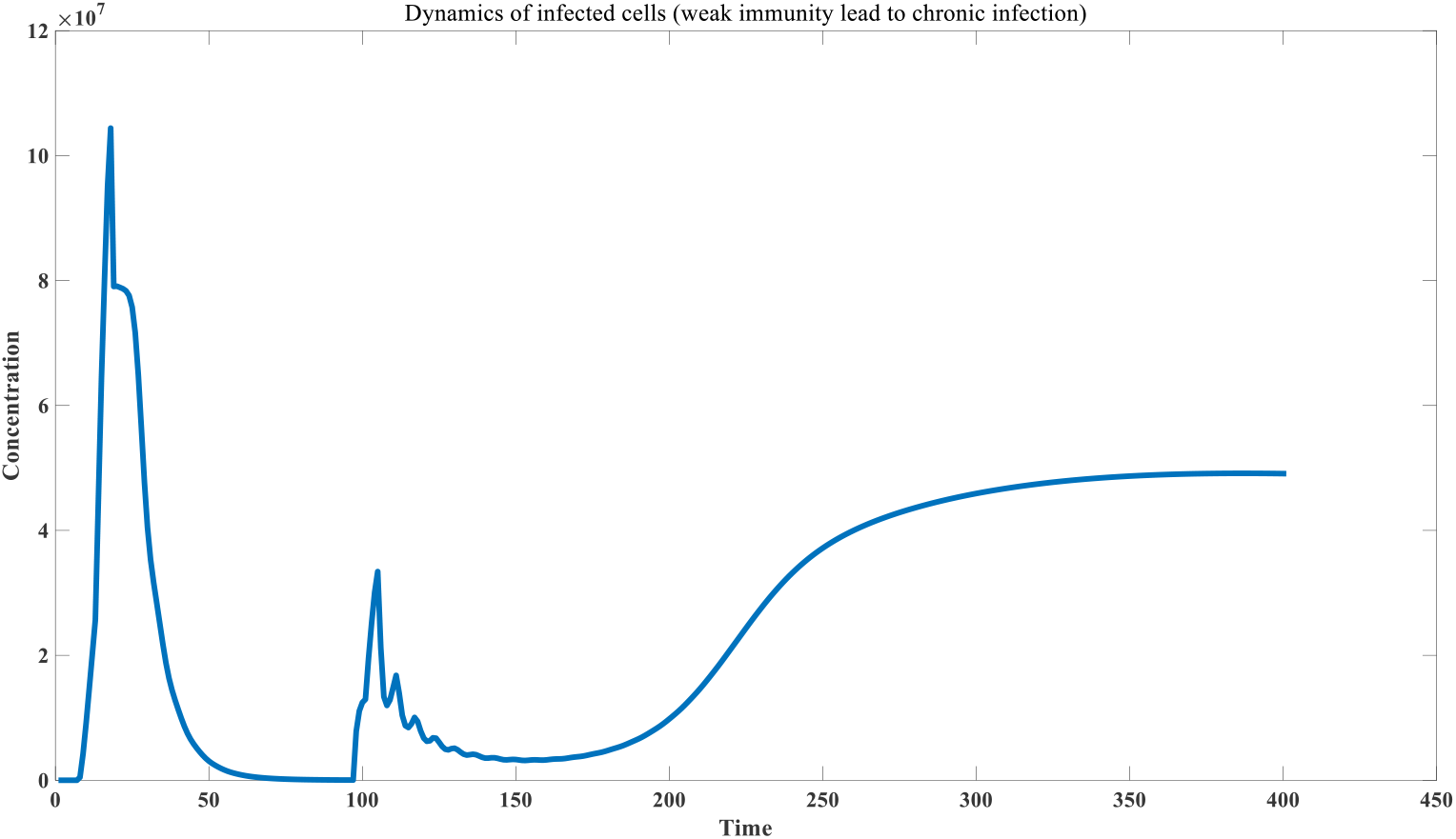
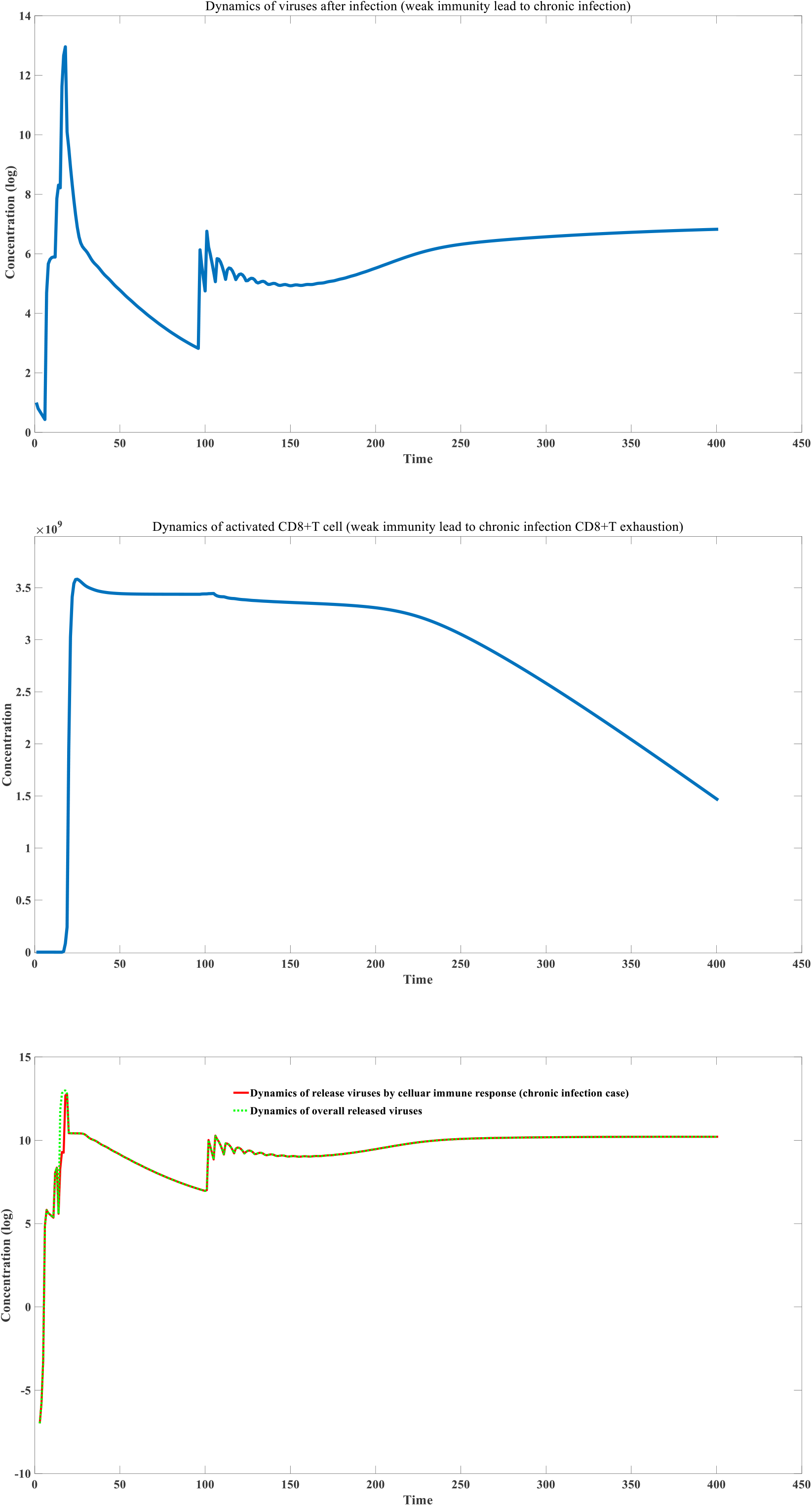
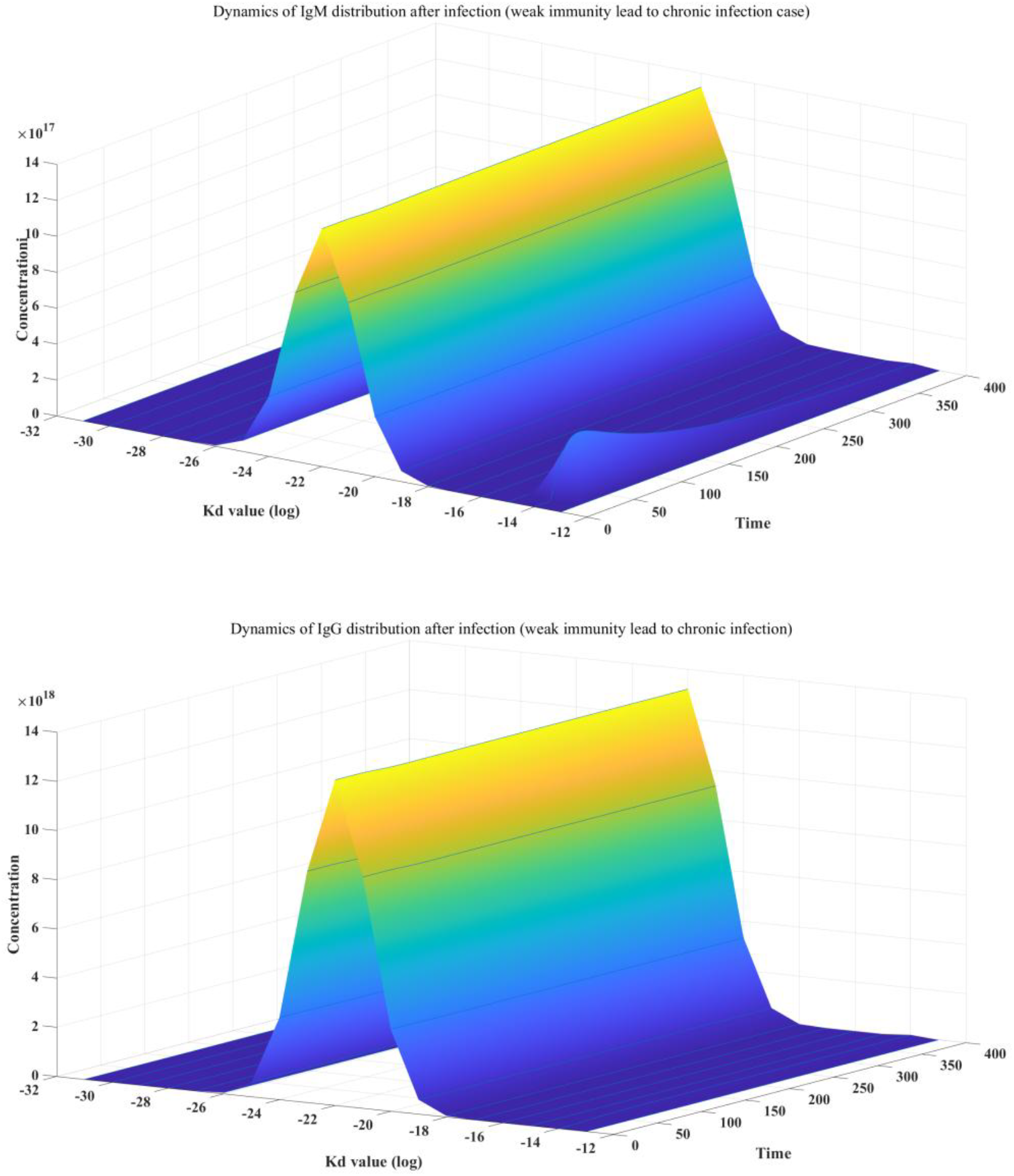
Virus-host interaction in chronic infection (chronic infection due to weak immunity)

Another common scenario can also lead to the occurrence of chronic infection. Clinical data show that secondary infection caused by mutant strains has a greater probability of causing chronic infection or leading to a longer infection cycle ^[64]^. Due to the existence of immune imprinting, the antibody distribution map against mutant strains has a higher proportion in the high affinity range than individuals who are infected with the virus for the first time, which leads to the initial antibody state being better than that of the naive group, which will lead to good inhibition of early viral reproduction and the peak concentration of the virus may be lower than that of the naive group. However, this also brings a new risk, that is, slower viral proliferation will weaker stimulate the proliferation of antibodies, resulting in a slow increase in antibody levels, which can easily fall into the threshold range of chronic infection. As can be seen from Figure 4D, secondary infection caused by mutant strains often does not change significantly in IgM, and the proliferation of specific antibodies is mainly driven by IgG.

**Figure 4D:**
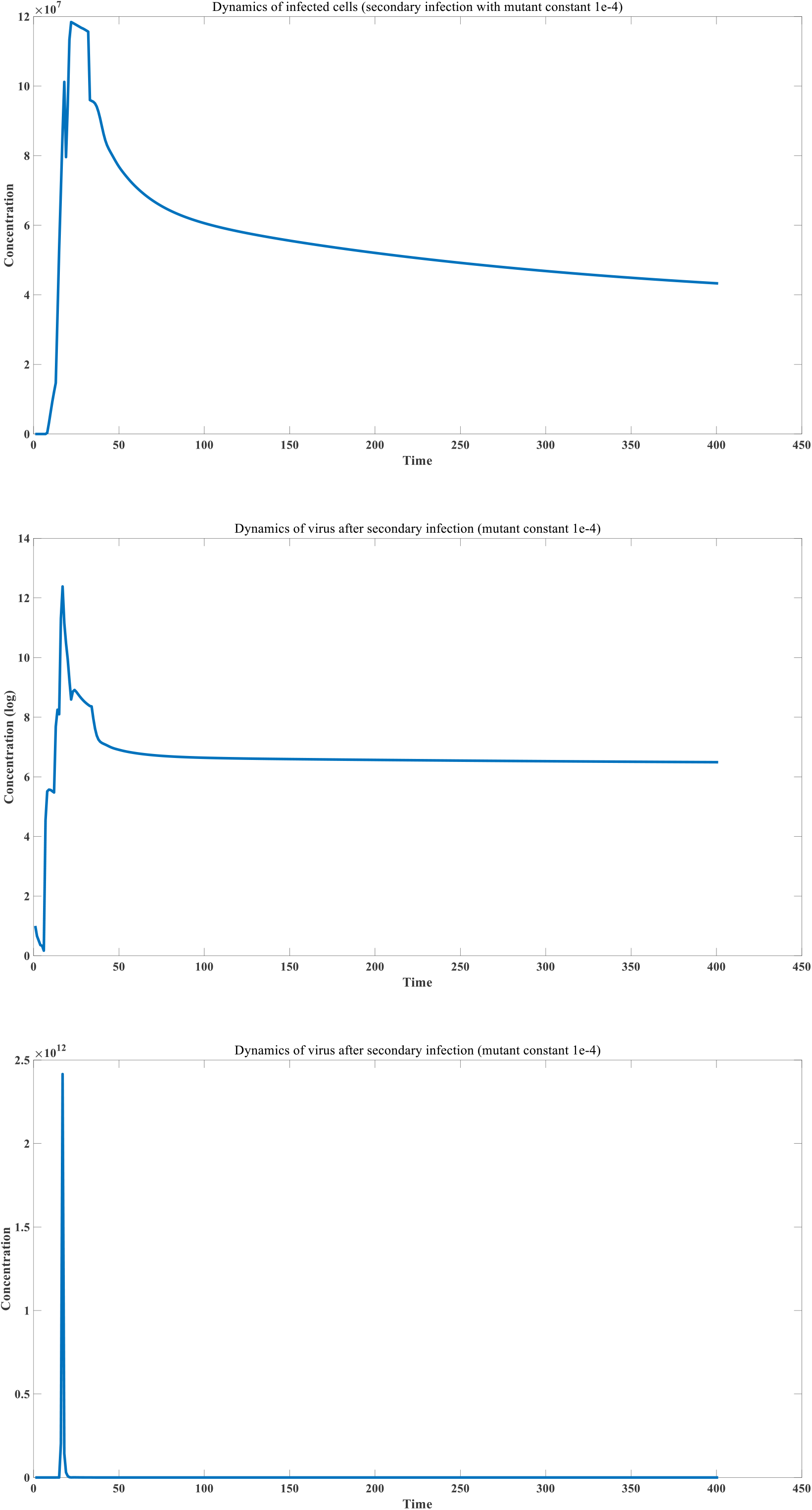
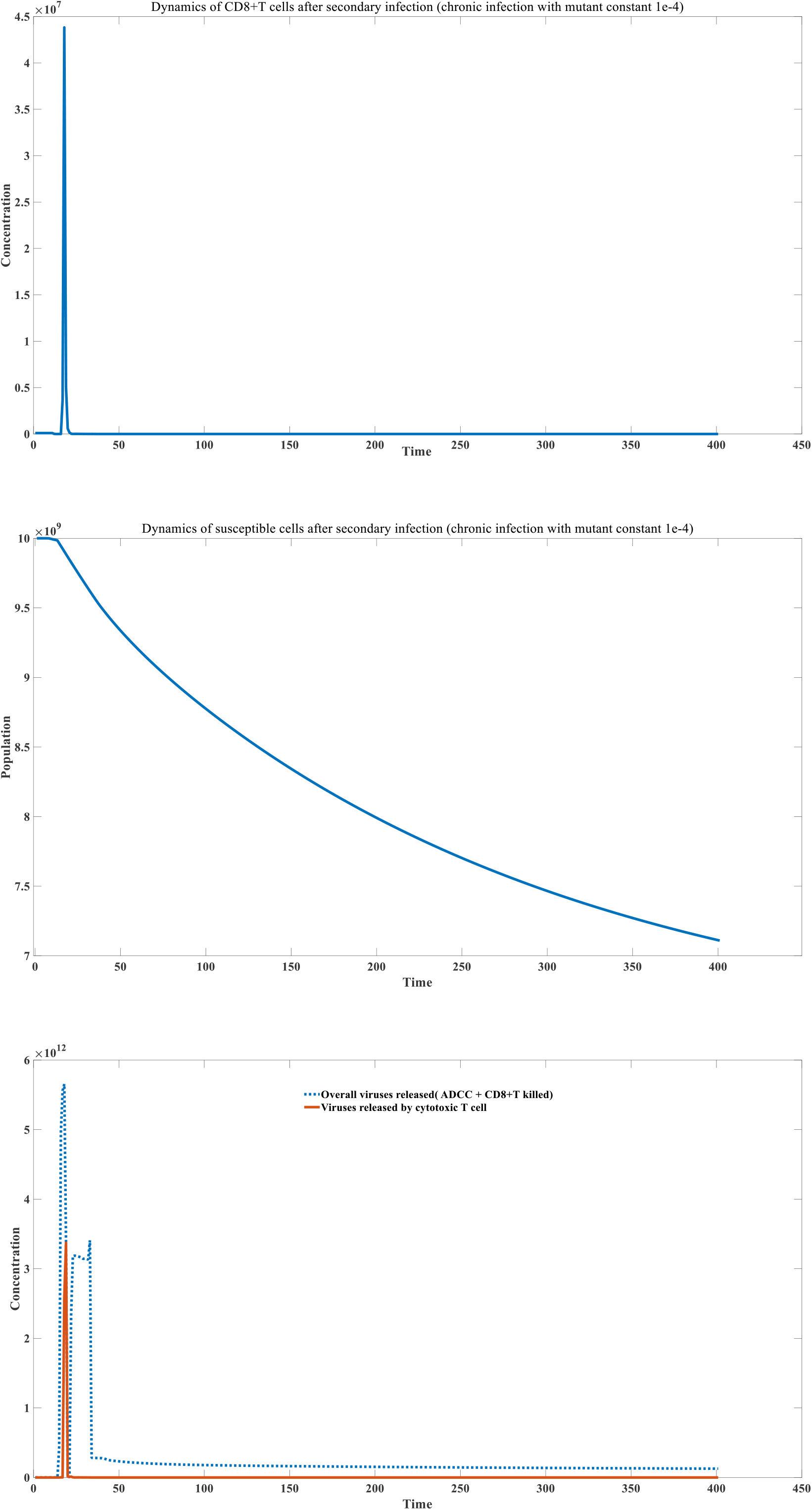
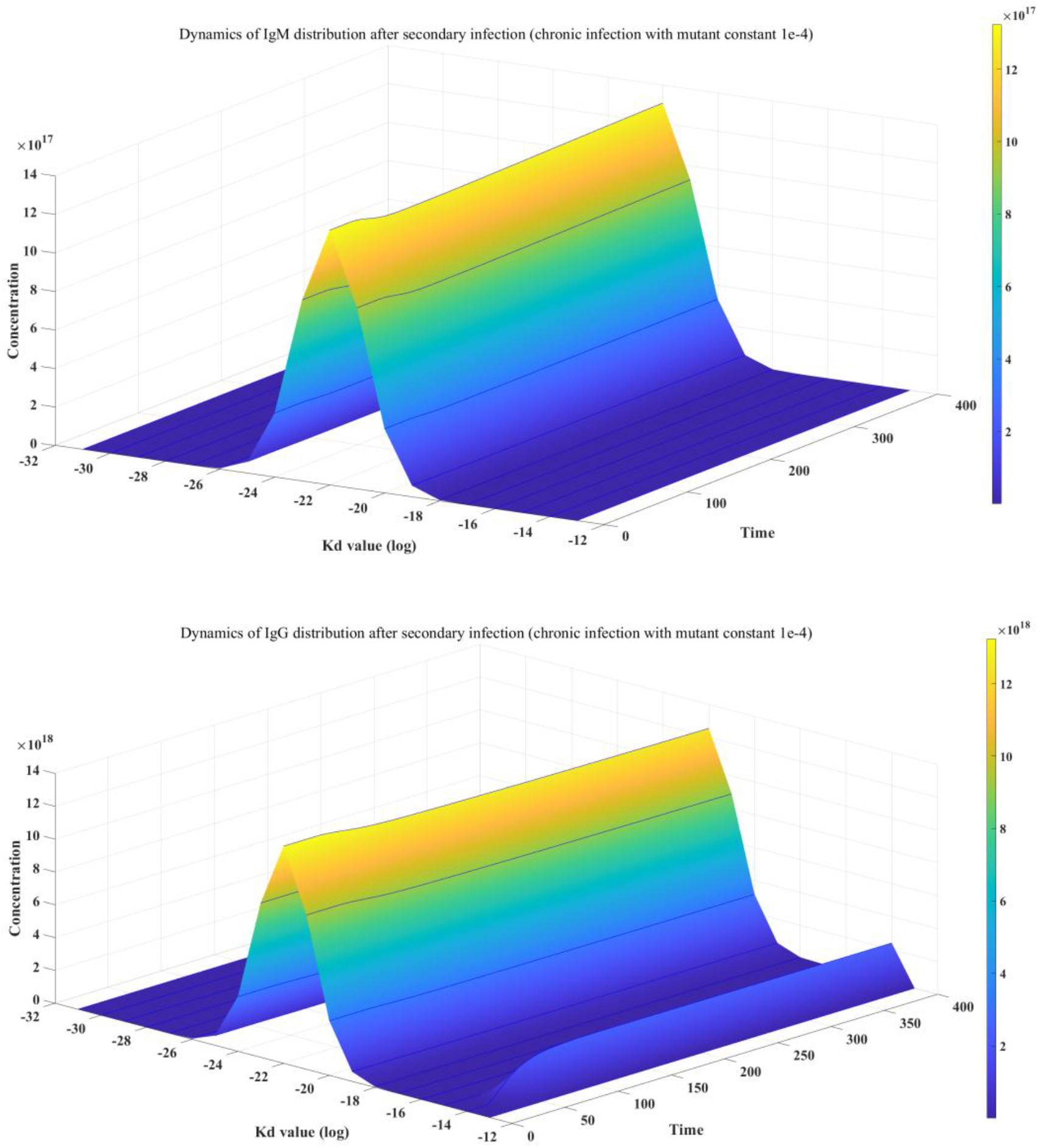
secondary infection (infection caused by mutant strain) tends to be chronic

We further investigated the impact of viral properties on chronic infection. Our simulation results show that viruses with longer replication cycles are more likely to cause chronic infection; the faster a virus replicates, the less likely it is to develop chronic infection. Figure 4E depicts infections caused by three viruses with different replication rates. As can be seen, viruses with slower replication rates are more likely to cause chronic infection. Real-world data supports our simulation results. Rapidly replicating viruses, such as influenza and COVID-19, are less likely to cause chronic infection, while viruses with longer replication cycles, such as HBV and HCV, are more likely to do so. Our model also demonstrates the feasibility of a kinetic explanation for the development of chronic infection. Viruses in the latent state may simply be early-stage viruses that have just invaded newly infected cells. The mechanism by which viruses with slow replication cycles cause chronic infection is as follows: Slowly replicating viruses cannot rapidly activate humoral and cellular immunity, resulting in levels of specific antibodies and specific CD8+ T cells falling within the threshold for chronic infection. Small molecule drugs for chronic viral infections, such as reverse transcriptase inhibitors, can alleviate infection symptoms ^[65–66]^. For acute viruses, small molecule drugs that inhibit viral replication, such as paxlovid, can also control the peak concentration of the virus and play a role in preventing severe illness ^[67]^. However, these small molecule inhibitors cannot fundamentally eliminate chronic infections.

**Figure 4E:**
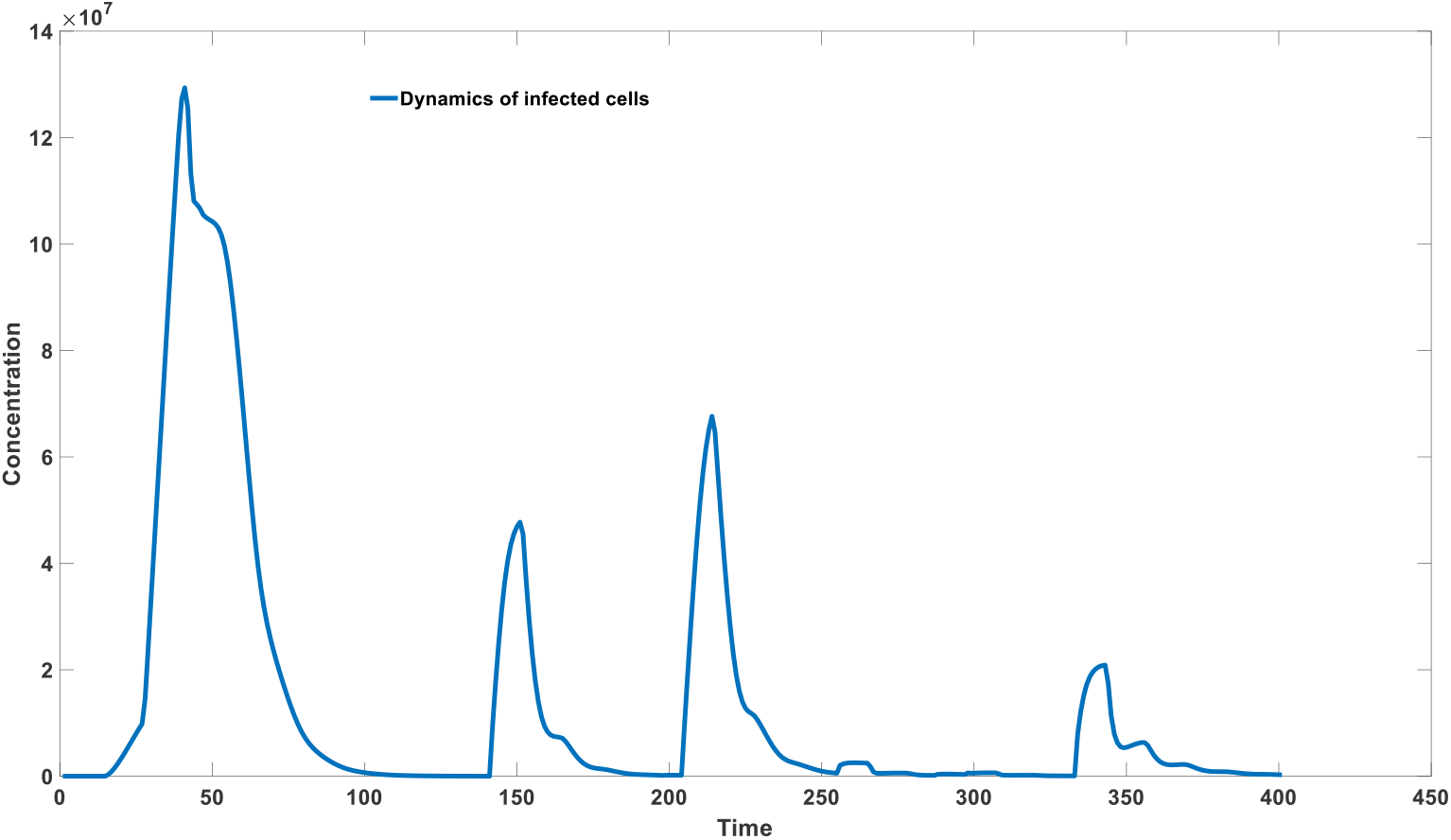
slow replication tends to form chronic infection (k_0_= 1)

After studying the causes of chronic infection, we further simulated the treatment of chronic infection. Here we mainly analyzed the therapeutic effects of three methods: monoclonal antibody therapy, increasing the concentration of environmental antigens, and therapeutic vaccine therapy. Monoclonal antibodies are widely used in the treatment of acute infections. In recent years, they have also been gradually used in the treatment of chronic infections, and have achieved very good results in the treatment of HIV ^[68–70]^. We simulated the effect of specific neutralizing antibodies on chronic infection during the chronic infection stage. As can be seen from Figure 4F, specific neutralizing antibody therapy can help overcome chronic infection, but it depends on the dosage. When the dosage is insufficient, it can only alleviate the symptoms in the short term and cannot fundamentally eliminate the chronic infection. However, a major problem with monoclonal antibody treatment of chronic infection is the high cost, because the manufacturing cost of monoclonal antibodies is much higher than the manufacturing cost of antigens. In addition, the required dosage is large. The dosage to eliminate chronic infection may be much higher than the dosage to alleviate the symptoms of acute infection, which greatly limits the feasibility of this method.

**Figure 4F:**
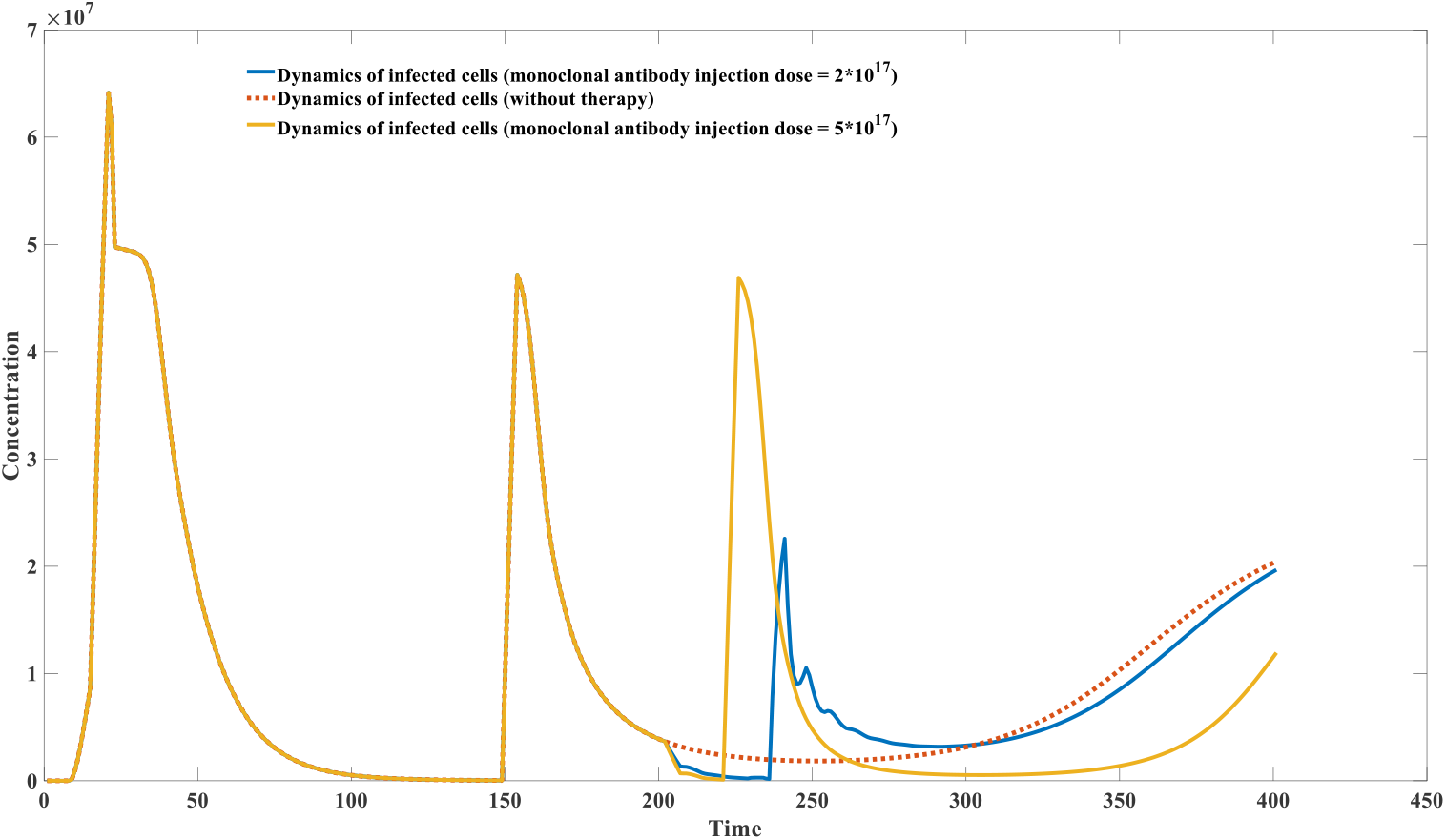
Effect of neutralizing antibody therapy in overcoming chronic infection

We also simulated another method that can increase antibody levels. This method is actually very old. There are cupping and scraping methods in Chinese traditional therapy, but the mechanism of this method lacks scientific research. Experiments have shown that scraping therapy can significantly increase the antibody level of mice after vaccination ^[71]^. Like those methods of taking peptidoglycan, we believe that the principle of scraping and cupping therapy is to increase the level of environmental antigens. They both induce a large number of cell lysis through physical methods, thereby leading to an increase in the concentration of self-antigen substances in the body fluid environment. High environmental antigens can increase the level of total antibodies in the short term. As shown in Figure 4G, by increasing the concentration of environmental antigen substances, all antibody levels (including IgM and IgG) will be improved, which helps to enhance the ADCC effect and the killing efficiency of CD8+T cells, and thus can play a positive role in reducing the number of cells in chronic infection. However, this approach is limited in its effectiveness, as the antibody boost isn’t concentrated in the high-binding region but rather uniformly increases all antibodies. As shown in Figure 4G, a short period of environmental antigen augmentation therapy failed to fundamentally reverse chronic infection; after cessation of environmental antigen administration, the number of infected cells reappeared. Completely reversing chronic infection requires the use of larger doses of autoantigens, which significantly increase overall antibody concentrations and, in practice, carries significant risk of side effects. Nevertheless, in certain specific cases, increasing the concentration of autoantigens can indeed accelerate the resolution of chronic infection. In the long term, the steady state of many chronic infections is not a continuous process, but rather a relatively stable plateau. During this plateau, as infected cells continue to lyse, antibody levels and distribution in the body undergo subtle changes. Ultimately, the body overcomes this chronic infection phase and achieves complete viral clearance. This stage of chronic infection is mathematically equivalent to a saddle point, which is not globally stable. Increasing the concentration of self-antigens can accelerate the process of equilibrium escaping from this saddle point, ultimately reaching a globally stable point of complete viral clearance. Many traditional therapies have demonstrated efficacy in accelerating the resolution of chronic infections.

Finally, we simulated the most critical treatment method, which is therapeutic vaccine therapy. Currently, therapeutic vaccines have achieved good results in the treatment of chronic viruses such as HBV, HCV and HPV ^[72–74]^. According to our model, the mechanism of therapeutic vaccine is that when the body enters the plateau phase of chronic infection, although the number of infected cells remains high, the virus concentration in the body is already at a low point. The lack of viral antigens cannot strongly induce the rapid proliferation of specific antibodies, thus forming a plateau phase. At this time, artificial injection of inactivated viral antigens can greatly stimulate the proliferation of antibodies with high binding activity, thereby breaking the plateau phase of chronic infection. Our simulation discovered an interesting and very important phenomenon, that is, unlike preventive vaccines, the efficacy of therapeutic vaccines shows a very strong positive correlation with dose. For preventive vaccines, regardless of the dose, the vaccine always plays a positive protective role in preventing infection. Of course, large doses can bring stronger protective effects. However, it is different for therapeutic vaccines. When the dose is low, therapeutic vaccines will aggravate chronic infection. This is because a small amount of antibody injection will consume the antibodies in the body. At the same time, due to the small dose, the BCR cannot effectively compete with the free antibodies to obtain antigenic substances, so the antibody regeneration function cannot be achieved, causing the effective antibodies in the body fluids to decrease, thereby aggravating the infection, which is also shown in Figure 4H. Moreover, for therapeutic vaccines, the dose required is usually much higher than that of preventive vaccines. Figure 4H also shows that when the dose is insufficient, it is difficult for therapeutic vaccines to fundamentally eliminate chronic infections, although it may bring about symptom relief in the short term. It can also be seen from Figure 4H that when the therapeutic vaccine dose is insufficient, the secondary antibody proliferation induced by vaccination is not significant, and it is impossible to quickly get rid of the plateau period of chronic infection. Compared to the first two approaches, therapeutic vaccines are the most viable option for combating common chronic infections. This is because the cost of preparing antigens is much lower than that of monoclonal antibodies. Furthermore, while the dosage of therapeutic vaccines is significantly higher than that of preventive vaccines, it is still significantly lower than that of monoclonal antibodies, and even lower than the dose of autoantigens.

**Figure 4G:**
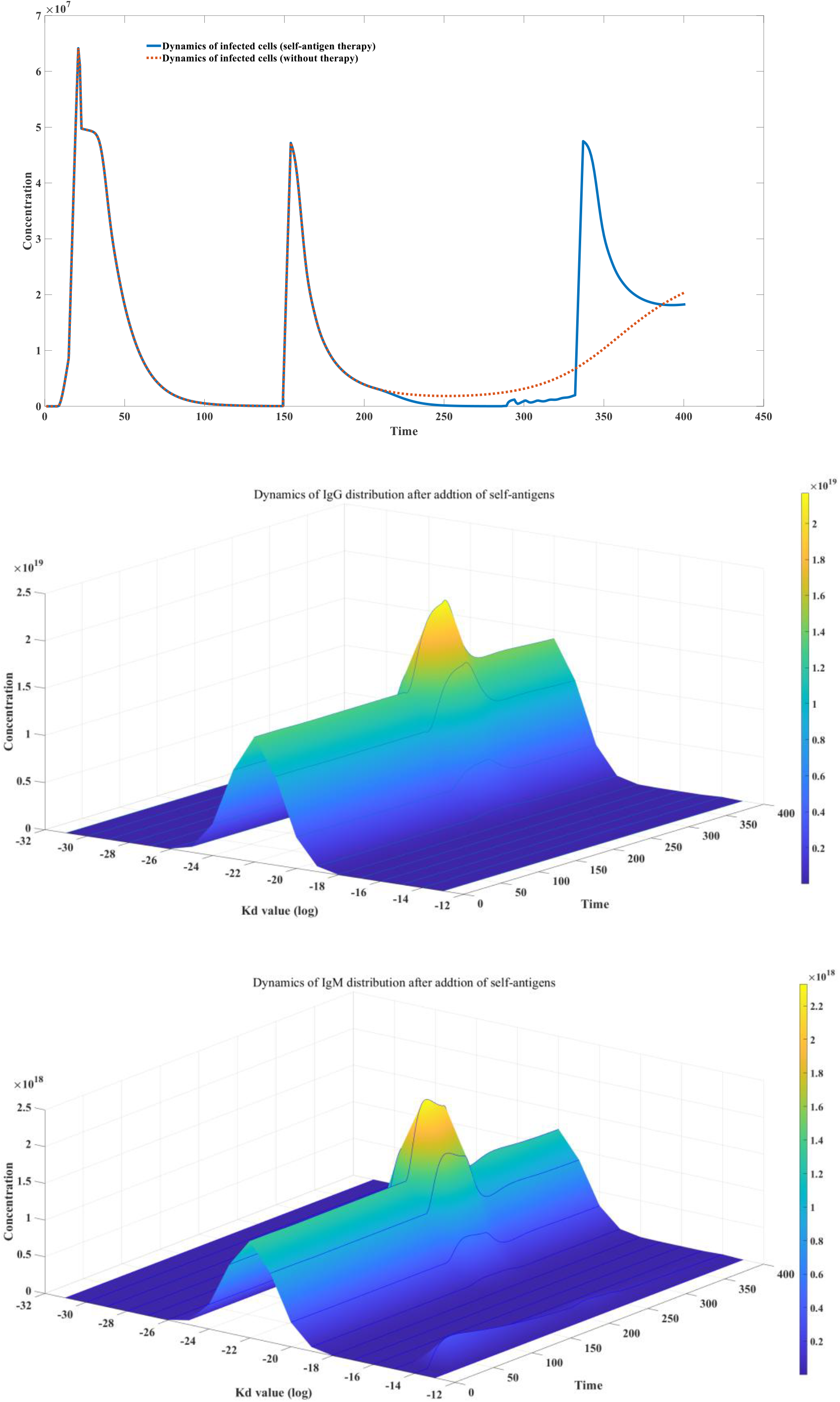
Effect of Self-antigen therapy in overcoming chronic infection

**Figure 4H:**
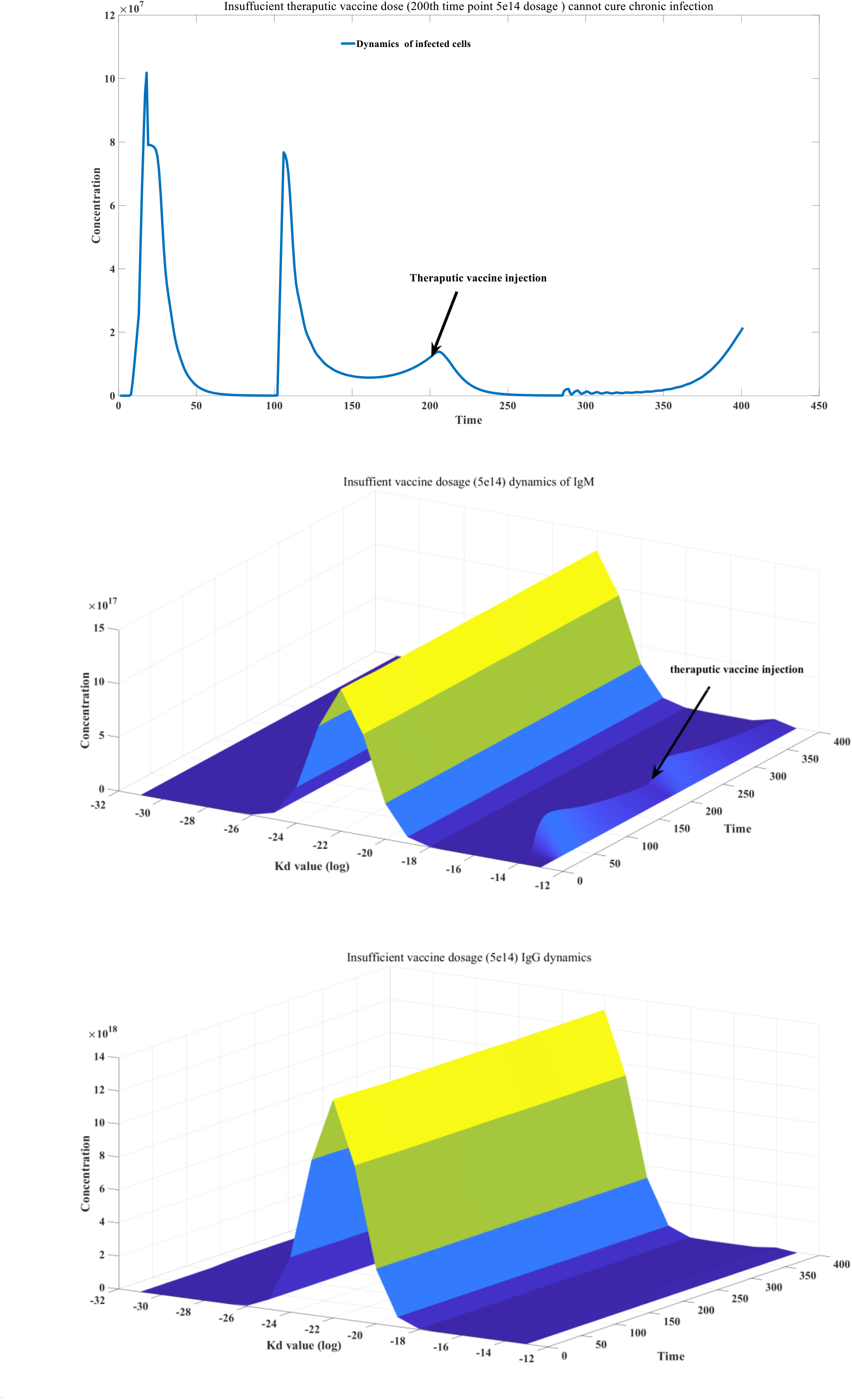
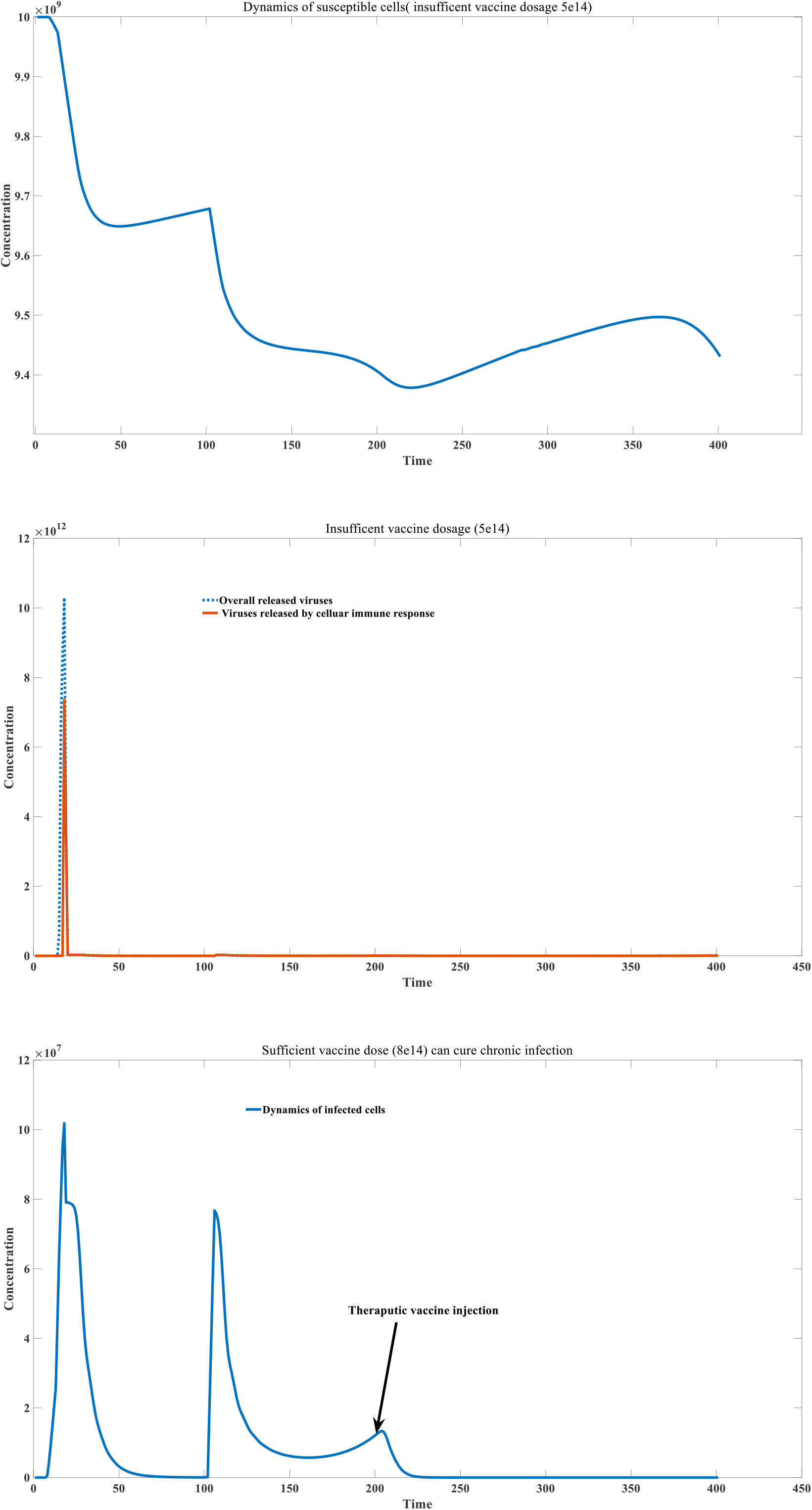
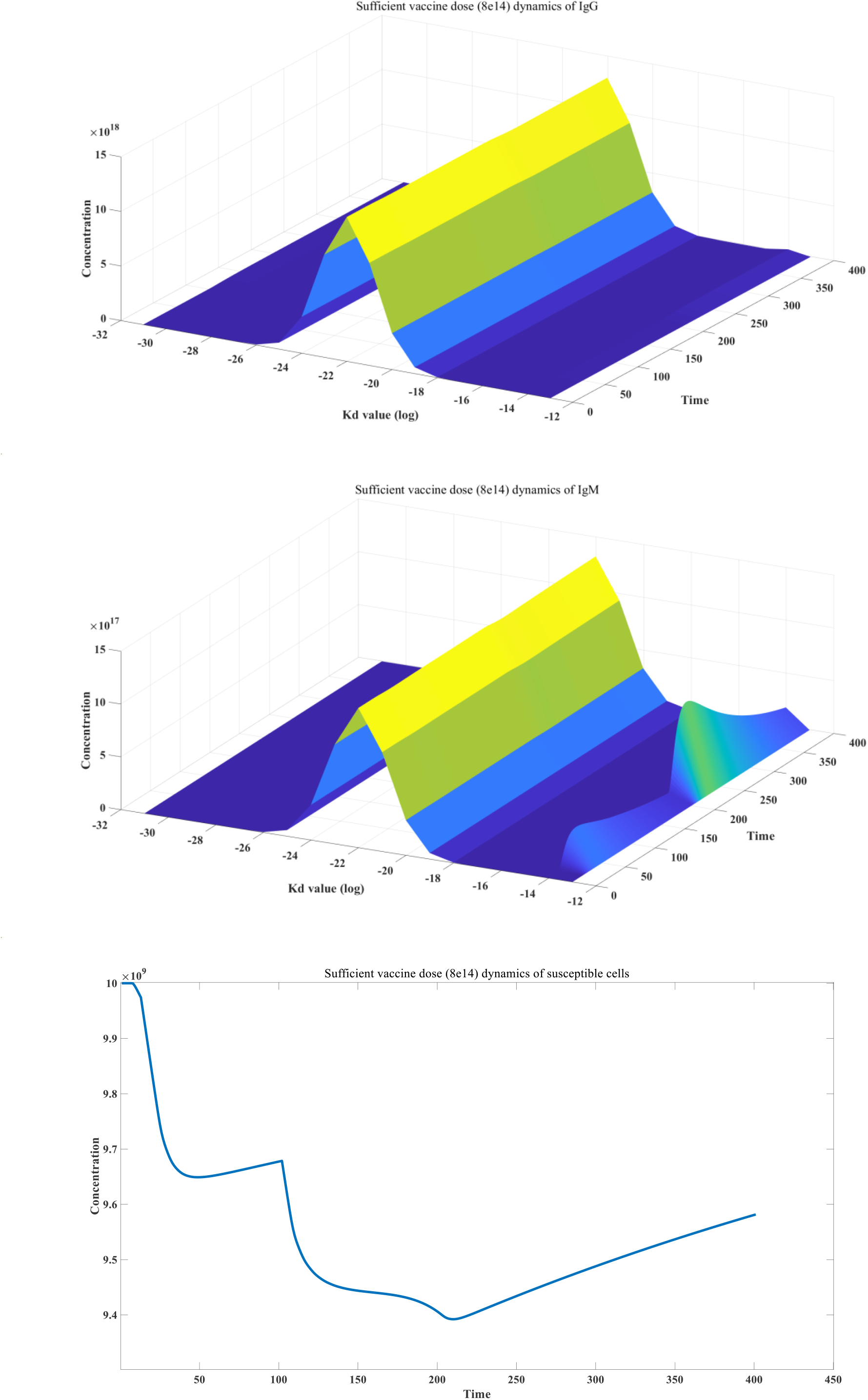
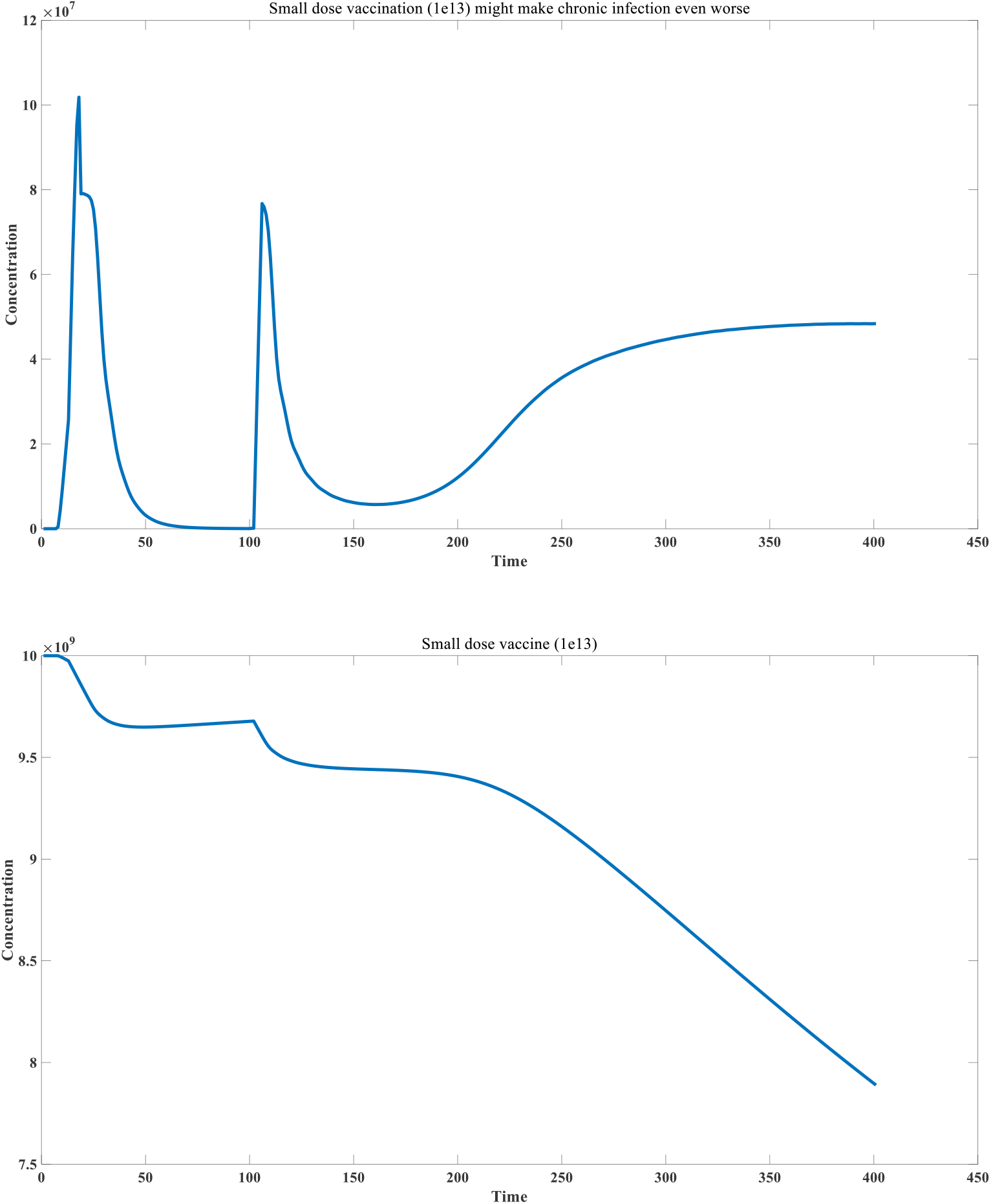
Effect of therapeutic vaccine in overcoming chronic infection

### 2.2 Establishment of cancer immunity model

Mathematical analysis of the cancer immunity model is provided in the supplementary material. In the fifth section, we established a new model for cancer-immune interaction. Although virus-immune system interaction models have been widely studied, cancer-immune system interaction models have been relatively less studied. We believe that the immune system recognizes cancer cells primarily through neoantigens. The traditional view is that neoantigens are mainly caused by gene mutations. Recent studies tend to believe that proteins with significantly increased expression in cancer cells (such as human endogenous retroviruses) ^[75–76]^ or proteins with large-scale sequence changes due to variable splicing ^[77–78]^ are high-quality neoantigens. This is caused by the recognition mechanism of the immune system. As we described in cellular immunity, cell lysis is not only dependent on the properties of the antigen, but is also positively correlated with the concentration of the internal antigen. Many antigens that are low in normal cells but highly expressed in cancer cells can become good neoantigens. Conventional point mutations, resulting in SNPs, cannot be converted into highly divergent peptide sequences after protease degradation. This is similar to the memory mechanism of CD4+ T cells. Therefore, they do not lead to selective recognition of cancer cells by CD8+ T cells or the selective production of cancer-specific antibodies by CD4+ T cells. However, mutations caused by alternative splicing are different. They can result in large-scale insertions of new proteins or frameshift mutations, which produce primary peptide sequences that are completely different from the wild-type. These novel MHC-I and MHC-II antigens activate the immune system to recognize and kill cancer cells. Cancer-immune system interaction models are best represented using ODEs because antigen concentrations within cancer cells tend to be stable, unlike viral infection models. Similar to agent-based models of viral infection, in Model 3.5.1, we assume that immune recognition of cancer cells relies on two effectors: ADCC and CD8+ T cell-mediated cytotoxicity. CD8+ T cells also have an exhaustion effect, and their activation and proliferation are closely related to the concentration of antibody-antigen complexes. Therefore, our model also explains the huge role of B cells in cancer immunity.

Based on Model 3.5.1, we simulated the mechanism of cancer development. Our model assumes that cancer cells are constantly present in the body, and the development of cancer depends on two factors: the proliferation capacity of cancer cells and the surveillance capacity of the immune system. In the supplementary materials, we use mathematical analysis to study the cancer-immune system interaction model. For most combinations of cancer cell proliferation capacity and immune surveillance capacity that are within a reasonable range, the cancer cell-immune system can reach an equilibrium state, in which cancer cells can persist stably at a low concentration. Therefore, the key question is whether the system undergoes significant fluctuations before reaching this equilibrium state, that is, whether cancer cells can proliferate significantly. If this proliferation is not significant, the body will not show symptoms of cancer. If it is significant, the cancer will be curable. If it is very significant, the cancer will be fatal. Different cancer treatments are not intended to change the final equilibrium state, but rather to mitigate the extent of cancer cell proliferation before reaching this equilibrium. Therefore, our model also supports the positive hypothesis that cancer can be completely cured because once the final equilibrium state is reached, cancer cells will remain at a low level for a long time through interaction with the immune system. However, due to the continuous decline in autoimmunity caused by aging and the continuous increase in the proliferation coefficient of cancer cells due to accumulated mutations, the risk of cancer continues to increase with age. Figure 5B illustrates the mechanism of cancer development. When immunity is strong or the cancer cell’s own proliferation coefficient is low, the cancer cells do not undergo significant proliferation to reach the final equilibrium state, which prevents us from showing obvious cancer symptoms. In our model, the strength of immunity is manifested in two aspects: the stimulation and regeneration coefficient of the antigen-antibody complex on the antibody and the stimulation and proliferation coefficient of the antigen-antibody complex on CD8+ T cells. These two aspects often change collectively. When immunity is weakened, cancer cells undergo a very significant proliferation process, thus showing cancer symptoms.

**Figure 5A:**
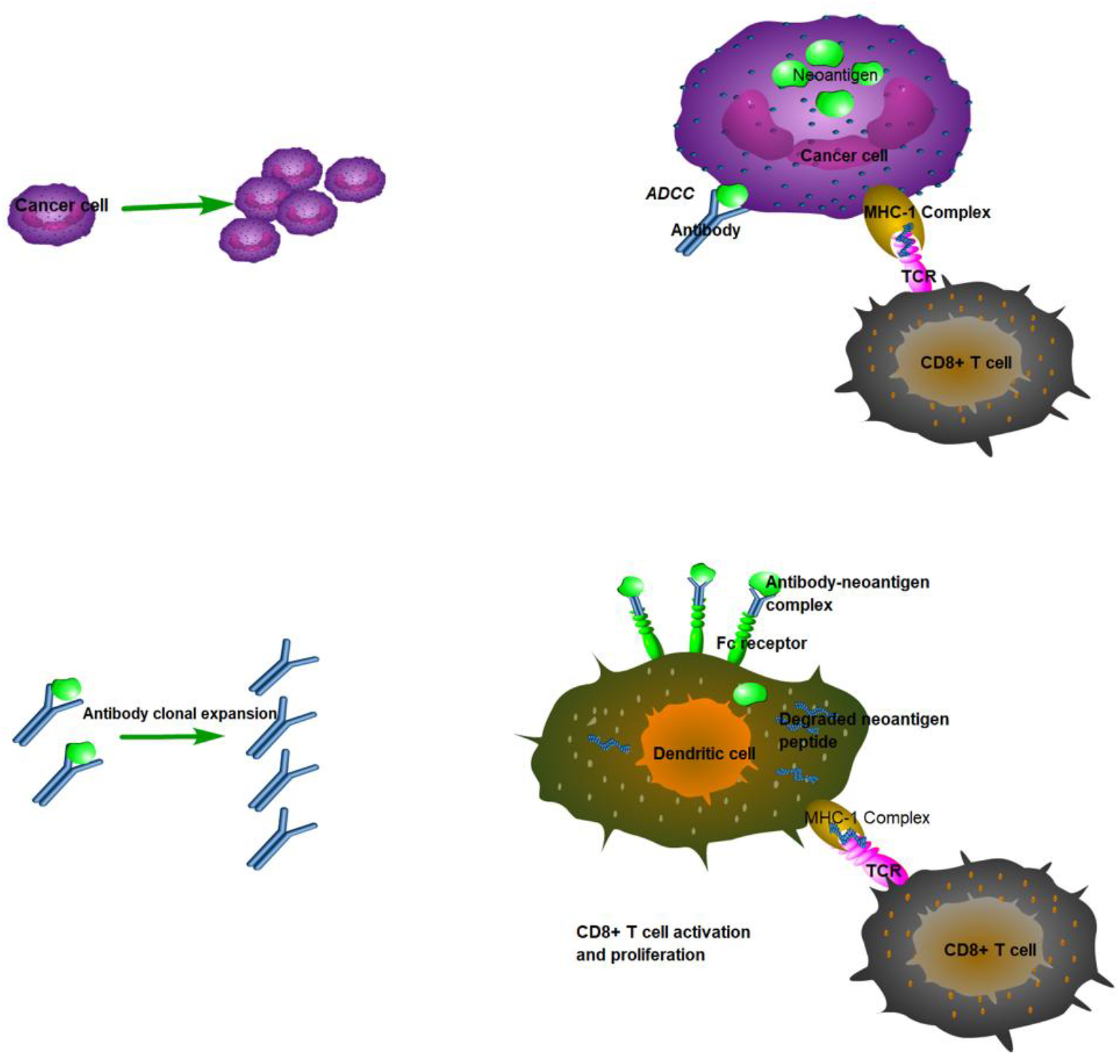
Cancer Immunity Model Flowchart

**Figure 5B:**
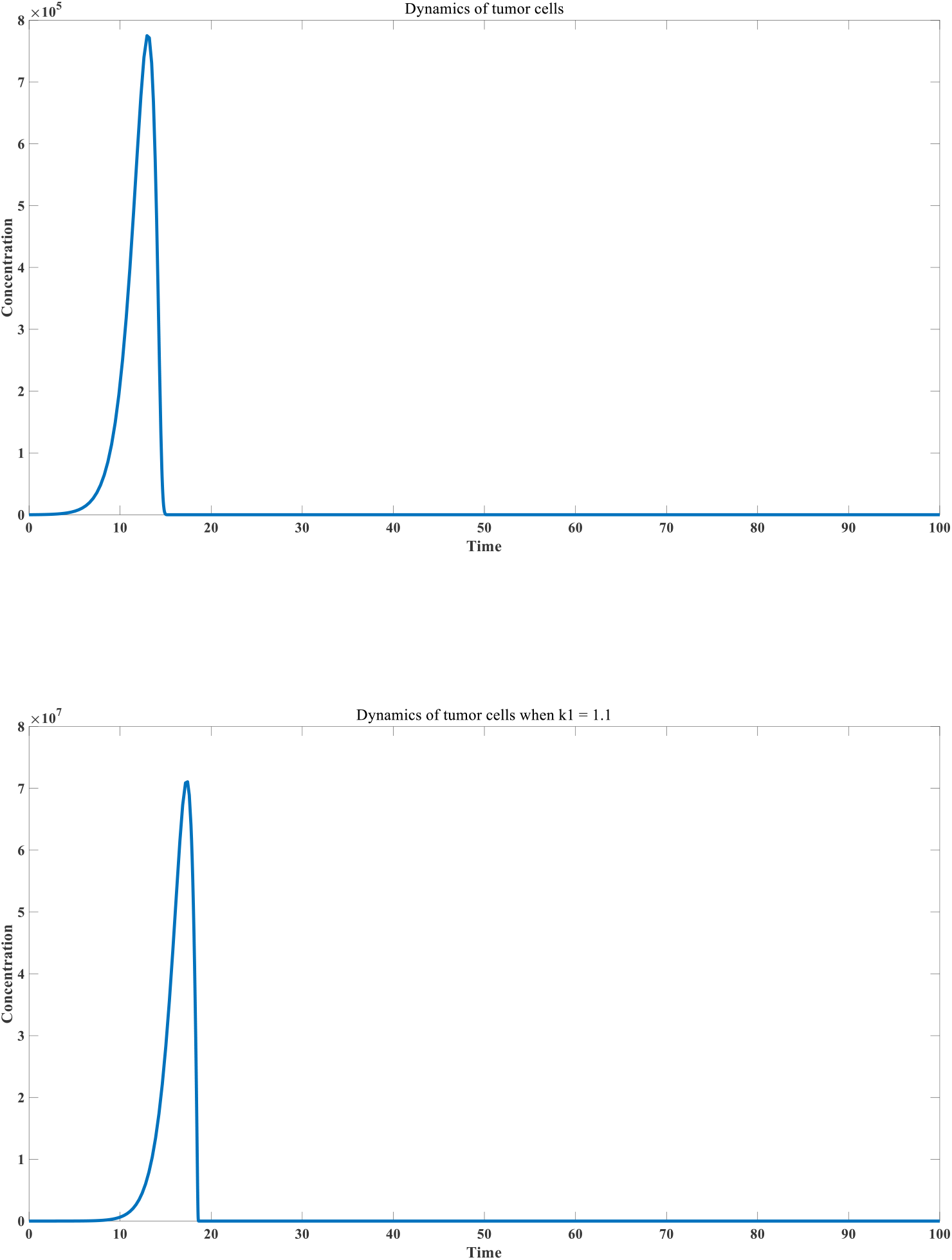
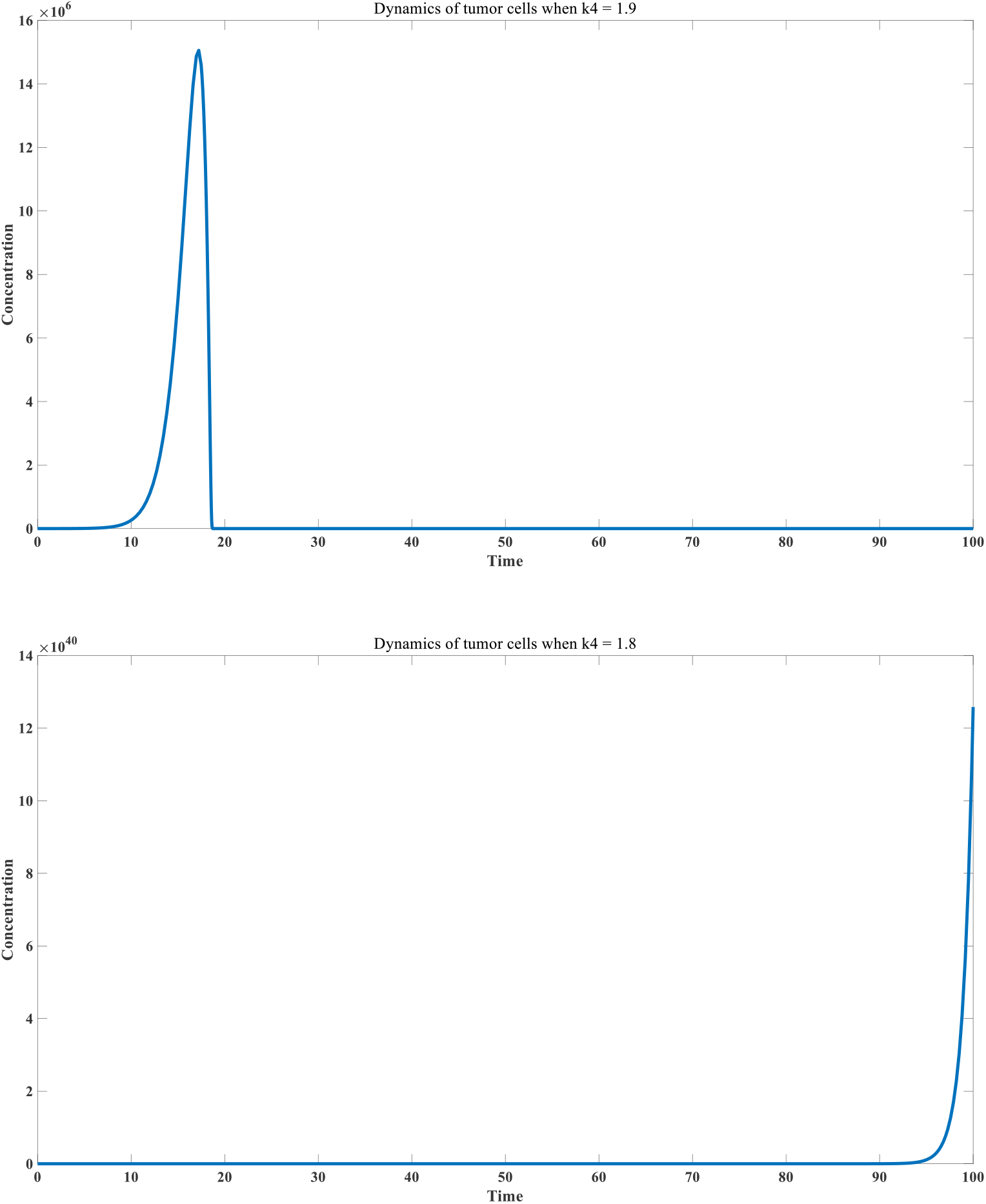
cancer induced immune response

We further simulated the effects of five commonly used immunotherapies on cancer cell proliferation: inhibitors that eliminate CD8+ T cell exhaustion (immune checkpoint inhibitors), monoclonal antibodies that specifically recognize neoantigens, B cell antigen-targeting immunotherapy, MHC-I antigen-targeting immunotherapy, and MHC-II antigen-targeting immunotherapy.

Since the discovery of PD-1 inhibitors, they have been widely used in cancer treatment and chronic infection treatment ^[79]^. Their specific mechanism of action is to reduce the expression of PD-1, thereby eliminating or alleviating the self-inactivation of CD8+ T cells during cell killing. In our model, *ρ*_2_the value is reduced, thereby reducing the loss of CD8+ T cells. As shown in Figure 5C, the addition of immune checkpoint inhibitors can significantly control the proliferation of cancer cells and achieve good therapeutic effects.

**Figure 5C:**
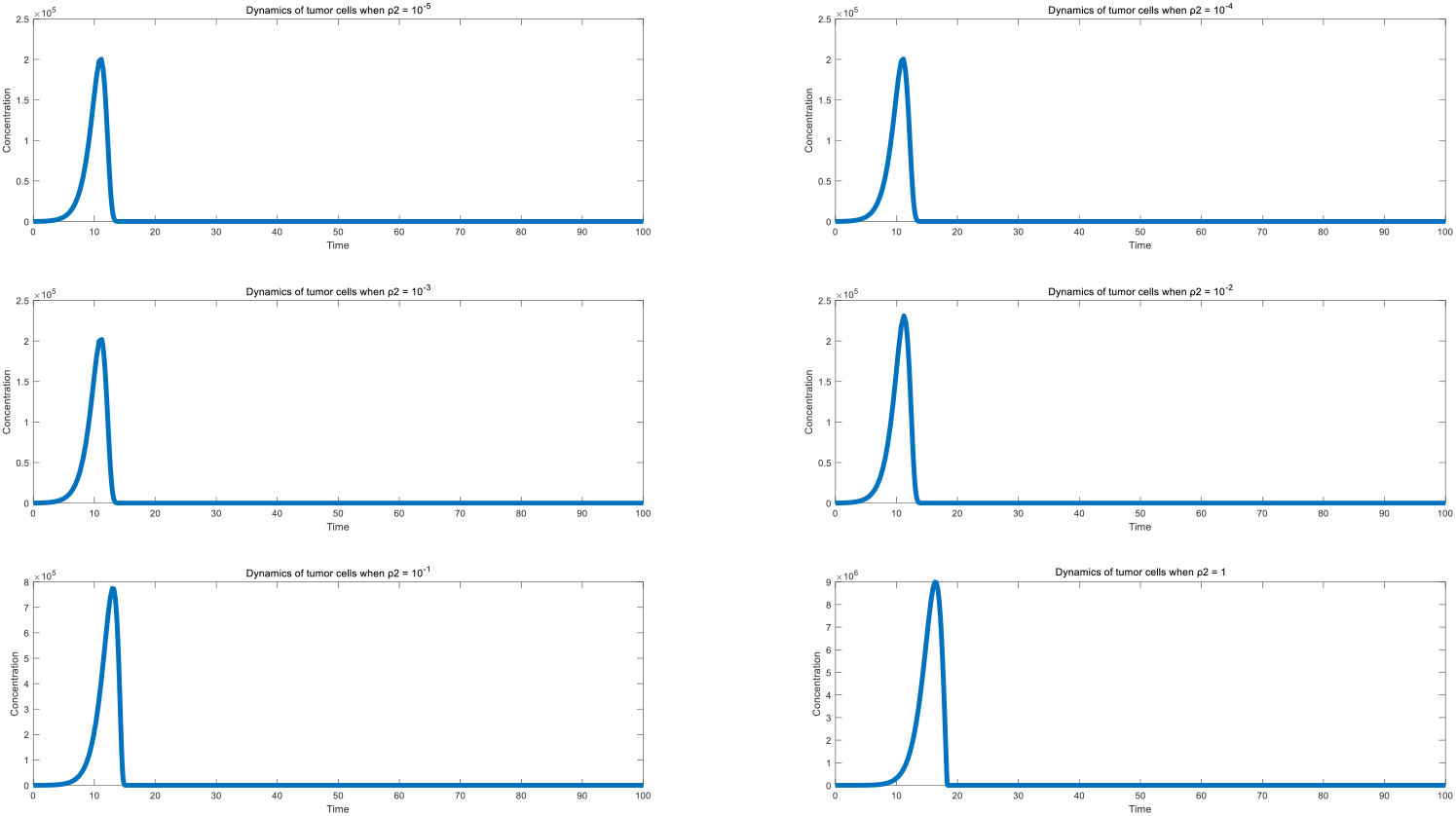
Effects of PD-1 inhibitor in cancer therapy

Monoclonal antibody therapy that specifically recognizes neoantigens has also achieved good results in cancer immunotherapy ^[80–81]^, especially when the neoantigen is very single and clear. Increasing the concentration of the antigen can directly enhance the ability of the ADCC effect to kill cancer cells, and can also indirectly generate a large number of antigen-antibody complexes to promote further enhancement of antibodies and rapid proliferation of CD8+ T cells. As shown in Figure 5D, the proliferation of cancer cells can be significantly inhibited after the addition of antibodies. However, the disadvantage of this method is that the cost of monoclonal antibodies is generally very high, and the premise is that the patient’s neoantigen is preferably very clearly expressed at a high level, and it is best to express a common neoantigen such as HERV. Otherwise, the preparation of antibodies in personalized treatment will cost a lot.

There has been a heated debate about the control of cancer cells by the addition of B cell antigens. This is because traditional radiotherapy and chemotherapy can lead to abnormal lysis of cancer cells, resulting in the release of a large amount of neoantigens into body fluids, and these neoantigens with complete protein structures can exist as B cell antigens. Some reports indicate that the use of B cell antigens (self-cancer cell lysates) can achieve a good inhibitory effect on certain cancers ^[82]^. Our simulation shows an interesting phenomenon, that is, the injection of different doses of B cell antigens at different times may have different effects on the development of cancer. In the early stage, moderate injection of B cell antigens can inhibit the development of cancer, but the injection of large amounts of B cell antigens in the middle and late stages may increase the proliferation of cancer cells. This is because the injection of large amounts of B cell antigens in the middle and late stages will consume a large amount of antibodies, thereby reducing the ADCC effect and leading to a decrease in the rate of cancer cell lysis. Figure 5E–F shows this process. This shows that the use of B cell antigen therapy is risky, so in recent years, research on cancer therapeutic vaccines has tended to favor MHC-I and MHC-II antigens.

**Figure 5D:**
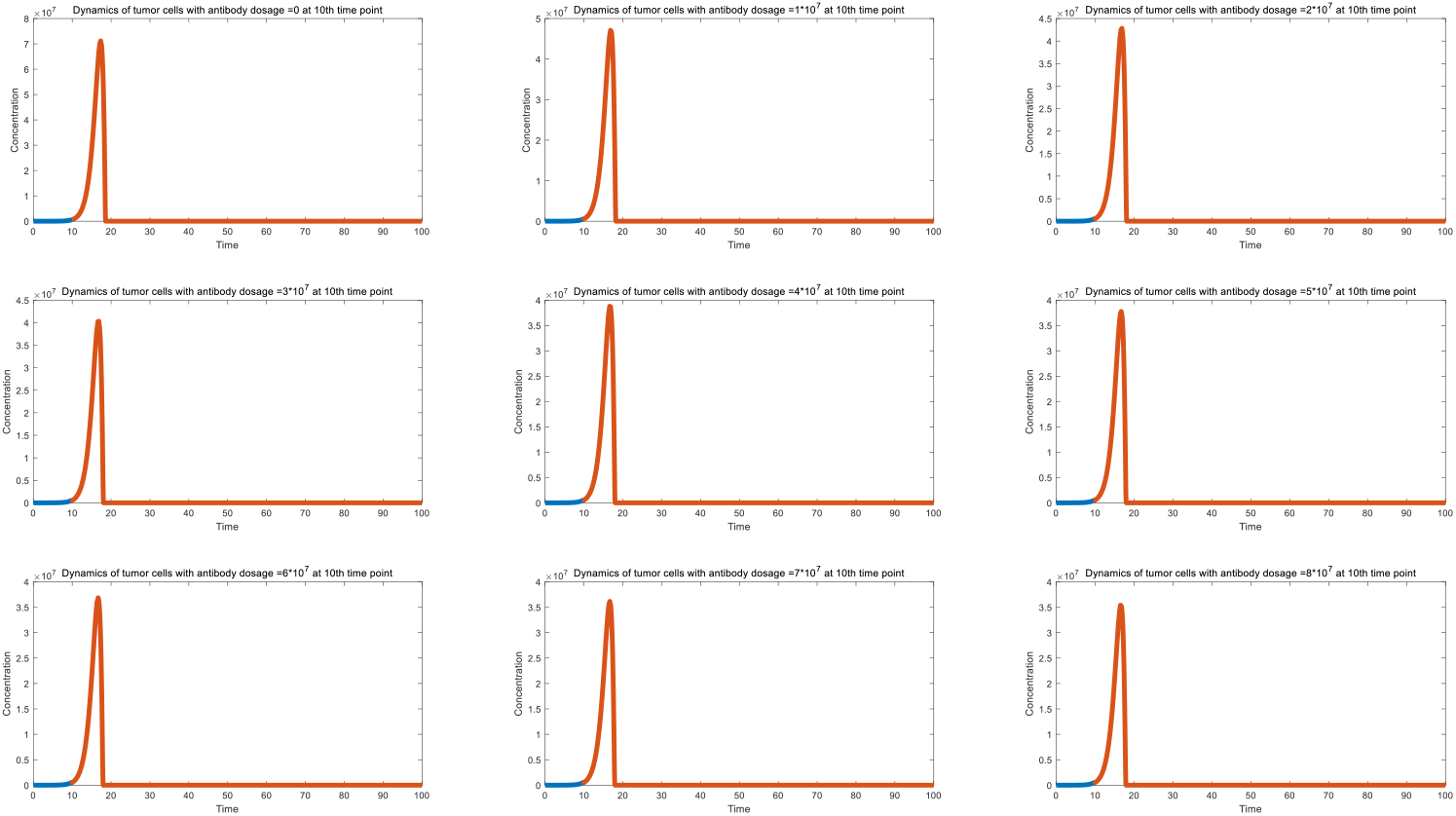
Effect of antibody therapy in cancer therapy

**Figure 5E:**
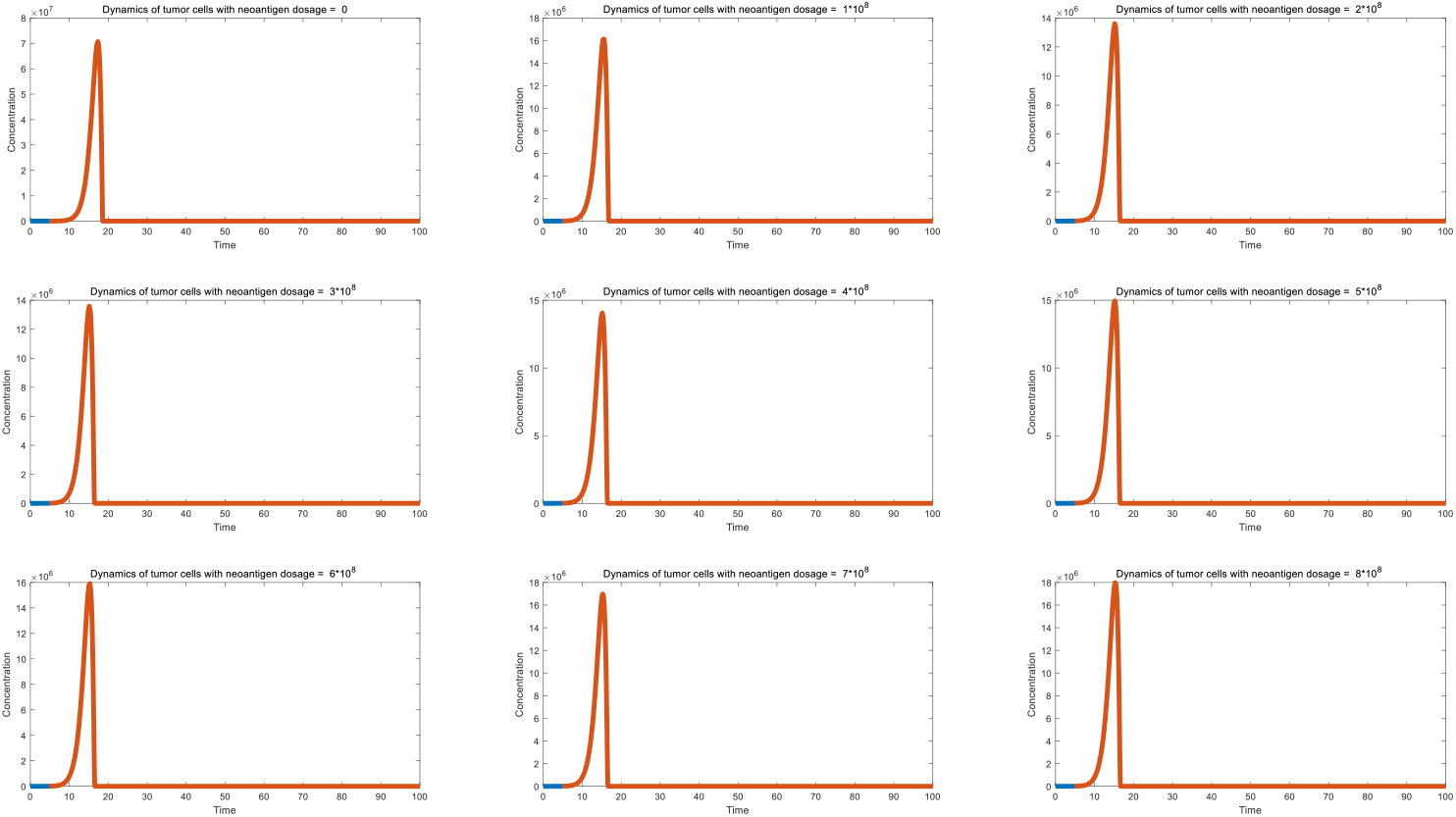
Effect of neoantigen addition on tumor control (vaccination at 5 ^th^time point)

**Figure 5F:**
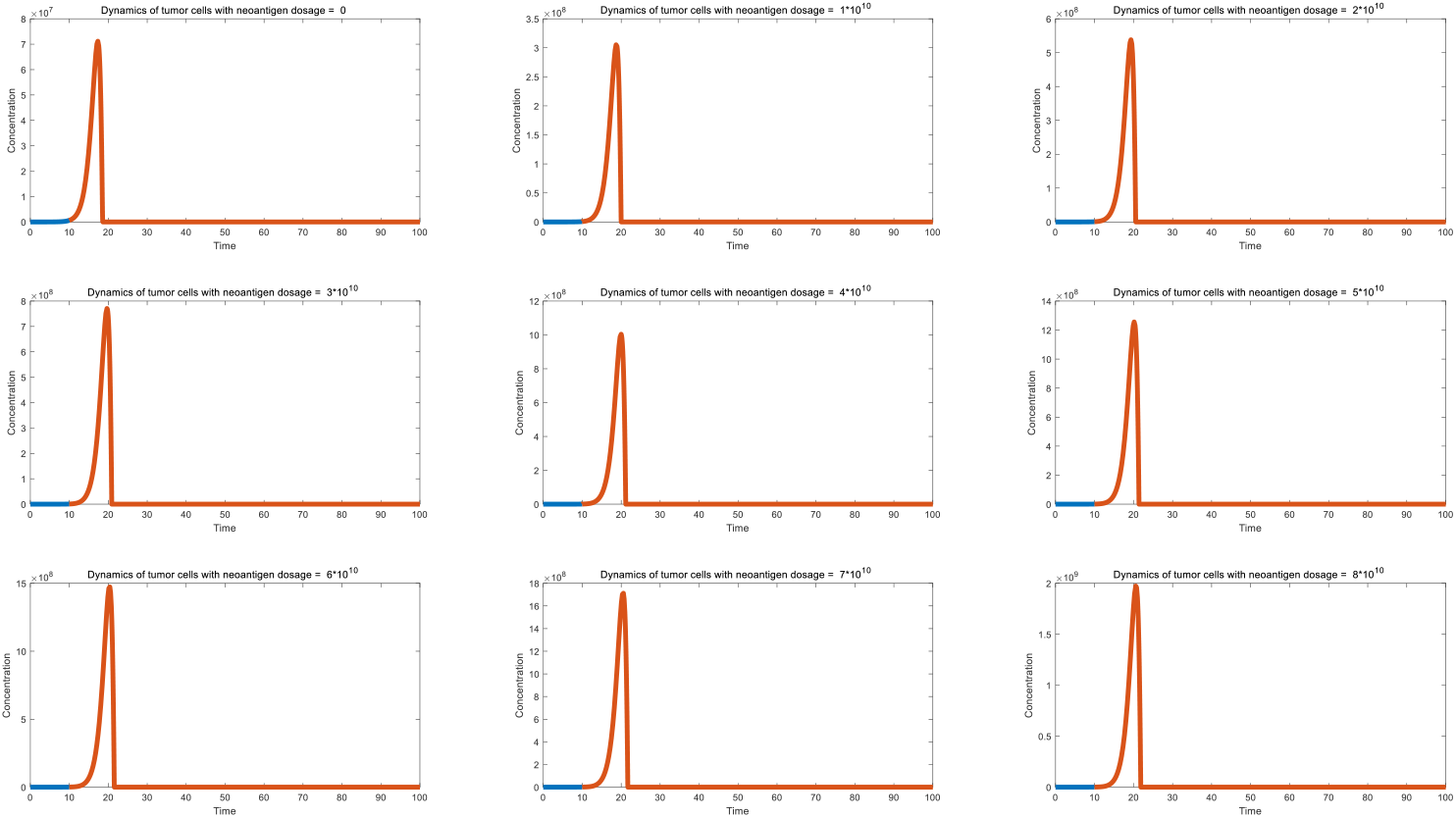
Effect of neoantigen addition on tumor control (vaccination at 10 ^th^time point)

The addition of MHC-I antigens can directly stimulate the ability of antigen-antibody complexes to feedback and generate CD8+ T cells, and has shown good effects in the treatment of many cancers ^[83–85]^. MHC-I antigens are polypeptide sequences that are derived from the cleavage of complete neoantigens and are polypeptide fragments that have strong binding activity with DC MHC-I complexes. Currently, many bioinformatics and artificial intelligence algorithms can effectively calculate the polypeptide sequences of MHC-I antigens. Figure 5G also shows the effect of adding MHC-I antigens on cancer cell control. The addition of MHC-I antigens can significantly increase the number of CD8+ T cells, thereby increasing the lysis of cancer cells mediated by them. The accelerated lysis of cancer cells can release more neoantigens, thereby accelerating the regeneration of antibodies and CD8+ T cells, playing a virtuous cycle. Like MHC-II antigens, MHC-I antigens are polypeptide fragments, so they will not directly bind to antibodies to cause antibody consumption, so they will not be infected with ADCC effects, and therefore they rarely bring negative effects.

In recent years, studies on MHC-II antigens have also shown that MHC-II antigen therapy has a good therapeutic effect on certain cancers ^[86–88]^. MHC-II antigens are polypeptide sequences that can bind to MHC-II complexes with high affinity after the original neoantigen is cleaved. Figure 5H shows that after the addition of MHC-II antigens, the proliferation of cancer cells can be significantly inhibited. This is because the addition of MHC-II antigens can greatly increase k_5_ value of specific antibodies without consuming antibodies, thereby greatly promoting the regeneration of specific antibodies, thereby enhancing the ADCC effect. More cancer cell lysis will play a virtuous cycle, leading to more antibody regeneration and CD8+ T cell regeneration. Therefore, MHC-II antigen therapy often achieves very good therapeutic effects. Like MHC-I antigens, they will not cause antibody exhaustion, so there are few reports of side effects.

**Figure 5G:**
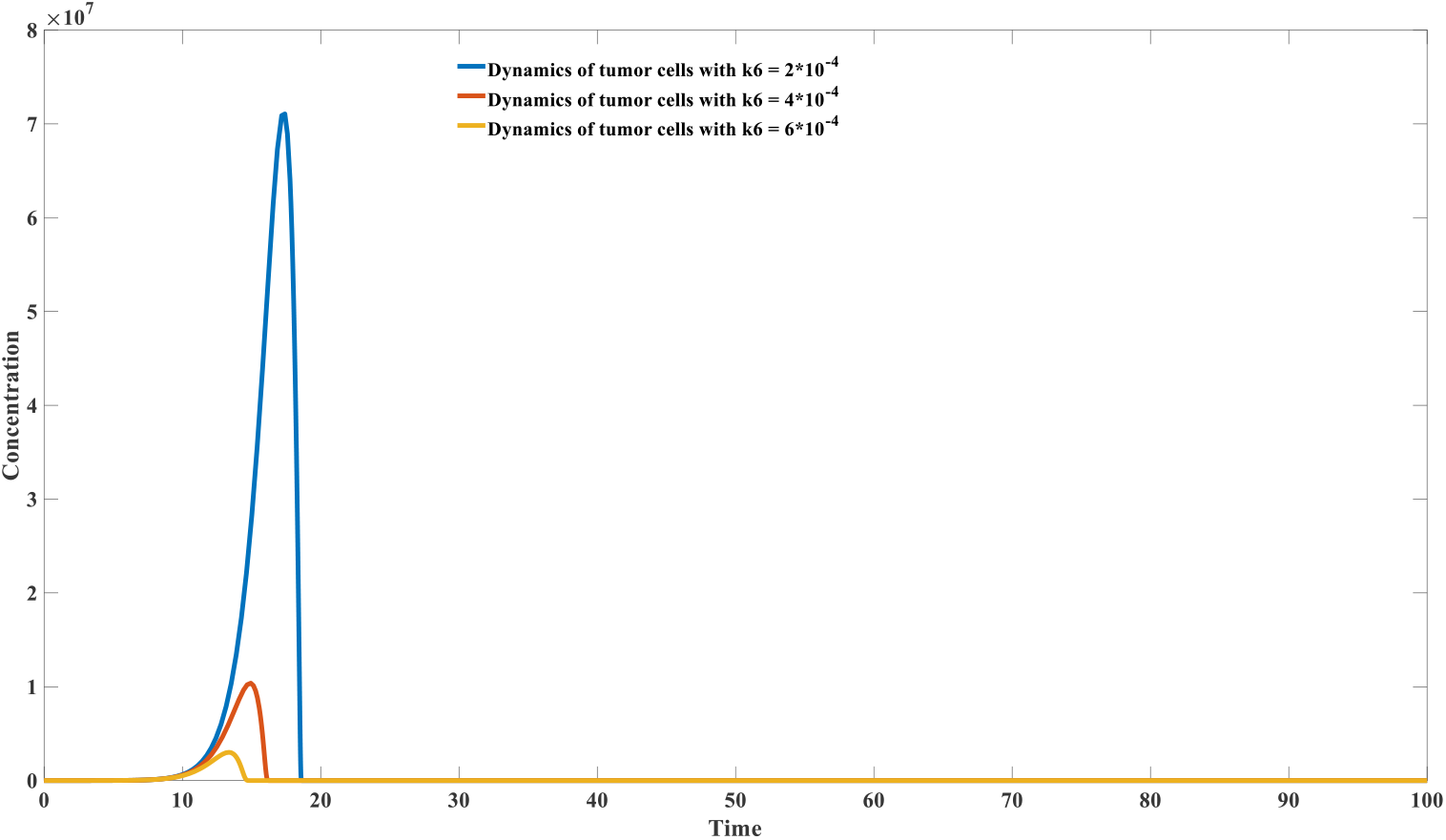
Effect of CD8+T neoantigen (MHC-1 neoantigen) in cancer therapy

**Figure 5H:**
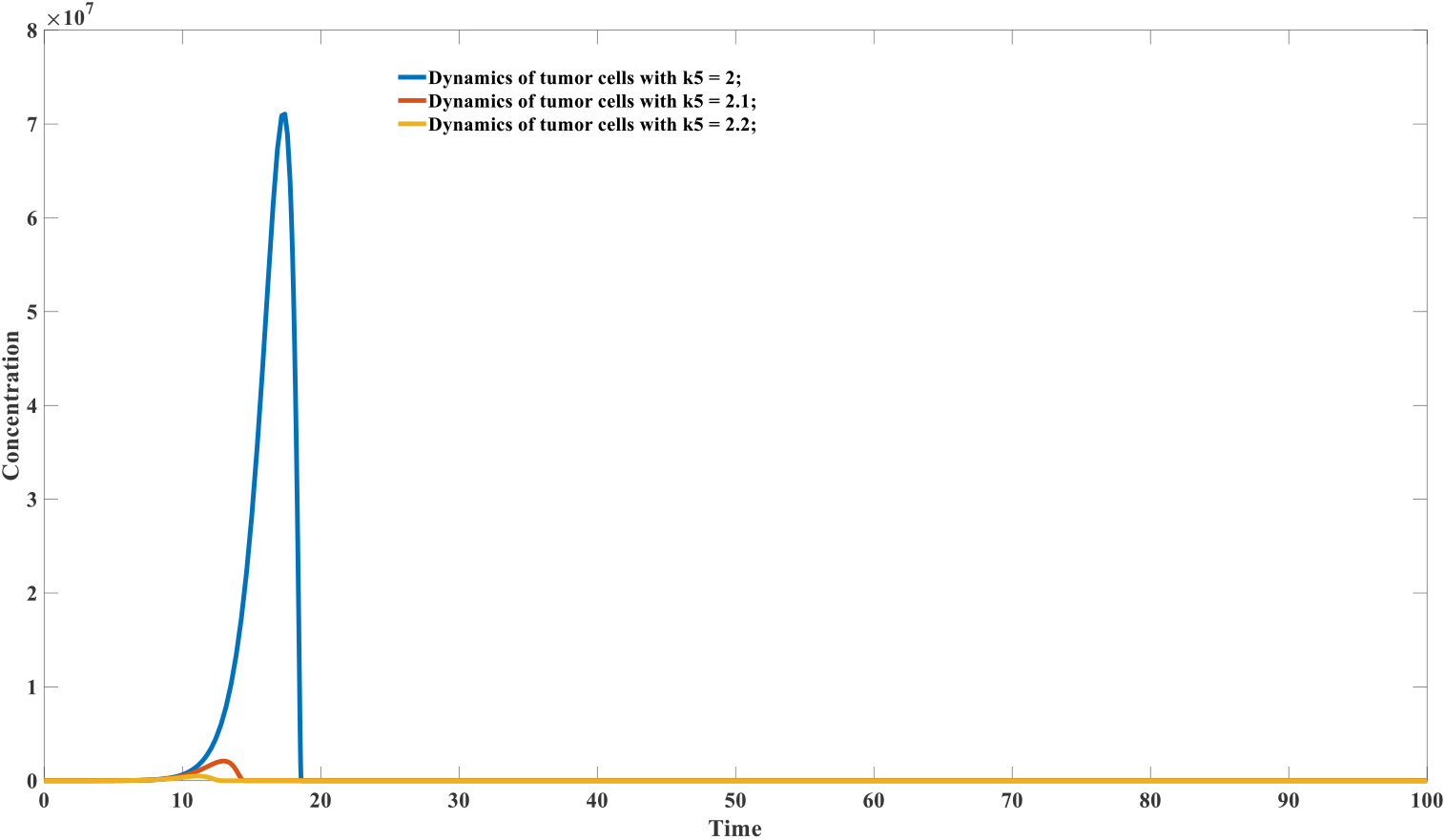
Effect of CD4+ neoantigen (MHC-IIneoantigen) in cancer therapy

## Methods

### 3.1 : Humoral immunity model in adaptive immunity

The core model of our humoral immunity model of adaptive immunity is as follows (Model 3.1.1) :

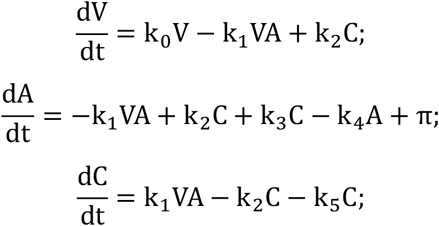

V stands for virus, A stands for antibody, and C stands for virus - antibody. complex. A mathematical analysis of this model is provided in the supplementary material. k_0_ represents the virus’s self-replication rate, k_1_ represents the positive binding parameter between the virus and the antibody, k_2_ represents the dissociation parameter of the virus-antibody complex; k_3_ represents the regeneration coefficient of the virus-antibody complex to the antibody; k_4_ represents the degradation rate of the antibody; π represents the antibody’s replenishment constant; k_5_ represents the clearance rate of the virus-antibody complex. A major problem with the above model is its inability to maintain immune memory. This is reflected in the fact that, during viral infection, although specific antibodies proliferate during infection (the presence of the second term −k_4_A + π, while both k_4_and π remain constant), the levels of all specific antibodies will eventually return to their original equilibrium state without increasing. This is inconsistent with the immune system’s long-term memory function. Therefore, we further modified the simple model by incorporating environmental antigens. The new model is as follows (Model 3.1.2) :

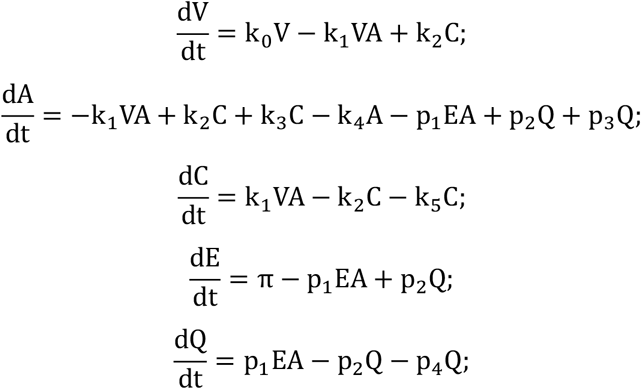

Among them, E represents environmental antigen substances, or self-antigen substances, its positive binding coefficient with antibodies is p_1_, Q represents the antibody-E binding complex, its dissociation rate is p_2_, the stimulation antibody regeneration coefficient is p_3_, and the self-degradation rate is p_4_.

To validate our core model, we simulated three scenarios: how the immune system selects antibodies with high binding activity in the presence of multiple antibodies; viral rebound during monoclonal antibody therapy; and viral rebound during viral replication inhibitor therapy. These three models are represented by ODEs. The immune system’s screening mechanism is modeled as follows (Model 3.1.3) :

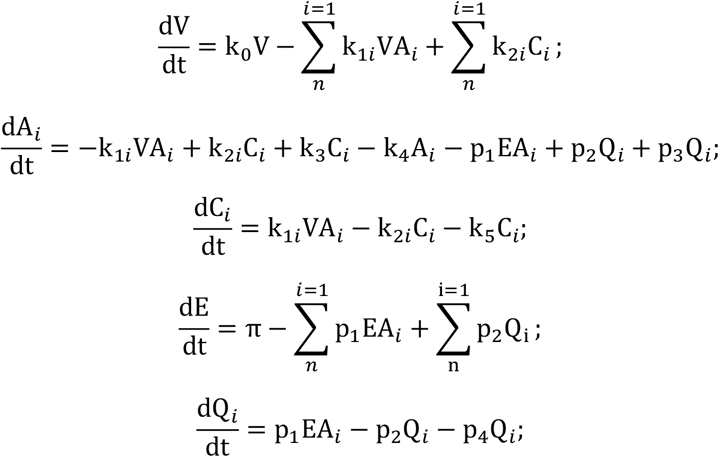

where 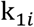 represents the positive binding coefficient of different antibodies to virus V; k_2*i*_ represents the dissociation coefficient of the virus-antibody complex; k_1*i*_/k_2*i*_ represents the equilibrium constant for antibody-virus binding Kd. Stronger binding forces indicating a larger k_1*i*_ and a smaller k_2*i*_. Due to the diversity of environmental antigens E, the overall binding and dissociation coefficients of different antibodies for environmental antigens E are constants, represented by p_1_ and p_2_ respectively. Simultaneously, the antibody regeneration coefficient and degradation rate are also constants, represented by p_3_ and p_4_ respectively. This model allows us to observe not only the proliferation of antibodies with different kinetic parameters during viral infection, but also the shift in the proportions of different antibodies as they reach a new equilibrium after viral clearance. This is the mechanism by which the immune system maintains long-term memory.

The viral rebound model in monoclonal antibody therapy is as follows (Model 3.1.4) :

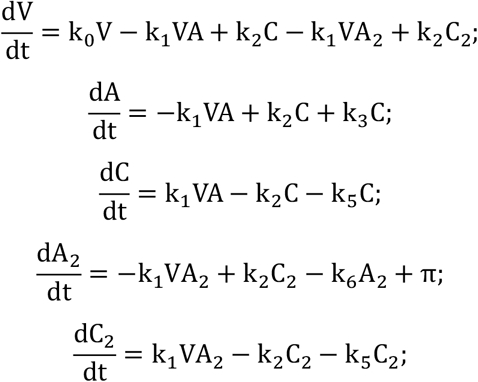

where A_2_ represents the injected monoclonal antibody. For simplicity, we assume that this monoclonal antibody has the same kinetic parameters k_1_ and k_2_ as the specific neutralizing antibody produced by the human body. However, the complex formed by this monoclonal antibody binding to the virus (C_2_)does not stimulate antibody A_2_regeneration and exhibits a faster degradation rate k_6_. At the time of injection, π is the injected dose. At other times π = 0.

ODEs model for viral replication inhibitor therapy is as follows (Model 3.1.5) :

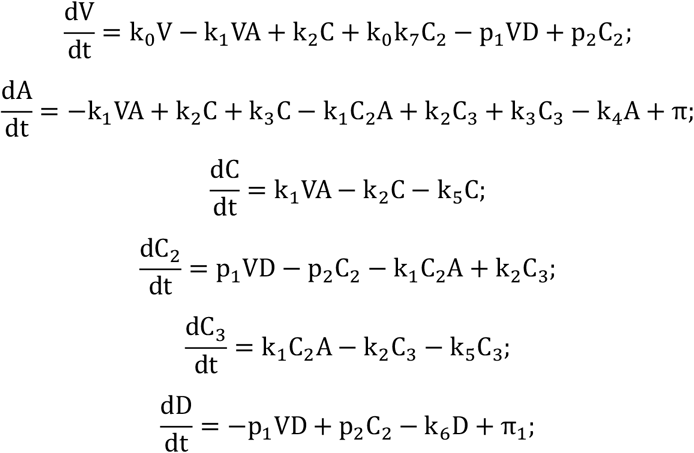

Where V represents virus, A represents antibody, C represents virus-antibody complex, C_2_ represents virus-inhibitor complex, and D represents small molecule inhibitor. C_3_represents the complex of virus-inhibitor and antibody. As long as the complex formed by binding to the antibody, including C and C_3_, has the same degradation coefficient k_5_, the degradation coefficient of the small molecule drug is k_6_, the degradation coefficient of the antibody is k_4_, and the antibody replenishment constant is π. The positive binding coefficient of the small molecule inhibitor with the virus is p_1_, the negative dissociation constant is p_2_, k_0_represents the viral replication rate, and k_7_represents the inhibitory coefficient of the small molecule on viral replication, which is a value between 0 and 1. The replenishment coefficient of the small molecule is π_1_, which is equal to the injected dose at the time of injection and π_1_ =0 at other times.

Further expand the antibody dynamics model to include more compartments, especially considering ASC (antibody secreting cell), a more precise model is expressed as follows (Model 3.1.6):

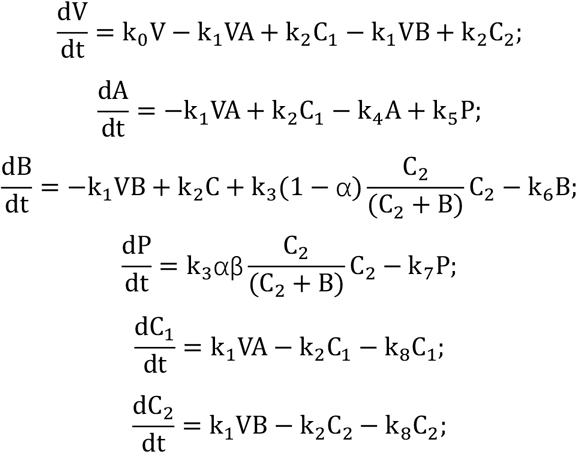

Wherein, A represents antibody, P represents ASC, B represents BCR on non-plasma B cells, which is proportional to the number of non-plasma B cells, k_3_ represents the positive feedback coefficient of BCR-Virus complex stimulating BCR regeneration, α represents the coefficient of non-plasma B cells converting to ASC after antigen stimulation, β is the corresponding coefficient of BCR and ASC, k_5_ represents the rate constant of ASC generating antibodies. When the virus is naturally infected, k_0_ is the virus reproduction coefficient, which is a positive value. When the vaccine is injected, k_0_ is the antigen degradation coefficient, which is a negative value. k_4_ is the antibody degradation coefficient, k_6_ is the non-plasma B cell death coefficient, k_7_ is the ASC death coefficient, and k_8_ is the degradation coefficient of the (antigen-antibody) or (antigen-BCR) complex.

When the above model is further expanded with more compartments, specifically dividing BCR cells into IgM-BCR and IgG-BCR, antibodies into IgM and IgG, and ASC cells into IgM-ASC and IgG-ASC, while also considering the diversity of IgG and IgM, a more complicated model is expressed as follows (Model 3.1.7):

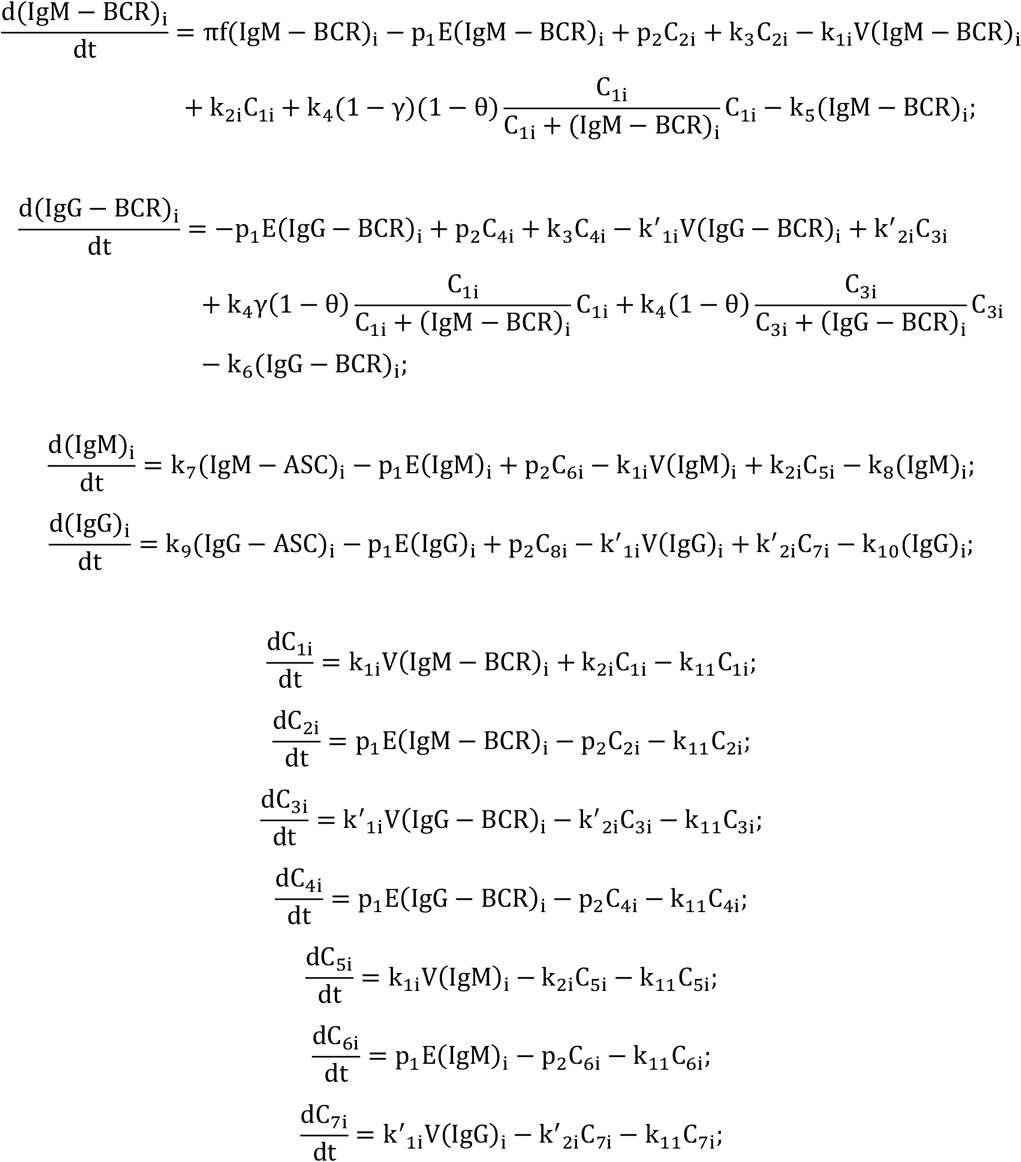

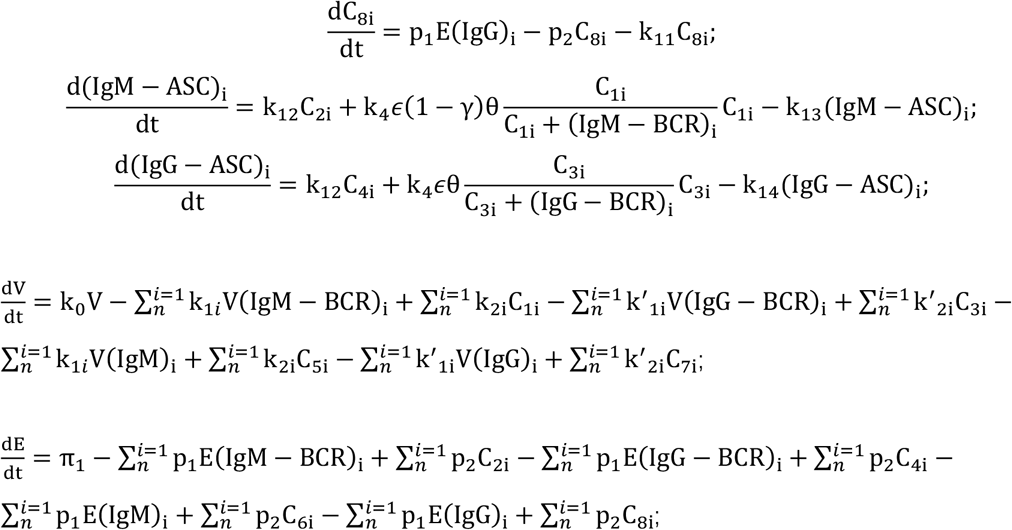

Here, f(IgM − BCR)_i_ represents the proportion of (IgM − BCR)_i_ among all IgM-BCRs, and this proportion follows a normal distribution based on its binding coefficient. The symbol π*π* denotes the replenishment constant for all IgM-BCRs. Different IgM-BCRs are denoted as (IgM − BCR)_i_, where the forward binding coefficient with environmental antigen E is p_1_, and the reverse dissociation coefficient of the resulting complex C_2i_ is p_2_. The degradation rate of all antigen-antibody complexes is k_11_.

Different IgG-BCRs are denoted as (IgG − BCR)_i_, where the forward binding coefficient with environmental antigen E is p_1_, and the reverse dissociation coefficient of the resulting complex C_4i_ is p_2_.

Different IgMs are denoted as (IgM)_i_, where the forward binding coefficient with environmental antigen E is p_1_, and the reverse dissociation coefficient of the resulting complex C_6i_ is p_2_.

Different IgGs are denoted as (IgG)_i_, where the forward binding coefficient with environmental antigen E is p_1_, and the reverse dissociation coefficient of the resulting complex C_8i_ is p_2_.

The complex formed between (IgM − BCR)_i_ and virus V is denoted as C_1i_, with a forward binding constant k_1*i*_ and a reverse dissociation constant k_2i_. The complex formed between (IgG − BCR)_i_ and virus V is denoted as C_3i_, with a forward binding constant k^′^_1i_ and a reverse dissociation constant k^′^_2i_.

For IgM and IgG of the same isotype, k_1*i*_ = k^′^_1i_, k_2i_ = k^′^_2i_. However, since free IgM predominantly exists in a pentameric form, its binding capacity to the virus is significantly stronger than that of IgG. In the simulation, k_1*i*_=5 * k^′^_1i_ and k_2i_ = k^′^_2i_ were used. Similarly, for IgM-BCR, which tends to form multimers on the B-cell surface, its viral affinity is also stronger than that of IgG-BCR. In the simulation, the same values k_1*i*_=5 * k^′^_1i_ and k_2i_ = k^′^_2i_ were applied. This effectively explains why, during initial viral infection, IgM often exhibits significant proliferation compared to IgG, even when their initial distributions are similar.

The meanings of other parameters are as follows:

- k_0_ : viral proliferation coefficient
- k_3_ : proliferation coefficient of environmental antigen-BCR complexes for BCRs
- k_4_: proliferation coefficient of viral antigen-BCR complexes for BCRs
- γ : conversion coefficient of IgM-BCR–type B cells to IgG-BCR–type B cells
- θ : maximum conversion coefficient of non-plasma B cells to antibody-secreting cells (ASCs)
- k_5_ : degradation rate of IgM-BCR
- k_6_: degradation rate of IgG-BCR
- k_7_: production rate of IgM by IgM-ASCs
- k_8_: degradation rate of IgM
- k_9_: production rate of IgG by IgG-ASCs
- k_10_: degradation rate of IgG
- k_11_: regeneration coefficient of environmental antigen-BCR complexes for ASCs
- k_13_: degradation coefficient of IgM-ASCs
- k_14_: degradation coefficient of IgG-ASCs
- ϵ*ϵ*: ratio coefficient of ASCs to BCRs
- π1*π*1: replenishment constant of environmental antigen E

### 3.2 Cellular immunity models in adaptive immunity

Cellular immunity models in adaptive immunity can be broadly divided into two categories based on the type of antigen: antibody-independent and antibody-dependent. Cellular immunity is primarily manifested through the activation and expansion of specific CD8+ T cells, which we believe are primarily driven by DCs. In the first category, because antigens (such as SARS-CoV-2) possess a unique structure that is recognized by pattern recognition receptors (TLRs and CLRs) on the surface of DCs, the antigens themselves can directly enter DCs, where they are degraded and ultimately presented to MHC-I complexes, triggering the activation and proliferation of CD8+ T cells. Furthermore, these antigens can bind to antibodies, forming an antigen-antibody complex. At this point, the antibody’s Fc region undergoes conformational changes, allowing it to bind to Fc receptors on the DC surface, thereby entering the DC cell and subsequently activating and proliferating CD8+ T cells. For other antigens (such as self-antigens and LMCV), the antigens themselves do not possess a specific structure, and therefore can only be recognized by DCs and further processed to promote CD8 + T cell proliferation after binding to antibodies. The humoral and cellular immunity models for these two types of antigens are as follows (Model 3.2.1) : Type I antigens:

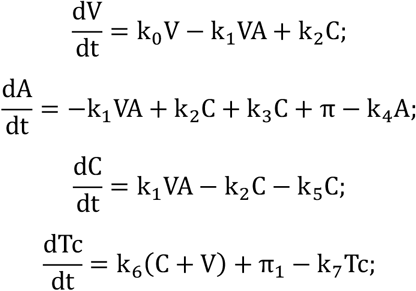

Second type of antigens:

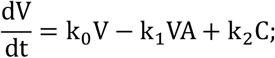

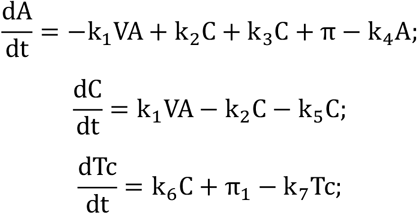

In the case of viral infection, k_0_ takes a relatively large positive value, whereas in the context of vaccination—where the antigen lacks self-replication capability— k_0_ is negative, representing the decay process of the antigen in vivo. The parameter k_1_ denotes the forward binding coefficient between the antigen and antibody, k_2_ represents the dissociation coefficient of the antigen–antibody complex, and k_3_ indicates the regeneration coefficient of antibodies induced by the antigen–antibody complex. For patients undergoing rituximab treatment, k_3_ is significantly lower than that in the normal population. The parameter k_5_ describes the clearance rate of antigen–antibody complexes, which is markedly higher than the degradation rate of the antigen itself. Detailed parameter selections are provided in the Supplementary Materials.

To investigate the relationship between the cytotoxic capacity of Tc cells and the intracellular antigen concentration in infected cells, we assume that at a given time point, N_i_represents the number of cells with a specific intracellular antigen concentration, and V_i_ denotes the total viral load. The antigen concentration is defined as V_i_/N_i_, and C_i_ refers to the number of complexes formed by the binding of N_i_ and Tc cells leading to target cell killing. Based on these definitions, we constructed the following system of ordinary differential equations (Model 3.2.2):

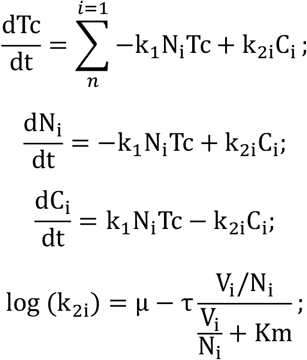

Here, k_2i_ represents the reverse dissociation coefficient between Tc cells and infected cells N_i_, which is a variable whose logarithm is related to the antigen concentration. The parameter μ denotes the maximum dissociation constant, corresponding to the dissociation coefficient in the absence of antigen, while μ − τ represents the minimum dissociation coefficient, i.e., the dissociation coefficient when the antigen concentration approaches infinity. Km is the antigen concentration at which the binding affinity is half of its maximum value.

We assume that the forward binding coefficient between Tc cells and infected cells N_i_ is a constant, k_1_, determined by the collision probability of the two cells. The equilibrium constant between Tc cells and infected cells, k_d_ = k_2i_/k_1_, is proportional to the binding energy. Since the binding energy increases with antigen concentration—due to the higher number of TCR– MHC class I complexes on the cell surfaces at elevated antigen levels—we derive the following expression for the dissociation coefficient:

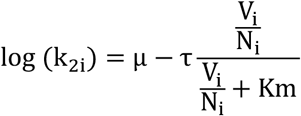

Detailed parameter settings are provided in the Supplementary Materials.

### 3.3 Humoral immunity model considering B-CD4+ T cell interactions

ODEs of the humoral immunity model considering B-CD4+ T cell interactions are as follows (Model 3.3.1) :

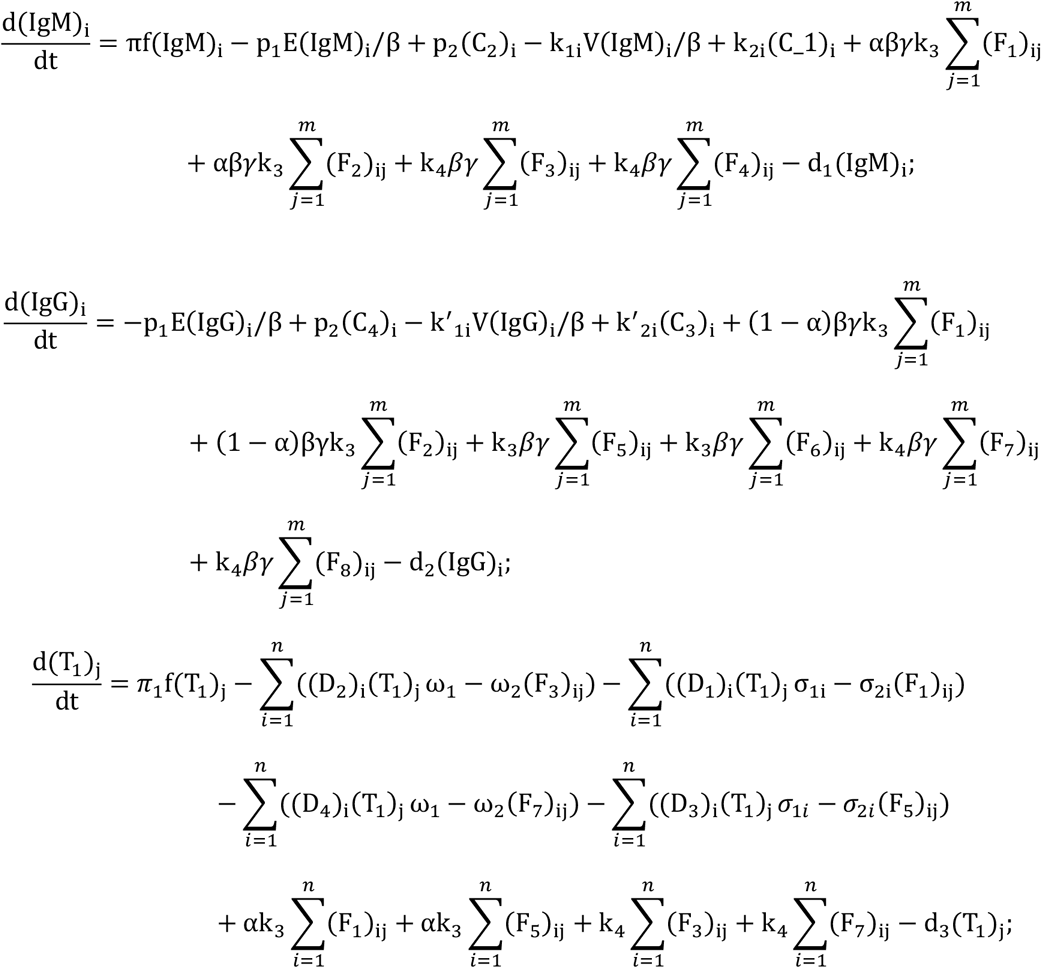

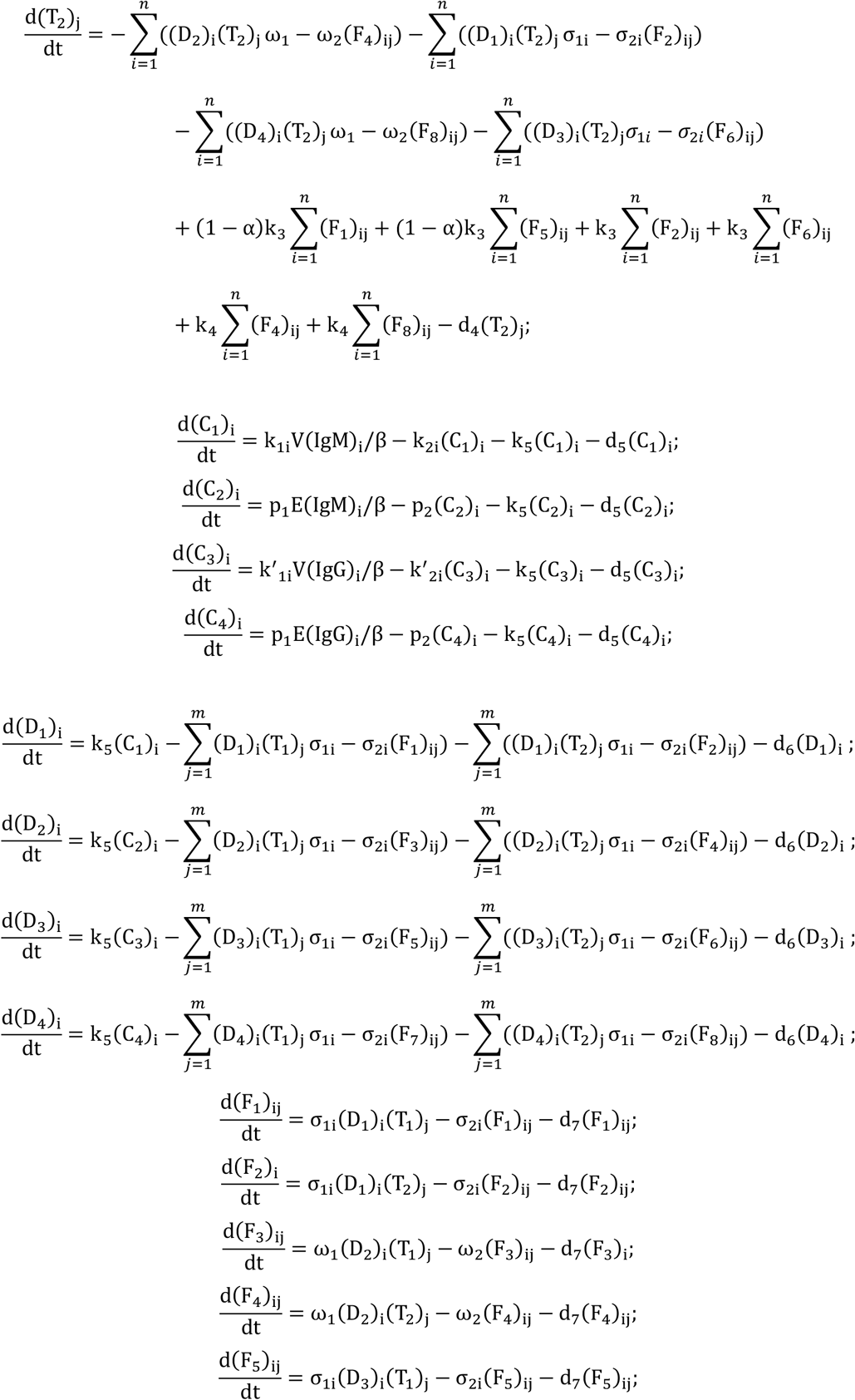

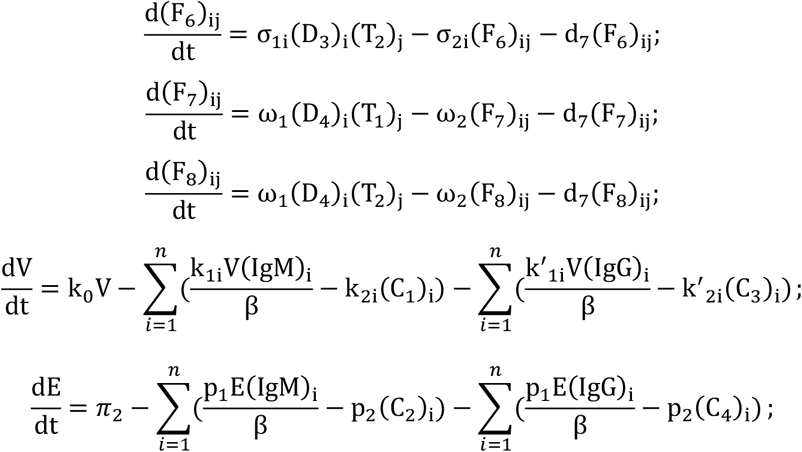

Here, (IgM)_i_ denotes the *i*-th type of IgM, and the corresponding IgG is denoted as (IgG)_i_. The complex formed by the binding of (IgM)_i_ to virus V is (C_1_)_i_, with a forward binding rate k_1i_ and a reverse dissociation rate k_2i_. After degradation, (C_1_)_i_ is presented on MHC class II complexes as (D_1_)_i_ with a conversion rate k_5_, while the intrinsic degradation rate of (C_1_)_i_ is d_5_. The complex formed between (D_1_)_i_ and non-memory CD4+ T cells (T_1_)_j_ is (F_1_)_ij_, and that with memory CD4+ T cells (T_2_)_j_ is (F_2_)_ij_, both with a forward binding rate σ_1i_ and a reverse dissociation rate σ_2i_. The degradation rate of (*D*)_i_ is d_6_, and that of (*F*)_ij_is d_7_.

The complex formed by the binding of (IgM)_i_ to environmental antigen E is (C_2_)_i_, with a forward binding rate p_1_ and a reverse dissociation rate p_2_. After degradation, (C_2_)_i_ is presented on MHC class II complexes as (D_2_)_i_, with a conversion rate k_5_, while the intrinsic degradation rate of (C_2_)_i_ is d_5_. The complex formed between (D_2_)_i_ and non-memory CD4+ T cells (T_1_)_j_ is (F_3_)_ij_, and that with memory CD4+ T cells (T_2_)_j_ is (F_4_)_ij_, both with a forward binding rate ω_1_ and a reverse dissociation rate ω_2_.

The *i*-th type of IgG, denoted as (IgG)_i_, binds to virus V to form the complex (C_3_)_i_, with a forward binding rate k^′^_1i_ and a reverse dissociation rate k^′^_2i_. After degradation, (C_3_)_i_ is presented on MHC class II complexes as (D_3_)_i_, with a conversion rate k_5_, while the intrinsic degradation rate of (C_3_)_i_ is d_5_. The complex formed between (D_3_)_i_ and non-memory CD4+ T cells (T_1_)_j_ is (F_5_)_ij_, and that with memory CD4+ T cells (T_2_)_j_ is (F_6_)_ij_, both with a forward binding rate σ_1i_ and a reverse dissociation rate σ_2i_.

The complex formed by the binding of (IgG)_i_ to environmental antigen E is (C_4_)_i_, with a forward binding rate p_1_ and a reverse dissociation rate p_2_. After degradation, (C_4_)_i_ is presented on MHC class II complexes as (D_4_)_i_, with a conversion rate k_5_, while the intrinsic degradation rate of (C_4_)_i_ is d_5_. The complex formed between (D_4_)_i_ and non-memory CD4+ T cells (T_1_)_j_ is (F_7_)_ij_, and that with memory CD4+ T cells (T_2_)_j_ is (F_8_)_ij_, both with a forward binding rate ω_1_ and a reverse dissociation rate ω_2_.

Other constants are defined as follows:

- α: conversion coefficient from IgM to IgG
- β: proportionality coefficient between BCR and B cells
- *γ*: expansion proportionality coefficient between B cells and CD4+ T cells
- *k*_3_: antibody regeneration coefficient induced by environmental antigen–CD4+ T cell complexes
- *k*_4_: antibody regeneration coefficient induced by viral antigen–CD4+ T cell complexes
- π: replenishment constant for IgM
- *π*_1_: replenishment constant for non-memory CD4+ T cells
- *π*_2_: replenishment constant for environmental antigens

The proportion of (IgM)_i_ in total IgM, denoted as f(IgM)_i_, follows a normal distribution and is related to its binding coefficient. Similarly, the proportion of non-memory CD4+ T cells of type (T_1_)_j_, denoted as f(T_1_)_j_, in the total non-memory CD4+ T cell population, also follows a normal distribution and depends on its binding coefficient.

Detailed parameter settings and simulation results are provided in the Supplementary Materials.

### 3.4 Agent - based model considering temporal infection sequence of infected cells

Pseudocodes of agent-based model considering the temporal infection sequence of infected cells are as follows:

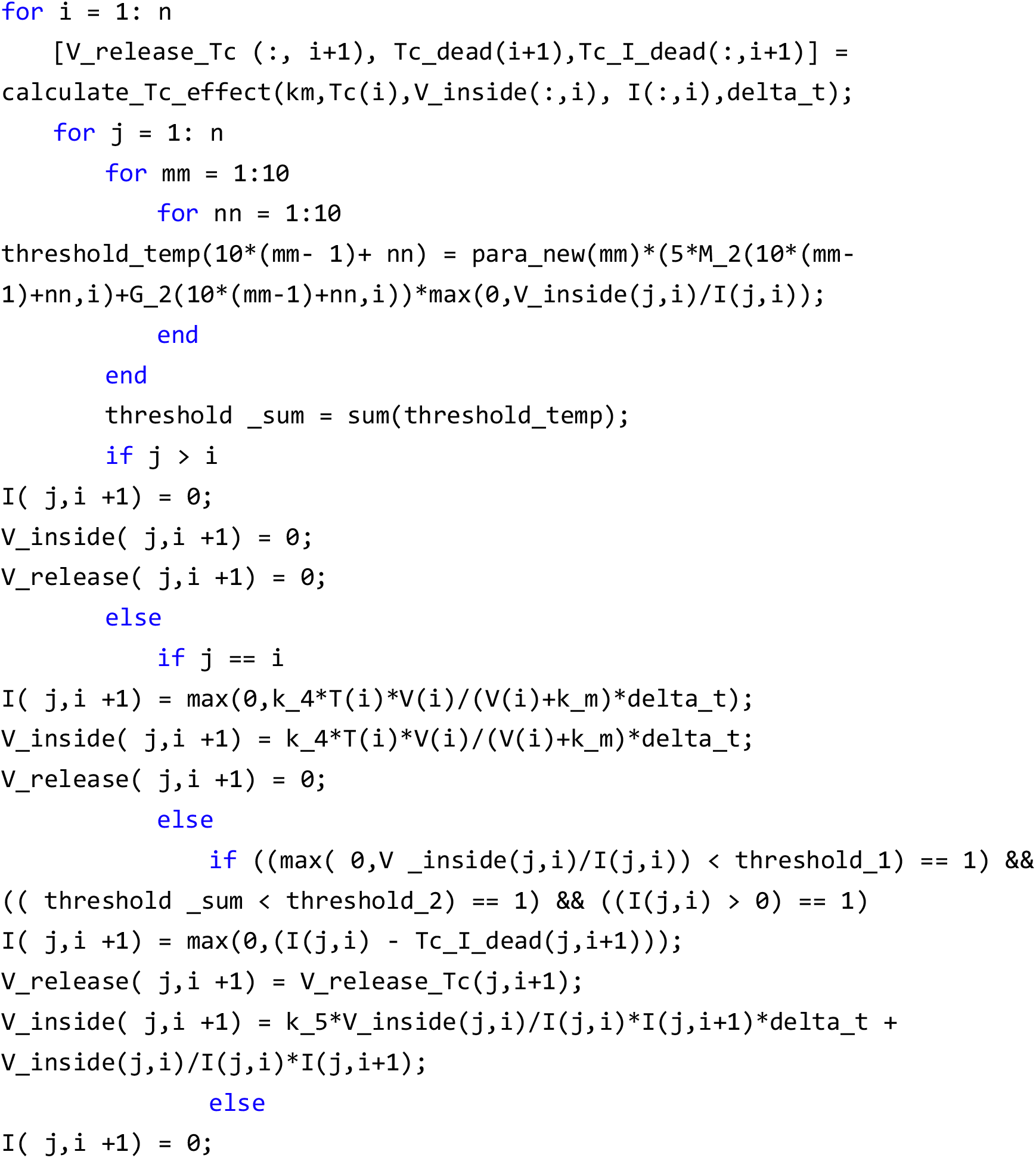
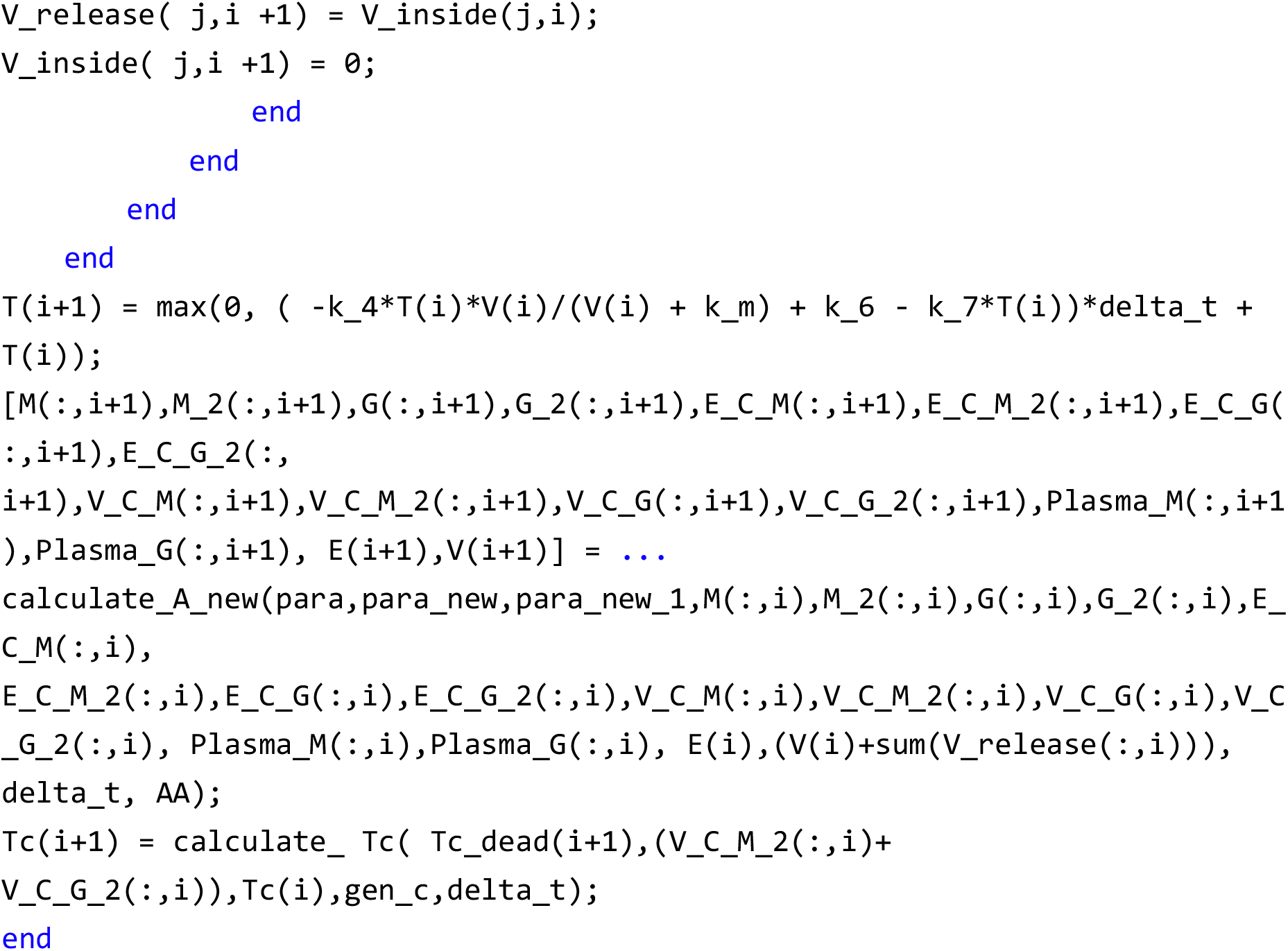

V_release_Tc is an *n * n* matrix, where V_release_Tc (i,j) represents the total amount of virus released by infected cells at time *i* through Tc cell lysis at time *j*.

Tc_dead is an *n *1* vector, where Tc_dead(i) represents the total number of cells killed by Tc cells at time *i*.

Tc_I_dead is an *n * n* matrix, where Tc_I_dead(i,j) represents the number of cells infected at time i that died through Tc cell lysis at time *j*.

Tc is an *n *1* vector, where Tc(i) represents the number of Tc cells at time *i*.

V_inside is an *n * n* matrix, where V_inside(i,j) represents the total amount of virus inside cells infected at time *i* at time *j*.

I is an *n * n* matrix, where I(i,j) represents the number of cells infected at time *i* at time *j*. The function calculate_Tc_effect calculates the lysis of infected cells caused by cell-mediated immunity over a fixed time period.

threshold_temp(10*(mm-1)+nn) represents the binding of a particular antibody to the virus on the cell surface at time *i*. It is equal to the sum of the positive binding coefficient of IgG * the intracellular virus concentration (max(0,V_inside(j,i)/I(j,i))) and the positive binding coefficient of IgM * the intracellular virus concentration. The positive binding coefficient of IgM is equal to 5 times the positive binding coefficient of IgG.

threshold_sum represents the binding of all types of antibodies to the virus on the cell surface at time *i*.

If time *j* is greater than *i*, it means that infection has not occurred yet, so I(j,i+1) = 0; V_inside(j,i+1) = 0; V_release(j,i+1) = 0; if j = i, it means that the infection occurred here, I(j,i+1) = max(0,k_4*T(i)*V(i)/(V(i)+k_m)*delta_t); V_inside(j,i+1) = k_4*T(i)*V(i)/(V(i)+k_m)*delta_t; V_release(j,i+1) = 0; here k_4 represents the virus invasion constant, V(i) represents the number of viruses in the body fluid at time *i*, T(i) represents the number of susceptible cells at time *i*, k_m is a coefficient equal to the number of viruses at half the maximum infection number, and delta_t represents the time interval.

If time j is less than i, then the following situations occur: If ((max(0,V_inside(j,i)/I(j,i)) < threshold_1) == 1) && ((threshold _sum < threshold_2) == 1) && ((I(j,i) > 0) == 1), where threshold_1 represents the virus concentration threshold at which the cell naturally lyses. If it is greater than this threshold, the cell naturally lyses, so threshold_1 is also a relatively large value. threshold_2 is the concentration threshold of the antigen-antibody complex when the ADCC effect occurs. When threshold_sum is greater than this threshold, the cell lyses due to ADCC. If both thresholds are not met at the same time and the cell has been infected (I(j,i) > 0), then I(j,i+1) = max(0,(I(j,i) - Tc_I_dead(j,i+1))) indicates that the number of infected cells at the next moment is equal to the number of infected cells at this moment minus the number of cells lysed by Tc cells at the next moment. At the same time, the amount of virus released into the body fluid at the next moment is equal to the total number of viruses in the cells lysed by the cytotoxic action of Tc cells at the next moment. V_release(j,i+1) = V_release_Tc(j,i+1); and the number of viruses in the body V_inside(j,i+1) = k_5*V_inside(j,i)/I(j,i)*I(j,i+1)*delta_t + V_inside(j,i)/I(j,i)*I(j,i+1), where k_5 represents the virus reproduction coefficient. Otherwise, once one of these two thresholds is met, all cells will undergo lysis, I(j,i+1) = 0; V_release(j,i+1) = V_inside(j,i); V_inside(j,i+1) = 0.

The number of susceptible cells T at the next moment is T(i+1) = max(0,(-k_4*T(i)*V(i)/(V(i) + k_m) + k_6 - k_7*T(i))*delta_t + T(i)), where k_6 represents the replenishment constant of susceptible cells and k_7 represents the attenuation coefficient of susceptible cells.

[M(:,i+1),M_2(:,i+1),G(:,i+1),G_2(:,i+1),E_C_M(:,i+1),E_C_M_2(:,i+1),E_C_G(:,i+1),E_C_G_2(:, i+1),V_C_M(:,i+1),V_C_M_2(:,i+1),V_C_G(:,i+1),V_C_G_2(:,i+1),Plasma_M(:,i+1),Plasma_G(:,i+1), E(i+1),V(i+1)] =calculate_A_new(para,para_new,para_new_1,M(:,i),M_2(:,i),G(:,i),G_2(:,i),E_C_M(:,i), E_C_M_2(:,i),E_C_G(:,i),E_C_G_2(:,i),V_C_M(:,i),V_C_M_2(:,i),V_C_G(:,i),V_C_G_2(:,i), Plasma_M(:,i),Plasma_G(:,i), E(i),(V(i)+sum(V_release(:,i))), delta_t, AA);

- M: A *t*×*n* matrix, where *t* is the total number of IgM-BCR variants; M(i,j) represents the quantity of the *i*-th IgM-BCR variant at time *j*.
- M_2: A *t*×*n* matrix; M_2(i,j) represents the quantity of the *i*-th IgM variant at time *j*.
- G: A *t*×*n* matrix; G(i,j) represents the quantity of the *i*-th IgG-BCR variant at time *j*.
- G_2: A *t*×*n* matrix; G_2(i,j) represents the quantity of the *i*-th IgG variant at time *j*.
- E_C_M(i,j): Number of complexes formed between the *i*-th IgM-BCR variant and environmental antigen *E* at time *j*.
- E_C_M_2(i,j): Number of complexes formed between the *i*-th IgM variant and environmental antigen *E* at time *j*.
- E_C_G(i,j): Number of complexes formed between the *i*-th IgG-BCR variant and environmental antigen *E* at time *j*.
- E_C_G_2(i,j): Number of complexes formed between the *i*-th IgG variant and environmental antigen *E* at time *j*.
- V_C_M(i,j): Number of complexes formed between the *i*-th IgM-BCR variant and virus *V* at time *j*.
- V_C_M_2(i,j): Number of complexes formed between the *i*-th IgM variant and virus *V* at time *j*.
- V_C_G(i,j): Number of complexes formed between the *i*-th IgG-BCR variant and virus *V* at time *j*.
- V_C_G_2(i,j): Number of complexes formed between the *i*-th IgG variant and virus *V* at time *j*.
- Plasma_M(i,j): Number of the *i*-th IgM plasma cell variant at time *j*.
- Plasma_G(i,j): Number of the *i*-th IgG plasma cell variant at time *j*.
- E(i): Quantity of environmental antigen E at time *i*.
- V(i): Viral load of virus *V* in body fluids at time *i*.
- AA: A given normal distribution probability function.

The function calculate_A_new is used to compute the quantities of all components after the next time step (delta_t), based on the principles outlined in Model 3.1.7.

Detailed parameter settings and simulation results are provided in the Supplementary Materials.

### 3.5 Tumor-host interaction model

The interaction model between cancer cells and the immune system is as follows (Model 3.5.1) :

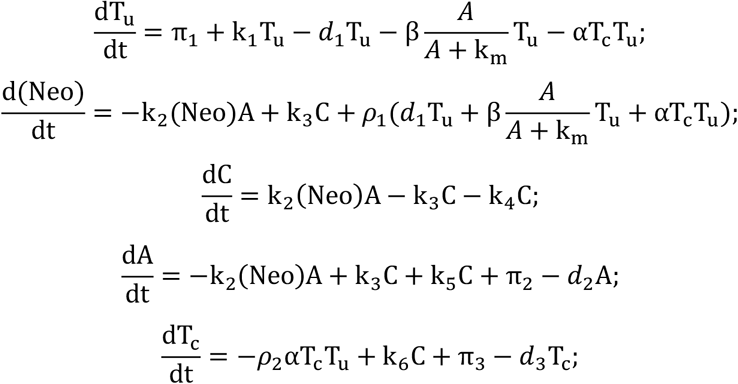

Which T_u_ means tumor cell, A represents the specific antibody, (Neo) represents the neoantigen, C represents the antibody-neoantigen complex, and T_c_ represents CD8+ T cells. π_1_ represents the cancer cell replenishment constant, k_1_ represents the cancer cell proliferation coefficient, *d*_1_ represents the cancer cell death coefficient, 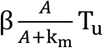 represents the killing effect of the ADCC on cancer cells, β represents the maximum killing coefficient, k_m_represents a constant, represents the antibody concentration at half the maximum killing capacity, αT_c_T_u_ represents the killing effect of cellular immunity on cancer cells, and α represents the killing coefficient. k_2_represents the positive binding coefficient between the neoantigen and the antibody, k_3_ represents the negative dissociation coefficient, *ρ*_1_represents the number of neoantigens per cell, k_4_represents the degradation rate of the antigen-antibody complex, k_5_ represents the regeneration coefficient of the antigen-antibody complex on the antibody, k_6_represents the coefficient of the antigen-antibody complex stimulating T_c_ regeneration, π_2_ represents the antibody replenishment constant, *d*_2_ represents the antibody attenuation coefficient, *ρ*_2_ represents the T_c_ exhaustion coefficient, π_3_ represents the replenishment constant of T_c_, and *d*_3_ represents the attenuation coefficient of T_c_.

## Discussion

Despite significant advances in mathematical modeling of virus-host interactions in recent years, several critical challenges remain unresolved. We summarize these key issues into three major aspects:

1. Distorted Kinetic Assumptions: Classical models employing the βTV term often overlook saturation effects in virus-cell interactions, leading to misinterpretations of infection clearance mechanisms. In the Supplementary Materials, we critique the classic cell depletion model, which utilizes the βTV term to describe the conversion of susceptible cells (T) to infected cells (I) by virus (V). This formulation appears consistent with second-order reaction kinetics, particularly when drawing analogies to SIR epidemic models. However, the actual process of viral infection does not adhere to second-order kinetics, which presupposes equivalent reacting entities (e.g., both being individuals or small molecules). In reality, viral particle counts typically vastly exceed cell counts, and the components are not homogeneous. Assuming a probability *p*_1_ that a single virion infects a cell per unit time, the probability for N virions becomes 1 − (1 − *p*_1_)^*N*^. Consequently, for virus concentration V infecting target cell concentration T, the conversion relationship should be (1 − (1 − *p*_1_)^*V*^)*T*, which deviates significantly from the βTV term. It more closely resembles Michaelis-Menten kinetics 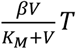, where the maximum conversion rate approaches βT, not infinity, even at saturating viral concentrations. Using the conventional βTV term incorrectly implies that susceptible cell depletion is the primary driver of infection termination, regardless of immune responses. Consequently, immune activation models incorporating βTV for infection, while potentially allowing a non-zero steady-state for susceptible cells, often predict an unrealistic, drastic decline in T. This contradicts biological observations where only a minimal fraction of susceptible cells becomes infected, and viral clearance is predominantly mediated by activated immune responses, fundamentally differing from SIR-type population dynamics. In essence, susceptible cell depletion is not the decisive factor for infection resolution. Furthermore, critical misconceptions exist in modeling adaptive immunity. For cellular immunity, Cytotoxic T Lymphocyte (CTL) activation is primarily mediated by Antigen-Presenting Cells (APCs), with later rapid proliferation driven by B cells (also APCs) presenting antigen. CTLs are not directly activated by ordinary infected cells (I); interaction with I typically leads to CTL exhaustion or depletion. Thus, equations like 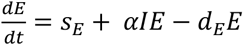 are fundamentally flawed. Accurate modeling requires incorporating an intermediate term representing virus-antibody complexes or virus-B cell interactions, which drive CTL proliferation. This approach better explains early CTL expansion and subsequent exhaustion during chronic infection. Similarly, for humoral immunity, modeling antibody (A) production simply via arbitrary functions like *h*(*V*(*t* − *τ*), *A*(*t* − *τ*)) based on mathematical fitting is inadequate. The direct driver of antibody proliferation is the concentration of virus-antibody complexes. Mechanistically grounded representations of biological processes are superior to purely phenomenological mathematical expressions. While complex formulas or additional parameters might improve curve fitting, they often fail to capture true biology, compromising the model’s predictive power.
2. Insufficient Immune Integration: Humoral and cellular immune responses are often studied in isolation, lacking quantitative descriptions of their dynamic synergy. Although Leon et al. (2020) developed a model integrating both effects^[44]^, such studies remain scarce. A comprehensive approach simultaneously considering both humoral and cellular immunity is crucial. CD8+ T cell-mediated cytotoxicity lyses infected cells, while B cell-mediated humoral immunity neutralizes virions and can lyse infected cells via Antibody-Dependent Cellular Cytotoxicity (ADCC). Isolating one immune arm fails to capture the integrated immune response and can lead to erroneous conclusions.
3. Neglect of Immune Diversity: Existing models largely ignore B Cell Receptor (BCR) and T Cell Receptor (TCR) diversity, precluding the simulation of affinity maturation and memory maintenance. Advances in experimental techniques, particularly single-cell sequencing of B and T cells, are rapidly mapping the diversity and evolution of BCR/TCR repertoires during infection. Coupled with abundant flow cytometry data, these resources offer multi-dimensional insights into how viral infection reshapes B and T cell populations. This demands models that move beyond bulk antibody/T cell counts towards incorporating subtyping and compartmentalization based on receptor diversity, albeit increasing computational complexity. Incorporating BCR/TCR diversity is essential for deeper mechanistic understanding of virus-host interactions.

Addressing these challenges, we developed a novel mathematical model firmly grounded in biological mechanisms. It explicitly incorporates BCR/TCR diversity and fully integrates both cellular and humoral immune processes. Our ultimate goal is to utilize this model to interpret immunological phenomena and guide vaccine development and immunotherapeutic strategies. This work represents not merely a model refinement but a paradigm shift “from reductionism to systems biology,” employing multi-compartment modeling to reveal emergent properties of immune responses and providing a “quantitative immunology” framework for rational intervention.

In essence, our model addresses several pivotal, unresolved questions in immunology:

- The mechanism underlying clonal expansion of high-affinity antibodies during humoral responses.
- The genesis of viral rebound during monoclonal antibody therapy or viral replication inhibitor treatments.
- The mechanism of antibody interference in vaccination and its implications for scheduling and dosing.
- The kinetics of antibody depletion and associated germinal center dysfunction.
- Distinct kinetic profiles of IgM and IgG during primary versus secondary infections.
- The dynamical mechanisms governing immune memory formation and original antigenic sin.
- Mechanisms of CD8+ T cell activation and the induction of exhaustion.
- The role of B cells in facilitating CD8+ T cell proliferation.
- Mechanisms driving the establishment of chronic infections.
- Mechanisms of oncogenesis and the dynamics of cancer cell-immune system interactions.

Furthermore, our model demonstrates the following capabilities:

- Using an agent-based modeling framework for viral infection, we identified conditions favoring the transition from acute to chronic infection: weak host immunity, relatively high initial specific antibody levels (e.g., from heterologous reinfection), and slow viral replication rates. These scenarios share the common feature of slow viral stimulation of specific antibody production.
- This agent-based model allows quantitative evaluation of different therapeutic strategies (e.g., specific antibody therapy, environmental antigen exposure, therapeutic vaccination) for resolving chronic infection. It suggests theoretical superiority for therapeutic vaccination, indicating required doses are significantly higher than for prophylactic vaccination.
- Through a modeled cancer-immune interaction system, we assessed the efficacy of various immunotherapies in controlling tumor growth. Our model theoretically supports therapeutic vaccine strategies targeting both MHC class I and MHC class II antigens.

Notwithstanding its novelty and potential utility, our model has inherent limitations, most notably the current lack of fitting to specific experimental datasets; its validity awaits further experimental verification. In the Supplementary Materials, we critique metaphysically-inspired parameter fitting practices and propose a new modeling strategy based on steady-state analysis, although necessary parameter estimation remains imperative. Importantly, future validation should utilize not just single-dimension clinical data (e.g., ELISA antibody levels, qPCR viral loads) but also higher-dimensional data, such as flow cytometry-derived BCR/TCR affinity landscapes. Future work aims to leverage such high-dimensional data to validate our antibody kinetics model and potentially establish a framework for refining immunological parameters based on evolving BCR/TCR repertoire patterns.

## Supplementary Materials

Supplementary Materials are attached. The MATLAB codes can be accessed through the following link: https://github.com/zhaobinxu23/adaptive_immunity_new.

## Author Contributions

Conceptualization, Z.X.; methodology, Z.X. and D.W.; writing—original draft preparation, Z.X.; writing—review and editing, D.W.; funding acquisition, Z.X. All authors have read and agreed to the published version of the manuscript.

## Institutional Review Board Statement

Not applicable.

## Informed Consent Statement

Not applicable. No wet experiments were performed in this study.

