## supplementary materials for "A Comprehensive Mathematical Model Simulating the Adaptive Immune Response"

**Mathematical analysis of model 3.1.1 :**

$$\frac{dV}{dt} = k_0V - k_1VA + k_2C;$$

$$\frac{dA}{dt} = -k_1VA + k_2C + k_3C - k_4A + \pi;$$

$$\frac{dC}{dt} = k_1VA - k_2C - k_5C;$$

Positive analysis shows that when V = 0 , $\frac{dV}{dt} \geq 0$; when A = 0 , $\frac{dA}{dt} \geq 0$; when C = 0 , $\frac{dC}{dt} \geq 0$.

Therefore, for any positive initial value, the system's positivity can be guaranteed.

Equilibrium analysis, for this simple system, we can easily deduce, for complex systems with more parameters, we will later use the numerical solution method after parameter assignment to solve its equilibrium state. The solution of equilibrium state is an important evaluation criterion for us to judge the rationality of parameter selection. For this system, we calculated two equilibrium states

a disease free equilibrium state (V = 0, A = $\pi/k_4$, C = 0 ), the other is endemic equilibrium balance point ( $V^* = \frac{k_0(k_2+k_5)k_4-\pi k_1k_5}{k_0k_1(k_3-k_5)}$ , $A^* = \frac{k_0(k_2+k_5)}{k_1k_5}$ , $C^* = \frac{k_0V^*}{k_5}$ ). We further used mathematical analysis to study the stability of the two equilibrium states.

The Jacobian matrix of the equilibrium point of Model 3.1.1 is as follows:

$$\begin{matrix} k_0 - k_1A^* & -k_1V^* & k_2 \\ -k_1A^* & -k_1V^* - k_4 & k_2 + k_3 \\ k_1A^* & k_1V^* & -k_2 - k_5 \end{matrix}$$

disease free equilibrium The Jacobian matrix of state is as follows:

$$\begin{matrix} k_0 - k_1\pi/k_4 & 0 & k_2 \\ -k_1\pi/k_4 & -k_4 & k_2 + k_3 \\ k_1\pi/k_4 & 0 & -k_2 - k_5 \end{matrix}$$

$$\lambda_1 = k_0 - \frac{k_1\pi}{k_4}; \lambda_2 = -k_4; \lambda_3 = -k_2 - k_5;$$

Therefore, disease free equilibrium stability depends on the positive or negative nature of $k_0 - \frac{k_1\pi}{k_4}$, when $k_0 < \frac{k_1\pi}{k_4}$ The system will eventually completely eliminate the virus. However, for most biological systems, because $k_1$it is a very small value, $k_0 > \frac{k_1\pi}{k_4}$ will be satisfied in most viral infections, which means that from a purely mathematical point of view, it is impossible to completely eliminate the virus. However, the biological system is not a purely mathematical system. The components in mathematics can approach zero infinitely in dynamic changes, but for biological components, since they exist discontinuously, they cannot approach zero infinitely. Therefore, in the real dynamic process, we will find that even if the disease free equilibrium The state is unstable, but we can still completely eliminate the virus. You will see it in the analysis.

endemic equilibrium

$$\begin{matrix} a & b & c \\ d & e & f \\ g & h & k \end{matrix}$$

$$a = k_0 - k_1 A^*;$$
$$b = -k_1 V^*;$$
$$c = k_2;$$
$$d = -k_1 A^*;$$
$$e = -k_1 V^* - k_4;$$
$$f = k_2 + k_3;$$
$$g = k_1 A^*;$$
$$h = k_1 V^*;$$
$$k = -k_2 - k_5;$$

$\lambda^3 - T\lambda^2 + M\lambda - D = 0$

- **Trace** : T = a+e+k
- **Sum of second-order principal minors** : M=(ae − bd)+( ak − cg)+( ek − fh )
- **Determinant** : D=det(J)=a( ek − fh ) − b(dk − fg )+c(dh − eg )

For a third-order system, the necessary and sufficient condition for the equilibrium point to be asymptotically stable is:

1. Trace is negative: T<0
2. The determinant is negative: D < 0
3. Stability condition: T * M>D

These conditions must be met simultaneously. If any one of them is not met, the equilibrium point is unstable.

Numerical simulations are as follows: When we use the following initial state and parameter combination

| | |
|---|---|
| $V_0$ | 1 |
| $A_0$ | $10^3$ |
| $C_0$ | 0 |
| $k_0$ | 1 |
| $k_1$ | $10^{-7}$ |
| $k_2$ | $10^{-14}$ |

| | |
|---|---|
| $k_3$ | 2 |
| $k_4$ | $10^{-2}$ |
| $k_5$ | 0.5 |
| π | 10 |

Table S1: parameter sets and initial value of model 3.1.1.

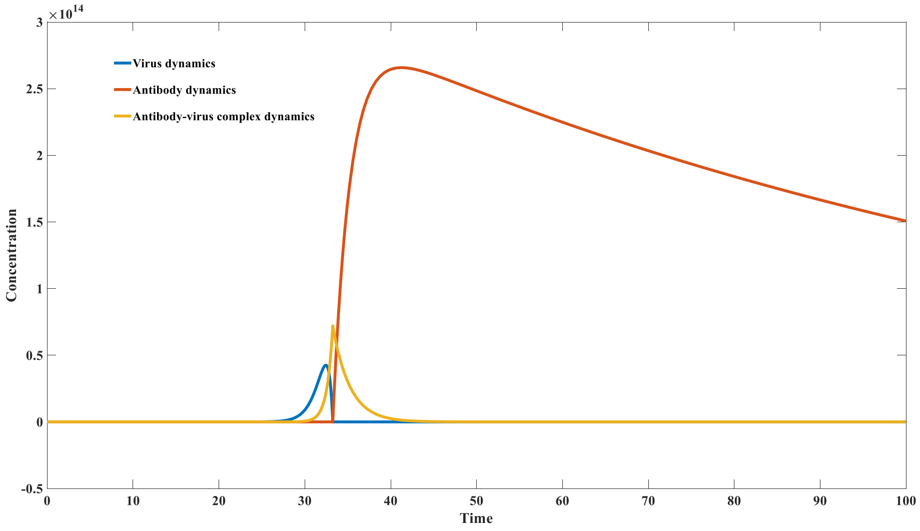


Figure S_1A: Dynamics of antibody - virus interaction（complete virus clearance

The simulation results are shown in Figure S_1A. From the figure, it can be seen that although from a mathematical analysis point of view $k_0 > \frac{k_1 \pi}{k_4}$ , the system cannot reach disease free equilibrium state. Endemic equilibrium stability analysis yields the equilibrium point (V = 33330; A = 107; C = 66660). The three eigenvalues of this equilibrium point are ( $\lambda_1 = -0.5369 + 0.0000i; \lambda_2 = 0.0118 + 0.0958i; \lambda_3 = 0.0118 - 0.0958i$), and the real parts of two eigenvalues are positive. Therefore, theoretically, the system will not be stable at any equilibrium point. However, the discontinuous characteristics of the components in the biological process allow the system to completely eliminate the virus. In the algorithm, we add a very small threshold. When V is less than this threshold, it will become 0, and we can obtain the result shown in Figure S_1A . In other words, the system has completely eliminated the virus and can ultimately achieve disease free state. Theoretically, the system can also reach endemic equilibrium state, for example, when we use the following parameter combination, the system can achieve endemic equilibrium state, the equilibrium point is (V = 5555.55 ; A = 12000000; C = 13333.333 ), and the three eigenvalues of the equilibrium point are ( $\lambda_1 = -0.5369 + 0.0000i; \lambda_2 = -0.0015 + 0.0444i; \lambda_3 = -0.0015 - 0.0444i$ ). The simulation results are shown in Figure S_1B . However, the occurrence of chronic infection in real-world infections cannot be fully determined based on this model, because real-world infections often occur through cell compartments, which can isolate antibodies from the virus. Therefore, we use Model 3.4.1 to explain a plausible mechanism of chronic infection .

| | |
|---|---|
| $V_0$ | 1 |
| $A_0$ | $10^{7}$ |

| | |
|---|---|
| $C_0$ | 0 |
| $k_0$ | 1.2 |
| $k_1$ | $10^{-7}$ |
| $k_2$ | $10^{-14}$ |
| $k_3$ | 2 |
| $k_4$ | $10^{-2}$ |
| $k_5$ | 0.5 |
| $\pi$ | $10^{5}$ |

Table S2: parameter sets and initial value of model 3.1.1.

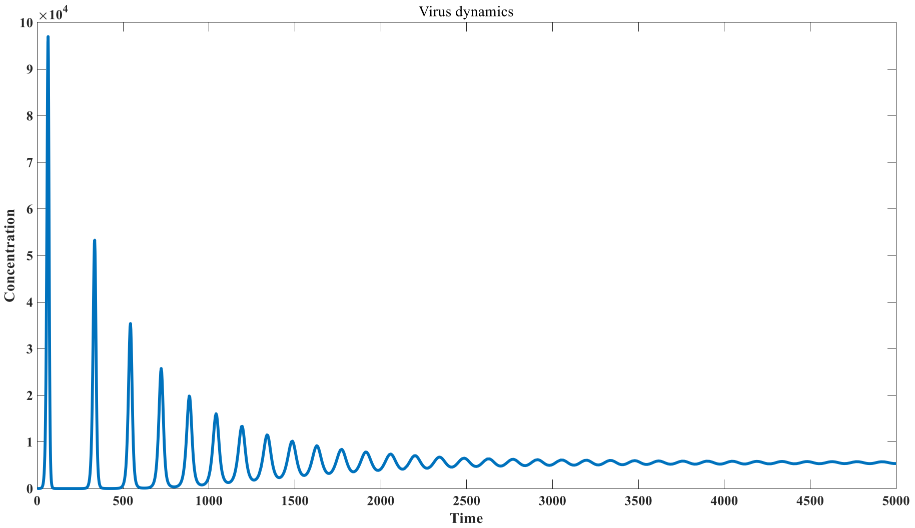


Figure S_1B: D ynamics of antibody - virus interaction (incomplete virus clearance)

There are three critical parameters here, one represents the characteristics of viral replication $k_0$ , and the other two describe the properties of the immune system. The parameter $k_3$ represents the ability of the antibody-antigen complex to feedback and generate antibodies, reflecting the activity of CD4 +T cells, which will gradually decline with aging. The third parameter $k_5$represents the degradation rate of the antigen-antibody complex, which is closely related to the number and activity of NK cells, and also shows significant individual differences. We focus on the relationship between the severity of infection and these three parameters. We specifically proposed the concept of antibody exhaustion, which means that when the virus replication coefficient $k_0$ is too large, or a small $k_3$, or a large $k_5$, the proliferation of antibodies will be affected. In this case, the proliferation rate of antibodies will be lower than the proliferation rate of viruses, resulting in the regeneration of antibodies being slower than the consumption of antibodies, thereby leading to the exhaustion of specific antibodies, which is reflected in the phenomenon of germinal center proliferation defects in severely infected patients. At this time, stability analysis shows that the system is in the disease free equilibrium state and endemic equilibrium states are all unstable. For example, when we use the following parameter combination, antibody exhaustion is likely to occur, leading to unlimited viral proliferation, as shown in Figures S_1C, S_1D, and S_1E .

| | | | |
|---|---|---|---|
| $V_0$ | 1 | 1 | 1 |
| $A_0$ | $10^3$ | $10^3$ | $10^3$ |
| $C_0$ | 0 | 0 | 0 |
| $k_0$ | 1 | 2 | 1 |
| $k_1$ | $10^{-7}$ | $10^{-7}$ | $10^{-7}$ |
| $k_2$ | $10^{-14}$ | $10^{-14}$ | $10^{-14}$ |
| $k_3$ | 2 | 2 | 1 |
| $k_4$ | $10^{-2}$ | $10^{-2}$ | $10^{-2}$ |
| $k_5$ | 1 | 0.5 | 0.5 |
| $\pi$ | $10^{4}$ | $10^{4}$ | $10^{4}$ |

Table S3: parameter sets and initial value with multiple antibodies.

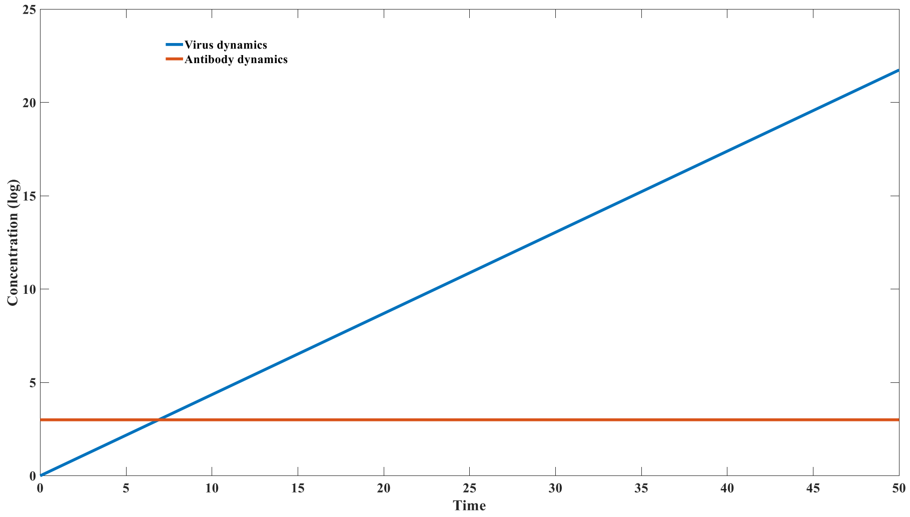


Figure S_1C: D efective antibody production due to fast clearance rate of antibody - virus complex( $k_5 = 1$)

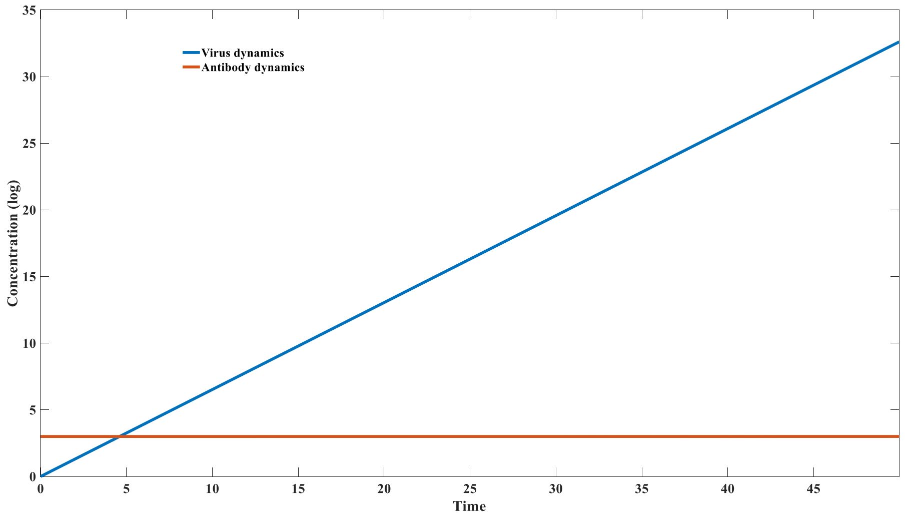


Figure S_1D: D efective antibody production due to fast virus replication rate( $k_0 = 1.5$)

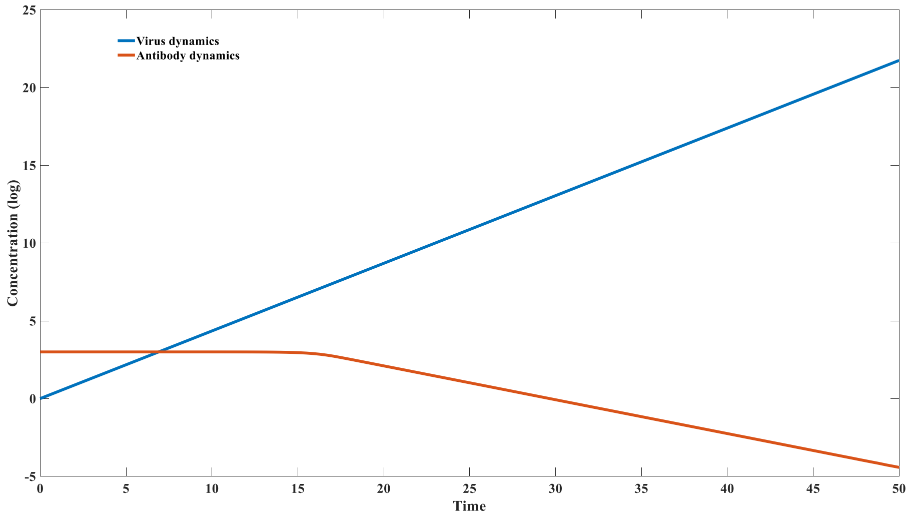


Figure S_1E: Defective antibody production due to slow antibody regeneration rate( $k_3 = 1$)

**Mathematical analysis and parameter setting of model 3.1.2**

One critical drawback in model 3.1.1 is that it cannot achieve immune memory. Especially when the virus is completely cleared, the specific neutralizing antibodies return to the initial disease-free equilibrium state without permanent upregulation in concentration. This doesn't occur in real infections. In real infections, memory B cells persist for a very long time, allowing specific IgG levels to remain elevated for extended periods, resulting in sustained protection. To explain this phenomenon, we introduced the concept of environmental antigens and reconstructed model 3.1.2 based on this concept.

$$\frac{dV}{dt} = k_0V - k_1VA + k_2C;$$

$$\frac{dA}{dt} = -k_1VA + k_2C + k_3C - k_4A - p_1EA + p_2Q + p_3Q;$$

$$\frac{dC}{dt} = k_1VA - k_2C - k_5C;$$

$$\frac{dE}{dt} = \pi - p_1EA + p_2Q;$$

$$\frac{dQ}{dt} = p_1EA - p_2Q - p_4Q;$$

One of the most significant differences between Model 3.1.2 and 3.1.1 is that it doesn't consider the supply of antibodies to be a fixed value, but rather a dynamic process driven by the interaction of environmental antigens and antibodies. In the absence of viruses, the maintenance of antibody homeostasis depends on interactions with environmental antigens. Here, $-p_1EA$ represents the process by which antibodies bind to environmental antigens E to form complex Q, $p_2Q$ represents the process by which complexes dissociate to generate antibodies, and most importantly $p_3Q$ represents the process by which complexes feed back to promote antibody regeneration. Positive analysis reveals that when V = 0 , $\frac{dV}{dt} \geq 0$; when

A = 0 , $\frac{dA}{dt} \geq 0$; when C = 0 , $\frac{dC}{dt} \geq 0$;when E = 0 , $\frac{dE}{dt} \geq 0$; when Q = 0 , $\frac{dQ}{dt} \geq 0$; Therefore, for any positive initial value, the system's positivity can be guaranteed. When the virus is 0, the system becomes:

$$\frac{dA}{dt} = -k_4A - p_1EA + p_2Q + p_3Q;$$

$$\frac{dE}{dt} = \pi - p_1EA + p_2Q;$$

$$\frac{dQ}{dt} = p_1EA - p_2Q - p_4Q;$$

We can calculate the initial state of the system based on the above equations. According to the following parameter combination, the initial state of the system is ( $A_0 = 10^7; E_0 = 10^8; Q_0 = 2 * 10^5$). The Jacobian matrix at the equilibrium point is as follows:

$$\begin{matrix} k_0 - k_1A^* & -k_1V^* & k_2 & 0 & 0 \\ -k_1A^* & -k_1V^* - k_4 - p_1E^* & k_2 + k_3 & -p_1A^* & p_2 + p_3 \\ k_1A^* & k_1V^* & -k_2 - k_5 & 0 & 0 \\ 0 & -p_1E^* & 0 & -p_1A^* & p_2 \\ 0 & p_1E^* & 0 & p_1A^* & -p_2 - p_4 \end{matrix}$$

The system also has two equilibrium states, namely disease-free equilibrium state and endemic equilibrium state. For a given parameter combination, we can also determine the stability of the equilibrium point. For example, when the system uses the following parameter combination, disease-free equilibrium state is ( $A^* = 10^7; E^* = 10^8; Q^* = 2 * 10^5; V^* = 0; C^* = 0$)and the eigenvalue at this equilibrium point is ( $\lambda_1 = 0.9; \lambda_2 = -0.5197; \lambda_3 = -0.5$; $\lambda_4 = 0.0087; \lambda_5 = 4.1 * 10^{-18}$ ). Endemic equilibrium state is ( $A^* = 10^8; E^* = 10^7; Q^* = 2 * 10^5; V^* = 3 * 10^5; C^* = 6 * 10^5$), and the eigenvalue value at this equilibrium point is ( $\lambda_1 = -0.5353 + 0.0000i; \lambda_2 = -0.5000 + 0.0000i; \lambda_3 = 0.0111 + 0.0911i$ ; $\lambda_4 = 0.0111 - 0.0911i; \lambda_5 = -0.0100 + 0.0000i$). The numerical simulation is shown in Figure S2. Although pure mathematical analysis shows that the system is in disease-free state is unstable, but when discontinuous constraints are added, the system can finally reach disease-free state with complete virus clearance.

| | |
|---|---|
| $V_0$ | 1 |
| $A_0$ | $10^7$ |
| $C_0$ | 0 |
| $E_0$ | $10^8$ |
| $Q_0$ | $2* 10^5$ |
| $k_0$ | 1 |
| $k_1$ | $10^{-8}$ |
| $k_2$ | $10^{-14}$ |
| $k_3$ | 2 |
| $k_4$ | $10^{-2}$ |

| | |
|---|---|
| $k_5$ | 0.5 |
| $\pi$ | $10^4$ |
| $p_1$ | $10^{-10}$ |
| $p_2$ | 0 |
| $p_3$ | 1 |
| $p_4$ | 0.5 |

Table S4: parameter sets and initial value in model 3.1.2.

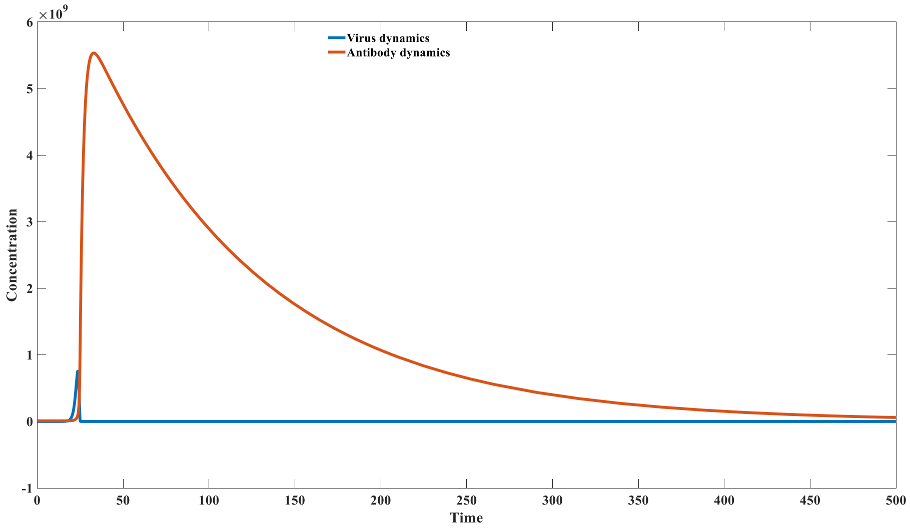


Figure S_2: Virus - antibody interaction dynamics using model 3.1.2

**Mathematical analysis and parameter setting of model 3.1.3**

Model 3.1.2 alone cannot effectively explain the issue of immune memory without the consideration of antibody diversity. When we take antibody diversity into consideration, we can use the updated model 3.1.3 to well explain the occurrence of antibody clonal expansion and the mechanism of immune memory. Model 3.1.3 still belongs to the category of simple models, but it can help us study the formation mechanism of immune memory more intuitively. The significance of the existence of environmental antigens is that they can maintain the stability of the overall antibodies, because their total supply $\pi$ is a fixed quantity, it can maintain a constant level of both its own and its total antibody content in the disease-free state. Because environmental antigens E are a broad category encompassing all different types of environmental antigens, the overall kinetic coefficients $p_1$ and $p_2$ do not vary depending on the antibody type. However, the kinetic coefficients $k_1$ and $k_2$ do exhibit $k_2$diverse differences. Model 3.1.3 is expressed as follows:

$$\frac{dV}{dt} = k_0 V - \sum_{n}^{i=1} k_{1i} V A_i + \sum_{n}^{i=1} k_{2i} C_i \,;$$

$$\frac{dA_i}{dt} = -k_{1i} V A_i + k_{2i} C_i + k_3 C_i - k_4 A_i - p_1 E A_i + p_2 Q_i + p_3 Q_i;$$

$$\frac{dC_i}{dt} = k_{1i}VA_i - k_{2i}C_i - k_5C_i;$$

$$\frac{dE}{dt} = \pi - \sum_{n}^{i=1} p_1EA_i + \sum_{n}^{i=1} p_2Q_i\,;$$

$$\frac{dQ_i}{dt} = p_1EA_i - p_2Q_i - p_4Q_i;$$

We simulated the changes of 4 different types of antibodies, and the specific parameter settings are as follows:

| | Antibody 1 | Antibody 2 | Antibody 3 | Antibody 4 |
|---|---|---|---|---|
| $V_0$ | 1 | 1 | 1 | 1 |
| $A_0$ | $2.5*10^6$ | $2.5*10^6$ | $2.5*10^6$ | $2.5*10^6$ |
| $C_0$ | 0 | 0 | 0 | 0 |
| $E_0$ | $2.5*10^7$ | $2.5*10^7$ | $2.5*10^7$ | $2.5*10^7$ |
| $Q_0$ | $5*10^4$ | $5*10^4$ | $5*10^4$ | $5*10^4$ |
| $k_0$ | 1 | 1 | 1 | 1 |
| $k_1$ | $10^{-8}$ | $10^{-8}$ | $0.9*10^{-8}$ | $0.8*10^{-8}$ |
| $k_2$ | $10^{-14}$ | $10^{0}$ | $0.9*10^{-14}$ | $0.8*10^{-14}$ |
| $k_3$ | 2 | 2 | 2 | 2 |
| $k_4$ | $10^{-2}$ | $10^{-2}$ | $10^{-2}$ | $10^{-2}$ |
| $k_5$ | 0.5 | 0.5 | 0.5 | 0.5 |
| $\pi$ | $10^4$ | $10^4$ | $10^4$ | $10^4$ |
| $p_1$ | $10^{-10}$ | $10^{-10}$ | $10^{-10}$ | $10^{-10}$ |
| $p_2$ | 0 | 0 | 0 | 0 |
| $p_3$ | 1 | 1 | 1 | 1 |
| $p_4$ | 0.5 | 0.5 | 0.5 | 0.5 |

Table S5: parameter sets and initial value in model 3.1.3.

The simulation results are shown in Figure S3. From the figure, it can be observed that, under conditions of identical forward binding coefficients, antibodies with lower dissociation coefficients (Antibody 1) proliferate faster than those with higher dissociation coefficients (Antibody 2). This indicates that antibodies with stronger binding affinity can achieve more rapid selective clonal expansion.

For antibodies with equivalent binding energies (Antibody 1 has the same Kd value as Antibodies 3 and 4), those with higher forward binding coefficients exhibit faster proliferation rates (Antibody 1 > Antibody 3 > Antibody 4). Moreover, the ultimate outcome of this antibody proliferation permanently alters the proportions of antibodies; as illustrated, in the initial state, the proportions of various antibodies are all 25%. After infection, Antibody 1 stabilizes permanently at 38%, Antibody 2 decreases to 8%, Antibody 3 changes to 31%, and Antibody 4 becomes 24%. Following infection, the total amount of antibodies will tend to stabilize, equaling the sum of the initial antibody concentrations. The permanent alteration of the antibody repertoire is significantly influenced by self-antigen substances.

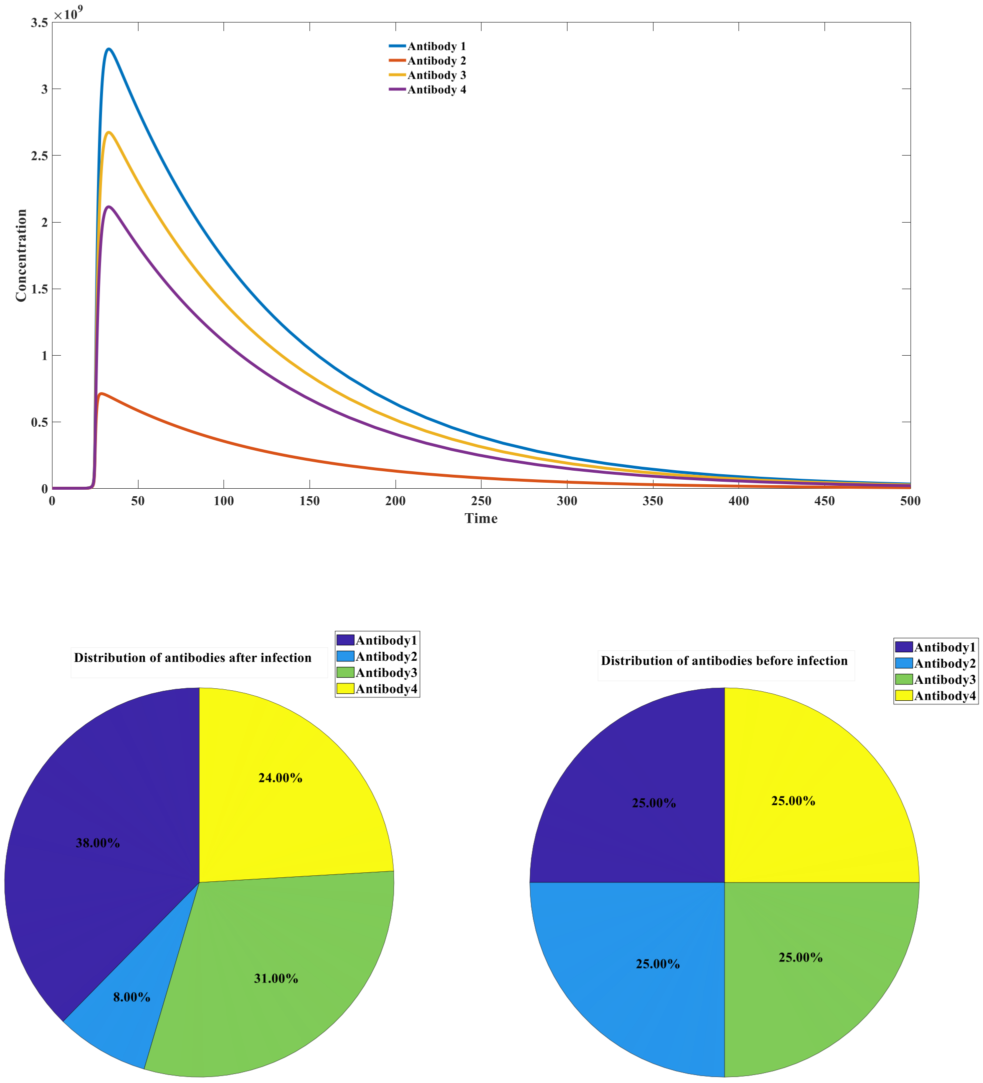


Figure S_3: Simulation of antibody dynamics using model 3.1.3

### Parameter settings and simulation results for model 3.1.4

Model 3.1.4 is as follows:

$$\frac{dV}{dt} = k_0 V - k_1 VA + k_2 C - k_1 VA_2 + k_2 C_2;$$

$$\frac{dA}{dt} = -k_1 VA + k_2 C + k_3 C;$$

$$\frac{dC}{dt} = k_1 VA - k_2 C - k_5 C;$$

$$\frac{dA_2}{dt} = -k_1 VA_2 + k_2 C_2 - k_6 A_2 + \pi;$$

$$\frac{dC_2}{dt} = k_1 VA_2 - k_2 C_2 - k_5 C_2;$$

We simulated the effects of antibody addition time and dosage on viral dynamics. The specific simulation parameters were set as follows:

| | |
|---|---|
| $V_0$ | 1 |
| $A_0$ | 10 |
| $C_0$ | 0 |
| $A2_0$ | 0 |
| $C2_0$ | 0 |
| $k_0$ | 0.5 |
| $k_1$ | $10^{-5}$ |
| $k_2$ | $10^{-14}$ |
| $k_3$ | 1.5 |
| $k_5$ | 0.5 |
| $k_6$ | 0.05 |
| π | $1.5 * 10^{6}$ |

Table S6: parameter sets and initial value in model 3.1.4.

The time of adding the antibody is ( 21 to 30 time point, the dose was $1.5 * 10^{6}$), the results are shown in Figure S4A . It can be seen from the figure that the earlier the monoclonal antibody is added, the better the treatment effect is, and the lower the peak concentration of the virus would be. Antibodies added before time point 28 and 29 will not trigger viral rebound . Adding monoclonal antibodies at this point will cause viral rebound, while adding monoclonal antibodies later will not cause viral rebound.

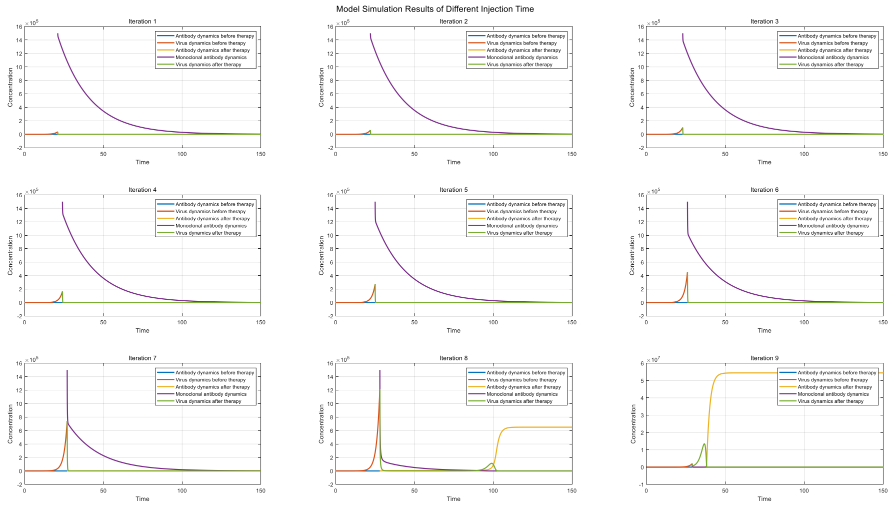


Figure S_4A: Simulation of H ost -virus Interaction at different injection time of monoclonal antibody

We further simulated the effect of antibody dosage on viral dynamics. We simulated the 28 th time point Virus kinetic curves with different doses of antibodies ($0.5 * 10^{6}$ to $4.5 * 10^{6}$). The results are shown in Figure S4B. As can be seen, viral rebound occurs within a certain range of monoclonal antibody doses. Below the minimum threshold, viral proliferation is unaffected. As shown in the figure, at a dose of $0.5 \times 10^{6}$ , viral growth is barely affected. At a dose of $10^{6}$, viral concentrations experience a short-term drop, followed by a strong rebound to a high point. At a dose of 1.5 $\times 10^{6}$ ' viral concentrations experience a significant drop, but then rebound significantly after a longer period. When the dose exceeds the maximum threshold, viral rebound does not occur.

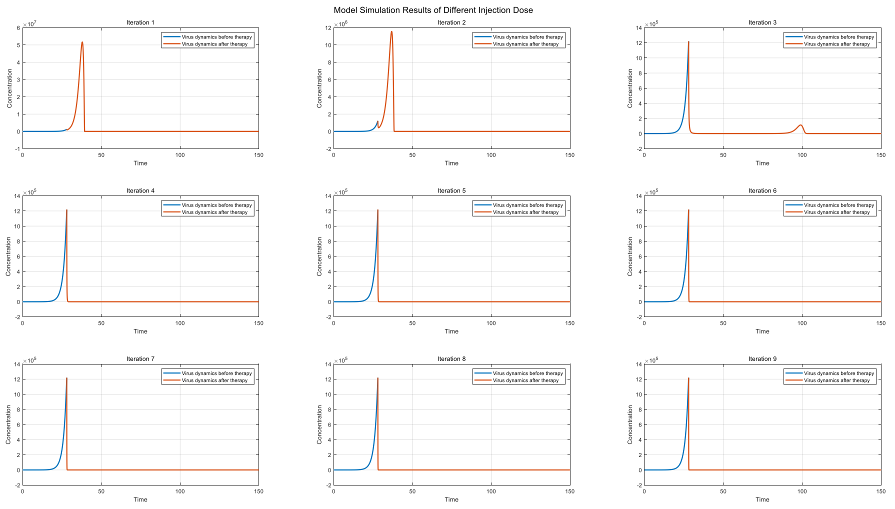


Figure S_4B: Simulation of H ost -virus Interaction at different injection Dose of monoclonal antibody

Alan Perelson 's model believes that the viral rebound phenomenon in monoclonal antibody therapy is caused by the generation of mutant strains. Their model is as follows:

$$\frac{\mathrm{dT}}{\mathrm{dt}} = -\beta_1 V_1 \mathrm{T} - \beta_2 V_2 \mathrm{T} + \gamma \mathrm{T}(1 - \frac{E_1 + E_2 + I_1 + I_2 + T}{T_M});$$

$$\frac{\mathrm{d}E_1}{\mathrm{dt}} = -\beta_1 V_1 \mathrm{T} - \mathrm{k}E_1;$$

$$\frac{\mathrm{d}E_2}{\mathrm{dt}} = -\beta_2 V_2 \mathrm{T} - \mathrm{k}E_2;$$

$$\frac{\mathrm{d}I_1}{\mathrm{dt}} = \mathrm{k}E_1 - \delta(\mathrm{t})I_1;$$

$$\frac{\mathrm{d}I_2}{\mathrm{dt}} = \mathrm{k}E_2 - \delta(\mathrm{t})I_2;$$

$$\frac{\mathrm{d}V_1}{\mathrm{dt}} = \pi(1-\mu)\mathrm{I}_1 - \mathrm{k}_{\mathrm{on},1}\mathrm{V}_1\mathrm{A} + \mathrm{k}_{\mathrm{off},1}\mathrm{C}_1 - \mathrm{cV}_1;$$

$$\frac{\mathrm{d}C_1}{\mathrm{dt}} = \mathrm{k}_{\mathrm{on},1}\mathrm{V}_1\mathrm{A} - \mathrm{k}_{\mathrm{off},1}\mathrm{C}_1 - \gamma_1\mathrm{C}_1;$$

$$\frac{\mathrm{d}V_2}{\mathrm{dt}} = \pi\mathrm{I}_2 + \pi\mu\mathrm{I}_1 - \mathrm{k}_{\mathrm{on},2}\mathrm{V}_2\mathrm{A} + \mathrm{k}_{\mathrm{off},2}\mathrm{C}_2 - \mathrm{cV}_2;$$

$$\frac{\mathrm{d}C_1}{\mathrm{dt}} = \mathrm{k}_{\mathrm{on},2}\mathrm{V}_2\mathrm{A} - \mathrm{k}_{\mathrm{off},2}\mathrm{C}_2 - \gamma_2\mathrm{C}_2;$$

$$\delta(\mathrm{t}) = \begin{cases} \delta, \mathrm{t} < \mathrm{t}^* \\ \delta_M - (\delta_M - \delta)e^{-\sigma(\mathrm{t}-\mathrm{t}^*)}, t \geq \mathrm{t}^* \end{cases};$$

$$A(t) = \begin{cases} 0, t < t_{inf} \\ \frac{A_{max}}{\Delta t}(t - t_{inf}), t_{inf} \le t < t_{inf} + \Delta t \\ A_{max} e^{-\alpha(t-(t_{inf}+\Delta t))}, t \ge t_{inf} + \Delta t \end{cases};$$

The specific meanings of each parameter can be found in the paper. We must point out the flaws of this model. The first is the use of an incorrect term for the depletion of susceptible cells $-\beta_1 V_1 T$. Because the sizes of viruses and cells are very unequal, a cell can be invaded and infected by multiple viruses simultaneously. Therefore, using a quadratic term to represent the rate of infection is inappropriate and will lead to accelerated depletion of susceptible cells. A second important flaw is that it ignores the role of autoantibodies and exogenously injected antibodies. Although both can specifically bind to the virus, it can be assumed that some autoantibodies are B cell BCRs. Therefore, autoantibody binding to antigen can lead to antibody regeneration, while exogenous neutralizing antibodies do not. Third, this model does not correctly simulate the dynamics of autoantibody production in response to viral infection, instead using several piecewise functions to approximate antibody dynamics. A fourth issue is the inadequate typing of autoantibodies and exogenous antibodies. The binding affinity of autoantibodies does not change with viral mutations, as high-affinity autoantibodies are selected by the virus. However, the affinity of exogenous monoclonal antibodies may be significantly reduced against mutant strains. Therefore, they observed that the phenomenon of viral rebound in patients infected with mutant strains is not caused by viral mutation during infection. A more likely explanation is that the mutated virus infects the patient, rather than completing the viral mutation in the patient's body. The affinity of this monoclonal antibody against mutant strains will show a significant decrease , so when the dose is insufficient, viral rebound is more likely to occur. At the 28th time after infection, the virus is more likely to rebound. The monoclonal antibody was injected into the point. The specific simulation parameters were set as follows, and the simulation results are shown in Figure S_4C .

| | Original strain | M utant strain |
|---|---|---|
| $V_0$ | 1 | 1 |
| $A_0$ | 10 | 10 |
| $C_0$ | 0 | 0 |
| $A2_0$ | 0 | 0 |
| $C2_0$ | 0 | 0 |
| $k_0$ | 0.5 | 0.5 |
| $k_1$ | $10^{-5}$ | $10^{-5}$ |
| $k_2$ | $10^{-14}$ | $10^{-14}$ |
| $k_1'$ (represents the positive binding constant of monoclonal antibody to | $10^{-5}$ | $10^{-6}$ |

| | | |
|---|---|---|
| virus ) | | |
| $k_2'$ (represents the negative dissociation constant of monoclonal antibody against virus ) | $10^{-14}$ | $10^{-14}$ |
| $k_3$ | 1.5 | 1.5 |
| $k_5$ | 0.5 | 0.5 |
| $k_6$ | 0.05 | 0.05 |
| $\pi$ | $2*10^{6}$ | $2*10^{6}$ |

Table S7: parameter sets and initial values in monoclonal antibody therapy induced virus rebound model.

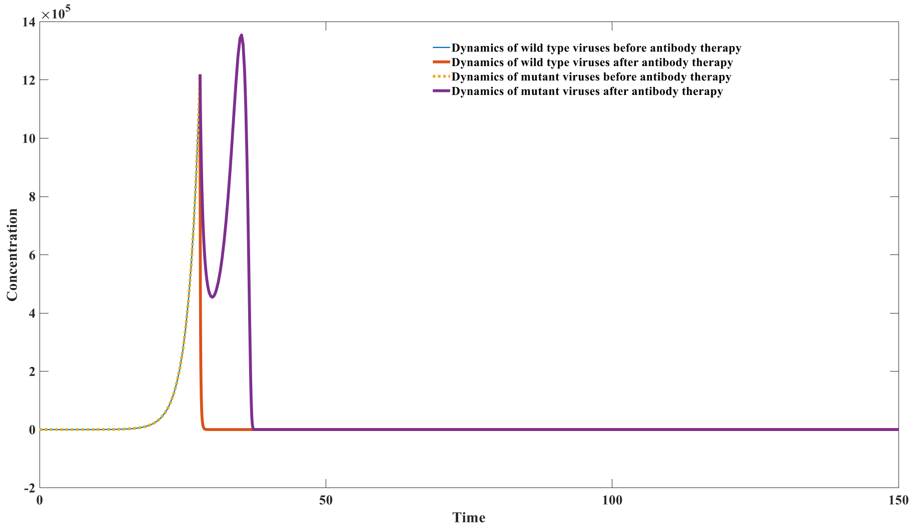


Figure S_4C: Simulation of monoclonal antibody therapy toward wild type virus and mutant strain

**Parameter settings and simulation results for model 3.1.5**

Model 3.1.5 is as follows:

$$\frac{dV}{dt} = k_0V - k_1VA + k_2C + k_0k_7C_2 - p_1VD + p_2C_2;$$

$$\frac{dA}{dt} = -k_1VA + k_2C + k_3C - k_1C_2A + k_2C_3 + k_3C_3 - k_4A + \pi;$$

$$\frac{dC}{dt} = k_1VA - k_2C - k_5C;$$

$$\frac{dC_2}{dt} = p_1VD - p_2C_2 - k_1C_2A + k_2C_3;$$

$$\frac{dC_3}{dt} = k_1C_2A - k_2C_3 - k_5C_3;$$

$$\frac{dD}{dt} = -p_1VD + p_2C_2 - k_6D + \pi_1;$$

We simulated the effects of the addition time and dosage of small molecule viral inhibitors on viral dynamics. The specific simulation parameters were set as follows:

| | |
|---|---|
| $V_0$ | 1 |
| $A_0$ | 1000 |
| $C_0$ | 0 |
| $C_{2_0}$ | 0 |
| $C_{3_0}$ | 0 |
| D | 0 |
| $k_0$ | 1 |
| $k_1$ | $10^{-7}$ |
| $k_2$ | $10^{-14}$ |
| $k_3$ | 2 |
| $k_4$ | 0.01 |
| $k_5$ | 0.5 |
| $k_6$ | 0.01 |
| $k_7$ | 0.1 |
| $\pi$ | 10 |
| $\pi_1$ | 0 |
| $p_1$ | $10^{-7}$ |
| $p_2$ | $10^{-14}$ |

Table S8: parameter sets and initial value in model 3.1.5.

The addition time of small molecule inhibitors was (16 to 24 time point, the dose was $7*10^8$), the results are shown in Figure S5A. It can be seen from the figure that good virus control is accompany with earlier drug administration. The use of small molecule inhibitors before the 20th time point will not cause significant viral rebound. The use of small molecule inhibitors at this point will lead to a significant rebound of the virus. Taking small molecule inhibitors later will not lead to a rebound of the virus, but will lead to an increase in the peak concentration of the virus, which will aggravate the symptoms of infection.

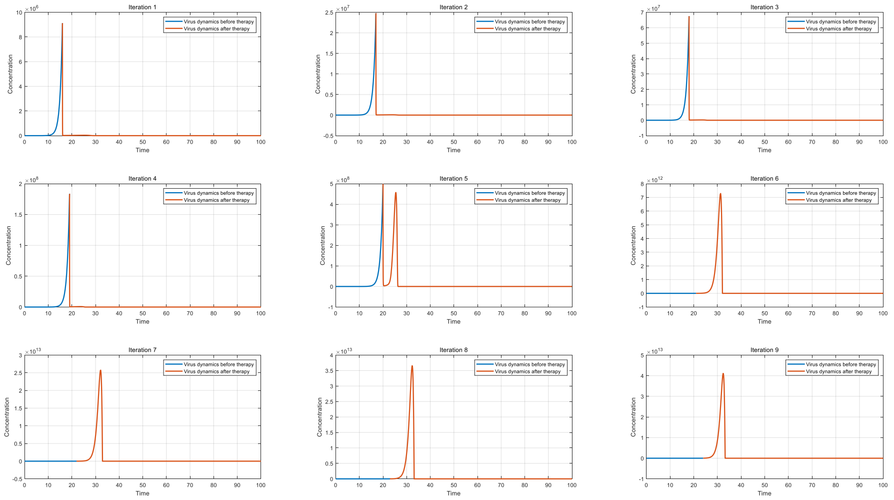


Figure S_5A: V iruses dynamics of different drug injection time

We further simulated the effect of small molecule inhibitor dosage on viral dynamics. Specifically, a gradient of drug dosage ($1*10^{8}$ to $12*10^{8}$) is administrated in-silicon to patients at 20 time point. The results are shown in Figure S5_B. As can be seen from the figure, viral rebound occurs within a certain dose range of the small molecule inhibitor. As shown in the figure, the peak viral concentration shows a significant downward trend with increasing dose. At a dose of $6\times10^{8}$, the viral concentration experiences a short-term drop, followed by a strong rebound to a high point. Further increases in the drug dose continue to reduce the peak viral concentration, and when the dose exceeds $10\times10^{8}$, the viral rebound phenomenon no longer occurs.

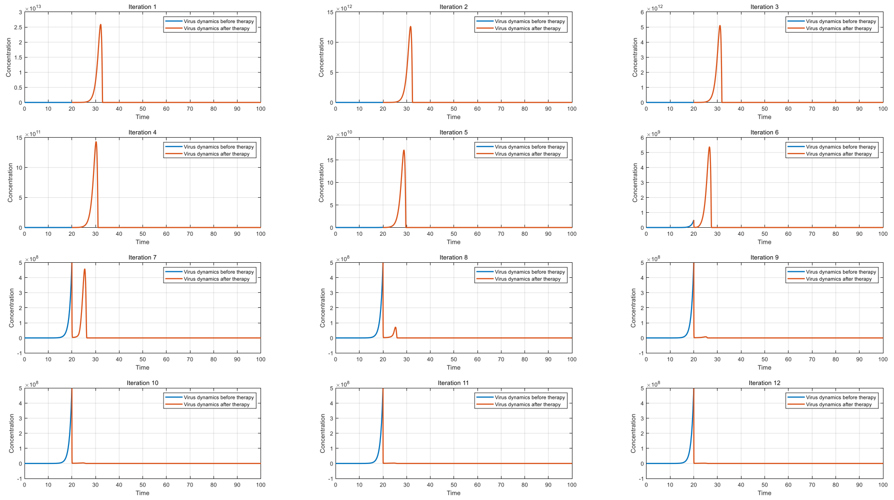


Figure S_5B: V iruses dynamics of different drug injection doses

**Parameter settings and simulation results for model 3.1.6**

Model 3.1.6 is as follows:

$$\frac{dV}{dt} = k_0V - k_1VA + k_2C_1 - k_1VB + k_2C_2;$$

$$\frac{dA}{dt} = -k_1VA + k_2C_1 - k_4A + k_5P;$$

$$\frac{dB}{dt} = -k_1VB + k_2C + k_3(1-\alpha)\frac{C_2}{(C_2+B)}C_2 - k_6B;$$

$$\frac{dP}{dt} = k_3\alpha\beta\frac{C_2}{(C_2+B)}C_2 - k_7P;$$

$$\frac{dC_1}{dt} = k_1VA - k_2C_1 - k_8C_1;$$

$$\frac{dC_2}{dt} = k_1VB - k_2C_2 - k_8C_2;$$

For vaccines, $k_0$it is a negative value, indicating that the viral antigen cannot replicate itself and will gradually degrade in body fluids. For viral infection, $k_0$it is a positive value. We simulated the effects of vaccination dose and vaccination interval on antibody dynamics. The specific parameters are as follows:

| | |
|---|---|
| $V_0$ | $10^8$ |
| $A_0$ | 0 |
| $B_0$ | 500 |
| $P_0$ | 0 |
| $C_{1_0}$ | 0 |
| $C_{2_0}$ | 0 |
| $k_0$ | -0.02 |
| $k_1$ | $10^{-7}$ |
| $k_2$ | $10^{-14}$ |
| $k_3$ | 2 |
| $k_4$ | 0.02 |
| $k_5$ | $10^5$ |
| $k_6$ | 0.005 |
| $k_7$ | 0.01 |
| $k_8$ | 0.5 |
| $\alpha$ | 0.5 |
| $\beta$ | $10^{-3}$ |

Table S9: parameter sets and initial value in model 3.1.6.

Both the first and second vaccination doses were set at $10^8$, with intervals varying from 50 to 600 time points (in increments of 50). The results are illustrated in Figure S_6A. It is evident from the figure that when the vaccination interval is set at 50 time points, there is a transient decline in antibody levels, followed by an upward trend. In contrast, when the intervals range from 100 to 250 time points, a significant decrease in antibody levels is observed after the second vaccination. As the interval continues to increase, starting from 300 time points, there is a notable resurgence in antibody concentration following the second vaccination. Moreover, from 450 time points onward, the peak antibody levels generated by the second vaccination surpass those produced by the initial vaccination. This finding highlights the impact of antibody interference on vaccination strategies, indicating that shorter vaccination intervals may actually accelerate the rate of antibody decay following the second dose.

We further investigated the influence of the second vaccination dose on antibody kinetics by administering the second vaccination at 100 time units, with doses ranging from $10^6$ to $10^{10}$, increasing incrementally by $10^{0.5}$. The results are shown in Figure S_6B. From the figure, it can be observed that when the second vaccination dose is extremely low, it does not significantly affect antibody dynamics. However, as the dose increases, the presence of antibody interference leads to an accelerated decline in antibody levels. When the vaccination dose is increased further, the effects of antibody interference diminish, allowing for a robust secondary response characterized by a strong boost in antibody production.

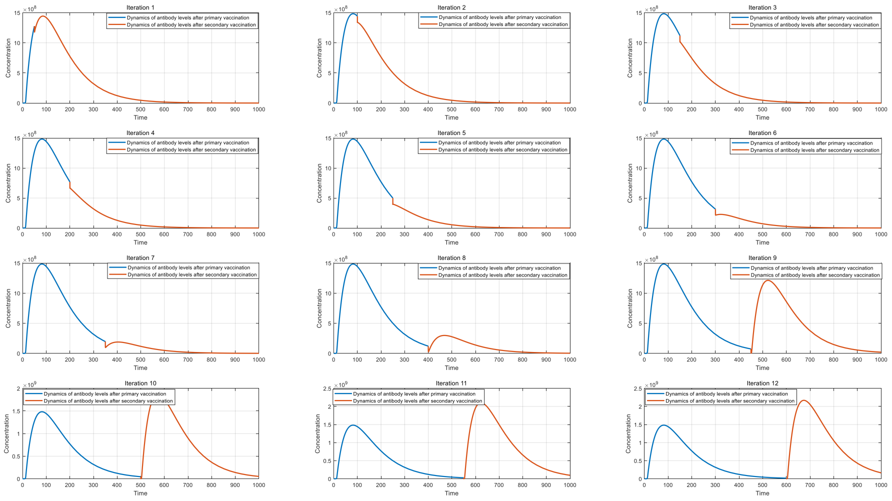


Figure S_6A: Antibody dynamics at different vaccination interval

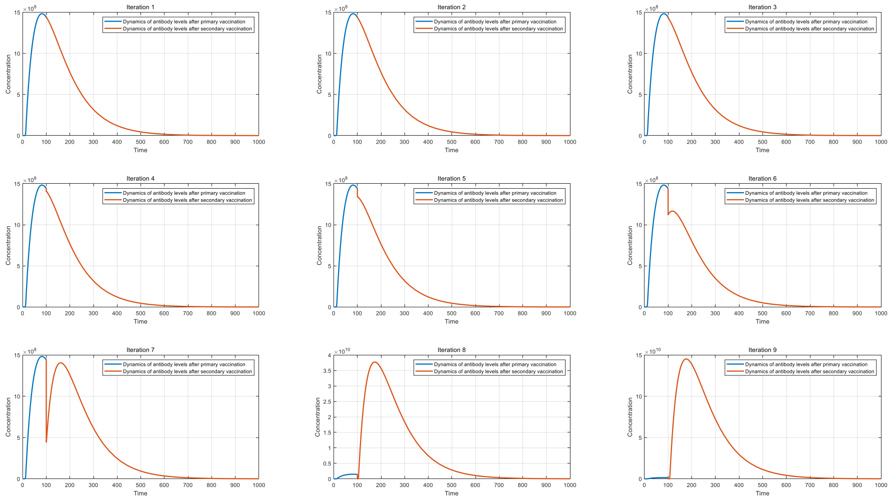


Figure S_6B: Antibody dynamics at different secondary vaccination dosage

Based on model 3.1.6, we further investigated the effect of the parameter α on the process of humoral immunity. The value of α ranges from 0 to 1. When α is excessively small, the conversion of non-plasma B cells into antibody-secreting cells (ASCs) is impaired, which subsequently affects antibody proliferation and facilitates viral replication. Conversely, when α exceeds a certain threshold, antibody proliferation is also adversely impacted. This occurs because the proliferation of non-plasma B cells becomes constrained, ultimately leading to a reduction in the number of conversions to ASCs.

We simulated viral dynamics under varying conditions of α, specifically ranging from 0.05 to 0.45 in increments of 0.05. The results are illustrated in Figure S_6C, with the following parameters set.

| | |
|---|---|
| $V_0$ | 1 |
| $A_0$ | 0 |
| $B_0$ | 500 |
| $P_0$ | 0 |
| $C_{1_0}$ | 0 |
| $C_{2_0}$ | 0 |
| $k_0$ | 0.5 |
| $k_1$ | $10^{-7}$ |
| $k_2$ | $10^{-14}$ |
| $k_3$ | 2 |
| $k_4$ | 0.02 |
| $k_5$ | $10^{5}$ |
| $k_6$ | 0.005 |
| $k_7$ | 0.01 |
| $k_8$ | 0.5 |
| $\beta$ | $10^{-3}$ |

Table S10: parameter sets and initial value in updated model 3.1.6.

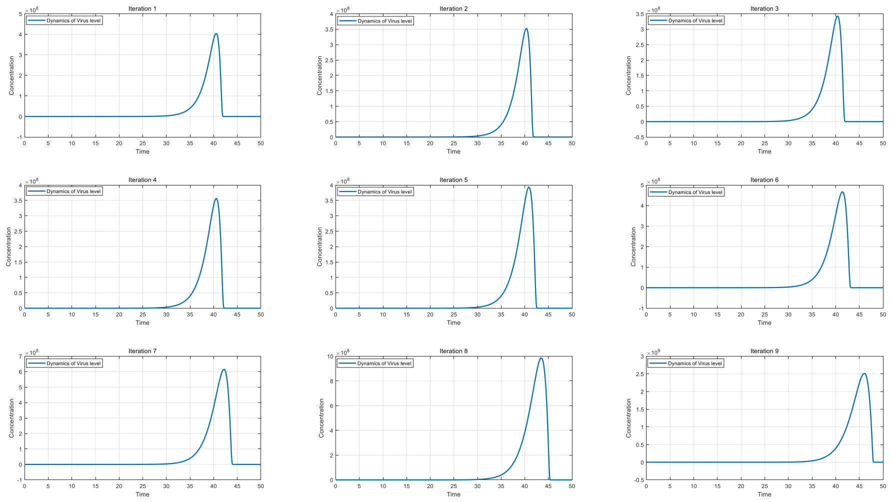

Figure S_6C: Virus dynamics at different α value

From Figure S_6C, it can be observed that as α increases, the peak viral concentration initially decreases before subsequently rising. The optimal value of α for controlling viral replication is influenced by both the viral proliferation coefficient ( $k_0$ ) and the antibody regeneration coefficient ($k_3$ ). This relationship is illustrated in Figure S_6D. The figure shows that the optimal α value for inhibiting viral replication significantly increases with an increase in the antibody regeneration coefficient ($k_3$), while it gradually decreases with an increase in the viral proliferation coefficient ( $k_0$ ).

This finding highlights two key points: First, in populations with strong immunity, a greater number of germinal centers facilitating the conversion of non-plasma B cells to ASCs will enhance resistance to infections. This scenario also applies to low-pathogenic (low replicative activity) viruses. Conversely, for populations with weaker immunity facing high-pathogenic viral infections, reducing the conversion rate to ASCs may provide better protection against viral infection.

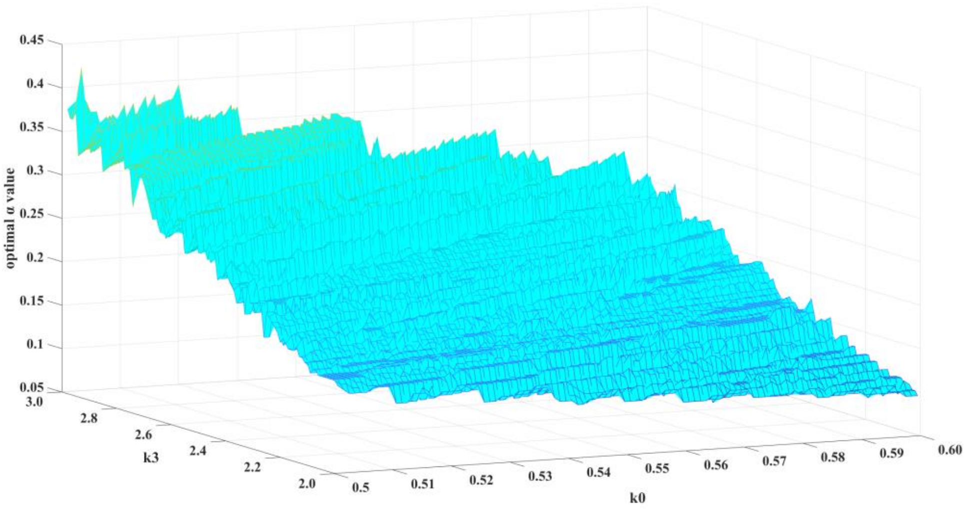

Figure S_6D: Distribution of optimal $\alpha$ value at different $k_0$and $k_3$ combinations

As in model 3.1.1 , for severe patients, there is a combination of $\alpha$, $k_0$and $k_3$that allows the virus to proliferate indefinitely, and the regeneration of antibodies is slower than the exhaustion rate of antibodies, resulting in antibody exhaustion and loss of germinal center activity. Excessively large $k_0$and small $k_3$values can lead to this phenomenon. We can also draw such a conclusion through equilibrium point analysis (disease free equilibrium state and endemic equilibrium state is in an unstable state), numerical When the simulation parameters are set as follows, antibody exhaustion occurs, as shown in Figure S6E . As can be seen in Figure S6E , when the $k_3$ value is relatively small and the $\alpha$ value is relatively large (0.5 ) , infected individuals will experience antibody exhaustion against a strongly replicating virus ( $k_0 = 0.55$). At this time, the virus concentration continues to increase while the antibody concentration remains at a very low point, indicating a regenerative impairment of antibodies in the germinal center.

Similar to model 3.1.1, in severe patients, there exists a combination of parameters α, $k_0$, and $k_3$ that allows for unlimited viral replication, where antibody regeneration is less than the rate of antibody depletion. This leads to antibody exhaustion and a loss of germinal center activity. Elevated values of $k_0$ and reduced values of $k_3$ contribute to this phenomenon.
We can also reach this conclusion through equilibrium point analysis, which indicates that both the disease-free equilibrium state and the endemic equilibrium state are unstable. Numerical simulations conducted under the specified parameter settings demonstrate the occurrence of antibody exhaustion, as shown in Figure S_6E.
From Figure S_6E, it is evident that when $k_3$ is relatively low, and α is relatively high (0.5), infected individuals experience antibody exhaustion in response to highly replicative viruses ( $k_0$= 0.55). During this period, the viral concentration continues to rise while the antibody concentration remains at a very low level, indicating impaired antibody regeneration within the germinal centers.

| | |
|---|---|
| $V_0$ | 1 |
| $A_0$ | 0 |
| $B_0$ | 500 |
| $P_0$ | 0 |
| $C_{1_0}$ | 0 |
| $C_{2_0}$ | 0 |
| $k_0$ | 0.55 |
| $k_1$ | $10^{-7}$ |
| $k_2$ | $10^{-14}$ |
| $k_3$ | 2 |
| $k_4$ | 0.02 |
| $k_5$ | $10^{5}$ |
| $k_6$ | 0.005 |
| $k_7$ | 0.01 |

| | |
|---|---|
| $k_8$ | 0.5 |
| $\beta$ | $10^{-3}$ |

Table S11: parameter sets and initial value in antibody depletion

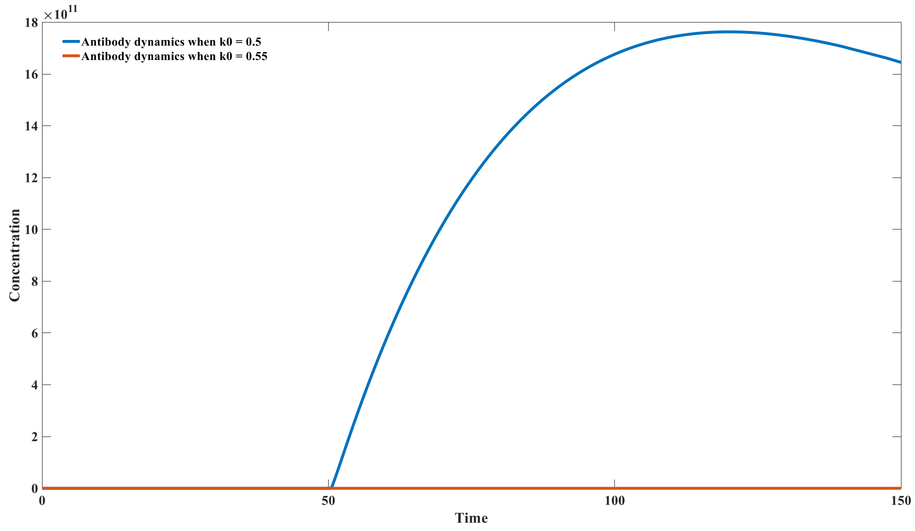


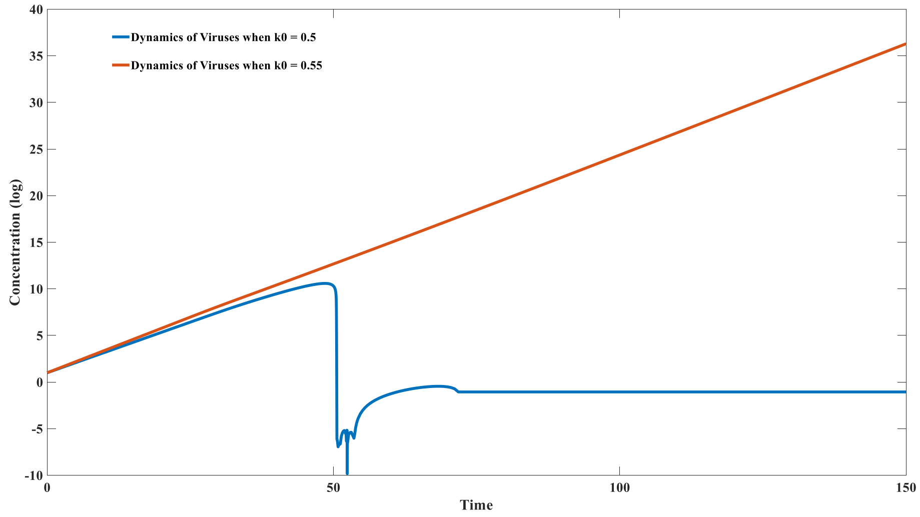


Figure S_6E: Antibody depletion can be induced in Infection caused by strong replication virus

**Parameter settings and simulation results for model 3.1.7**

Model 3.1.7 is as follows:

$$\frac{d(IgM-BCR)_i}{dt} = \pi f(IgM-BCR)_i - p_1E(IgM-BCR)_i + p_2C_{2i} + k_3C_{2i} - k_{1i}V(IgM-BCR)_i + k_{2i}C_{1i} + k_4(1-\gamma)(1-\theta)\frac{C_{1i}}{C_{1i}+(IgM-BCR)_i}C_{1i} - k_5(IgM-BCR)_i;$$

$$\frac{d(IgG-BCR)_i}{dt} = -p_1E(IgG-BCR)_i + p_2C_{4i} + k_3C_{4i} - k'_{1i}V(IgG-BCR)_i + k'_{2i}C_{3i} + k_4\gamma(1-\theta)\frac{C_{1i}}{C_{1i}+(IgM-BCR)_i}C_{1i} + k_4(1-\theta)\frac{C_{3i}}{C_{3i}+(IgG-BCR)_i}C_{3i} - k_6(IgG-BCR)_i;$$

$$\frac{d(IgM)_i}{dt} = k_7(IgM-ASC)_i - p_1E(IgM)_i + p_2C_{6i} - k_{1i}V(IgM)_i + k_{2i}C_{5i} - k_8(IgM)_i;$$

$$\frac{d(IgG)_i}{dt} = k_9(IgG-ASC)_i - p_1E(IgG)_i + p_2C_{8i} - k'_{1i}V(IgG)_i + k'_{2i}C_{7i} - k_{10}(IgG)_i;$$

$$\frac{dC_{1i}}{dt} = k_{1i}V(IgM-BCR)_i + k_{2i}C_{1i} - k_{11}C_{1i};$$

$$\frac{dC_{2i}}{dt} = p_1E(IgM-BCR)_i - p_2C_{2i} - k_{11}C_{2i};$$

$$\frac{dC_{3i}}{dt} = k'_{1i}V(IgG-BCR)_i - k'_{2i}C_{3i} - k_{11}C_{3i};$$

$$\frac{dC_{4i}}{dt} = p_1E(IgG-BCR)_i - p_2C_{4i} - k_{11}C_{4i};$$

$$\frac{dC_{5i}}{dt} = k_{1i}V(IgM)_i - k_{2i}C_{5i} - k_{11}C_{5i};$$

$$\frac{dC_{6i}}{dt} = p_1E(IgM)_i - p_2C_{6i} - k_{11}C_{6i};$$

$$\frac{dC_{7i}}{dt} = k'_{1i}V(IgG)_i - k'_{2i}C_{7i} - k_{11}C_{7i};$$

$$\frac{dC_{8i}}{dt} = p_1E(IgG)_i - p_2C_{8i} - k_{11}C_{8i};$$

$$\frac{d(IgM-ASC)_i}{dt} = k_{12}C_{2i} + k_4\epsilon(1-\gamma)\theta\frac{C_{1i}}{C_{1i}+(IgM-BCR)_i}C_{1i} - k_{13}(IgM-ASC)_i;$$

$$\frac{d(IgG-ASC)_i}{dt} = k_{12}C_{4i} + k_4\epsilon\theta\frac{C_{3i}}{C_{3i}+(IgG-BCR)_i}C_{3i} - k_{14}(IgG-ASC)_i;$$

$$\frac{dV}{dt} = k_0V - \sum_n^{i=1}k_{1i}V(IgM-BCR)_i + \sum_n^{i=1}k_{2i}C_{1i} - \sum_n^{i=1}k'_{1i}V(IgG-BCR)_i + \sum_n^{i=1}k'_{2i}C_{3i} - \sum_n^{i=1}k_{1i}V(IgM)_i + \sum_n^{i=1}k_{2i}C_{5i} - \sum_n^{i=1}k'_{1i}V(IgG)_i + \sum_n^{i=1}k'_{2i}C_{7i};$$

$$\frac{dE}{dt} = \pi_1 - \sum_n^{i=1}p_1E(IgM-BCR)_i + \sum_n^{i=1}p_2C_{2i} - \sum_n^{i=1}p_1E(IgG-BCR)_i + \sum_n^{i=1}p_2C_{4i} - \sum_n^{i=1}p_1E(IgM)_i + \sum_n^{i=1}p_2C_{6i} - \sum_n^{i=1}p_1E(IgG)_i + \sum_n^{i=1}p_2C_{8i};$$

The parameter settings that are independent of the antibody type are as follows:

| Parameter Name | Meaning | Value |
|---|---|---|
| $k_0$ | Virus replication | 1.2 |

| | | |
|---|---|---|
| | constant | |
| $k_3$ | Antibody regeneration constant of environmental antigen - antibody complex | 1 |
| $k_4$ | Antibody regeneration constant of Virus - antibody complex | 2 |
| $k_5$ | Decay constant of IgM-BCR B cell | 0.01 |
| $k_6$ | Decay constant of IgG-BCR B cell (memory B cell) | 0.005 |
| $k_7$ | IgM production rate of IgM-ASC | $4.4*10^{9}$ |
| $k_8$ | Decay constant of IgM | 0.05 |
| $k_9$ | IgG production rate of IgG-ASC | $1.2*10^{10}$ |
| $k_{10}$ | Decay constant of IgG | 0.025 |
| $k_{11}$ | Decay constant of antibody-antigen complex | 0.5 |
| $k_{12}$ | Regeneration constant of ASC from environmental antigen-antibody complex | $5*10^{-7}$ |
| $k_{13}$ | Decay constant of IgM-secreting plasma B cell | 0.1 |
| $k_{14}$ | Decay constant of IgG-secreting plasma B cell (memory plasma B cell) | 0.05 |
| $\gamma$ | Transformation ratio of IgG-BCR B cell from IgM- | 0.02 |

| | | |
|---|---|---|
| | BCR B cell | |
| θ | Maximal transformation ratio of ASC from non-plasma B cell | 0.1 |
| $\epsilon$ | ASC/BCR ratio | $10^5$ |
| $p_1$ | Forward binding constant of antibody and environmental antigen | $10^{-20}$ |
| $p_2$ | Dissociation constant of antibody-environmental antigen complex | 0.5 |
| π | Overall supplement of non-plasma IgM-BCR B cell | $0.5 * 10^{13}$ |
| $\pi_1$ | Supplement of environmental antigens | $2.2 * 10^{17} + 10^{13}$ |

Table S12: parameter sets and initial value in model 3.1.7

Due to the involvement of numerous parameters, we employed a novel approach for selecting these parameters by deriving their values based on the state of the disease-free equilibrium state. When there is no viral infection, the system transforms to:

$$\frac{d(IgM - BCR)_i}{dt} = \pi f(IgM - BCR)_i - p_1 E(IgM - BCR)_i + p_2 C_{2i} + k_3 C_{2i} - k_5 (IgM - BCR)_i;$$

$$\frac{d(IgG - BCR)_i}{dt} = -p_1 E(IgG - BCR)_i + p_2 C_{4i} + k_3 C_{4i} - k_6 (IgG - BCR)_i;$$

$$\frac{d(IgM)_i}{dt} = k_7 (IgM - ASC)_i - p_1 E(IgM)_i + p_2 C_{6i} - k_8 (IgM)_i;$$

$$\frac{d(IgG)_i}{dt} = k_9 (IgG - ASC)_i - p_1 E(IgG)_i + p_2 C_{8i} - k_{10} (IgG)_i;$$

$$\frac{dC_{2i}}{dt} = p_1 E(IgM - BCR)_i - p_2 C_{2i} - k_{11} C_{2i};$$

$$\frac{dC_{4i}}{dt} = p_1 E(IgG - BCR)_i - p_2 C_{4i} - k_{11} C_{4i};$$

$$\frac{dC_{6i}}{dt} = p_1 E(IgM)_i - p_2 C_{6i} - k_{11} C_{6i};$$

$$\frac{dC_{8i}}{dt} = p_1 E(IgG)_i - p_2 C_{8i} - k_{11} C_{8i};$$

$$\frac{d(IgM - ASC)_i}{dt} = k_{12} C_{2i} - k_{13} (IgM - ASC)_i;$$

$$\frac{d(IgG - ASC)_i}{dt} = k_{12} C_{4i} - k_{14} (IgG - ASC)_i;$$

$$\frac{dE}{dt} = \pi_1 - \sum_n^{i=1} p_1 E(IgM - BCR)_i + \sum_n^{i=1} p_2 C_{2i} - \sum_n^{i=1} p_1 E(IgG - BCR)_i + \sum_n^{i=1} p_2 C_{4i} - \sum_n^{i=1} p_1 E(IgM)_i + \sum_n^{i=1} p_2 C_{6i} - \sum_n^{i=1} p_1 E(IgG)_i + \sum_n^{i=1} p_2 C_{8i};$$

These equations consist of linear and quadratic terms, making the derivation of parameters quite straightforward. In total, there are 11 equations, corresponding to 11 equilibrium state values and 15 parameters (namely $k_3, k_5, k_6, k_7, k_8, k_9, k_{10}, k_{11}, k_{12}, k_{13}, k_{14}, p_1, p_2, \pi, \pi_1$). Based on literature reports and searches via ChatGPT, we can obtain some initial value information. We can then set certain parameters based on prior models and assumptions, while the remaining parameters will be determined through equilibrium point analysis. The initial value information and the specified parameters are as follows:

| Parameter Name | Known value |
|---|---|
| $IgM - BCR$ | $10^{15}$ |
| $IgG - BCR$ | $10^{15}$ |
| IgM | $4*10^{18}$ |
| IgG | $4*10^{19}$ |
| $C_{2i}$ | $10^{13}$ |
| $C_{4i}$ | $10^{13}$ |
| $C_{6i}$ | $4*10^{16}$ |
| $C_{8i}$ | $4*10^{17}$ |
| $IgM - ASC$ | $5*10^{7}$ |
| $IgG - ASC$ | $10^{8}$ |
| E | $10^{18}$ |
| $k_3$ | 1 |
| $k_5$ | 0.01 |
| $k_6$ | 0.005 |
| $k_8$ | 0.05 |
| $k_{10}$ | 0.025 |
| $k_{11}$ | 0.5 |
| $k_{13}$ | 0.1 |
| $k_{14}$ | 0.005 |
| $p_1$ | $10^{-20}$ |
| $p_2$ | 0.5 |

Table S13: Known parameter and initial value in model 3.1.7

The coefficients associated with antibodies are set as follows:
Here we used 100 IgMs and 100 IgGs , as well as 100 IgM -BCRs and IgG -BCRs , and 100 IgM -ASCs and IgG -ASCs . Antibody type i = 10* (m -1 ) + n (m and n are integers from 1 to 10 ) , and the *i*-th IgM-BCR has a positive affinity coefficient $k_{1i}$of 5 * 10 $^{(m - 23)}$, The negative dissociation coefficient $k_{2i}$is 10 $^{(n -1)}$ , and the distribution probability of the initial IgM -BCR is$f(IgM - BCR)_i$ = P(m-1 ⩽ X ⩽ m)* P( n -1⩽X⩽n ) , the log values of the positive affinity coefficient and negative dissociation coefficient follow a normal distribution ( $N(5,0.8^2)$ ) which we derived based on experimental reports and previous studies. The i- th IgG - BCR antibody has a positive affinity coefficient $k_{1i}'$of 10 $^{(m -23)}$, The negative dissociation coefficient $k_{2i}'$is 10 $^{(n -1)}$, and the distribution probability of the initial IgG - BCR is $f(IgG - BCR)_i$ = P(m-1 ⩽ X ⩽ m)* P( n -1 ⩽ X ⩽ n ) ; the i - th IgM has a positive affinity coefficient $k_{1i}$of 5 * 10 $^{(m -23)}$, The negative dissociation coefficient $k_{2i}$is 10 $^{(n -1)}$, and the initial IgM distribution probability is $f(IgM)_i$ = P(m-1 ⩽ X ⩽ m)* P( n -1 ⩽ X ⩽ n ) ; the *i*-th IgG antibody has a positive affinity coefficient $k_{1i}'$of 1 0 $^{(m -23)}$, The negative dissociation coefficient $k_{2i}'$is 10 $^{(n -1)}$, and the initial IgG distribution probability is $f(IgG)_i$ = P(m-1 ⩽ X ⩽ m)* P( n -1 ⩽ X ⩽ n ) . A point of clarification is that because IgM -BCRs are often present on the B cell surface as multimers, the binding energy of an IgM- BCR multimer to a homopolymeric viral antigen is n times greater than that of a monomer (n is the polymerization number of the IgM -BCR multimer). Therefore, we consider the positive binding coefficient of IgM -BCR to be five times greater than that of IgG -BCR . Similarly, in body fluids, IgM exists as a pentamer, so we also consider the positive binding coefficient of IgM to be five times greater than that of IgG. This allows us to calculate the initial values of the various compartments for each antibody type based on the total initial number of IgM , IgG , IgM - BCR , IgG -BCR , IgM - ASC , and IgG -ASC .

The coefficients related to antibodies are set as follows:
We utilize 100 types of IgM and 100 types of IgG, along with 100 types of IgM-BCR and IgG-BCR, as well as 100 types of IgM-ASC and IgG-ASC. The antibody type *i* is defined as ( 10* (*m*-1) +*n*), where *m* and *n* are integers ranging from 1 to 10. The forward affinity coefficient $k_{1i}$ for the $i^{th}$ IgM-BCR is 5*10 $^{(m-23)}$ , and the negative dissociation coefficient $k_{2i}$ is 10 $^{(n-1)}$ . The initial distribution probability of IgM-BCR is given by ($f(IgM - BCR)_i$ = P(*m*-1 ⩽ X ⩽ *m*)* P(*n*-1 ⩽ X ⩽ *n*)) . The logarithmic values of the forward affinity coefficients and negative dissociation coefficients follow a normal distribution ($N(5, 0.8^2)$), which is derived based on experimental reports and prior studies.
For the $i^{th}$ IgG-BCR antibody, the forward affinity coefficient $k_{1i}'$ is 10 $^{(m-23)}$ , and the negative dissociation coefficient $k_{2i}'$ is 10 $^{(n-1)}$ . The initial distribution probability of IgG-BCR is given by ( $f(IgG - BCR)_i$ = P(*m*-1 ⩽ X ⩽ *m*)* P(*n*-1 ⩽ X ⩽ *n*)) .
The $i^{th}$ IgM possesses a forward affinity coefficient $k_{1i}$ = 5*10$^{(m-23)}$ and a negative dissociation coefficient $k_{2i}$ = 10 $^{(n-1)}$ . The initial distribution probability of IgM is ( $f(IgM)_i$ = P(*m*-1 ⩽ X ⩽ *m*)* P(*n*-1 ⩽ X ⩽ *n*)).
For the $i^{th}$ IgG antibody, the forward affinity coefficient $k_{1i}'$ = 10 $^{(m-23)}$ and the negative dissociation coefficient $k_{2i}'$ = 10 $^{(n-1)}$ . The initial distribution probability of IgG is ( $f(IgG)_i$ = P(*m*-1 ⩽ X ⩽ *m*)* P(*n*-1 ⩽ X ⩽ *n*) ).
It is important to note that because IgM-BCR predominantly exists on the surface of B cells

in a multimeric form, the binding energy between IgM-BCR multimers and homologous polymeric viral macromolecular antigens is *N* times greater than that of the monomer (where *N* is the degree of polymerization of the IgM-BCR multimer). We assume that the forward binding coefficient of IgM-BCR is five times that of IgG-BCR. Similarly, in bodily fluids, IgM also exists in the form of pentamers; therefore, we also assume that the forward binding coefficient of IgM to the virus is five times that of IgG.
Thus, we can calculate the initial values for various compartments of each antibody type based on the total initial quantities of IgM, IgG, IgM-BCR, IgG-BCR, IgM-ASC, and IgG-ASC.

The simulation results are shown in Figure S_7A:

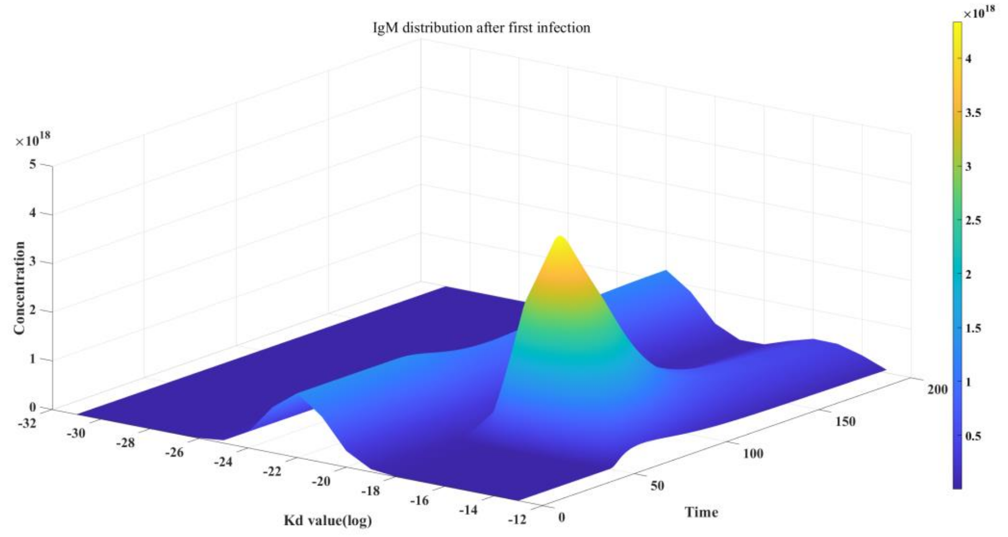


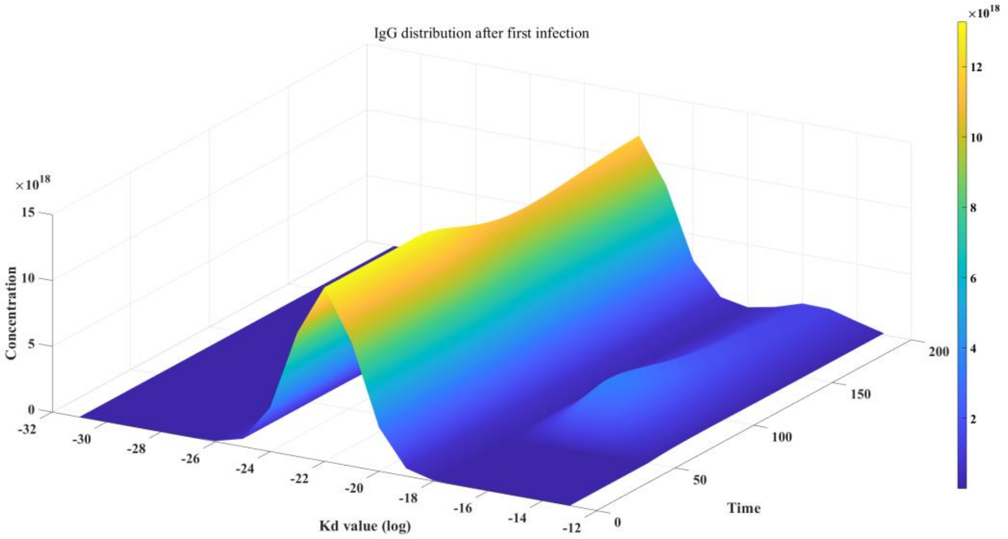


Figure S_7A: Antibody distribution after virus infection

From the figure, it is evident that the proliferation rate of IgM significantly outpaces that of IgG following the initial infection; however, IgM subsequently declines rapidly and cannot maintain a stable peak in the high-affinity region over an extended period. In contrast, while the proliferation rate of IgG is relatively slower, it can sustain a stable peak in the high-affinity region for a prolonged duration. Stability analysis of this new disease-free equilibrium state reveals that all eigenvalues are negative, indicating that it is a stable state, which forms the dynamic basis for immune memory. Given that IgG ultimately derives from the conversion of IgM, the initial distributions of both IgM and IgG, as well as those of IgM-BCR and IgG-BCR, are similar. This similarity contributes to the rapid response of IgM and IgM-BCR to viral infections due to their multimeric structure, which results in kinetics that are significantly faster than those of IgG and IgG-BCR. This rapid response facilitates the host's ability to quickly proliferate specific antibodies.

Moreover, IgG predominantly exists in monomeric form, allowing the host to maintain immune memory with a lower synthetic cost. The same mass of IgG clearly represents a greater number of molecules compared to IgM, leading to enhanced viral neutralization activity. Therefore, the formation of IgM multimers and IgG monomers in humoral immunity is not a coincidence but rather a result of long-term evolution. This differentiation aids in the swift activation of the immune system while also efficiently sustaining immune memory.

We further simulated the determination of IgM and IgG levels using the ELISA method. The principle of ELISA involves adding antigen components to measure the quantity of antibodies that bind to them, making the dosage of the antigen critical. If the dosage is too low, even with very low titers of specific antibodies, most antigens can still bind to antibodies due to the presence of a large pool of moderately binding antibodies in the humoral fluid. Conversely, if the antigen dosage is excessively high, most antigens will remain unneutralized, affecting the outcome of the assay. The antigen-antibody reaction in ELISA closely resembles our model 3.1.7, with the sole difference being that this model only considers IgM and IgG since cells have been filtered out, resulting in the absence of IgM-BCR, IgG-BCR, IgM-ASC, and IgG-ASC, and the antigen used is not a live virus, thus ( $k_0$ = 0). Using a dosage of $10^{16}$, we obtained the results shown in Figure S7_B. As illustrated in Figure S7_B, there is a notable increase in IgM during the primary infection, while the increase in IgG during secondary infection is more pronounced, reflecting the kinetic characteristics of humoral immunity.

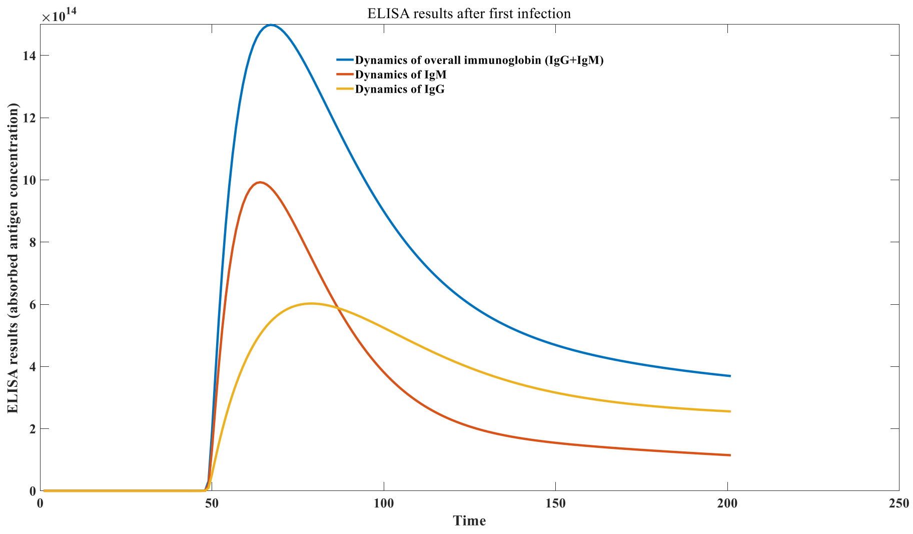

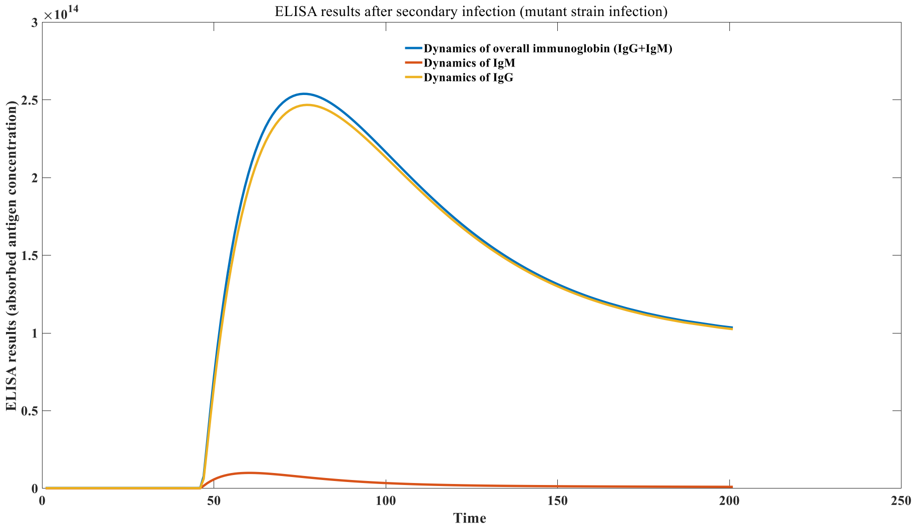


Figure S7_B: Different dynamics of IgM and IgG in primary infection and secondary infection

To further investigate the principles of immune memory and the theory of discontinuous antibody decay, we increased the complexity of the model. We classified the newly formed IgG-BCR (memory B cells) into two types: one type is derived from the proliferation of IgG-BCR, while the other type arises from the conversion of IgM-BCR. Consequently, our model has evolved to:

$$\frac{d(\mathrm{IgM}-\mathrm{BCR})_i}{dt} = \pi f(\mathrm{IgM}-\mathrm{BCR})_i - p_1 E(\mathrm{IgM}-\mathrm{BCR})_i + p_2 C_{2i} + k_3 C_{2i} - k_{1i} V(\mathrm{IgM}-\mathrm{BCR})_i + k_{2i} C_{1i} + k_4(1-\gamma)(1-\theta)\frac{C_{1i}}{C_{1i}+(\mathrm{IgM}-\mathrm{BCR})_i} C_{1i} - k_5(\mathrm{IgM}-\mathrm{BCR})_i;$$

$$\frac{d(\mathrm{IgG}-\mathrm{BCR})_{1i}}{dt} = -p_1 E(\mathrm{IgG}-\mathrm{BCR})_{1i} + p_2 C_{4i} + k_3 C_{4i} - k'_{1i} V(\mathrm{IgG}-\mathrm{BCR})_{1i} + k'_{2i} C_{3i} + k_4(1-\theta)\frac{C_{3i}}{C_{3i}+(\mathrm{IgG}-\mathrm{BCR})_{1i}+(\mathrm{IgG}-\mathrm{BCR})_{2i}} C_{3i} - k_6(\mathrm{IgG}-\mathrm{BCR})_{1i};$$

$$\frac{d(\mathrm{IgG}-\mathrm{BCR})_{2i}}{dt} = -p_1 E(\mathrm{IgG}-\mathrm{BCR})_{2i} + p_2 C_{4i}' + k_3 C_{4i}' - k'_{1i} V(\mathrm{IgG}-\mathrm{BCR})_{2i} + k'_{2i} C_{3i}' + k_4 \gamma(1-\theta)\frac{C_{1i}}{C_{1i}+(\mathrm{IgM}-\mathrm{BCR})_{1i}} C_{1i} + k_4(1-\theta)\frac{C_{3i}'}{C_{3i}++(\mathrm{IgG}-\mathrm{BCR})_{1i}+(\mathrm{IgG}-\mathrm{BCR})_{2i}} C_{3i}' - k_6(\mathrm{IgG}-\mathrm{BCR})_{2i};$$

$$\frac{d(\mathrm{IgM})_i}{dt} = k_7(\mathrm{IgM}-\mathrm{ASC})_i - p_1 E(\mathrm{IgM})_i + p_2 C_{6i} - k_{1i} V(\mathrm{IgM})_i + k_{2i} C_{5i} - k_8(\mathrm{IgM})_i;$$

$$\frac{d(\mathrm{IgG})_i}{dt} = k_9(\mathrm{IgG}-\mathrm{ASC})_i - p_1 E(\mathrm{IgG})_i + p_2 C_{8i} - k'_{1i} V(\mathrm{IgG})_i + k'_{2i} C_{7i} - k_{10}(\mathrm{IgG})_i;$$

$$\frac{dC_{1i}}{dt} = k_{1i}V(IgM - BCR)_i + k_{2i}C_{1i} - k_{11}C_{1i};$$

$$\frac{dC_{2i}}{dt} = p_1E(IgM - BCR)_i - p_2C_{2i} - k_{11}C_{2i};$$

$$\frac{dC_{3i}}{dt} = k'_{1i}V(IgG - BCR)_i - k'_{2i}C_{3i} - k_{11}C_{3i};$$

$$\frac{dC_{4i}}{dt} = p_1E(IgG - BCR)_i - p_2C_{4i} - k_{11}C_{4i};$$

$$\frac{dC_{3i}'}{dt} = k'_{1i}V(IgG - BCR)_i - k'_{2i}C_{3i} - k_{11}C_{3i};$$

$$\frac{dC_{4i}'}{dt} = p_1E(IgG - BCR)_i - p_2C_{4i} - k_{11}C_{4i};$$

$$\frac{dC_{5i}}{dt} = k_{1i}V(IgM)_i - k_{2i}C_{5i} - k_{11}C_{5i};$$

$$\frac{dC_{6i}}{dt} = p_1E(IgM)_i - p_2C_{6i} - k_{11}C_{6i};$$

$$\frac{dC_{7i}}{dt} = k'_{1i}V(IgG)_i - k'_{2i}C_{7i} - k_{11}C_{7i};$$

$$\frac{dC_{8i}}{dt} = p_1E(IgG)_i - p_2C_{8i} - k_{11}C_{8i};$$

$$\frac{d(IgM - ASC)_i}{dt} = k_{12}C_{2i} + k_4\epsilon(1-\gamma)\theta\frac{C_{1i}}{C_{1i} + (IgM - BCR)_i}C_{1i} - k_{13}(IgM - ASC)_i;$$

$$\frac{d(IgG - ASC)_i}{dt} = k_{12}C_{4i} + k_4\epsilon\theta\frac{C_{3i}}{C_{3i} + (IgG - BCR)_i}C_{3i} - k_{14}(IgG - ASC)_i;$$

$$\frac{dV}{dt} = k_0V - \sum_n^{i=1} k_{1i}V(IgM - BCR)_{1i} + \sum_n^{i=1} k_{2i}C_{1i} - \sum_n^{i=1} k'_{1i}V(IgG - BCR)_{1i} + \sum_n^{i=1} k'_{2i}C_{3i} - \sum_n^{i=1} k'_{1i}V(IgG - BCR)_{2i} + \sum_n^{i=1} k'_{2i}C_{3i}{}' - \sum_n^{i=1} k_{1i}V(IgM)_i + \sum_n^{i=1} k_{2i}C_{5i} - \sum_n^{i=1} k'_{1i}V(IgG)_i + \sum_n^{i=1} k'_{2i}C_{7i};$$

$$\frac{dE}{dt} = \pi_1 - \sum_n^{i=1} p_1E(IgM - BCR)_i + \sum_n^{i=1} p_2C_{2i} - \sum_n^{i=1} p_1E(IgG - BCR)_{1i} + \sum_n^{i=1} p_2C_{4i} - \sum_n^{i=1} p_1E(IgG - BCR)_{2i} + \sum_n^{i=1} p_2C_{4i}{}' - \sum_n^{i=1} p_1E(IgM)_i + \sum_n^{i=1} p_2C_{6i} - \sum_n^{i=1} p_1E(IgG)_i + \sum_n^{i=1} p_2C_{8i};$$

In this model, we introduced several compartments: $(IgG - BCR)_{1i}$ represents IgG-BCR derived from IgG-BCR proliferation, while $(IgG - BCR)_{2i}$ denotes IgG-BCR converted from IgM-BCR. $C_{3i}'$ signifies the complexes formed between $(IgG - BCR)_{2i}$ and the virus, and $C_{4i}'$ represents the complexes formed between $(IgG - BCR)_{2i}$ and environmental antigens E. Based on this model, we can quantitatively simulate the changes in $(IgG - BCR)_{1i}$ and $(IgG - BCR)_{2i}$ during viral infection, resulting in Figure S7_C.
From Figure S7_C, it is evident that a significant proportion of the newly formed IgG-BCR originates from the conversion of IgM-BCR. These converted IgG-BCRs are high-affinity BCRs,

which form the foundation for the theory of discontinuous antibody decay. Specifically, when an infection by a particular virus concludes, IgM will gradually disappear over time, while IgG stabilizes at a constant level. However, this scenario assumes that no further viral infections occur, which is clearly unrealistic. When another new virus infects the host, if all newly formed IgG-BCRs arise solely from the original IgG-BCR, the concentration of specific IgG against the original antibodies would remain unchanged, as environmental antigens consistently maintain a stable overall antibody level.

The crux of the issue lies in the fact that a portion of IgG-BCR always originates from the conversion of IgM-BCR, and the proportion of high-affinity IgG-BCR targeting the original virus among those derived from IgM-BCR conversion is extremely low. This is because IgM-BCR does not have immune memory; it quickly reverts to its original normal distribution after the infection ends and does not maintain a peak in the high-affinity region. Consequently, the final population of IgG-BCR will exhibit a decreased titer of specific neutralizing antibodies against the original virus. The more frequently the host is infected with heterologous viruses subsequently, the greater the decline in specific IgG against the original virus will be. This leads to the phenomenon where the titers of specific neutralizing antibodies display a trend of discontinuous stepwise decline. This declining trend may be rejuvenated following secondary infections caused by variants of the original strain.

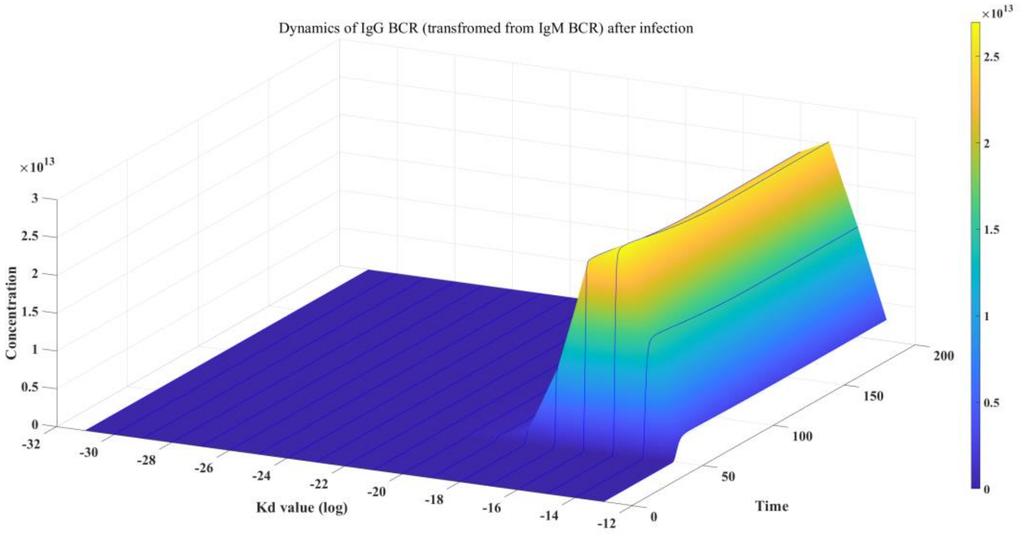

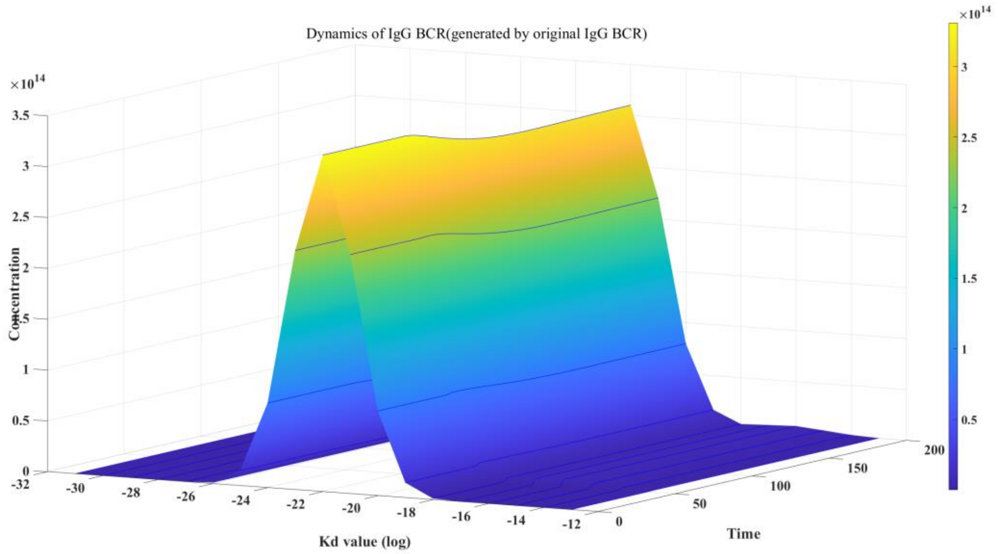


Figure S7_C: Dynamics of IgG with different originality

#### The principle of immunoblotting and its numerical simulation

To study immune imprinting, it is necessary to first simulate the mechanisms of secondary infection. Generally, for DNA viruses that do not mutate easily, although antibodies exhibit a trend of stepwise decline over time and with subsequent heterologous infections, this discontinuous decrease is relatively slow, allowing antibodies to provide protection for an extended period, with some even offering lifelong immunity. However, the situation is different for rapidly mutating RNA viruses. While the antibody levels against the original strain may maintain a peak within the high-affinity region, the altered properties of the virus mean that the antibody distribution categorized by binding strength to the original strain will no longer apply to the mutant strains. The antibody distribution targeting the mutant strains will instead become an intermediate distribution, lying between the initial normal distribution and the bimodal distribution against the original virus.

Therefore, we introduce a parameter α to define the mutation coefficient of the virus. When the mutation coefficient is α, the antibody distribution against the mutant strain can be expressed as:

[ $10^{-\alpha}$*(antibody distribution against the original strain) + ($1 - 10^{-\alpha}$) *(initial normal distribution).]

Of course, this linear approximation has its drawbacks, but it also has the advantage of being computationally convenient. Thus, when the mutation coefficient is 0, this new antibody distribution is equivalent to the original bimodal distribution, while as α approaches infinity, it approximates the initial normal distribution.

We denote $f$ as the reference normal distribution, $F$ as the distribution of antibody binding parameters for the wild-type antigen, $N$ as the total number of antibodies, and $F_{New}$ as the new distribution of antibody binding parameters for the mutated antigen.

$$f(x; \mu, \ \ \sigma) = \frac{1}{\sigma\sqrt{2\Pi}} \exp\left(\frac{-(x-\mu)^2}{2\sigma^2}\right)$$

$$F_{New} = (1 - 10^{-\alpha})fN + 10^{-\alpha}FN$$

We studied the relationship between viral proliferation and the mutation coefficient during secondary infection, after the initial infection ends and antibodies reach a steady state, as shown in Figure S_8A . As shown in Figure S_8A , when the mutation coefficient is low (α <= 3 ), viral proliferation is prevented, meaning that secondary infection symptoms do not occur. However, as the mutation coefficient increases, the peak viral concentration gradually increases. Therefore, the primary driving force behind secondary infection with variant strains is viral mutation, not a decrease in specific antibody levels against the original strain.

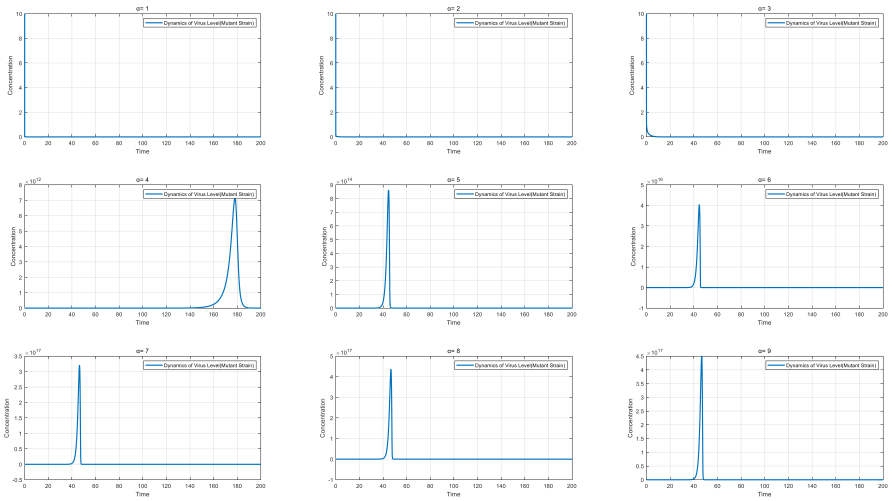


Figure S8_A: Dynamics of mutant viruses in secondary Infection with different α value

We further investigated the principles of immune imprinting and defined two numerical values for cross-imprinting. Here, $Cross\ reaction_1$ represents the commonly used percentage of cross-reaction. The formula for calculating the proportion of cross-reaction is expressed in equations (22-25), where $Cross\ reaction_1$ indicates the proportion of cross-reaction with the wild-type antigen, and $Cross\ reaction_2$ denotes the proportion of cross-reaction with the mutant strain antigen. $antibody(i)$ refers to the concentration of antibodies with strong affinity to the mutant strain after the second infection, specifically indicating antibodies 51, 61-62, 71-73, 81-84, and 91-95.

The term "strong affinity" here is defined based on the Kd value of the antibodies; we can also use stricter Kd values to calculate the immune imprinting levels of those super high-affinity antibodies. $aantibody(i)_{initial}$ represents the initial concentration of antibodies with strong affinity to the mutant strain at the time of the second infection, while $baseline$ denotes the reference concentration. Additionally, $antibody_{\_first\ infection}(i)$ indicates the concentration of antibodies with strong affinity to the wild type after the first infection.

The relationship between different mutation coefficients α and immune imprinting during natural infections is shown in Figure S_8B, while the relationship between the mutation coefficient and immune imprinting during the re-vaccination with mutant strains after natural infection is illustrated in Figure S_8C.

$$Cross\ reaction_1 = \sum_{i=1}^{n} antibody(i) * \frac{antibody(i)_{initial} - baseline}{antibody(i)_{initial}} / \sum_{i=1}^{n} antibody(i)$$

$$Cross\ reaction_2 = \sum_{i=1}^{n} antibody(i) * \frac{antibody(i)_{initial} - baseline}{antibody(i)_{initial}} / \sum_{i=1}^{n} antibody_{\_first\ infection}(i)$$

$$antibody(i)_{initial} = antibody_{\_first\ infection}(i) * (10^{-\alpha})$$

$$baseline = (1 - 10^{-\alpha})fN$$

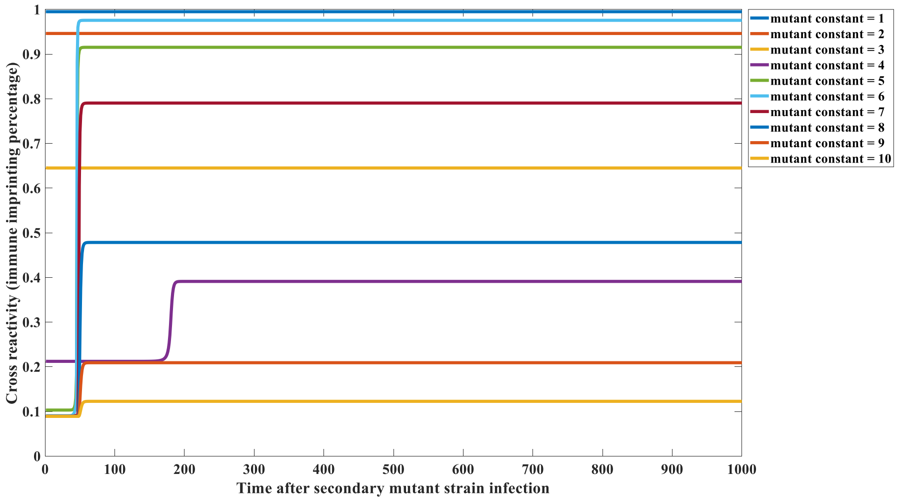

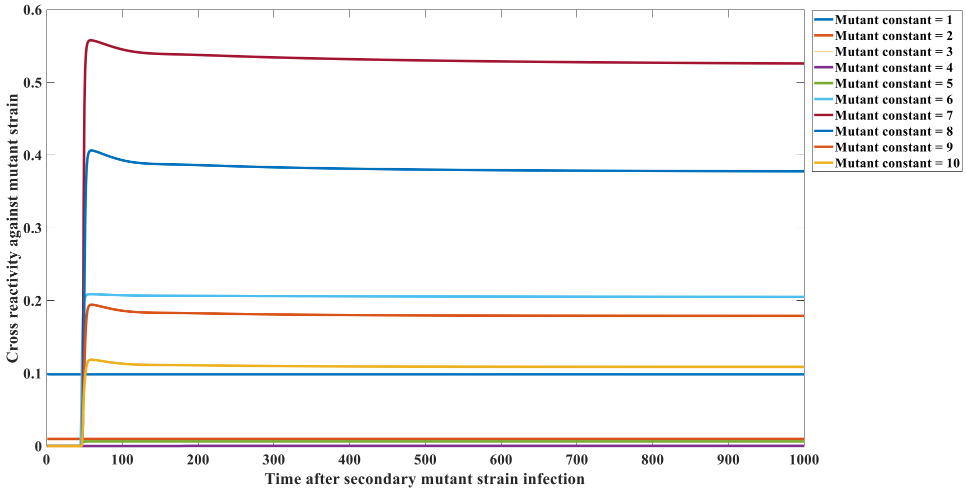


Figure S8_B: Relationship between α value and immune imprinting cross reactivity after natural Infection

From Figure S8_B, an interesting phenomenon is observed: the level of immune imprinting first decreases, then increases, and finally decreases again as the viral mutation coefficient α increases. This occurs because when the α value is less than or equal to 3, it does not lead to significant infections, and therefore, the antibody repertoire is not reshaped. As a result, immune imprinting appears as a horizontal line, although this level will decline with increasing α, since the immune imprinting level of specific antibodies approximates to $\frac{10^{-\alpha}FN}{(1-10^{-\alpha})fN+10^{-\alpha}FN}$. However, when α is greater than or equal to 4, secondary infections occur, resulting in the reshaping of antibody levels. In this case, an increase in the α value leads to a strong proliferation of specific antibodies, thus increasing the level of immune imprinting. As the mutation coefficient α continues to rise, the main source of specific antibodies targeting the mutant strains gradually shifts toward those that do not cross-react with the original strain. Consequently, the immune imprinting level begins to show a declining trend once again. This phenomenon also occurs in the affinity cross-reaction ($Cross\ reaction_2$) between specific antibodies against the original strain and the mutant strain.

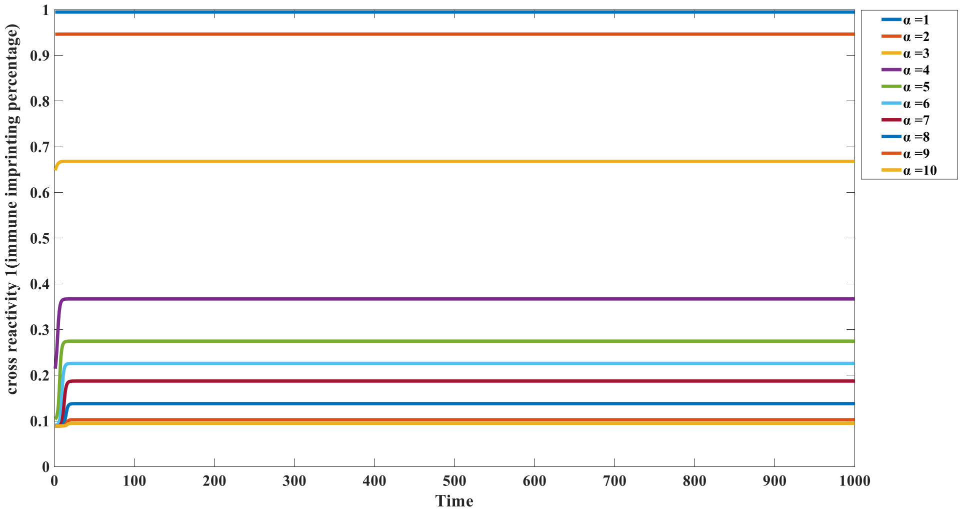


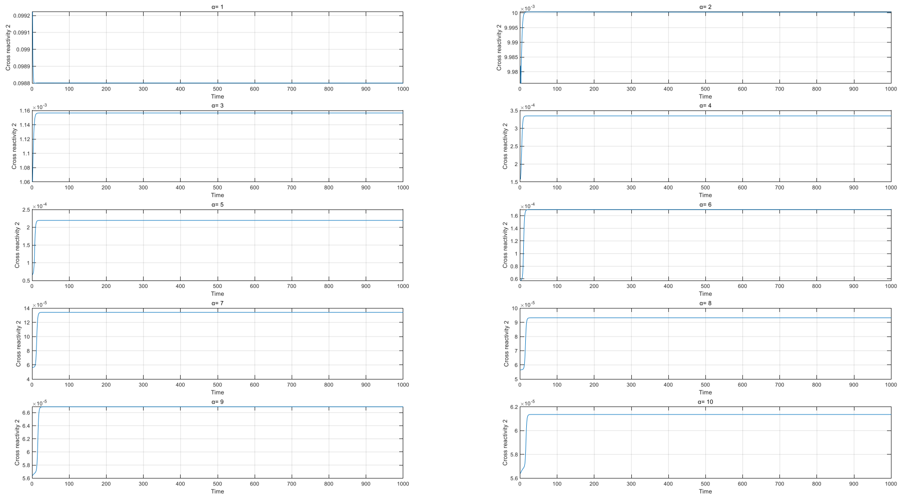

Figure S8_C: Relationship between α value and immune imprinting cross reactivity after mutant vaccination

However, vaccination can lead to different outcomes. As shown in Figure S8_C, when the vaccination dose is $10^{14}$, both $Cross\,reaction_1$ and $Cross\,reaction_2$ exhibit a downward trend as the viral mutation coefficient α increases, particularly the value of $Cross\,reaction_1$, which represents the level of immune imprinting. This indicates that the greater the mutation coefficient of the mutant strain, the lower the proportion of immune imprinting after vaccination.

Another interesting phenomenon is that even when the mutation coefficient approaches infinity (meaning the two viruses are completely from different origins), immune imprinting still exists at a certain level. For example, in this case, the minimum level is approximately around 10%.

**Parameter setting of model 3.2.1**

Model 3.2.1 is as follows:

Type I antigens:

$$\frac{dV}{dt} = k_0V - k_1VA + k_2C;$$

$$\frac{dA}{dt} = -k_1VA + k_2C + k_3C + \pi - k_4A;$$

$$\frac{dC}{dt} = k_1VA - k_2C - k_5C;$$

$$\frac{dTc}{dt} = k_6(C + V) + \pi_1 - k_7Tc;$$

Second type of antigens:

$$\frac{dV}{dt} = k_0V - k_1VA + k_2C;$$

$$\frac{dA}{dt} = -k_1VA + k_2C + k_3C + \pi - k_4A;$$

$$\frac{dC}{dt} = k_1VA - k_2C - k_5C;$$

$$\frac{dTc}{dt} = k_6C + \pi_1 - k_7Tc;$$

The first type of antigen refers to antigens that can be recognized by Pattern Recognition Receptors (such as TLR and CLR) on the surface of DC cells, such as the new coronavirus; the second type of antigen refers to antigens that cannot be recognized by the pattern recognition receptors on the surface of DC cells. recognition Antigens recognized by receptors need to first bind to antibodies to form antigen-antibody complexes, and then be detected by Fc Receptor recognition includes self-antigens and LMCV viruses. Therefore, the cellular immunity effect stimulated by these two types of viruses, that is, the proliferation capacity of Tc cells, has a different relationship with humoral immunity. The parameter settings are as follows:

| V ariable Name | Initial Value |
|---|---|
| $V_0$ | $10^{10}$ |
| $A_0$ | $10^{5}$ |
| $C_0$ | 0 |
| $Tc_0$ | $10^{3}$ |
| Parameter Name | Value |
| $k_0$ | - 0.05 |
| $k_1$ | $10^{-7}$ |
| $k_2$ | $10^{-14}$ |
| $k_3$ | 5 in normal group, 3 in rituximab treatment group |

| | |
|---|---|
| $k_4$ | 0.01 |
| $k_5$ | 2 |
| $k_6$ | $10^{-5}$ |
| $k_7$ | 0.01 |
| $\pi$ | $10^{3}$ |
| $\pi_1$ | 10 |

Table S14: parameter and initial value in model 3.2.1

The simulation results of vaccination with the first type of antigen are shown in Figure S_9A, while the results for the second type of antigen are presented in Figure S_9B. From Figure S_9A, it can be observed that for the first type of antigen, patients with suppressed humoral immunity during vaccination show a significantly smaller proliferation of specific antibodies compared to the control group. However, their CD8+ T cell proliferation is significantly stronger than that of the control group. This is because the decay rate of the antigen is notably lower than the rate at which the antigen-antibody complexes disappear. Due to the presence of a large number of Fc receptors in the body, such as those on NK cells, the degradation rate of antigen-antibody complexes is relatively rapid. The production of a large number of antibodies leads to the formation of antigen-antibody complexes with the antigens in the vaccine. Although these complexes can stimulate dendritic cell (DC) proliferation and CD8+ T cell activation, their short lifespan results in fewer CD8+ T cells being produced compared to populations with restricted humoral immunity.
In contrast, for the second type of antigen, the situation is entirely different. Since DCs can only recognize antigen-antibody complexes, the activation level of cellular immunity shows a positive correlation with the magnitude of antibody production in humoral immunity. Therefore, in vaccinated individuals with restricted humoral immunity, the activation of their cellular immune function is also suppressed.

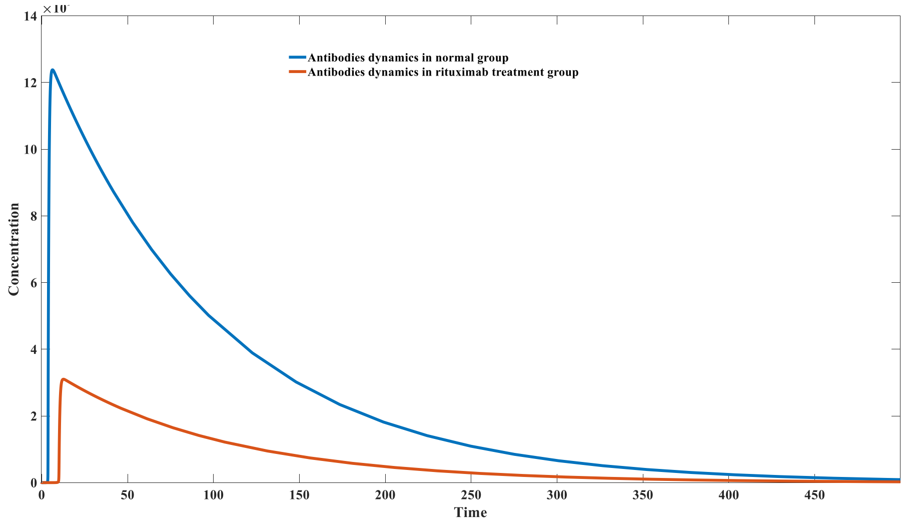

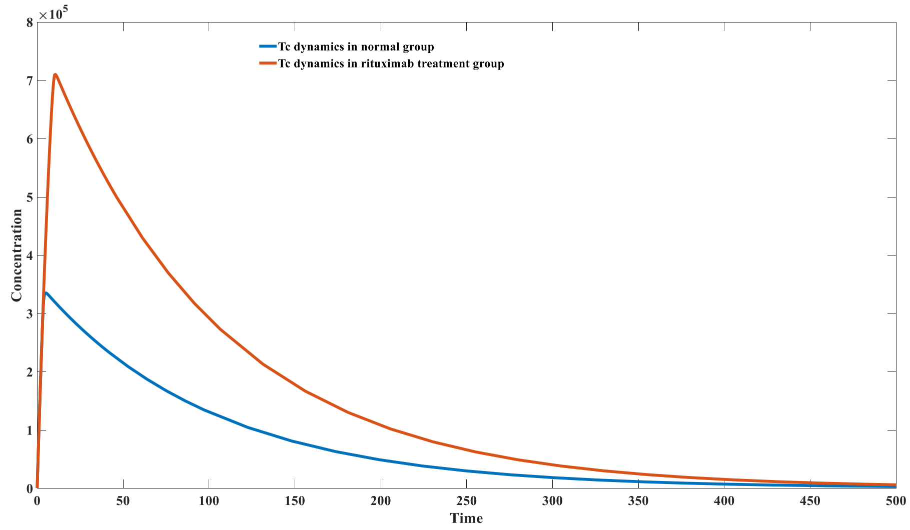


Figure S9_A: Opposite trends of humoral and cellular immune response after rituximab treatment after first type antigen vaccination.

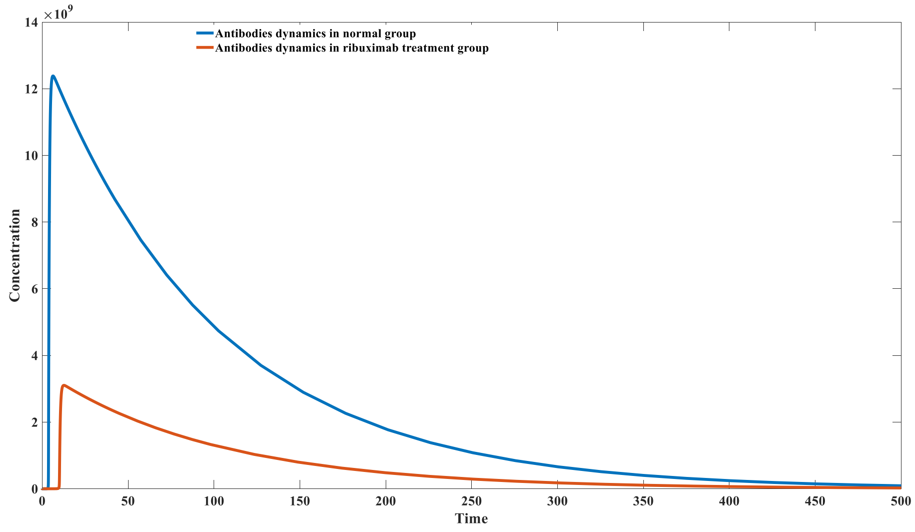

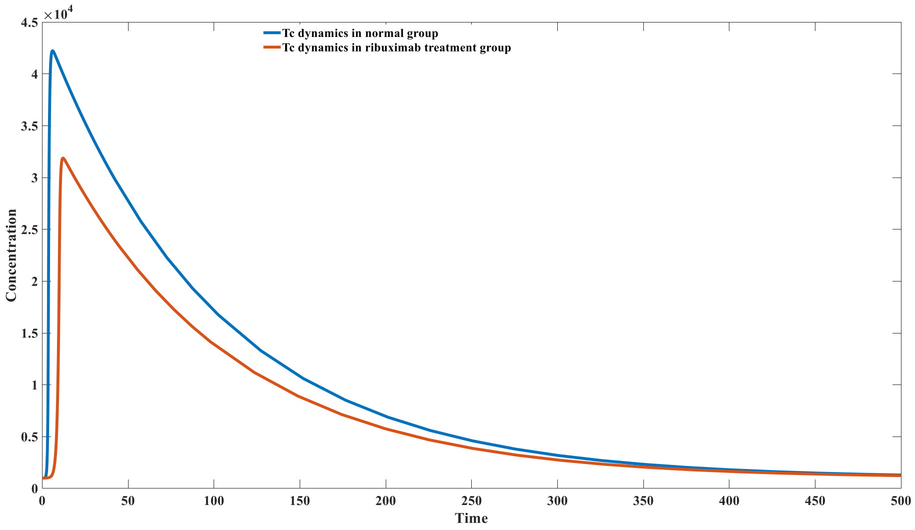


F igure S9_B: Decreased humoral and cellular immune response after rituximab treatment after second type antigen vaccination.

### Model 3.2.2 Parameter setting

Model 3.2.2 is as follows:

$$\frac{dTc}{dt} = \sum_{n}^{i=1} -k_1 N_i Tc + k_{2i} C_i \, ;$$

$$\frac{dN_i}{dt} = -k_1 N_i Tc + k_{2i} C_i;$$

$$\frac{dC_i}{dt} = k_1 N_i Tc - k_{2i} C_i;$$

$$\log(k_{2i}) = \mu - \tau \frac{V_i/N_i}{\frac{V_i}{N_i} + Km};$$

Model 3.2.2 is primarily used to study the relationship between the cytotoxicity of CD8+ T cells and the concentration of antigens within infected cells. It actually forms part of the cellular immunity model in Model 3.4.1 . Its parameters are set as follows:

| Parameter Name | Value |
|---|---|
| $k_1$ | 0.01 |
| μ | 12 |
| τ | 16 |
| Km | $10^3$ |

Table S15: parameter and initial value in model 3.2.2

### Mathematical analysis and parameter setting of model 3.3.1

Model 3.3.1 is as follows:

$$\frac{d(IgM)_i}{dt} = \pi f(IgM)_i - p_1E(IgM)_i/\beta + p_2(C_2)_i - k_{1i}V(IgM)_i/\beta + k_{2i}(C\_1)_i + \alpha\beta\gamma k_3\sum_{j=1}^{m}(F_1)_{ij}$$
$$+ \alpha\beta\gamma k_3\sum_{j=1}^{m}(F_2)_{ij} + k_4\beta\gamma\sum_{j=1}^{m}(F_3)_{ij} + k_4\beta\gamma\sum_{j=1}^{m}(F_4)_{ij} - d_1(IgM)_i;$$

$$\frac{d(IgG)_i}{dt} = -p_1E(IgG)_i/\beta + p_2(C_4)_i - k'_{1i}V(IgG)_i/\beta + k'_{2i}(C_3)_i + (1-\alpha)\beta\gamma k_3\sum_{j=1}^{m}(F_1)_{ij}$$
$$+ (1-\alpha)\beta\gamma k_3\sum_{j=1}^{m}(F_2)_{ij} + k_3\beta\gamma\sum_{j=1}^{m}(F_5)_{ij} + k_3\beta\gamma\sum_{j=1}^{m}(F_6)_{ij} + k_4\beta\gamma\sum_{j=1}^{m}(F_7)_{ij}$$
$$+ k_4\beta\gamma\sum_{j=1}^{m}(F_8)_{ij} - d_2(IgG)_i;$$

$$\frac{d(T_1)_j}{dt} = \pi_1 f(T_1)_j - \sum_{i=1}^{n}((D_2)_i(T_1)_j\,\omega_1 - \omega_2(F_3)_{ij}) - \sum_{i=1}^{n}((D_1)_i(T_1)_j\,\sigma_{1i} - \sigma_{2i}(F_1)_{ij})$$
$$- \sum_{i=1}^{n}((D_4)_i(T_1)_j\,\omega_1 - \omega_2(F_7)_{ij}) - \sum_{i=1}^{n}((D_3)_i(T_1)_j\,\sigma_{1i} - \sigma_{2i}(F_5)_{ij})$$
$$+ \alpha k_3\sum_{i=1}^{n}(F_1)_{ij} + \alpha k_3\sum_{i=1}^{n}(F_5)_{ij} + k_4\sum_{i=1}^{n}(F_3)_{ij} + k_4\sum_{i=1}^{n}(F_7)_{ij} - d_3(T_1)_j;$$

$$\frac{d(T_2)_j}{dt} = -\sum_{i=1}^{n}((D_2)_i(T_2)_j\,\omega_1 - \omega_2(F_4)_{ij}) - \sum_{i=1}^{n}((D_1)_i(T_2)_j\,\sigma_{1i} - \sigma_{2i}(F_2)_{ij})$$
$$- \sum_{i=1}^{n}((D_4)_i(T_2)_j\,\omega_1 - \omega_2(F_8)_{ij}) - \sum_{i=1}^{n}((D_3)_i(T_2)_j\sigma_{1i} - \sigma_{2i}(F_6)_{ij})$$
$$+ (1-\alpha)k_3\sum_{i=1}^{n}(F_1)_{ij} + (1-\alpha)k_3\sum_{i=1}^{n}(F_5)_{ij} + k_3\sum_{i=1}^{n}(F_2)_{ij} + k_3\sum_{i=1}^{n}(F_6)_{ij}$$
$$+ k_4\sum_{i=1}^{n}(F_4)_{ij} + k_4\sum_{i=1}^{n}(F_8)_{ij} - d_4(T_2)_j;$$

$$\frac{d(C_1)_i}{dt} = k_{1i}V(IgM)_i/\beta - k_{2i}(C_1)_i - k_5(C_1)_i - d_5(C_1)_i;$$
$$\frac{d(C_2)_i}{dt} = p_1E(IgM)_i/\beta - p_2(C_2)_i - k_5(C_2)_i - d_5(C_2)_i;$$

$$\frac{d(C_3)_i}{dt} = k'_{1i}V(IgG)_i/\beta - k'_{2i}(C_3)_i - k_5(C_3)_i - d_5(C_3)_i;$$

$$\frac{d(C_4)_i}{dt} = p_1E(IgG)_i/\beta - p_2(C_4)_i - k_5(C_4)_i - d_5(C_4)_i;$$

$$\frac{d(D_1)_i}{dt} = k_5(C_1)_i - \sum_{j=1}^{m}(D_1)_i(T_1)_j\,\sigma_{1i} - \sigma_{2i}(F_1)_{ij}) - \sum_{j=1}^{m}((D_1)_i(T_2)_j\,\sigma_{1i} - \sigma_{2i}(F_2)_{ij}) - d_6(D_1)_i\,;$$

$$\frac{d(D_2)_i}{dt} = k_5(C_2)_i - \sum_{j=1}^{m}(D_2)_i(T_1)_j\,\sigma_{1i} - \sigma_{2i}(F_3)_{ij}) - \sum_{j=1}^{m}((D_2)_i(T_2)_j\,\sigma_{1i} - \sigma_{2i}(F_4)_{ij}) - d_6(D_2)_i\,;$$

$$\frac{d(D_3)_i}{dt} = k_5(C_3)_i - \sum_{j=1}^{m}(D_3)_i(T_1)_j\,\sigma_{1i} - \sigma_{2i}(F_5)_{ij}) - \sum_{j=1}^{m}((D_3)_i(T_2)_j\,\sigma_{1i} - \sigma_{2i}(F_6)_{ij}) - d_6(D_3)_i\,;$$

$$\frac{d(D_4)_i}{dt} = k_5(C_4)_i - \sum_{j=1}^{m}(D_4)_i(T_1)_j\,\sigma_{1i} - \sigma_{2i}(F_7)_{ij}) - \sum_{j=1}^{m}((D_4)_i(T_2)_j\,\sigma_{1i} - \sigma_{2i}(F_8)_{ij}) - d_6(D_4)_i\,;$$

$$\frac{d(F_1)_{ij}}{dt} = \sigma_{1i}(D_1)_i(T_1)_j - \sigma_{2i}(F_1)_{ij} - d_7(F_1)_{ij};$$

$$\frac{d(F_2)_i}{dt} = \sigma_{1i}(D_1)_i(T_2)_j - \sigma_{2i}(F_2)_{ij} - d_7(F_2)_{ij};$$

$$\frac{d(F_3)_{ij}}{dt} = \omega_1(D_2)_i(T_1)_j - \omega_2(F_3)_{ij} - d_7(F_3)_i;$$

$$\frac{d(F_4)_{ij}}{dt} = \omega_1(D_2)_i(T_2)_j - \omega_2(F_4)_{ij} - d_7(F_4)_{ij};$$

$$\frac{d(F_5)_{ij}}{dt} = \sigma_{1i}(D_3)_i(T_1)_j - \sigma_{2i}(F_5)_{ij} - d_7(F_5)_{ij};$$

$$\frac{d(F_6)_{ij}}{dt} = \sigma_{1i}(D_3)_i(T_2)_j - \sigma_{2i}(F_6)_{ij} - d_7(F_6)_{ij};$$

$$\frac{d(F_7)_{ij}}{dt} = \omega_1(D_4)_i(T_1)_j - \omega_2(F_7)_{ij} - d_7(F_7)_{ij};$$

$$\frac{d(F_8)_{ij}}{dt} = \omega_1(D_4)_i(T_2)_j - \omega_2(F_8)_{ij} - d_7(F_8)_{ij};$$

$$\frac{dV}{dt} = k_0V - \sum_{i=1}^{n}(\frac{k_{1i}V(IgM)_i}{\beta} - k_{2i}(C_1)_i) - \sum_{i=1}^{n}(\frac{k'_{1i}V(IgG)_i}{\beta} - k'_{2i}(C_3)_i)\,;$$

$$\frac{dE}{dt} = \pi_2 - \sum_{i=1}^{n}(\frac{p_1E(IgM)_i}{\beta} - p_2(C_2)_i) - \sum_{i=1}^{n}(\frac{p_1E(IgG)_i}{\beta} - p_2(C_4)_i)\,;$$

The initial values and parameters settings unrelated to BCR and TCR typing are as follows:

| Variable Name | Initial Value |
| --- | --- |

| | |
|---|---|
| $V_0$ | 10 |
| $E_0$ | $10^{15}$ |
| IgM | $10^{15}$ |
| IgG | $10^{15}$ |
| $T_1$ | $10^{10}$ |
| $T_2$ | $10^{10}$ |
| $C_1$ | 0 |
| $C_2$ | $3.6 * 10^{13}$ |
| $C_3$ | 0 |
| $C_4$ | $3.6 * 10^{13}$ |
| $D_1$ | 0 |
| $D_2$ | $10^{13}$ |
| $D_3$ | 0 |
| $D_4$ | $10^{13}$ |
| $F_1$ | 0 |
| $F_2$ | 0 |
| $F_3$ | $10^{8}$ |
| $F_4$ | $10^{8}$ |
| $F_5$ | 0 |
| $F_6$ | 0 |
| $F_7$ | $10^{8}$ |
| $F_8$ | $10^{8}$ |
| Parameter Name | Value |
| $k_0$ | 0.8 |
| $k_3$ | 2 |
| $k_4$ | 1 |
| $k_5$ | 0.5 |
| $p_1$ | $10^{-16}$ |
| $p_2$ | 10/3.6-1 |
| $d_1$ | 0.008 |
| $d_2$ | 0.004 |
| $d_3$ | 0.016 |
| $d_4$ | 0.008 |
| $d_5$ | 0.5 |
| $d_6$ | 0.6 |
| $d_7$ | 0.6 |
| $\omega_1$ | $10^{-4}$ |
| $\omega_2$ | $10^{6}$-0.6 |
| $\alpha$ | 0.95 |
| $\beta$ | $10^{5}$ |
| $\gamma$ | 2 |
| $\pi$ | $0.1 * 10^{13}$ |
| $\pi_1$ | $0.8 * 10^{8}$ |

| | |
|---|---|
| $\pi_2$ | $7.2 * 10^{13}$ |

Table S16: parameter and initial value in model 3.3.1

The selection principles for the parameters are the same as those described in model 3.1.7, ensuring that the system can achieve a balanced state when disease-free. The estimates of substance concentrations at equilibrium were obtained through literature searches and ChatGPT queries. We generally consider that the ratio of T cells to B cells at equilibrium is 1:1, and similarly, the ratio of memory cells to effector cells is also 1:1. Since there are typically about $10^5$ BCRs on a single B cell, the number of BCRs is $10^5$ times that of T cells. The parameters set in the table are highlighted in blue, while those estimated based on the equilibrium state are highlighted in red. Both mathematical analysis and numerical simulations indicate that when the initial virus concentration is zero, the system can maintain a stable disease-free state without fluctuations in the values of various compartments.

The coefficients related to BCR and TCR are set as follows: Due to computational speed constraints, we did not use combinations of 100 different BCRs and 100 different TCRs. Instead, we adopted two schemes: one with a small diversity of BCRs (2 types of BCRs) combined with a large diversity of TCRs (100 types of TCRs), and another with a small diversity of TCRs (2 types of TCRs) combined with a large diversity of BCRs (100 types of BCRs).

For the first case, the affinity coefficients of the two IgM-BCRs are( $k_{11} = 5 * 10^{-12}; k_{21} = 10^{-3}; k_{12} = 5 * 10^{-16}; k_{22} = 10;$) and the affinity coefficients of the two IgG-BCRs are( $k_{11}' = 10^{-12}; k_{21}' = 10^{-3}; k_{12}' = 10^{-16}; k_{22}' = 10;$). The proportion of strong binding BCRs is $10^{-5}$, and the proportion of weak binding BCRs is ($1-10^{-5}$). 100 TCRs (including T CRs of memory CD4+ T cells and TCRs of non-memory CD4 + T cells). TCR type $i = 10*(m-1) + n$ ($m$ and $n$ are integers from 1 to 10 ) , the positive affinity coefficient of the $i^{th}$ TCR to the MHC- II complex $\sigma_{1i}$ is $10^{(m-13)}$, The negative dissociation coefficient $\sigma_{2i}$ is $10^{(n-4)}$, and the distribution probability of initial effector CD4 + T cells and memory CD4 + T cells is $f(T)_i = P(m-1 \leqslant X \leqslant m) * P(n-1 \leqslant X \leqslant n)$, with the log values of the positive affinity coefficient and negative dissociation coefficient following a normal distribution ($N(5,0.8^2)$) . The equilibrium simulation results for this case are shown in Figure S10_A , and the virus infection simulation results are shown in Figure S10_B .

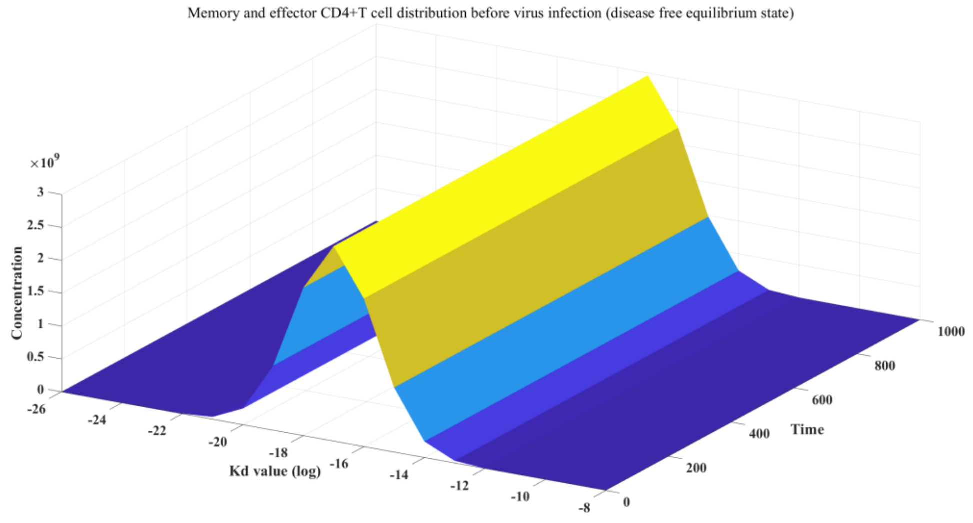

Figure S10_A: Disease free equilibrium state in B-CD4+T interaction model before infection (BCR diversity = 2; TCR diversity = 100).

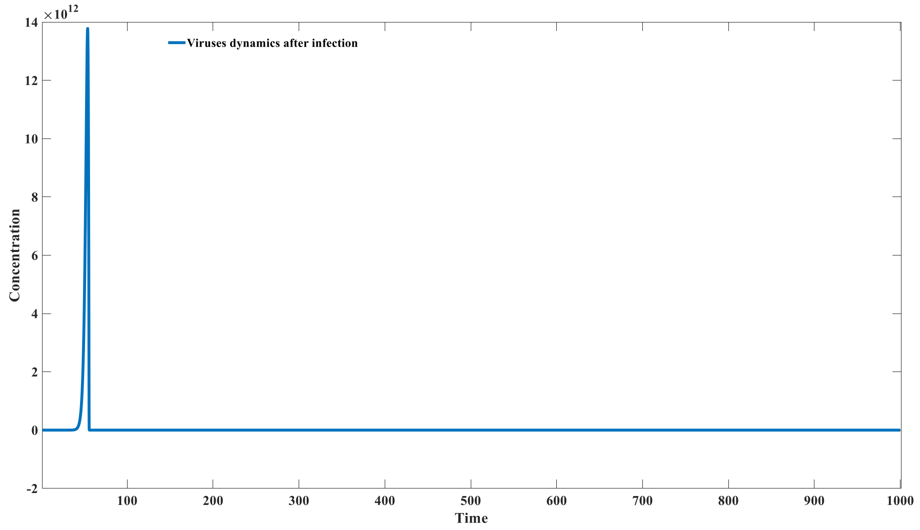


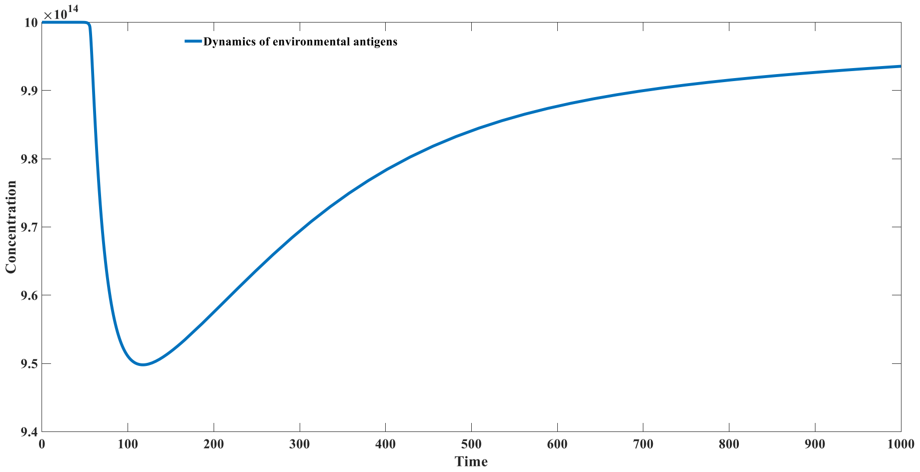

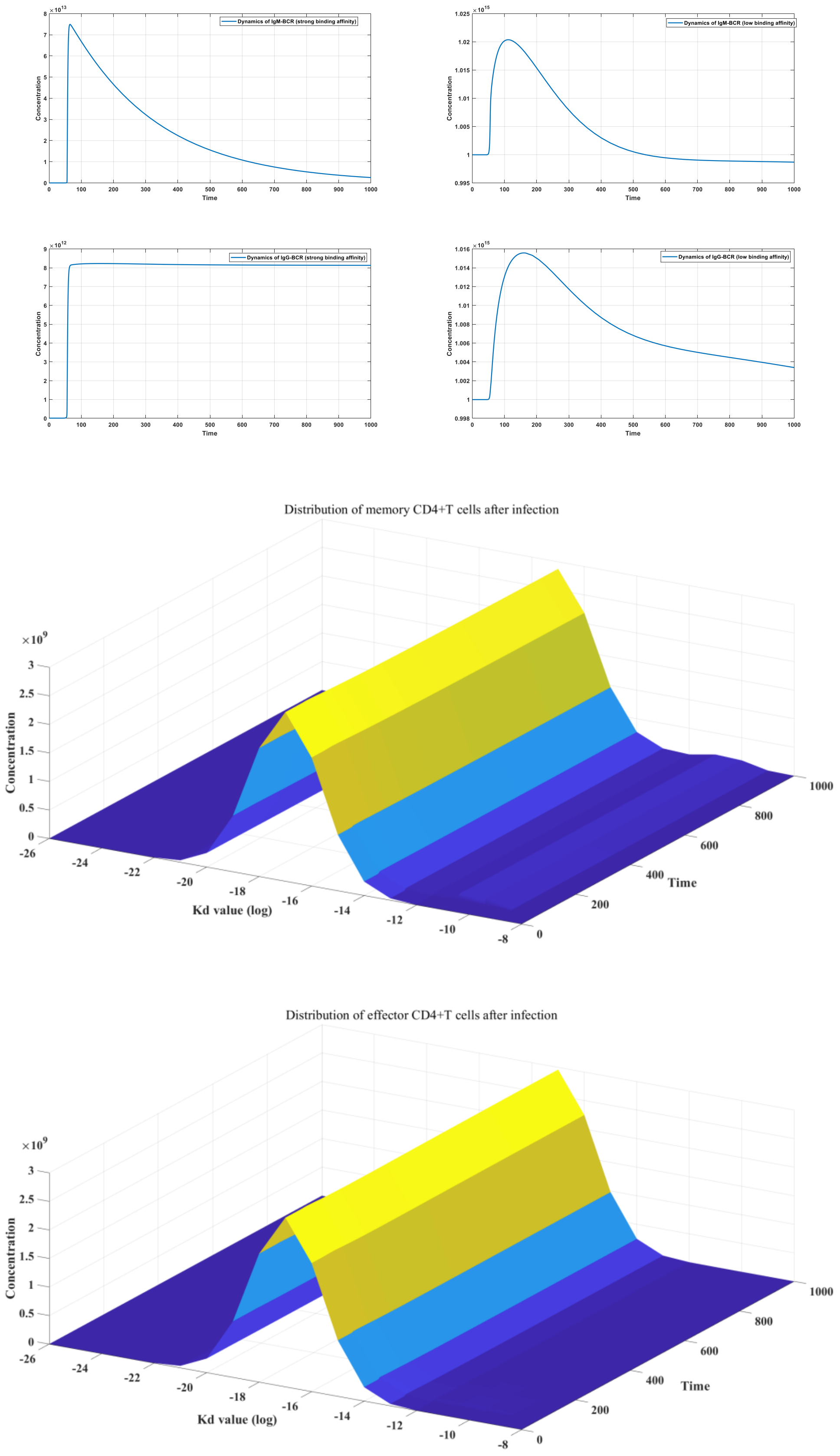


Figure S10_B: Dynamics of host-virus interaction in B-CD4+T interaction model after infection (BCR diversity = 2; TCR diversity = 100).

For the second case, the affinity coefficients of the two TCRs are ( $\sigma_{11} = 5 * 10^{-12}; \sigma_{21} = 10^{-3}; \sigma_{12} = 5 * 10^{-16}; \sigma_{22} = 10;$), where the proportion of strong binding TCR is $10^{-5}$ and the proportion of weak binding TCR is ($1 - 10^{-5}$).

There are 100 BCRs ( including IgG-BCRs of memory B cells and IgM-BCRs of non-memory B cells ). BCR type antibody type $i = 10*(m-1) + n$ ($m$ and $n$ are integers from 1 to 10 ) , the $i^{th}$ IgM- BCR has a positive affinity coefficient $k_{1i}$ of $5 * 10^{(m-22)}$, The negative dissociation coefficient $k_{2i}$ is $10^{(n-4)}$, and the initial IgM - BCR distribution probability is $f(IgM)_i = P(m-1 \leqslant X \leqslant m) * P(n-1 \leqslant X \leqslant n)$ , the log values of its positive affinity coefficient and negative dissociation coefficient follow a normal distribution ( $N(5,0.8^2)$ ) . The $i^{th}$ IgG -BCR antibody has a positive affinity coefficient $k_{1i}'$ of $10^{(m-22)}$, The negative dissociation coefficient $k_{2i}'$ is $10^{(n-4)}$, and the distribution probability of the initial IgG - BCR is $f(IgG)_i = P(m-1 \leqslant X \leqslant m) * P(n-1 \leqslant X \leqslant n)$. The simulation results of viral infection in this case are shown in Figure S10_C .

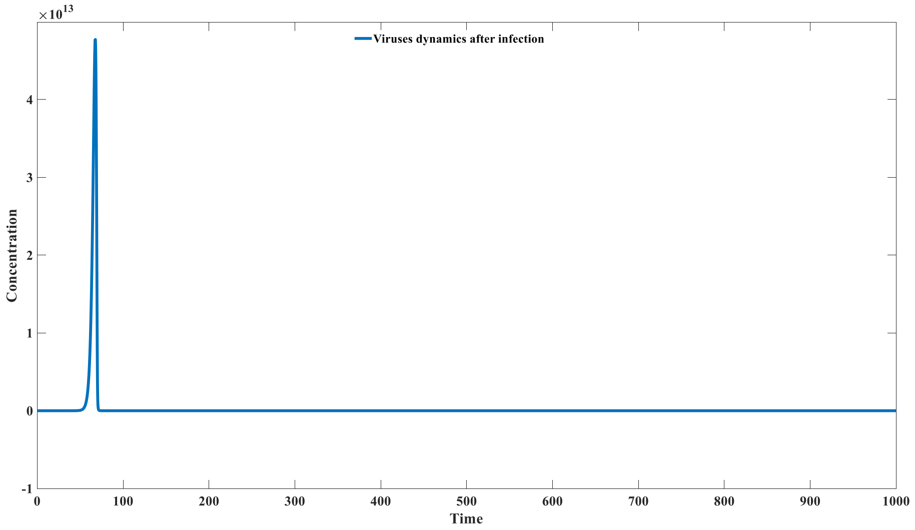


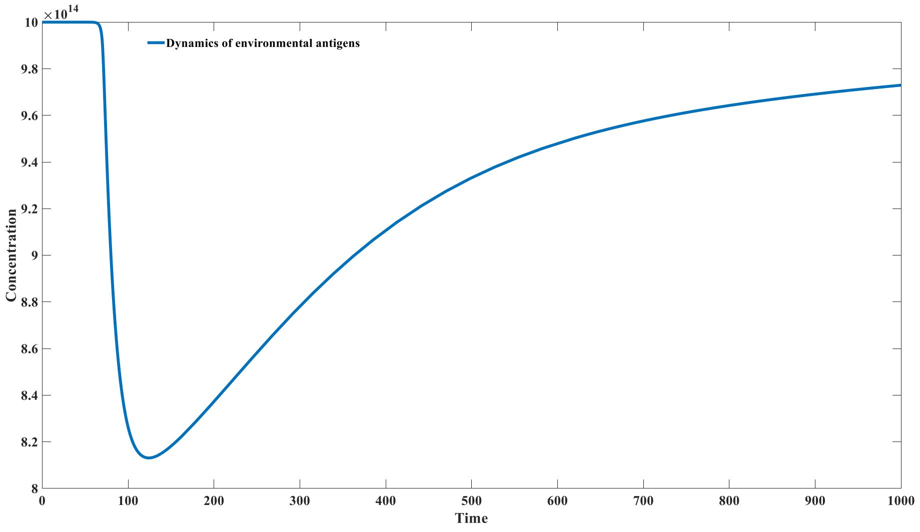

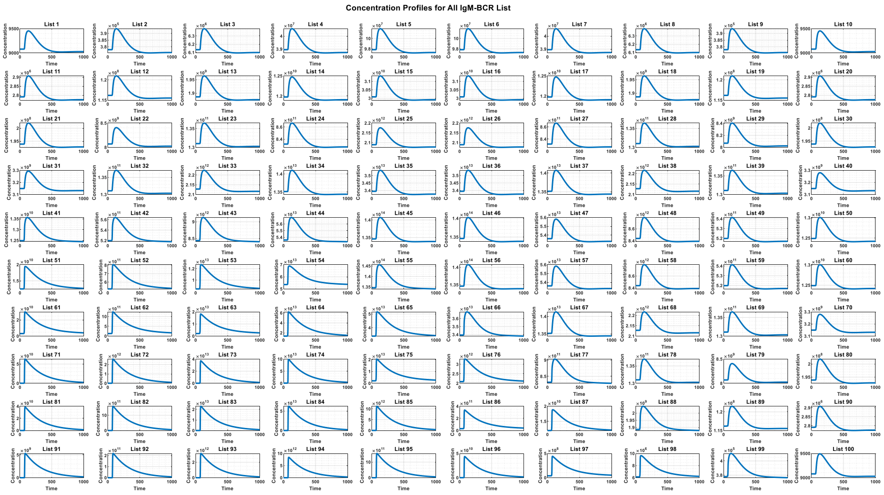
Concentration Profiles for All IgM-BCR List

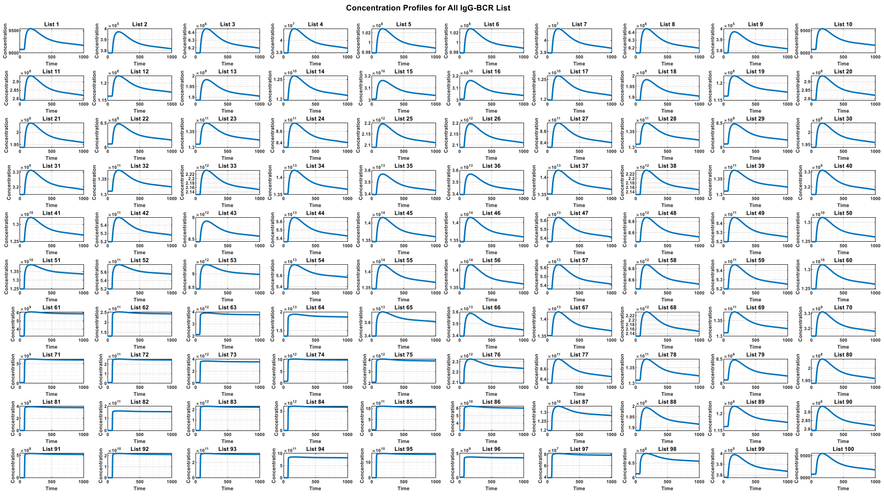
Concentration Profiles for All IgG-BCR List

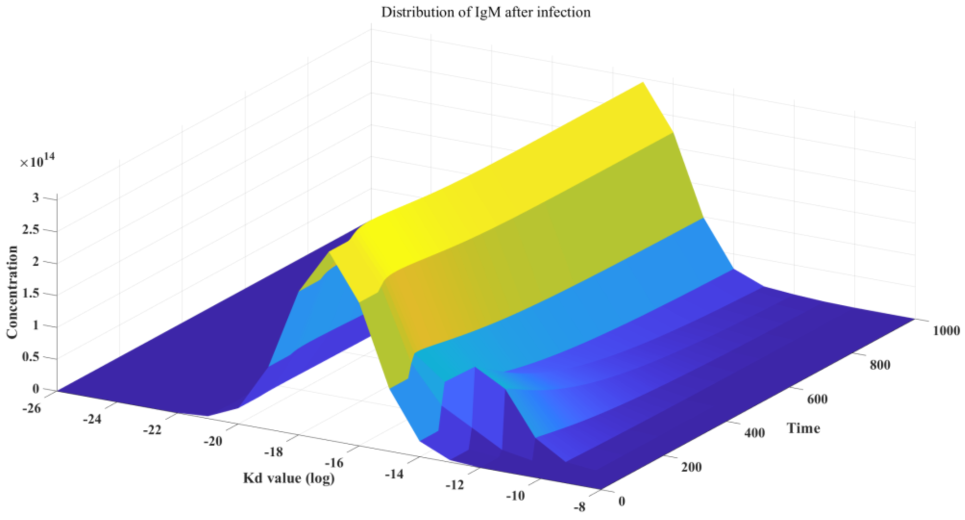
Distribution of IgM after infection
×10^14
3
2.5
2
1.5
1
0.5
0
Concentration
-26
-24
-22
-20
-18
-16
-14
-12
-10
-8
Kd value (log)
0
200
400
600
800
1000
Time

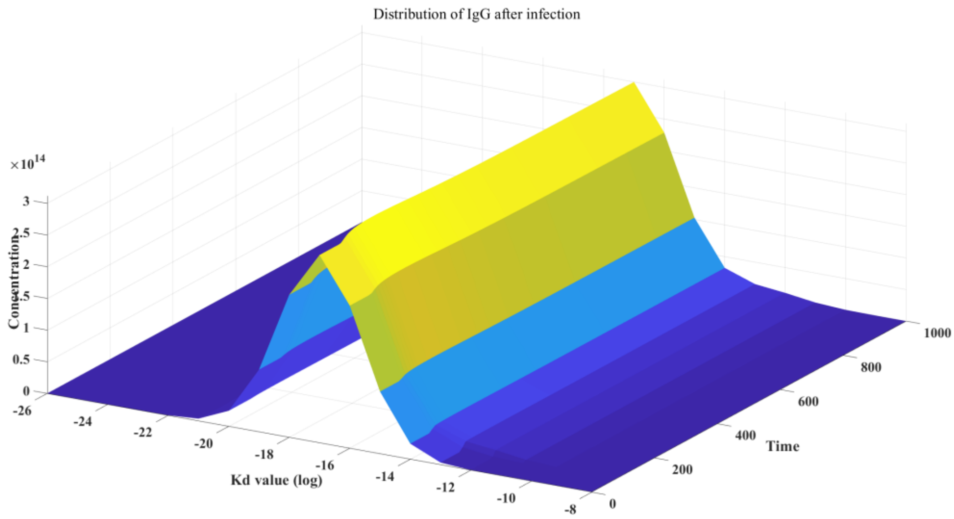


Figure S10_C: Dynamics of host-virus interaction in B-CD4+ T interaction model after Infection (BCR diversity = 100; TCR diversity = 2).

We further investigated the preventive capability of CD4+ T cell memory and B cell memory against secondary infections. For the first scenario, we studied the preventive capacity of CD4+ T cell memory against secondary infections. We first simulated the primary infection over a time span from 0 to 1000 time points. Then, we modeled the situation with CD4+ T cell memory by setting the concentrations of all CD4+ T cell components (100 memory CD4+ T cells and 100 effector CD4+ T cells) to be equal to the concentrations at the 1000th time point while initializing the B cell components. Subsequently, we conducted simulations for secondary infections, and the results are shown in S10_D.

From the figure, it can be seen that although CD4+ T cell memory can reduce the peak viral concentration during secondary infections, it cannot completely prevent the occurrence of secondary infections.

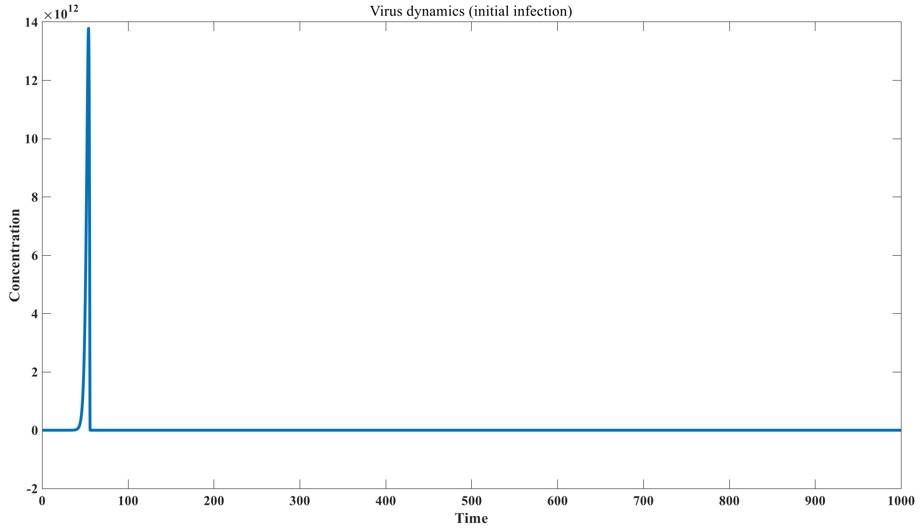

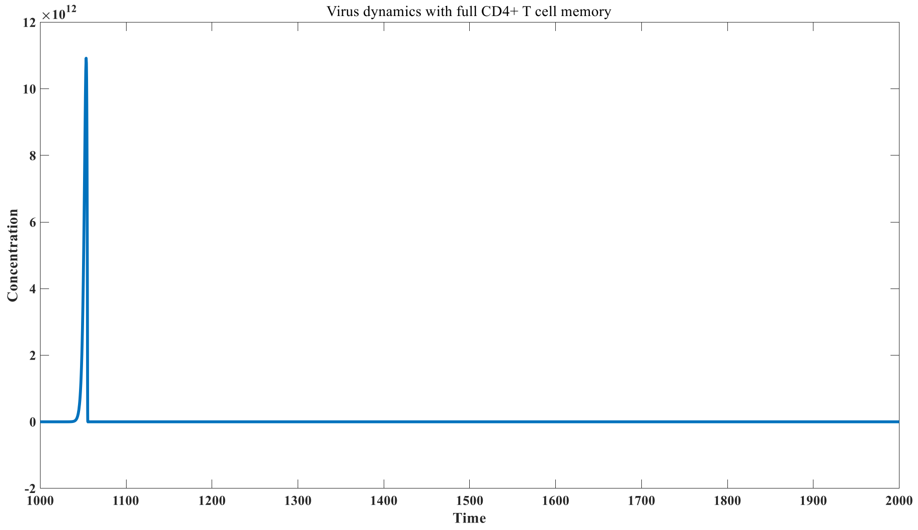


Figure S10_D: effect of CD4+ T cell memory in preventing secondary infection

We used the second scenario to study the preventive capacity of B cell memory against secondary infections. We first simulated the primary infection over a time span from 0 to 1000 time points. Then, we modeled the situation with B cell memory by setting the concentrations of all B cell components (100 IgM B cells and 100 IgG B cells) to be equal to the concentrations at the 1000th time point while initializing the T cell concentrations. Subsequently, we conducted simulations for secondary infections, and the results are shown in S10_E.

From the figure, it is evident that complete B cell memory can fully prevent the occurrence of secondary infections. However, as described in the main text, B cell memory relies heavily on the tertiary structure of antigenic epitopes for B cells, making it vulnerable to point mutations that can significantly weaken its original memory function. The mutational effects caused by point mutations can lead to a rapid decline in B cell memory. Therefore, while B cell memory is powerful, it is also very transient and fragile, quickly becoming ineffective when faced with rapidly mutating RNA viruses.

Nonetheless, as discussed in our immunological imprinting section, the retained B cell immune memory imprints can still rapidly initiate humoral immunity, effectively reducing the severity of secondary infections. T cell memory includes two types: CD4+ T cell memory and CD8+ T cell memory. Here, we only discuss CD4+ T cell memory; however, both types of T cell memory target the primary sequences of antigens, making them less sensitive to point mutations. This contributes to the long-lasting nature of T cell memory, which can reduce the severity of secondary infections, although it cannot completely prevent their occurrence.

When discussing the protective efficacy of vaccines, especially regarding those targeting rapidly mutating RNA viruses, we cannot rely solely on traditional parameters such as the duration of protection for evaluation.

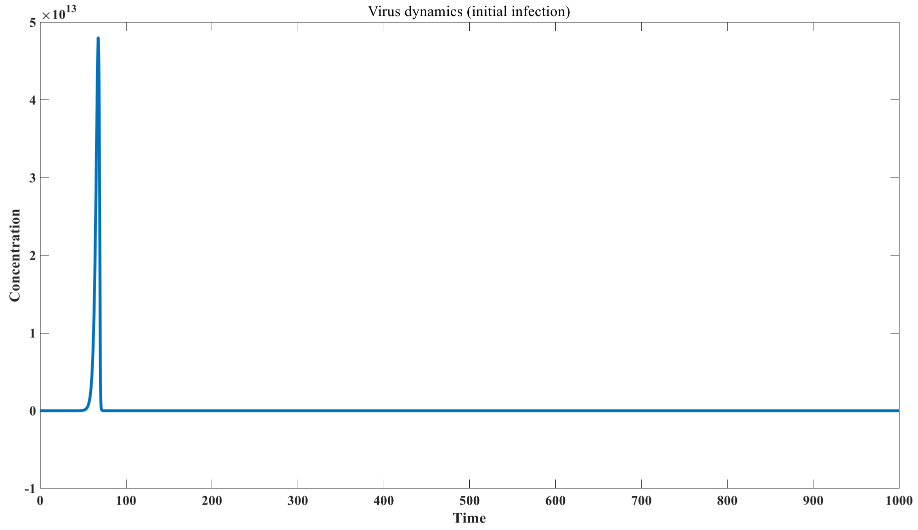


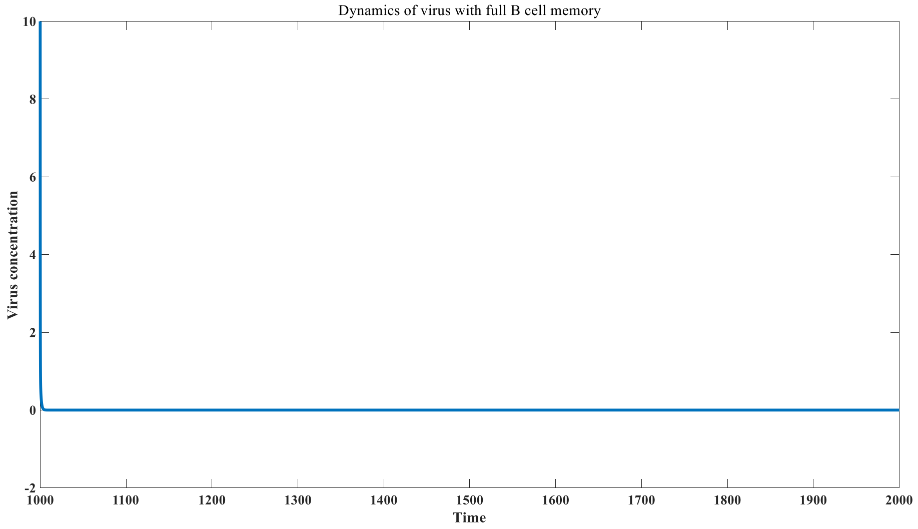


Figure S10_E: effect of B cell memory in preventing secondary infection

### Model 3.4.1 parameter settings and simulation results

Pseudocodes of the agent-based model considering the temporal infection sequence of infected cells are as follows:

```
for i = 1: n

    [V_release_Tc (:, i+1), Tc_dead (i+1), Tc_I_dead (:,i+1)] =
calculate_Tc_effect ( km,Tc ( i ), V_inside (:, i ), I(:, i ), delta_t );

    for j = 1: n

        for mm = 1:10
            for nn = 1:10
```

```
                threshold_temp (10*(mm- 1)+ nn ) =
para_new(mm)*(5*M_2(10*(mm-1)+nn,i)+G_2(10*(mm-
1)+nn,i))*max(0,V_inside(j,i)/I(j,i));
            end
        end
        threshold _sum = sum( threshold_temp );

        if j > i
I( j,i +1) = 0;
            V_inside ( j,i +1) = 0;
            V_release ( j,i +1) = 0;
        else
            if j == i
I( j,i +1) = max(0,k_4*T( i )*V( i )/(V( i )+ k_m )* delta_t );
                V_inside ( j,i +1) = k_4*T( i )*V( i )/(V( i )+ k_m )*
delta_t ;
                V_release ( j,i +1) = 0;
            else

                if ((max( 0,V _inside( j,i )/I( j,i )) < threshold_1) == 1)
&& (( threshold _sum < threshold_2) == 1) && ((I( j,i ) > 0) == 1)

I( j,i +1) = max(0,(I( j,i ) - Tc_I_dead (j,i+1)));
                    V_release ( j,i +1) = V_release_Tc (j,i+1);
                    V_inside ( j,i +1) = k_5* V_inside
( j,i )/I( j,i )*I(j,i+1)* delta_t + V_inside ( j,i )/I( j,i )*I(j,i+1);
                else
I( j,i +1) = 0;
                    V_release ( j,i +1) = V_inside ( j,i );
                    V_inside ( j,i +1) = 0;
                end
            end
        end
    end

T(i+1) = max( 0,( -k_4*T( i )*V( i )/(V( i ) + k_m ) + k_6 - k_7*T( i ))*
delta_t + T( i ));


[M(:,i+1),M_2(:,i+1),G(:,i+1),G_2(:,i+1),E_C_M(:,i+1),E_C_M_2(:,i+1),E_C_G(
:,i+1),E_C_G_2(:,
i+1),V_C_M(:,i+1),V_C_M_2(:,i+1),V_C_G(:,i+1),V_C_G_2(:,i+1),Plasma_M(:,i+1
),Plasma_G(:,i+1), E(i+1),V(i+1)] = ...
```

```
calculate_A_new(para,para_new,para_new_1,M(:,i),M_2(:,i),G(:,i),G_2(:,i),E_
C_M(:,i),E_C_M_2(:,i),E_C_G(:,i),E_C_G_2(:,i),V_C_M(:,i),V_C_M_2(:,i),V_C_G
(:,i),V_C_G_2(:,i), Plasma_M (:, i ), Plasma_G (:, i ),
E( i ),(V( i )+sum( V_release (:, i ))), delta_t , AA);
Tc(i+1) = calculate_ Tc ( Tc_dead (i+1),(V_C_M_2(:, i )+ V_C_G_2(:,
i )),Tc( i ), gen_c,delta_t );

End
```

| Parameter Name | Value |
|---|---|
| km | 2 *10 $^{6}$ |
| k_4 | 1 0 $^{-3}$ |
| k_5 | 5 |
| k_6 | 1 0 $^{7}$ |
| k_7 | 1 0 $^{-3}$ |
| gen_c | 5 *10 $^{-5}$ |
| threshold_1 | 1 0 $^{14}$ |
| threshold_2 | 1 0 $^{9}$ |

Table S17: parameter and initial value in model 3.4.1

When we used the function calculate_A_new to calculate the concentrations of various substances after the time point (delta_t), we employed the model parameters based on 3.1.7. The difference here is that since this process occurs in a non-cellular environment, the viral replication coefficient $k_0$ becomes 0. Additionally, we took into account the additive effect of CD4+ T cells on B cell clonal expansion; therefore, we introduced an additional coefficient $\delta$ (where ( $\delta$ = 1.2)). The system of ordinary differential equations then become:

$$\frac{d(IgM-BCR)_i}{dt} = \pi f(IgM-BCR)_i - p_1 E(IgM-BCR)_i + p_2 C_{2i} + k_3 C_{2i} - k_{1i} V(IgM-BCR)_i + k_{2i} C_{1i} + k_4(1-\gamma)(1-\theta)\frac{C_{1i}}{C_{1i}+(IgM-BCR)_i} C_{1i}{}^{\delta} - k_5(IgM-BCR)_i;$$

$$\frac{d(IgG-BCR)_i}{dt} = -p_1 E(IgG-BCR)_i + p_2 C_{4i} + k_3 C_{4i} - k'_{1i} V(IgG-BCR)_i + k'_{2i} C_{3i} + k_4 \gamma(1-\theta)\frac{C_{1i}}{C_{1i}+(IgM-BCR)_i} C_{1i}{}^{\delta} + k_4(1-\theta)\frac{C_{3i}}{C_{3i}+(IgG-BCR)_i} C_{3i}{}^{\delta} - k_6(IgG-BCR)_i;$$

$$\frac{d(IgM)_i}{dt} = k_7(IgM-ASC)_i - p_1 E(IgM)_i + p_2 C_{6i} - k_{1i} V(IgM)_i + k_{2i} C_{5i} - k_8(IgM)_i;$$

$$\frac{d(IgG)_i}{dt} = k_9(IgG-ASC)_i - p_1 E(IgG)_i + p_2 C_{8i} - k'_{1i} V(IgG)_i + k'_{2i} C_{7i} - k_{10}(IgG)_i;$$

$$\frac{dC_{1i}}{dt} = k_{1i}V(IgM - BCR)_i + k_{2i}C_{1i} - k_{11}C_{1i};$$

$$\frac{dC_{2i}}{dt} = p_1E(IgM - BCR)_i - p_2C_{2i} - k_{11}C_{2i};$$

$$\frac{dC_{3i}}{dt} = k'_{1i}V(IgG - BCR)_i - k'_{2i}C_{3i} - k_{11}C_{3i};$$

$$\frac{dC_{4i}}{dt} = p_1E(IgG - BCR)_i - p_2C_{4i} - k_{11}C_{4i};$$

$$\frac{dC_{5i}}{dt} = k_{1i}V(IgM)_i - k_{2i}C_{5i} - k_{11}C_{5i};$$

$$\frac{dC_{6i}}{dt} = p_1E(IgM)_i - p_2C_{6i} - k_{11}C_{6i};$$

$$\frac{dC_{7i}}{dt} = k'_{1i}V(IgG)_i - k'_{2i}C_{7i} - k_{11}C_{7i};$$

$$\frac{dC_{8i}}{dt} = p_1E(IgG)_i - p_2C_{8i} - k_{11}C_{8i};$$

$$\frac{d(IgM - ASC)_i}{dt} = k_{12}C_{2i} + k_4\epsilon(1-\gamma)\theta\frac{C_{1i}}{C_{1i} + (IgM - BCR)_i}{C_{1i}}^{\delta} - k_{13}(IgM - ASC)_i;$$

$$\frac{d(IgG - ASC)_i}{dt} = k_{12}C_{4i} + k_4\epsilon\theta\frac{C_{3i}}{C_{3i} + (IgG - BCR)_i}{C_{3i}}^{\delta} - k_{14}(IgG - ASC)_i;$$

$$\frac{dV}{dt} = k_0V - \sum_n^{i=1} k_{1i}V(IgM - BCR)_i + \sum_n^{i=1} k_{2i}C_{1i} - \sum_n^{i=1} k'_{1i}V(IgG - BCR)_i + \sum_n^{i=1} k'_{2i}C_{3i} - \sum_n^{i=1} k_{1i}V(IgM)_i + \sum_n^{i=1} k_{2i}C_{5i} - \sum_n^{i=1} k'_{1i}V(IgG)_i + \sum_n^{i=1} k'_{2i}C_{7i};$$

$$\frac{dE}{dt} = \pi_1 - \sum_n^{i=1} p_1E(IgM - BCR)_i + \sum_n^{i=1} p_2C_{2i} - \sum_n^{i=1} p_1E(IgG - BCR)_i + \sum_n^{i=1} p_2C_{4i} - \sum_n^{i=1} p_1E(IgM)_i + \sum_n^{i=1} p_2C_{6i} - \sum_n^{i=1} p_1E(IgG)_i + \sum_n^{i=1} p_2C_{8i};$$

We used this model to first study the impact of individual immune strength on the dynamics of virus-host interactions. The antigen-BCR complex's regeneration coefficient $k_4$ is an important indicator reflecting the strength of individual immunity. We found that when $k_4$ is reduced, it becomes easier to trigger the occurrence of chronic infections. We investigated the infection characteristics under two different scenarios: $k_4$ = 2 and $k_4$ = 3. When $k_4$ = 2, chronic infection occurred, as shown in Figure S11. However, when the immune response was strong, chronic infection did not occur, as illustrated in Figure S12.

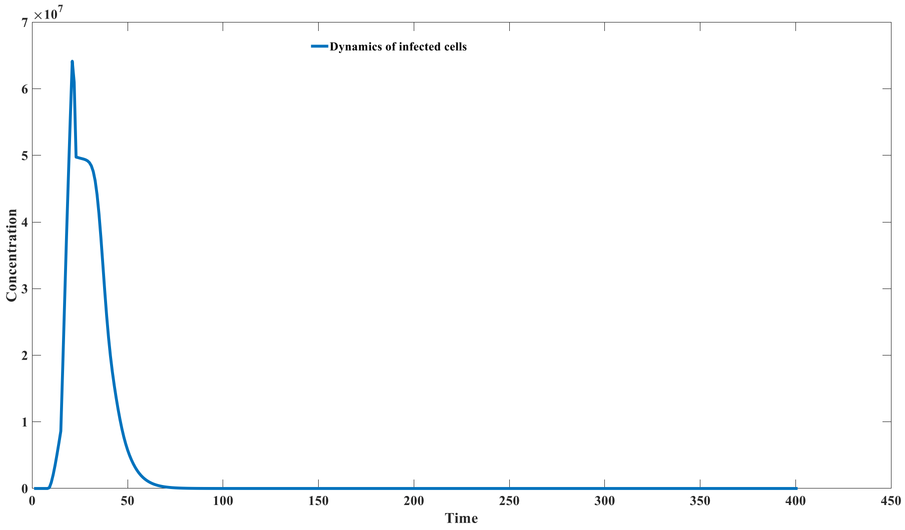


Figure S11_A: Dynamics of infected cells ( $k_4$= 3 )

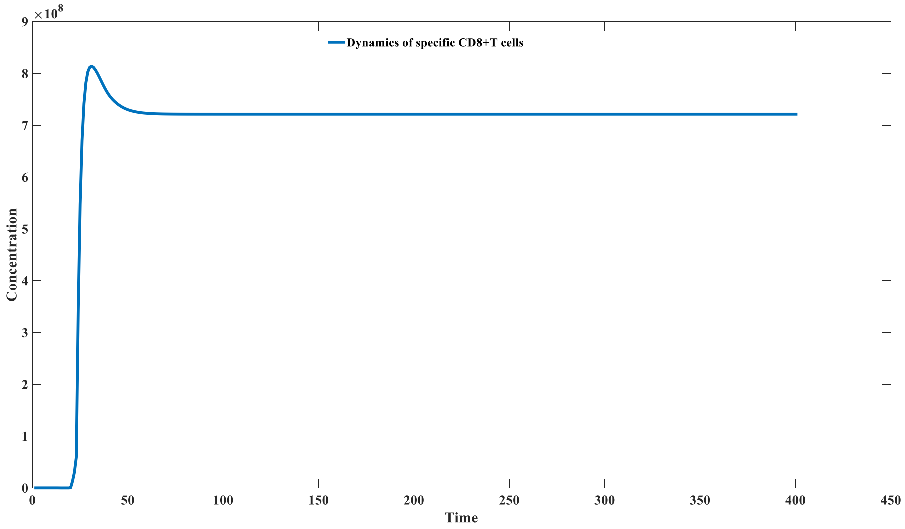


Figure S11_B: Dynamics of specific CD8+T cells ( $k_4$= 3 )

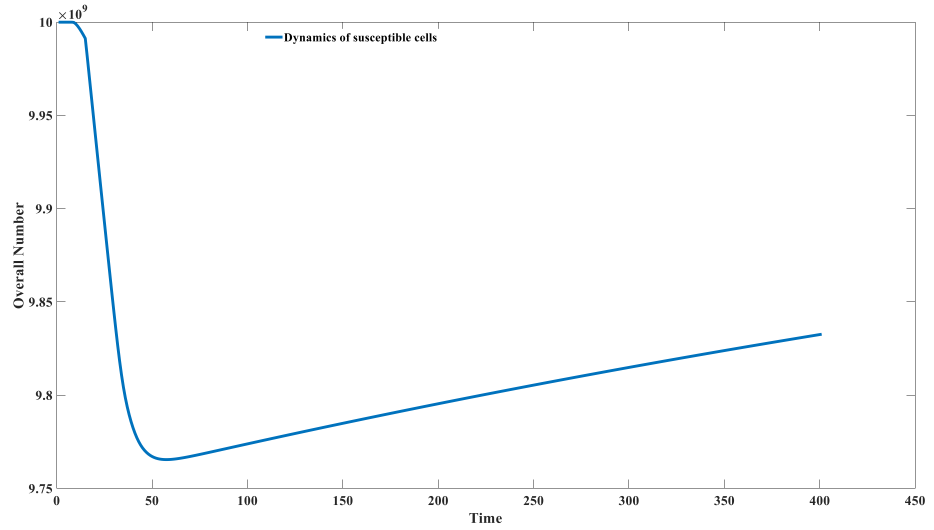


Figure S11_C: Dynamics of susceptible cells ( $k_4$= 3 )

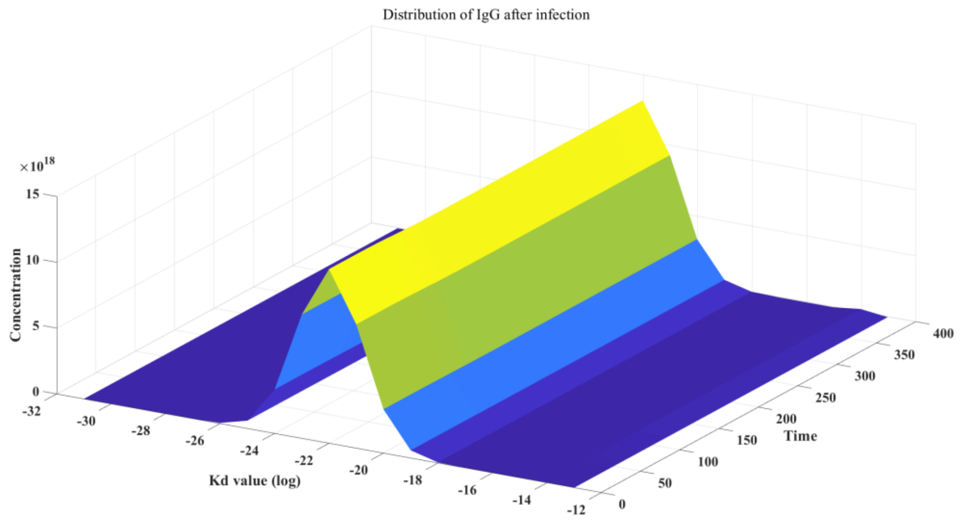


Figure S11_D: Distribution of IgG after infection ( $k_4$= 3 )

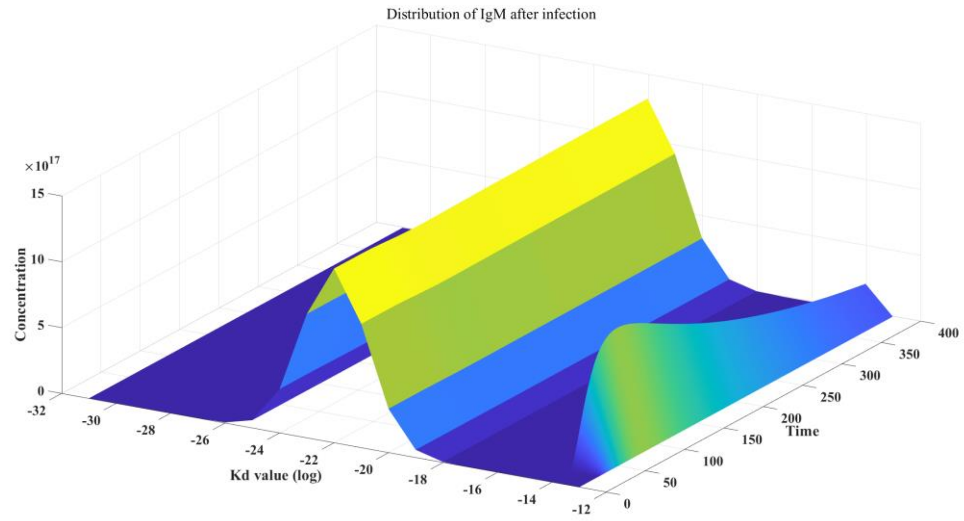


Figure S11_E: Distribution of IgM after infection ($k_4$=3)

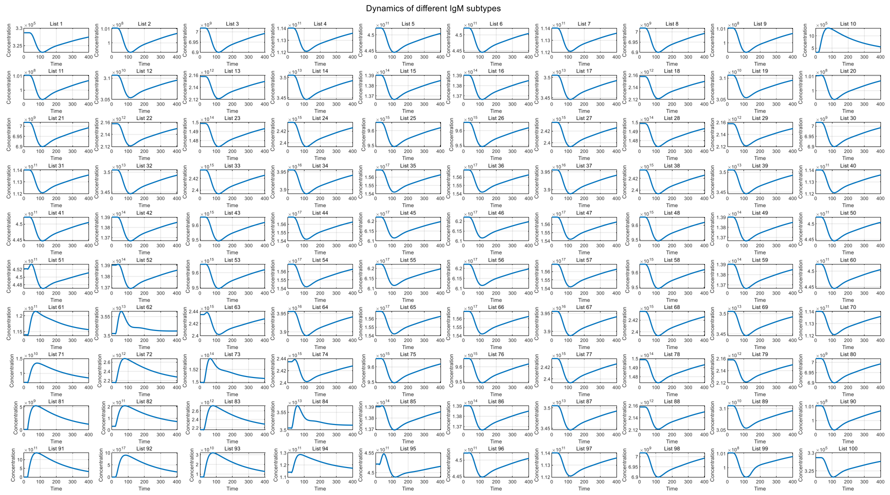


Figure S11_F: Dynamics of different IgM subtypes ($k_4$=3)

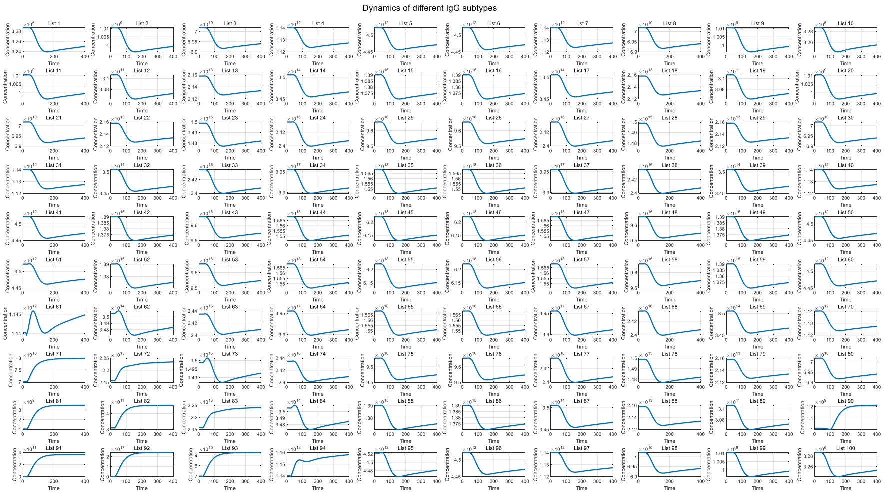


Figure S11_G: Dynamics of different IgG subtypes ($k_4$=3)

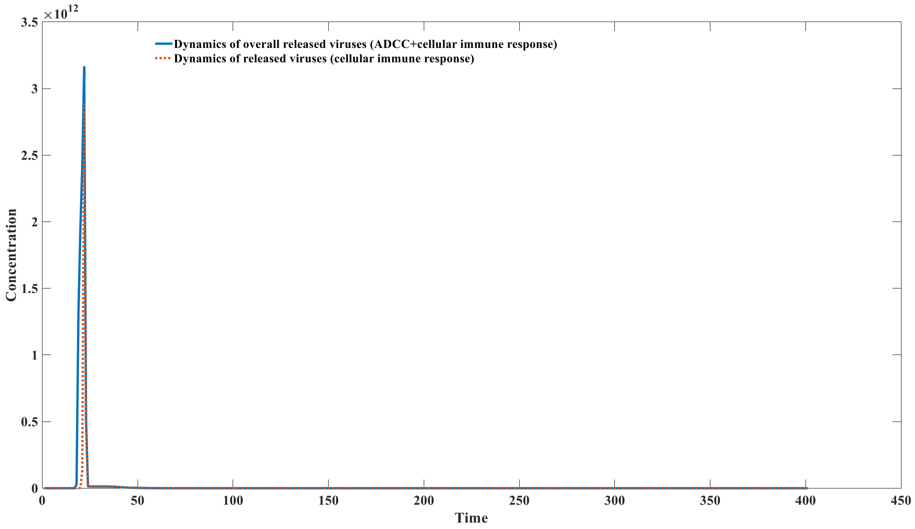


Figure S11_H: Dynamics released viruses by ADCC and cellular immune response ( $k_4$= 3 )

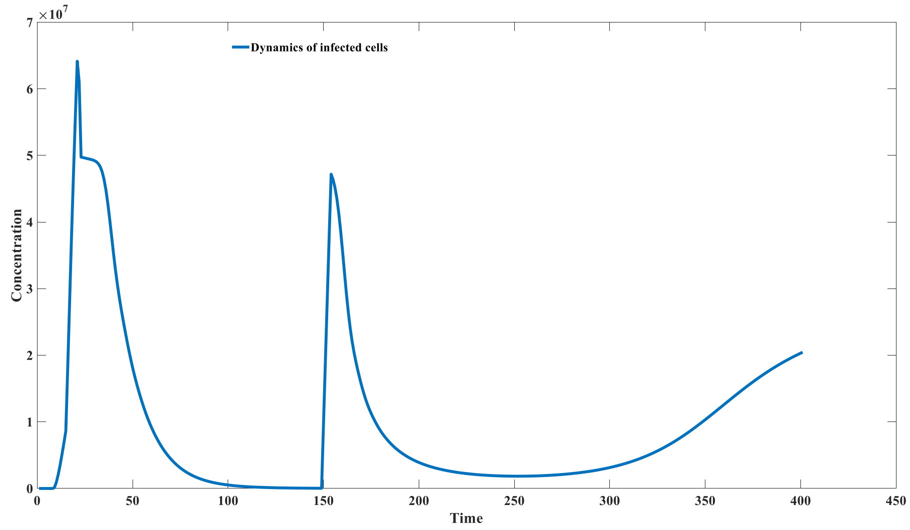


Figure S12_A: Dynamics of infected cells ( $k_4$= 2 )

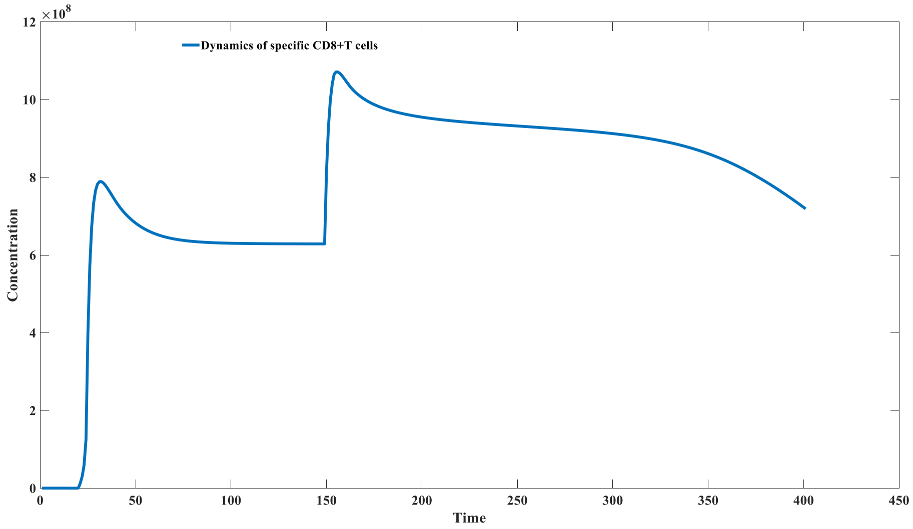


Figure S12_B: Dynamics of specific CD8+T cells ( $k_4$= 2 )

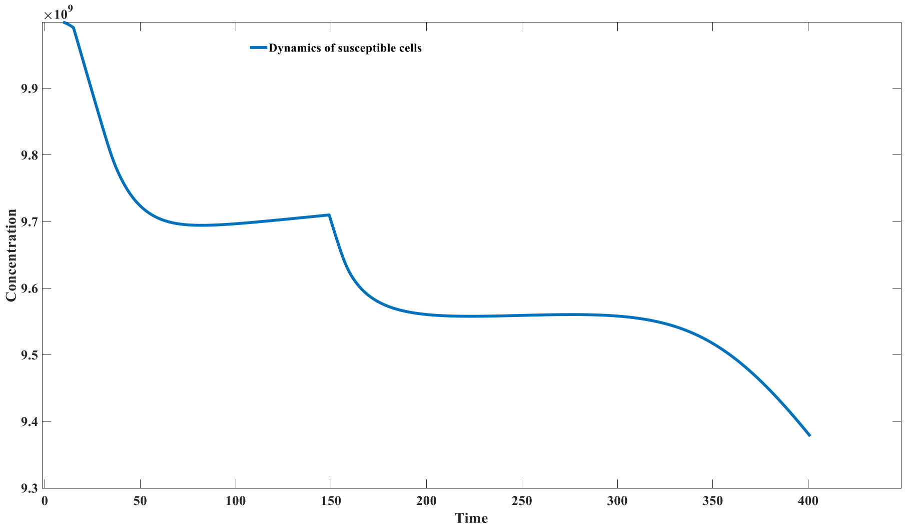


Figure S12_C: Dynamics of susceptible cells ( $k_4$= 2 )

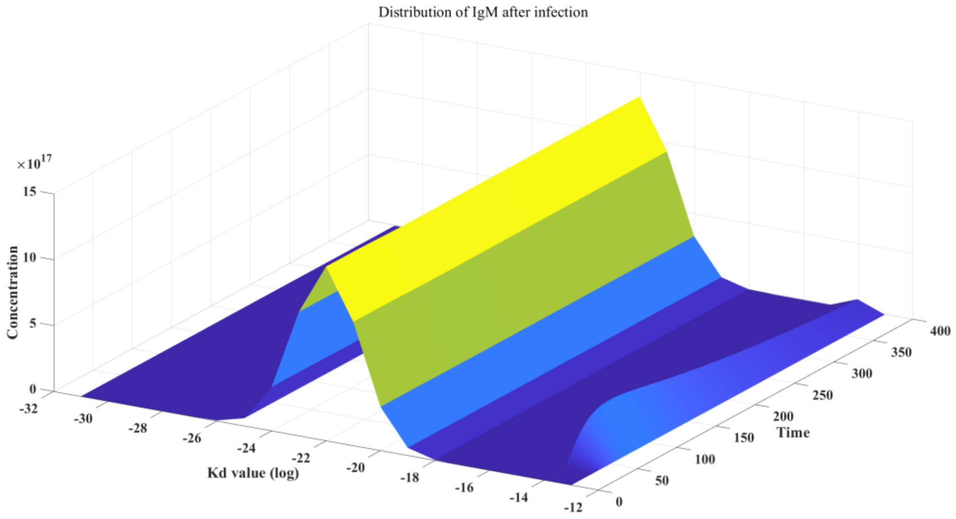


Figure S12_D: Distribution of IgM after infection ( $k_4$= 2 )

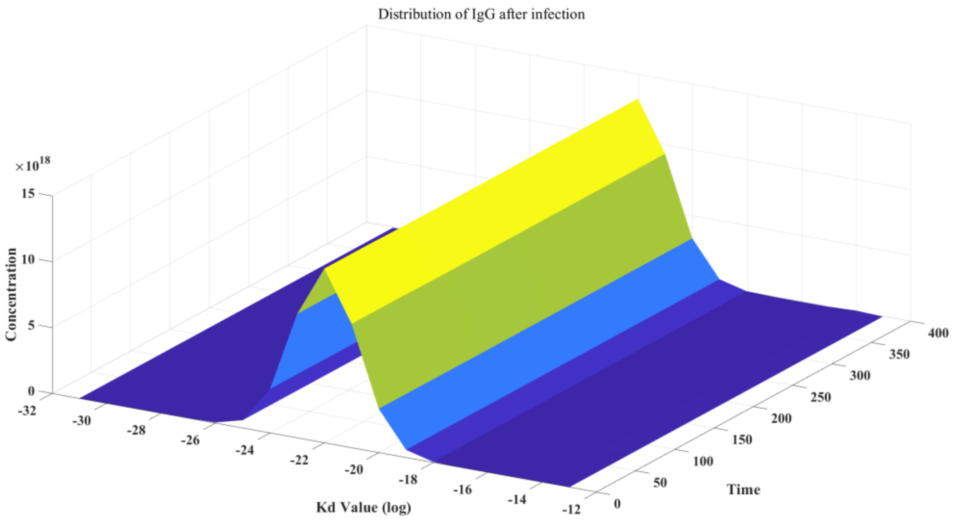


Figure S12_E: Distribution of IgG after infection ($k_4$=2)

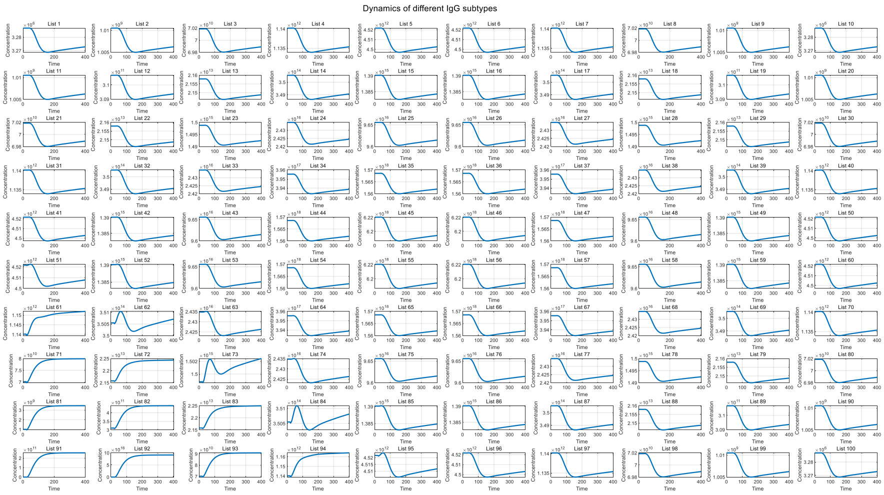


Figure S12_F: Dynamics of different IgG subtypes after infection ($k_4$=2)

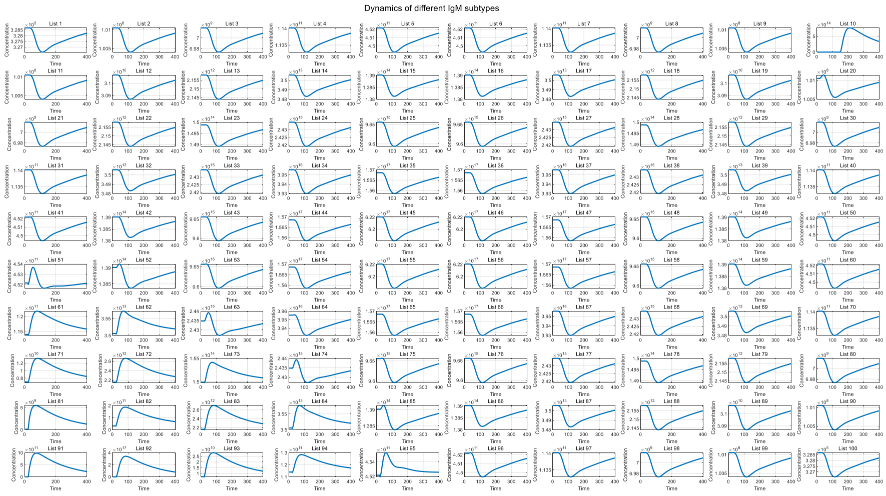


Figure S12_G: Dynamics of different IgM subtypes after infection ($k_4$=2)

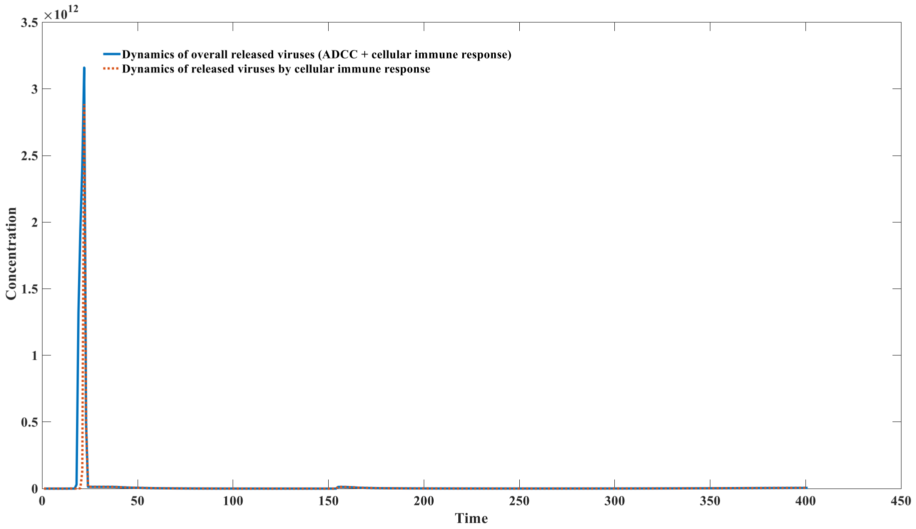


Figure S12_H: Dynamics released viruses by ADCC and cellular immune response ( $k_4$= 2)

We further investigated the impact of the viral replication coefficient $k_0$ on chronic infections. We found that a smaller viral replication coefficient makes it easier to trigger the occurrence of chronic infections. The dynamics of infected cells when the viral replication coefficient $k_0 = 1$ are shown in Figure S13. In contrast, the dynamics of infected cells when the viral replication coefficient $k_0 = 5$ are illustrated in Figure S11_A.

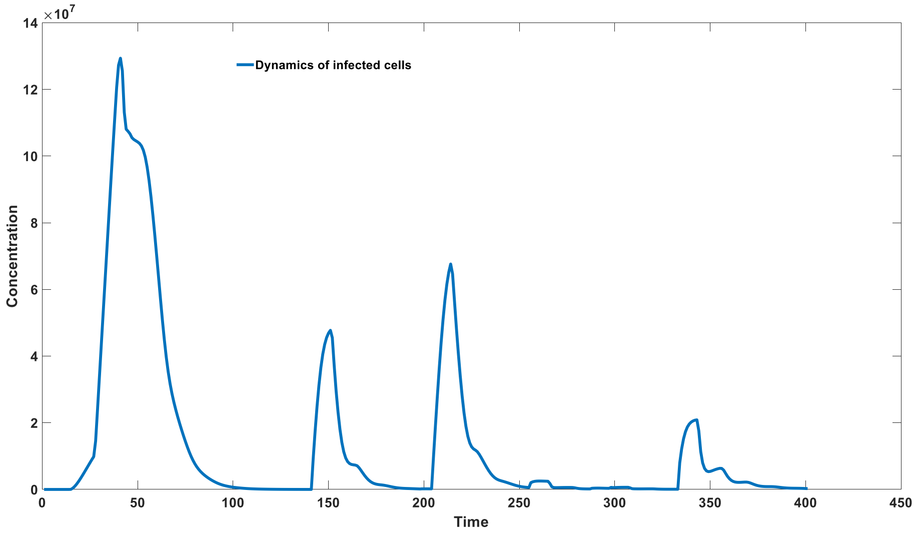


Figure S13: Slow replication virus tends to cause chronic infection ($k_0$ = 1)

Due to the presence of immune imprinting, secondary infections caused by mutant strains may also trigger the occurrence of chronic infections or prolong the infection cycle. We investigated the impact of infections from mutant strains with different values of ( \alpha ) on the dynamics of infected cells, and the results are shown in Figure S14. From S14, it can be seen that as the mutation coefficient increases, the likelihood of chronic infections may rise; however, with a further increase in the mutation coefficient, the risk of chronic infections decreases. This is because when the mutation coefficient is very low, the antibodies formed during the primary infection are sufficient to clear the virus. However, as the mutation coefficient increases, the antibodies generated from the initial infection become inadequate to cope with the occurrence of secondary infections. At the same time, due to the initially high concentration of antibodies, there is effective suppression of viral proliferation in the early stages of infection, which hinders the rapid regeneration of antibodies. This creates a platform for an endemic equilibrium phase, making it easier for chronic infections to occur.

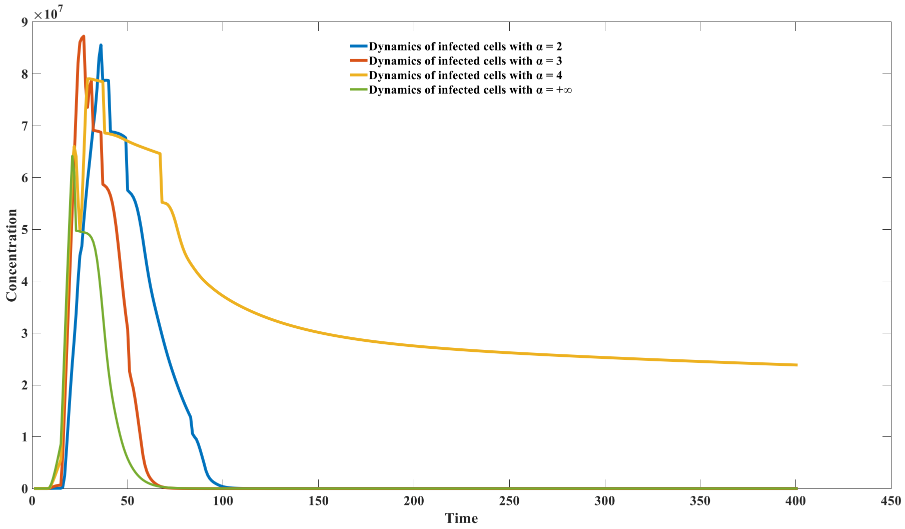

Figure S14: Secondary Infection tends to be chronic

We further investigated three different methods for treating chronic infections: monoclonal antibody therapy, self-antigen substance therapy, and therapeutic vaccine therapy.

For monoclonal antibody therapy, the forward affinity coefficient of the antibodies we added was $10^{-13}$, and the reverse dissociation coefficient was 1 (these properties are identical to those of antibody 91). We administered a relatively large dose of monoclonal antibodies at the 200th time point (specifically, $2 \times 10^{17}$ and $5 \times 10^{17}$), and the results are shown in Figure S15.

From Figure S15, it can be observed that the addition of monoclonal antibodies can effectively kill infected cells in a short period. However, this effect is not durable within this dosage range; as the monoclonal antibodies are consumed, chronic infection re-emerges.

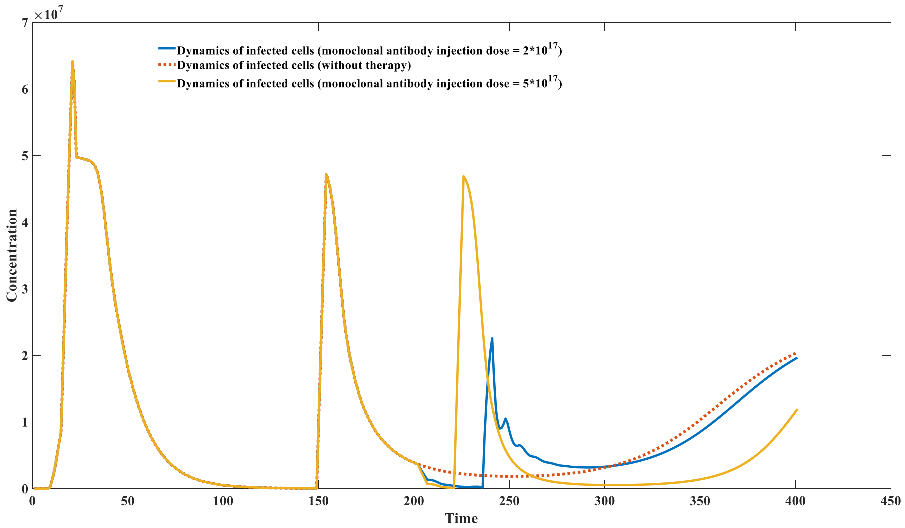


Figure S15: Effects of monoclonal antibody therapy on chronic infection

For self-antigen substance therapy, we continuously added self-antigen substances between the 200th and 250th time points at a concentration of $5 \times 10^{17}$. The results are shown in Figure S16.

From Figure S16, it can be seen that the continuous addition of a large dose of self-antigen substances can help alleviate chronic infections; however, this effect is also not durable.

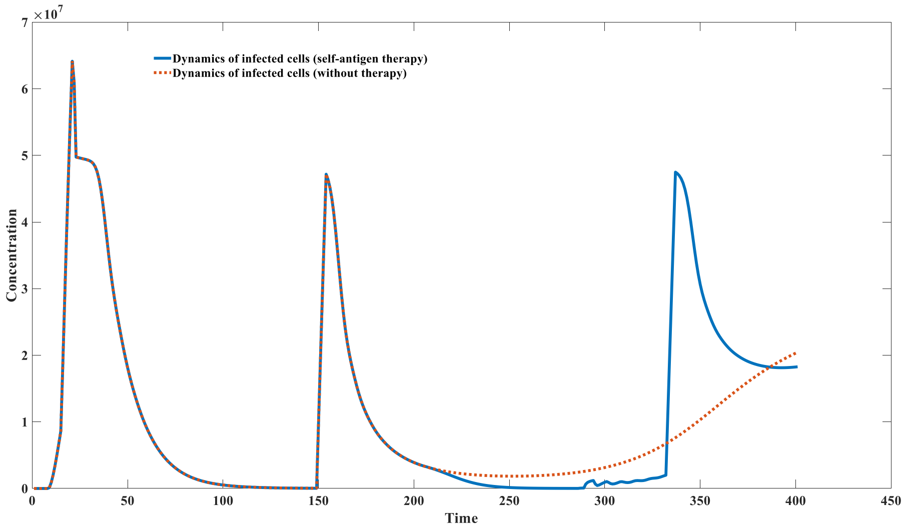


Figure S16: Effects of self - antigen therapy on chronic infection

Finally, we simulated the therapeutic effect of vaccine therapy on chronic infections. We administered viral antigens at doses of $5 * 10^{14}$ and $8 * 10^{14}$ at the 200th time point, and the simulation results are shown in Figure S17. From the figure, it can be observed that when a sufficient dose of the therapeutic vaccine is added, chronic infections can be fundamentally cleared, as indicated by the red solid line in the figure.

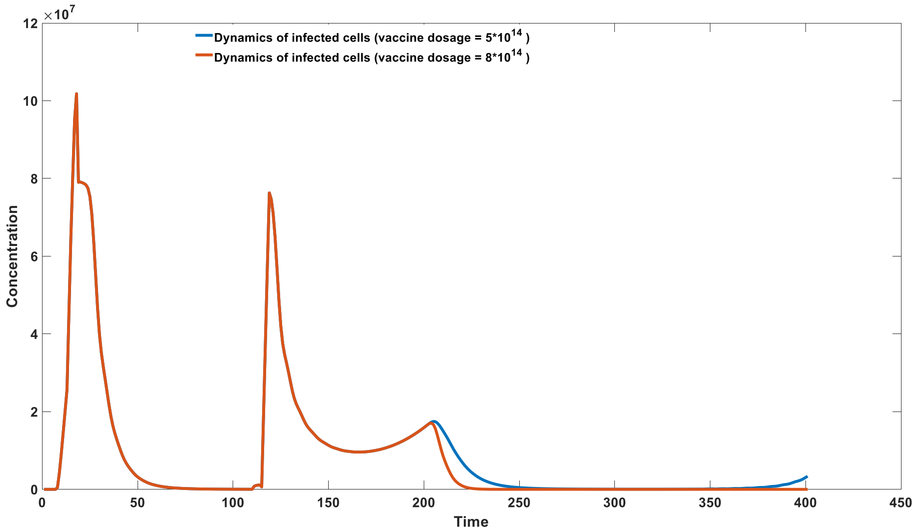


Figure S17: Effects of therapeutic vaccine on chronic infection

We further investigated the dosage effect of therapeutic and preventive vaccines. To overcome chronic infections, the vaccination dose of therapeutic vaccines needs to be significantly higher than that of preventive vaccines. We used the same low dose of vaccine (both at $10^{13}$); one served as a preventive vaccine, administered at time point 0, followed by infection with the actual virus at time point 200. The dynamics of infected cells are shown in Figure S18. From the figure, it can be observed that the low-dose preventive vaccine effectively prevents infection.

We then applied the same low dose of vaccine for therapeutic purposes, where we infected with the actual virus at time point 0 and administered the vaccine at time point 200. The dynamics of infected cells are shown in Figure S19. It is clear from the figure that the same dose of therapeutic vaccine is insufficient to eradicate chronic infections. To achieve complete elimination of chronic infections, the dose of therapeutic vaccines generally needs to be significantly higher than the dose used for preventive vaccines.

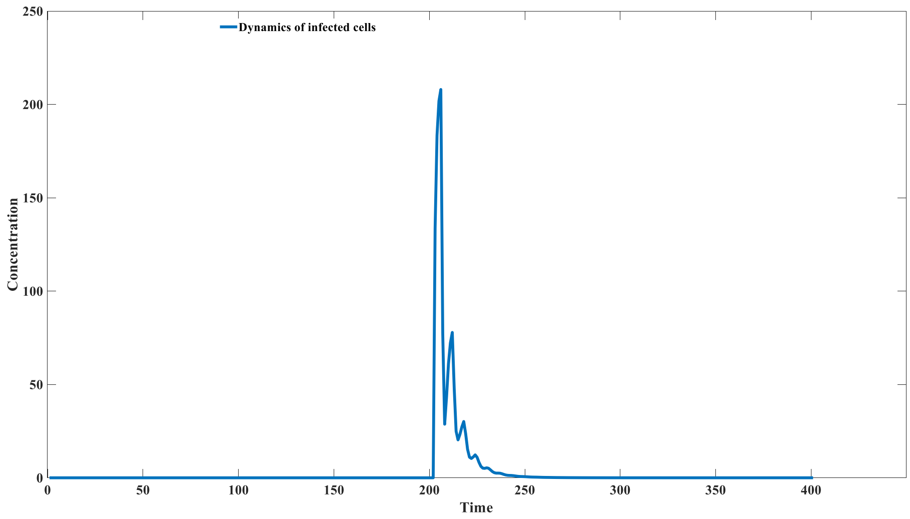


Figure S18: Effects of prophylactic vaccine in preventing Infection

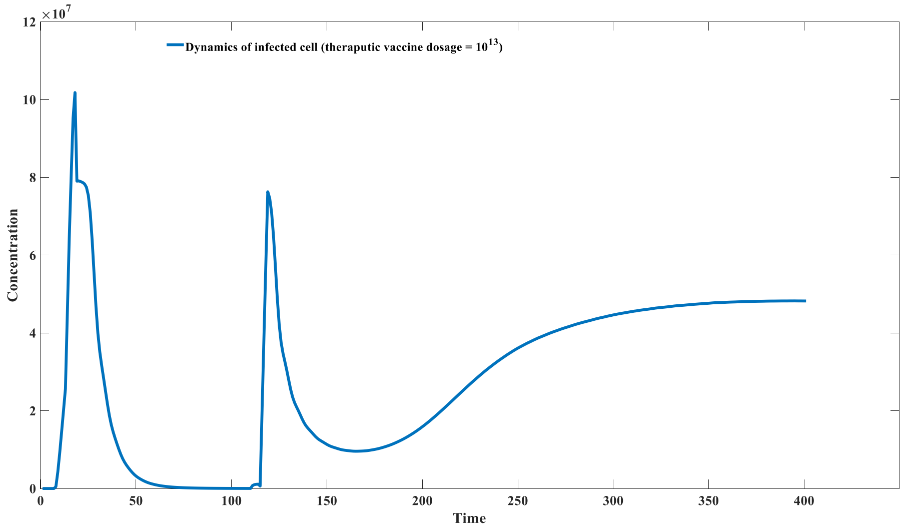


Figure S19: Insufficient therapeutic vaccine dosage cannot cure chronic Infection

**Mathematical analysis and simulation results of Model 3.5.1**

Model 3.5.1 is as follows:

$$\frac{dT_u}{dt} = \pi_1 + k_1 T_u - d_1 T_u - \beta\frac{A}{A + k_m} T_u - \alpha T_c T_u;$$

$$\frac{d(Neo)}{dt} = -k_2(Neo)A + k_3 C + \rho_1\left(d_1 T_u + \beta\frac{A}{A + k_m} T_u + \alpha T_c T_u\right);$$

$$\frac{dC}{dt} = k_2(Neo)A - k_3 C - k_4 C;$$

$$\frac{dA}{dt} = -k_2(Neo)A + k_3 C + k_5 C + \pi_2 - d_2 A;$$

$$\frac{dT_c}{dt} = -\rho_2 \alpha T_c T_u + k_6 C + \pi_3 - d_3 T_c;$$

The initial values and parameter selections are as follows:

| | |
|---|---|
| $T_{u0}$ | 0 |
| $A_0$ | $10^{5}$ |
| $C_0$ | 0 |
| $Neo_0$ | 0 |
| $T_{c0}$ | $10^{3}$ |
| $k_1$ | 1 |
| $k_2$ | 10-7 |
| $k_3$ | 0 |
| $k_4$ | 1 |
| $k_5$ | 2 |
| $k_6$ | $2*10^{-4}$ |
| $\pi_1$ | 100 |
| $\pi_2$ | 100 |
| $\pi_3$ | 0.1 |
| $d_1$ | 0.05 |
| $d_2$ | 0.001 |
| $d_3$ | 0.001 |
| $\rho_1$ | $10^{4}$ |
| $\rho_2$ | 0.1 |
| $\alpha$ | $10^{-4}$ |
| $\beta$ | 10 |
| $k_m$ | $10^{7}$ |

Table S18: parameter and initial value in model 3.5.1

There exists a balance point of an endemic equilibrium state ($T_c^* = 200966.589$; $T_u^* = 3.4420904$; $Neo^* = 9999.033503$; $C^* = 1034420.904$; $A^* = 1034520904.481$). Due to the presence of many parameters, our parameter selection process is based on the reverse deduction of the equilibrium point. For instance, when using the above parameters, the resulting equilibrium point aligns well with the equilibrium state observed in actual biological processes. Therefore, we consider these parameter combinations to be relatively reasonable.

Stability analysis of equilibrium point:

The Jacobian matrix of the equilibrium point is as follows:

$$\begin{matrix} k_1 - d_1 - \beta\frac{A}{A+k_m} - \alpha T_c & 0 & 0 & \frac{-\beta k_m T_u}{(A+k_m)^2} & -\alpha T_u \\ \rho_1(d_1 + \beta\frac{A}{A+k_m} + \alpha T_c) & -k_2 A & k_3 & -k_2(\mathrm{Neo}) + \frac{\rho_1 \beta k_m T_u}{(A+k_m)^2} & \rho_1 \alpha T_u \\ 0 & k_2 A & -k_3 - k_4 & k_2(\mathrm{Neo}) & 0 \\ 0 & -k_2 A & k_3 + k_5 & -k_2(\mathrm{Neo}) - d_2 & 0 \\ -\rho_2 \alpha T_c & 0 & 0 & k_6 & -\rho_2 \alpha T_u - d_3 \end{matrix}$$

The eigenvalue of the equilibrium point is ( $\lambda_1 = -103.452; \lambda_2 = -8.954; \lambda_3 = -0.998;$ $\lambda_4 = -0.0026; \lambda_5 = -0.0020;$), so the equilibrium point is stable.

**Numerical Simulation:** The initial value selection for numerical simulation involves choosing a disease-free state, where it is assumed that no cancer cells are present. The initial values are listed as shown. The results of the numerical simulation are presented in Figure S20. From Figure S20, it can be observed that the system will ultimately reach an equilibrium state; however, in the early stages, cancer cells will undergo a significant proliferation phase. Therefore, the outbreak of cancer does not depend on the stability of the equilibrium state.

In the majority of cases, with the accumulation of mutation coefficients and the aging of the immune system, the growth coefficient of cancer cells $k_1$ will continuously increase, while the coefficients $k_5$ and $k_6$, which represent immunity strength, will continuously decrease. At this point, when cells undergo carcinogenesis, cancer cells will experience a substantial and significant proliferation process, leading to the manifestation of typical symptoms of cancer in the body, indicating the occurrence of clinical cancer. However, the system can ultimately reach a new equilibrium state, which implies that such cancers can theoretically be completely cured.

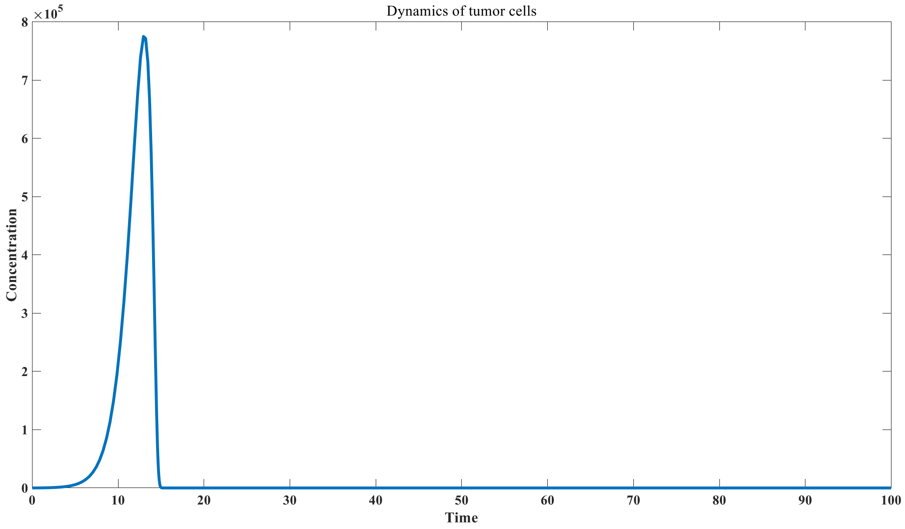


Figure S20: Dynamics of tumor cells when $k_1$ = 1.0

For example, when the cancer cell proliferation coefficient $k_1$ increases to 1.1, the numerical simulation results are shown in Figure S21. Compared with S20, the cancer cells have been significantly expanded in the early stage. At this time, the new equilibrium point is ($T_c{}^* = 201632.956$; $T_u{}^* = 3.446151$; $Neo^* = 9999.0366$; $C^* = 1037907.667$; $A^* = 1038007667.074$), and the eigenvalue of the equilibrium point is ( $\lambda_1 = -103.800; \lambda_2 = -8.855; \lambda_3 = -0.998;$ $\lambda_4 = -0.0027; \lambda_5 = -0.0020;$). At this time, the equilibrium point is still stable.

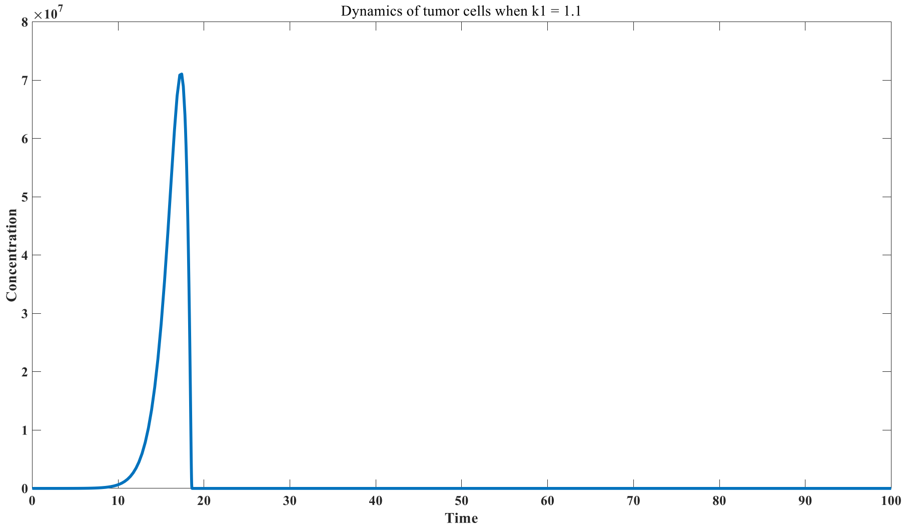


Figure S21: D ynamics of tumor cells when $k_1$= 1.1

When the immunity coefficient $k_5$ drops to 1.9, the numerical simulation results are shown in Figure S22. Compared to S20, the peak concentration of cancer cells is significantly increased, which causes the individual to exhibit typical characteristics of cancer. The new equilibrium point is ( $T_c{}^* = 200966.711$; $T_u{}^* = 3.443$; $Neo^* = 11109.917$; $C^* = 1034434.773$; $A^* = 931091296.350$), and the eigenvalue of the equilibrium point is ( $\lambda_1 = -93.109; \lambda_2 = -8.944; \lambda_3 = -0.998;$ $\lambda_4 = -0.0026; \lambda_5 = -0.0021$). The system remains stable at the equilibrium point.

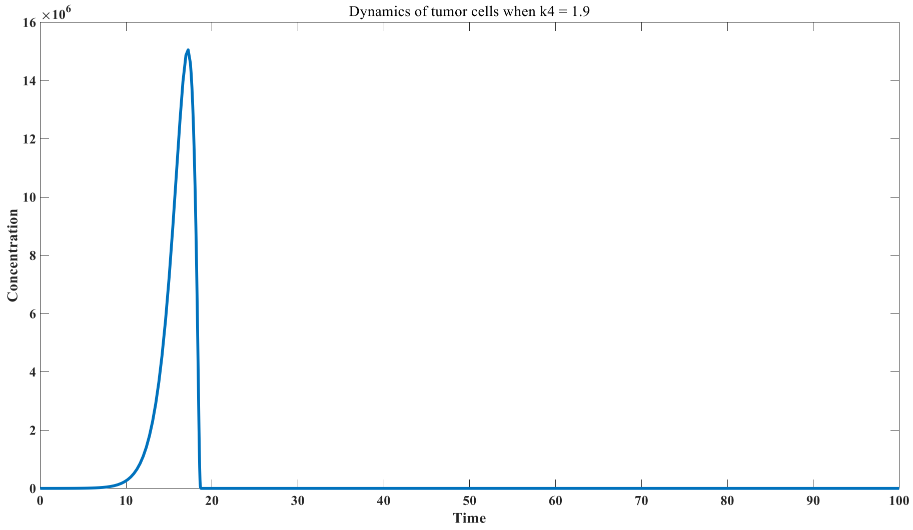


Figure S22: D ynamics of tumor cells when $\mathrm{k}_5$= 1.9

When the immunity coefficient $\mathrm{k}_5$ continues to decrease, for instance, when it drops to 1.8, the proliferation of cancer cells becomes even more pronounced, as shown in Figure S23. At this point, the system exhibits a new equilibrium point ( $\mathrm{T_c}^* = 200966.6969$ ; $\mathrm{T_u}^* = 3.4450340$; $\mathrm{Neo}^* = 12498.48971849$; $\mathrm{C}^* = 1034450.340189$; $\mathrm{A}^* = 827660272.1514$ ), with eigenvalues ( $\lambda_1 = -82.7660; \lambda_2 = -8.9310; \lambda_3 = -0.9980$; $\lambda_4 = -0.0027; \lambda_5 = -0.0022$). At this time, the equilibrium point remains stable; however, the proliferation of cancer cells in the actual simulation is remarkably striking.

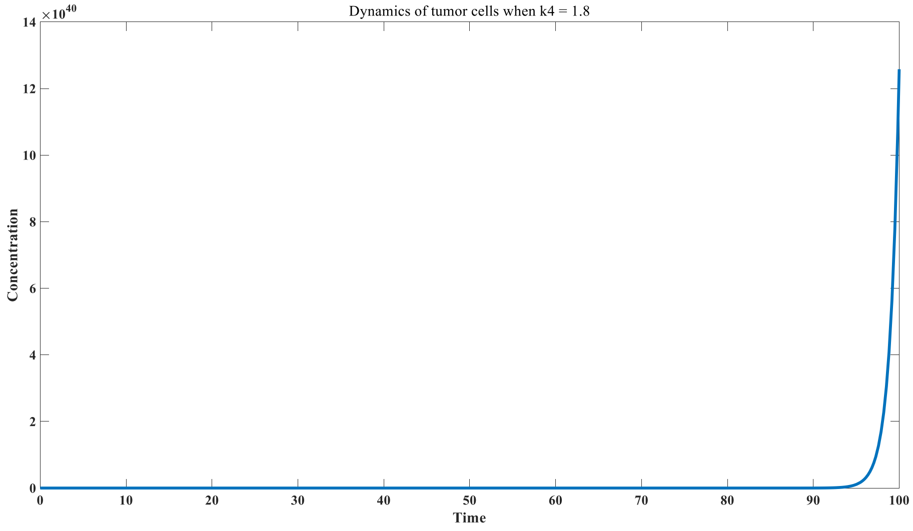


Figure S23: D ynamics of tumor cells when $\mathrm{k}_5$= 1.8

We further simulated the inhibitory effects of several common cancer immunotherapy regimens on cancer cell proliferation, where the immune checkpoint inhibitor therapy corresponds to changes in $\rho_2$ within the system. The specific simulation results are shown in

Figure S24. From Figure S24, it can be observed that reducing $\rho_2$ can significantly control the extent of cancer cell proliferation.

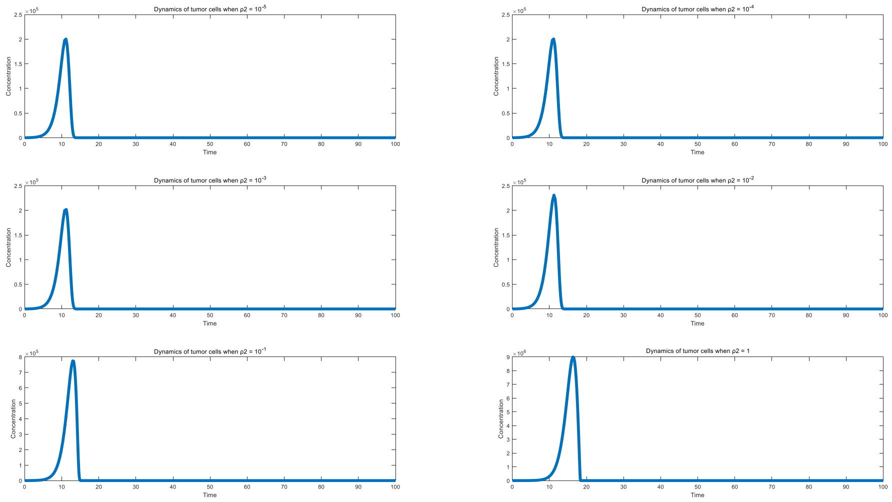

Figure S24: Mechanism of PD-1 inhibitor on tumor control

We further simulated the impact of the addition of monoclonal antibodies on cancer cell dynamics. We extended Model 3.5.1 by incorporating a set of ordinary differential equations for the monoclonal antibody component, as follows:

$$\frac{dT_u}{dt} = \pi_1 + k_1T_u - d_1T_u - \beta\frac{A + A'}{A + A' + k_m}T_u - \alpha T_cT_u;$$

$$\frac{d(Neo)}{dt} = -k_2(Neo)A + k_3C + \rho_1(d_1T_u + \beta\frac{A + A'}{A + A' + k_m}T_u + \alpha T_cT_u);$$

$$\frac{dC}{dt} = k_2(Neo)A - k_3C - k_4C;$$

$$\frac{dA}{dt} = -k_2(Neo)A + k_3C + k_5C + \pi_2 - d_2A;$$

$$\frac{dT_c}{dt} = -\rho_2\alpha T_cT_u + k_6C + \pi_3 - d_3T_c;$$

$$\frac{dA'}{dt} = -k_2(Neo)A' + k_3C' - d_4A';$$

The parameters and initial values used are as follows:

| | |
|---|---|
| $T_{u0}$ | 0 |
| $A_0$ | $10^5$ |
| $C_0$ | 0 |
| $Neo_0$ | 0 |
| $T_{c0}$ | $10^3$ |
| $A'_0$ | 0 |
| $C'_0$ | 0 |
| $k_1$ | 1.1 |

| | |
|---|---|
| $k_2$ | 10-7 |
| $k_3$ | 0 |
| $k_4$ | 1 |
| $k_5$ | 2 |
| $k_6$ | $2*10^{-4}$ |
| $\pi_1$ | 100 |
| $\pi_2$ | 100 |
| $\pi_3$ | 0.1 |
| $d_1$ | 0.05 |
| $d_2$ | 0.001 |
| $d_3$ | 0.001 |
| $d_4$ | 0.05 |
| $\rho_1$ | $10^{4}$ |
| $\rho_2$ | 0.1 |
| $\alpha$ | $10^{-4}$ |
| $\beta$ | 10 |
| $k_m$ | $10^{7}$ |

Table S19: parameter and initial value in monoclonal antibody therapy model

We introduced different doses of monoclonal antibody $A'$ at the 10th time point. The simulation results are shown in Figure S25. From Figure S25, it can be observed that the addition of monoclonal antibodies targeting neoantigens effectively controls cancer cell proliferation, with the inhibitory effect showing a positive correlation with the added dose.

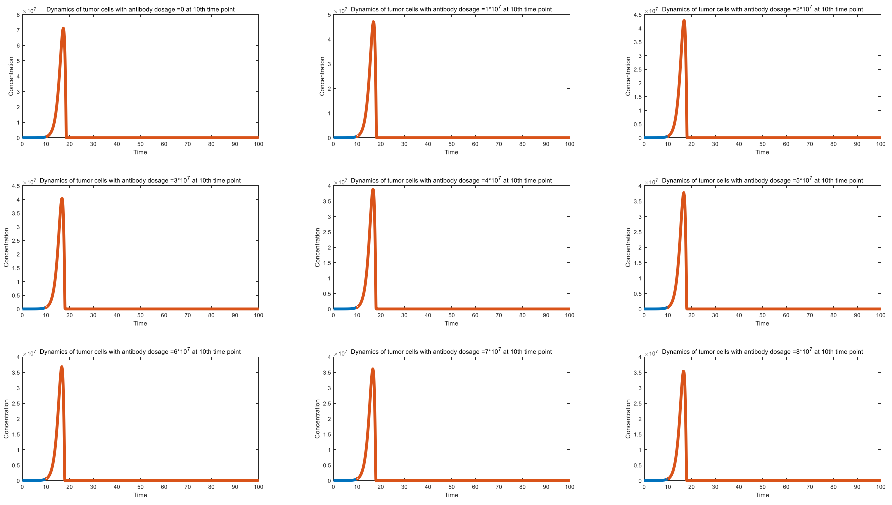


Figure S25: Effect of monoclonal antibody on tumor control

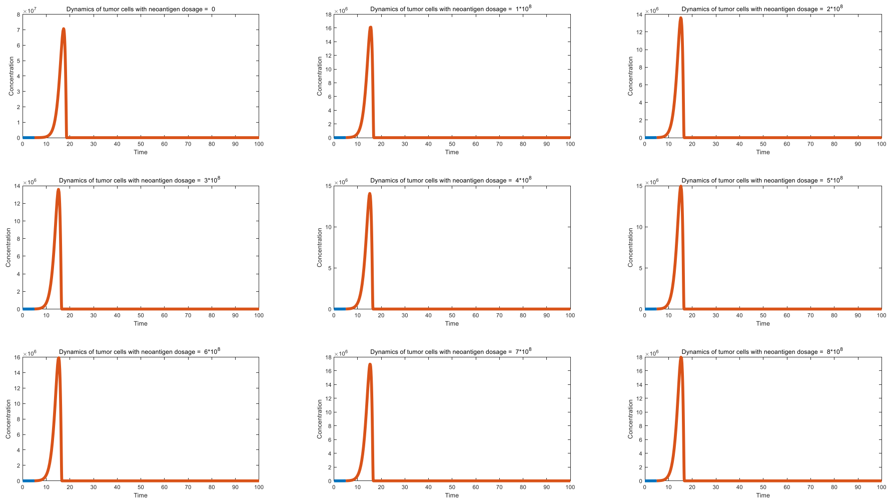


Figure S26_A: Effect of neoantigen addition on tumor control（vaccination at 5 th time point)

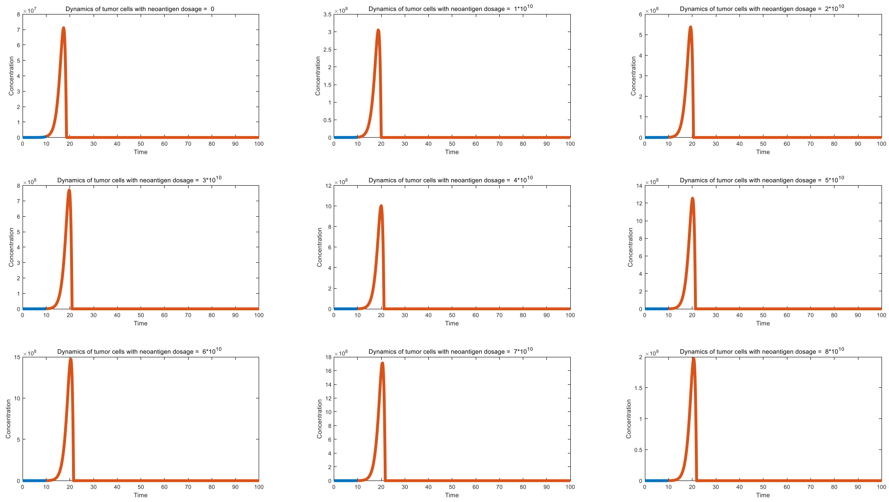


Figure S26_B: Effect of neoantigen addition on tumor control（vaccination at 10 th time point)

We further simulated the impact of the addition of neoantigens on tumor cell dynamics. The timing of neoantigen administration is crucial; we found that early administration of neoantigens can control the proliferation of tumor cells, while late administration can promote tumor cell proliferation.

We introduced different doses of neoantigens at the 5th time point, and the results are shown in Figure S26_A. From the figure, it can be seen that earlier administration of a lower dose of neoantigen has a certain controlling effect on cancer cells.

We also introduced different doses of neoantigens at the 10th time point, with results presented in Figure S26_B. From this figure, it can be observed that later use of neoantigens stimulates tumor proliferation. This is because the added neoantigens can lead to the depletion of antibodies.

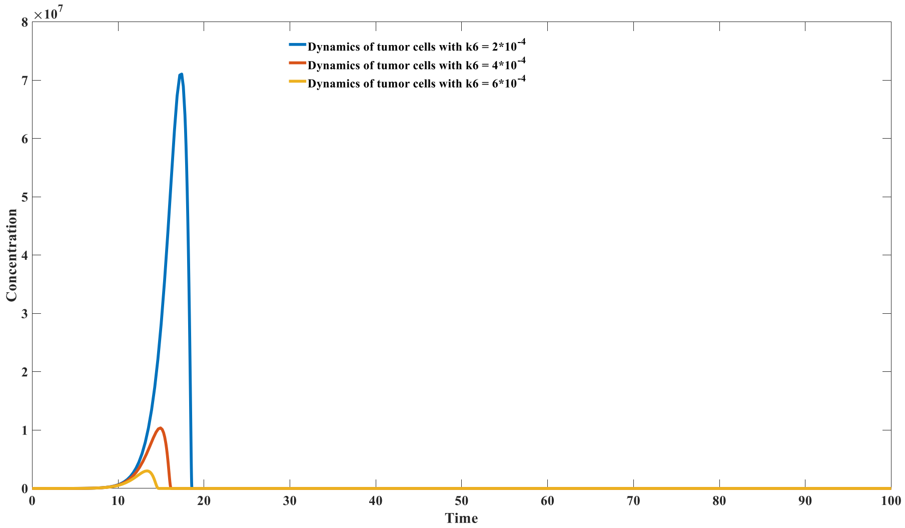


Figure S27: Mechanism of MHC-1 neoantigen vaccine on tumor control

We further investigated the control effect of MHC-I class antigens on cancer cell proliferation. Unlike neoantigens that bind to B cells, MHC-I class antigens do not bind to antibodies and therefore do not lead to antibody depletion. Their mechanism of action lies in promoting the proliferation of CD8+ T cells, which effectively increases the value of $k_6$.
We simulated the dynamics of cancer cells under different proliferation capacities of CD8+ T cells, as shown in Figure S27. From the figure, it can be observed that with the enhancement of CD8+ T cell activation capacity, the early proliferation of cancer cells can be better controlled.

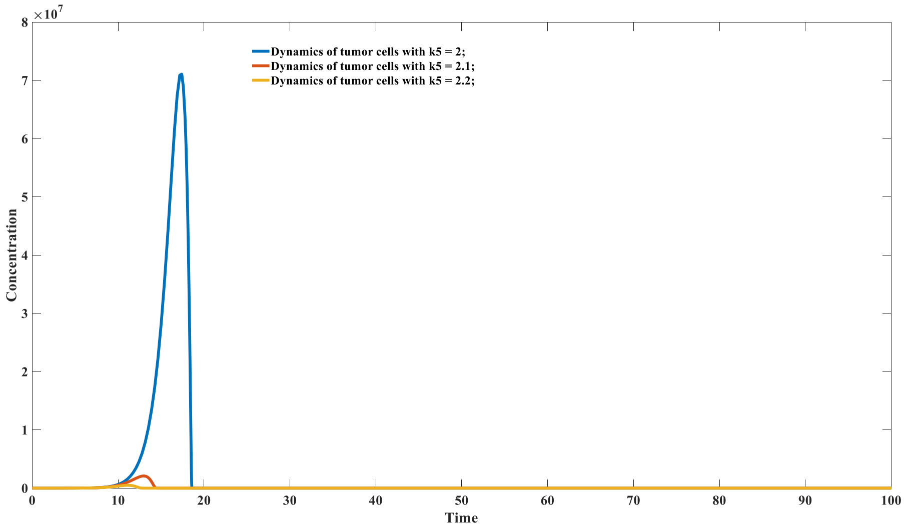


Figure S28: Mechanism of MHC- II neoantigen vaccine on tumor control

Finally, we studied the control effect of MHC-II class antigens on cancer cell proliferation. Similar to MHC-I class antigens, MHC-II class antigens do not bind to antibodies and therefore do not lead to antibody depletion. Their mechanism of action lies in promoting the proliferation of antibodies, which effectively increases the value of $k_5$.

We simulated the dynamics of cancer cells under different antibody proliferation capacities, as shown in Figure S28. From the figure, it can be observed that with the enhancement of antibody proliferation capacity, the early proliferation of cancer cells can be better controlled.
